# Discovery and biosynthesis of imidazolium antibiotics from a probiotic *Bacillus licheniformis*

**DOI:** 10.1101/2022.10.05.511033

**Authors:** Song Lim Ham, Tae Hyun Lee, Kyung Jun Kim, Jung Ha Kim, Su Jung Hwang, Sun Ho Lee, Wonsik Lee, Hyo Jong Lee, Chung Sub Kim

## Abstract

Antibiotic resistance is one of the world’s most urgent public health problems and therefore novel antibiotics to kill drug-resistant bacteria are desperately needed. So far, natural product-derived small molecules have been the major sources for new antibiotics. Here we describe a family of antibacterial metabolites isolated from a probiotic bacterium *Bacillus licheniformis*. Cross-streaking assay followed by activity-guided isolation yielded a novel antibacterial metabolite bacillimidazole G, which possesses a rare imidazolium ring in the structure, showing MIC values of 0.7–2.6 *μ*g/mL against human pathogenic Gram-positive and Gram-negative bacteria including methicillin-resistant *Staphylococcus aureus* (MRSA) and a lipopolysaccharide(LPS)-lacking *Acinetobacter baumannii* Δ*lpxC*. Bacillimidazole G also lowered MICs of colistin, a Gram-negative antibiotic, up to 8-fold against wild-type *E. coli* MG1655 and *Acinetobacter baumannii*. We propose biosynthetic pathway of the characterized metabolites based on the precursor-feeding studies, chemical biological approach, biomimetic total synthesis, and biosynthetic genes knockout method.

**TOC/Abstract Graphic:** 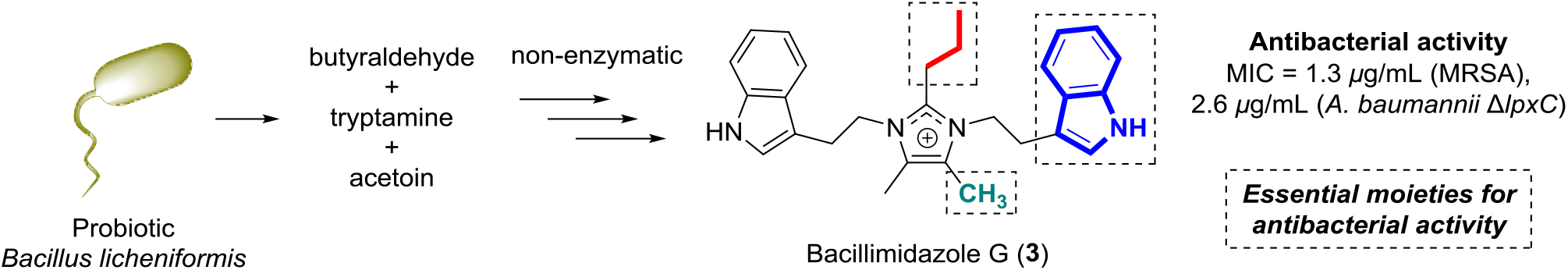

## Introduction

Bacteria evolve resistance to antibiotics that is a major global challenge for public health and more than 700,000 people die each year worldwide due to drug-resistance.^[1]^ The overuse of antibiotics is the main reason for this phenomenon and current antibiotics are becoming less and less effective. Moreover, many pharmaceutical companies have stopped to develop novel antibiotics with only 30– 40 new antibacterial agents in the clinical trial phases while about 4,000 new immuno-oncology agents are in development.^[2]^. Since 1986, no new class of antibiotic has been discovered,^[3]^ and therefore, the need for antibiotics with new chemistry and biology is urgent.

Probiotics are live microorganisms that, when administered to hosts, provide health benefits especially by improving or restoring the gut flora. *Bifidobacterium* and *Lactobacillus* bacteria are the most widely used probiotics and also exist predominantly in the human.^[4-7]^ These beneficial bacteria have been reported to show the antagonistic effects on other microorganisms by modulating the immune system, strengthening the gut epithelial barrier, competitively adhering to the mucosa and epithelium, and secreting antimicrobial substances.^[8]^ Several *Bacillus* species such as *B. subtilis, B. licheniformis, B. cereus, B. clausii, B. polyfermenticus, B. pumilus*, and *B. coagulans* are the commercial probiotics in use worldwide^[9-10]^ and these bacteria are reported to produce antibacterial molecules. For examples, bacitracin, bacteriocin, surfactin, fengycin, and iturin A are well-known natural antibiotics from *Bacillus* species.^[11-13]^ However, most of these *Bacillus*-derived antibiotics are peptide-based which indicated that they would lose their antibacterial activity by rapid protease-mediated degradation and by hepatic and renal clearance in the hosts. *B. licheniformis* is a Gram-positive, endospore-forming organism isolated from soil, human feces, and marine environments.^[14-16]^ This bacterium has been used as a probiotic combined with other probiotic *Bacillus* strains with product names of Biosporin^®^ in Ukraine and Russia and MegaSporeBiotic in the United States,^[9]^ and also used to ferment many Asian foods which showed anti-obesity, anti-diabetic, and anti-Alzheimer’s disease properties.^[17-18]^

In this study, we describe structural and biosynthetic characterization of a new family of non-peptide imidazolium molecules and related analogs in a probiotic bacteria *B. licheniformis* and explore their antibiotic properties against human pathogenic Gram-positive and Gram-negative bacteria as well as their cytotoxic and anti-inflammatory activities. Among the isolated metabolites of *B. licheniformis*, bacillimidazole G (**3**) exhibited strong antibacterial activity against the human pathogenic Gram-positive bacterium, methicillin-resistant *Staphylococcus aureus* (MRSA), and potentiated colistin’s antibacterial effect on the two Gram-negative bacteria, *Acinetobacter baumannii* and *Escherichia coli*.

## Results and Discussion

### Activity-Guided Isolation and Structure Elucidation of Metabolites 1–13

To discover new non-peptide antibacterial agents from *Bacillus* species, first we performed cross-streaking assay with *B. licheniformis* ATCC14580 (KCTC 1918) against several bacteria our research group owns. We observed significant growth inhibition against a Gram positive bacteria *Lacticaseibacillus paracasei* subsp. *paracasei* ATCC25302 (KCTC 3510) (Figure 1A). Then, we cultured 6 L of *B. licheniformis* and organic metabolites were extracted into EtOAc. The dried organic extract was fractionated by HPLC and a disk diffusion assay was carried out for the 26 resulting fractions against *L. paracasei* subsp. *paracasei* ATCC25302 (Figure 1B). The most active fraction #16 showed 0.95 cm of inhibition zone (Figure 1C) and the LC-MS analysis suggested that the major compound (**1**) in this fraction was a not peptide-based molecule deduced by absence of oxygen atom in the expected molecular formula, C_26_H_29_N_4_^+^ (observed mass: *m*/z 397.2403; calculated mass: *m*/z 397.2387). Therefore, we isolated this molecule (**1**) and confirmed it to be the potent antibacterial molecule in the fraction #16 by the same disk diffusion assay (Figure 1D).

**Figure 1.**
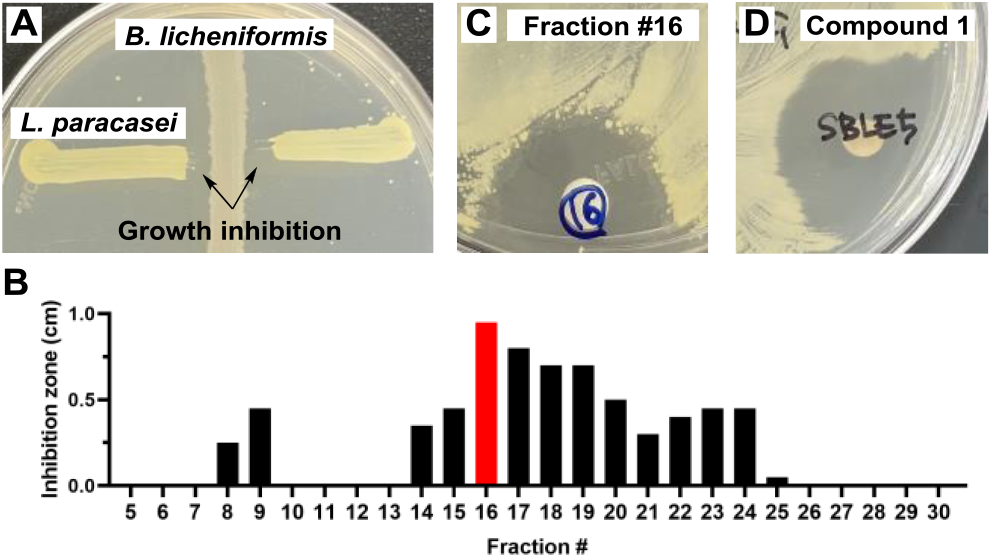
Antibacterial assay-guided isolation of bacillimidazole D (**1**) from *B. licheniformis* extract. (A) Cross-streaking assay result that showed growth inhibitory effect of *B. licheniformis* against *L. paracasei*. (B) Inhibition zone data of subfractions of *B. licheniformis* extract. Fraction #16 showing the most powerful antibacterial activity is marked with red color. (C, D) Pictures of the disk diffusion assay on the fraction #16 and bacillimidazole D (**1**, 200 *μ*g/disk), purified from the fraction #16.

The chemical structure of **1** was determined to be a rare imidazolium-containing molecule as shown in Figure 2A by conventional 1D and 2D NMR data analysis and chemical synthesis (Supporting Information). While we prepare this manuscript, Yan et al. reported the same molecule isolated from marine *Bacillus* sp. WMMC1349 and named it bacillimidazole D.^[19]^ However, interestingly, in the previous study by Yan et al. bacillimidazole D did not show antibacterial activity [MICs > 50 *μ*M against methicillin-resistant *Staphylococcus aureus* (MRSA), *B. subtilis*, and *E. coli*]. During the course of our efforts, we additionally isolated 13 structurally related metabolites (**2–8, 9a, 9b**, and **10– 13**, Figure 2A) using HPLC techniques including chiral separation. Their structures were similarly characterized via NMR analysis and/or synthesis (Figure 2B and SI), and ECD simulation (Figure 2C). We found that eight compounds (**3**–**5, 8, 9a, 9b, 10**, and **12**) were not previously reported and named them bacillimidazoles G–J (**3**–**5** and **8**), bacillindoles A–C (**9a, 9b**, and **10**), and bacillipyrrole B (**12**) based on their core scaffold and structurally similar compounds previously reported.^[19-20]^ The other five known compounds were identified as bacillimidazoles D–F (**2, 6**, and **7**),^[19]^ tryptamine (**11**),^[21]^ and (2-phenylethyl)acetamide (**13**)^[22]^ by comparison of their NMR data with the literature values.

**Figure 2.**
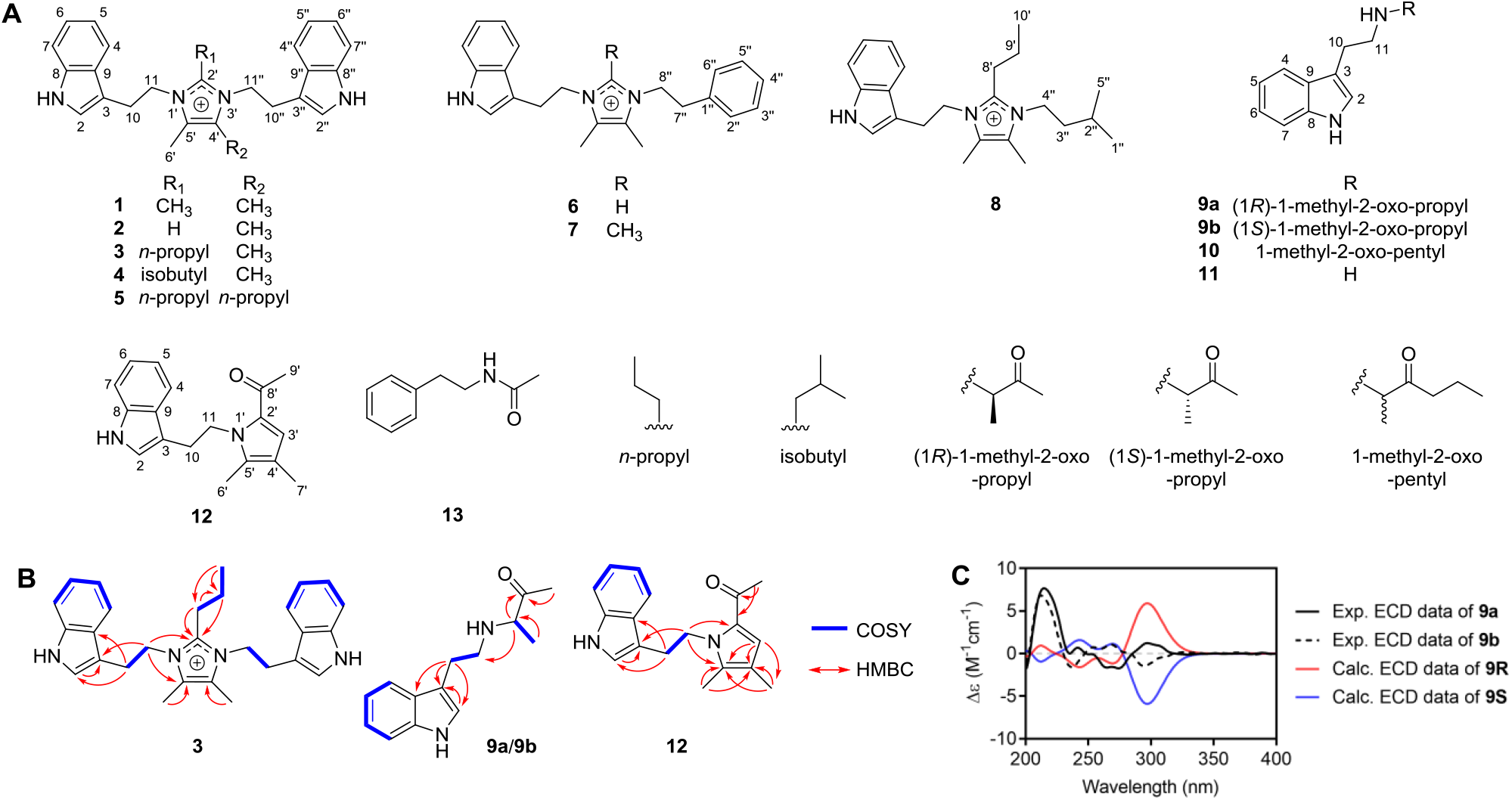
Structure characterization of isolated compounds. (A) Chemical structures of compounds **1**–**13**. (B) Key COSY (blue bold) and HMBC (red arrows) correlations of new compounds **3, 9a/9b**, and **12**. (C) Experimental and calculated ECD spectra of **9a** and **9b** (**9R**: 3′*R*-form, **9S**: 3′*S*-form)

### Biosynthetic Precursors Identification and Biomimetic Synthesis of the Metabolites

Given that the previous studies that reported bacillimidazoles A–F^[19]^ and discolin A ^[23]^ having the same imidazolium core structure proposed that diacetyl is a biosynthetic precursor, we fed diacetyl with four different concentrations individually (0.01, 0.1, 1, 10 mM) in the *B. licheniformis* culture to see if it could enhance the production of metabolites. Surprisingly, there was no significant change in the production level of **1, 2, 6**, and **7** at 0.01 and 0.1 mM of diacetyl (Figures 3A and S2) while there were cell toxicity at 1 and 10 mM of diacetyl. In *Bacillus* spp., diacetyl is produced via nonenzymatic oxidative decarboxylation process from (*S*)-2-acetolactic acid whereas acetoin, a reduced analog of diacetyl, is enzymatically produced from the same precursor (*S*)-2-acetolactic acid.^[24-25]^ Considering this in vivo biological context we assumed that acetoin would be more relevant precursor for synthesizing the imidazolium structure than diacetyl. As expected, we observed ∼4.4-fold enhanced production of **2** from the *B. licheniformis* culture fed with 10 mM of acetoin (Figure 3A). We also observed ∼13.7-fold increase in metabolite **2** production from the tryptamine-fed culture (Figure 3A) suggesting tryptamine as the other biosynthetic precursor of **2**. The last expected biosynthetic precursor for imidazolium scaffold is an aldehyde and we assumed that formaldehyde would be used for production of 2-unsubstituted imidazolium metabolites such as **2** and **6**. To test formaldehyde production from *B. licheniformis* we cultured *B. licheniformis* in presence of Fluoral-P, which reacts with formaldehyde to yield 3,5-diacetyl-1,4-dihydro-2,6-lutidine (DDL) (Figure 3B, left).^[26-27]^ We confirmed DDL production from the *B. licheniformis* culture supplemented with Fluoral-P by comparing its extracted ion chromatogram (EICs; 194.1176, [M+H]^+^ of DDL) with that of standard DDL obtained from reaction product of formaldehyde with Fluoral-P in LC-MS analysis (Figure 3B, right). Metabolites **4** and **8** possessed isobutyl and isopentyl groups, respectively, which indicated that these two substitutes could be derived from L-Leu. Therefore, we supplemented *B. licheniformis* cultures with L-[1,2-^13^C]-Leu and analyzed their metabolite extracts by LC-MS. Expected 1 Da mass shifts were observed for both **4** and **8** (Figure 3C), establishing their L-Leu-derived origin.

**Figure 3.**
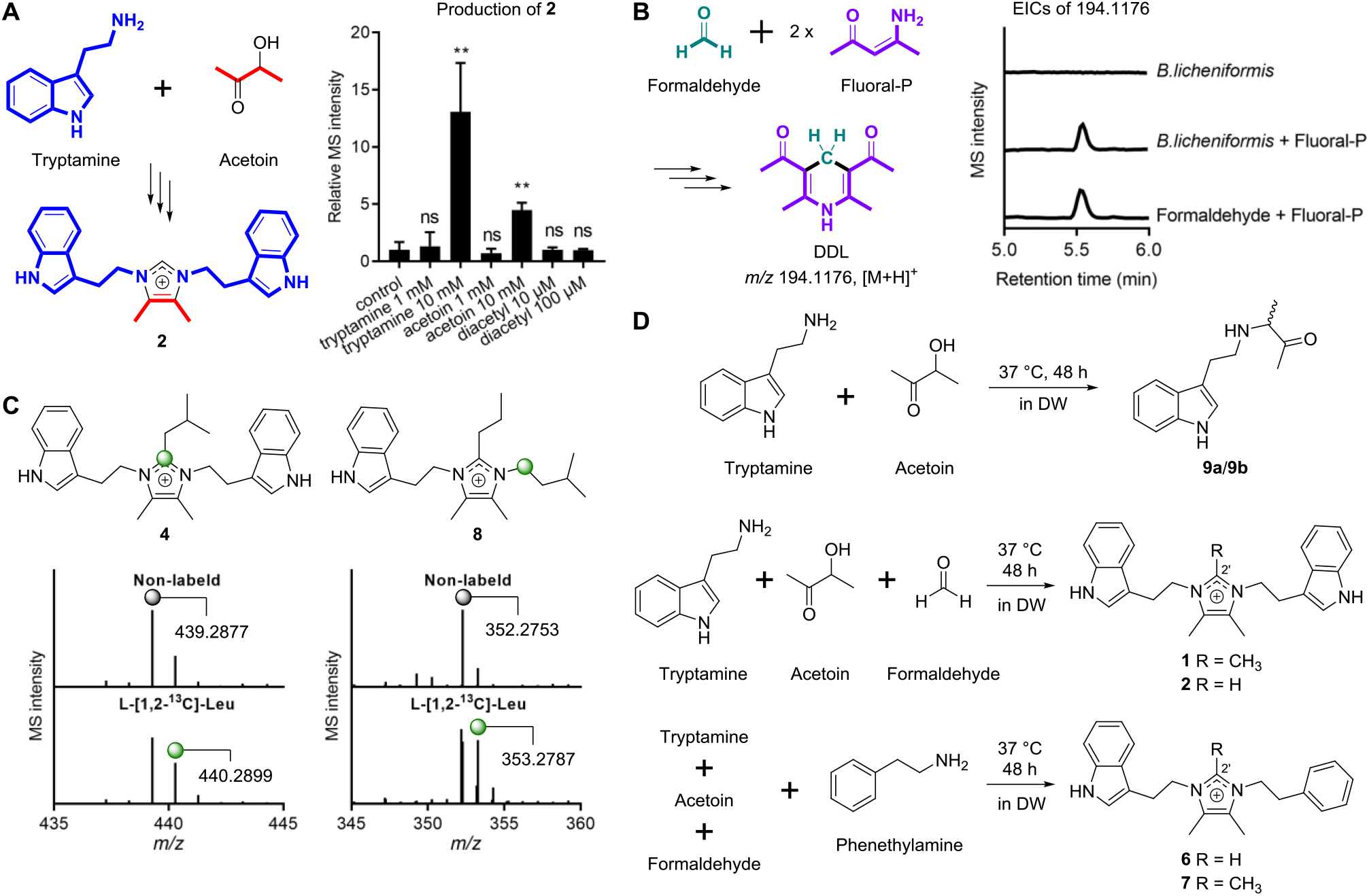
Identification of biosynthetic origins of the isolated compounds. (A) Individual tryptamine, acetoin, and diacetyl feeding experiments results indicating tryptamine and acetoin as dominant precursor of **2**. The bar graphs were generated based on the extracted ion chromatograms (EICs) of *m*/*z* 383.2230 [M]^+^. *n* = 3 biological replicates. Data are mean ± SD. \*\**p* < 0.01. ns, not significant.Two-tailed *t*-test. (B) Confirmation of formaldehyde as a metabolite produced from *B. licheniformis*. 3,5-diacetyl-1,4-dihydro-2,6-lutidine (DDL) is produced from formaldehyde and fluoral-P by spontaneous reaction (left). Production of DDL was observed from *B. licheniformis* cultures supplemented with fluoral-P by EICs of *m*/*z* 194.1176 (right). (C) Proposed ^13^C-isotope labeling pattern of **4** and **8** (top) and their HRMS spectra from organic extracts of *B. licheniformis* cultivated with L-[1,2-^13^C]-Leu (bottom). (D) Biomimetic total synthesis study for **1, 2, 6, 7**, and **9a**/**9b**. Retention times in LC-MS experiments and ^1^H NMR of all the synthesized compounds were identical with those of natural compounds.

Upon characterizing the structures and biosynthetic precursors of the isolated metabolites as described above, we proposed that two biogenic amines (e.g. tryptamine, phenethylamine, and isopentylamine), an α-hydroxy ketone (e.g. acetoin and 3-hydroxyhexan-2-one), and an aldehyde (e.g. formaldehyde, butyraldehyde, and isovaleraldehyde) could spontaneously react to produce the imidazolium metabolites **1–8**. Also, three linear indole metabolites **9a, 9b**, and **10** were assumed to be generated from a tryptamine and an acetoin/a 3-hydroxyhexan-2-one. To test this proposal, we incubated the precursors in three different combinations in the absence of bacteria (37 °C, 48 h, in DW); 1) tryptamine and acetoin, 2) tryptamine, acetoin, and formaldehyde, and 3) tryptamine, phenethylamine, acetoin, and formaldehyde (Figure 3D). As anticipated, compounds **9a/9b, 2**, and **6** were produced from each reaction mixture, and unexpectedly, compounds **1** and **7**, 2′-methylated analogs of **2** and **6**, respectively, were also detected indicating that the methyl group in **1** and **7** would be derived from formaldehyde. Structures of all the synthesized compounds **1, 2, 6, 7**, and **9a/9b** were identified to be the same as the bacteria-derived metabolites by comparing their ^1^H NMR data and LC-MS retention times (Supporting Information).

### Biosynthetic Proposal of the Metabolites

Acetoin is a well-known compound widely used in foods, cosmetics, detergents, plant growth promoters and biological pest controls and produced from mammals including human, plants, fungi, and bacteria.^[24]^ Two biosynthetic genes for production of acetoin, *alsS* and *alsD*, are conserved in many bacteria such as *Bacillus* spp. (*B. subtilis, B. licheniformis, B. amyloliquefaciens*, and *B. pumilus*), *Paenibacillus polymyxa, Serratia marcescens, Enterobacter cloacae, E. aerogenes, Klebsiella oxytoxa*, and *K. pneumoniae* (Figure 4A).^[24]^ In *Bacillus* spp., AlsS, 2-acetolactate synthase, produces (*S*)-2-acetolactic acid from two pyruvic acid molecules via decarboxylative condensation reaction and AlsD, 2-acetolactate decarboxylase, decarboxylates (*S*)-2-acetolactic acid to generate (*R*)-acetoin (Figure 4B).^[24]^ As mentioned above, diacetyl is a non-enzymatic spontaneous degradation product of (*S*)-2-acetolactic acid. To confirm that these two enzymes are involved in biosynthesis of the isolated metabolites we analyzed the culture extracts of wildtype (WT) strain of *B. subtilis* and its two single mutant strains of Δ*alsD* or Δ*alsS* by LC-MS. We observed ∼3.0-fold decrease in the production of **1** in the Δ*alsD* mutant strain compared to WT strain while deletion of *alsS* dramatically impaired the production of **1** (Figure 4C). The other isolated metabolites also showed the similar patterns (Figure S3). This data indicated that, in the Δ*alsD* mutant strain, diacetyl produced non-enzymatically from (*S*)-2-acetolactic acid would still contribute to synthesizing the metabolites. Consequently, we suggest that the enzymatic product (*R*)-acetoin is the major biosynthetic precursor while the non-enzymatic product diacetyl is also the minor contributor. Here we report the first genetic supports that (*R*)-acetoin and diacetyl are involved in biosynthesis of bacterial imidazolium metabolites.

**Figure 4.**
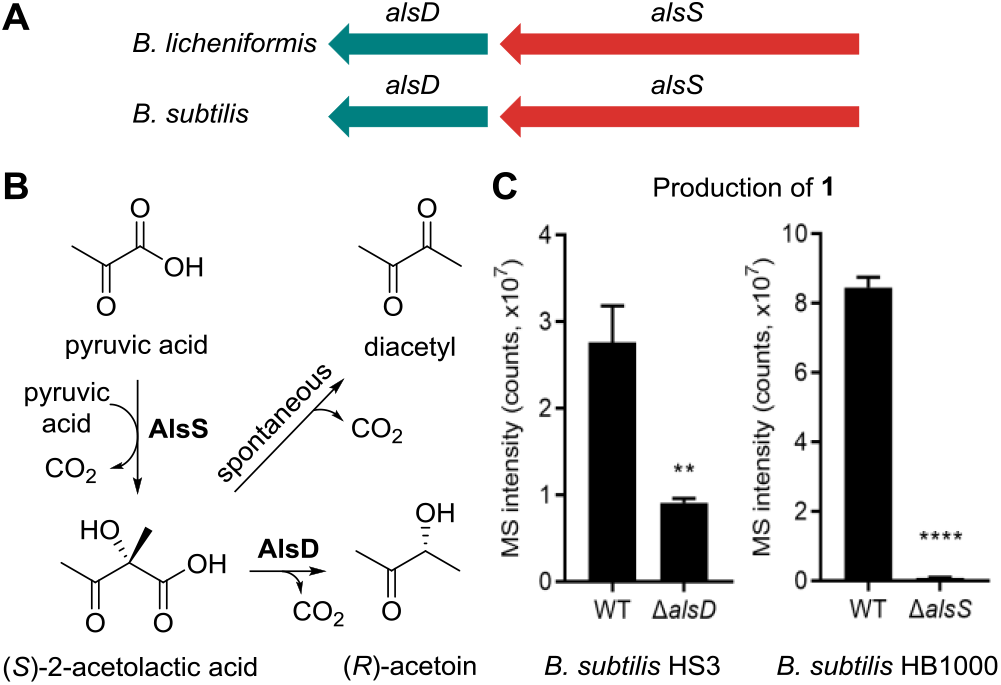
Characterization of biosynthetic genes for the isolated compounds. (A) Two genes for biosynthesis of (*R*)-acetoin (*alsD* and *alsS*) conserved in *B. licheniformis* and *B. subtilis*. (B) 2-acetolactate synthase (AlsS) and 2-acetolactate decarboxylase (AlsD) produce (*R*)-acetoin from two pyruvic acid molecules. Diacetyl is produced from by non-enzymatical decarboxylation. (C) Comparison of production levels of **1** between WT *B. subtilis* HS3 and its *alsD* mutant strain (left) and between WT *B. subtilis* HB1000 and its *alsS* mutant strain (right). The bar graphs were generated based on the EICs of *m*/*z* 397.2387 [M]^+^. *n* = 3 biological replicates. Data are mean ± SD. \*\**p* < 0.01, \*\*\*\**p* < 0.0001. Two-tailed *t*-test.

With these diverse chemical and biological data on the isolated metabolites in hand, we then able to propose their biosynthetic pathway (Figure 5). (*R*)-acetoin, the AlsS/AlsD product, would react with tryptamine (**11**) produced from L-Trp to form an imine-containing linear molecule **i**, which then converts to racemate **9a/9b** by α-ketol rearrangement. Addition of one more tryptamine molecule to **9a/9b** followed by tautomerization would generate an intermediate **ii** and this would be coupled with formaldehyde to produce the imidazolium metabolite **2** through iminium formation, cyclization, and oxidation. Compound **1** would be formed by addition of a formaldehyde molecule to **2** (Figure 5A). Compounds **3** and **4** possess a *n*-propyl and an isobutyl group, respectively, instead of the methyl group in **1** suggesting that butyraldehyde and isovaleraldehyde derived from L-Leu would couple to **ii** to generate **3** and **4**, respectively (Figure 5B). Similar to the biosynthetic proposal of **9a/9b, 10** would be formed from **11** and 3-hydroxyhexan-2-one, and then **5** would be produced in the presence of tryptamine and butyraldehyde (Figure 5C). The other two biogenic amines, phenethylamine and isopentylamine, than tryptamine would contribute to synthesize **6/7** and **8**, respectively (Figure 5D and 5E).

**Figure 5.**
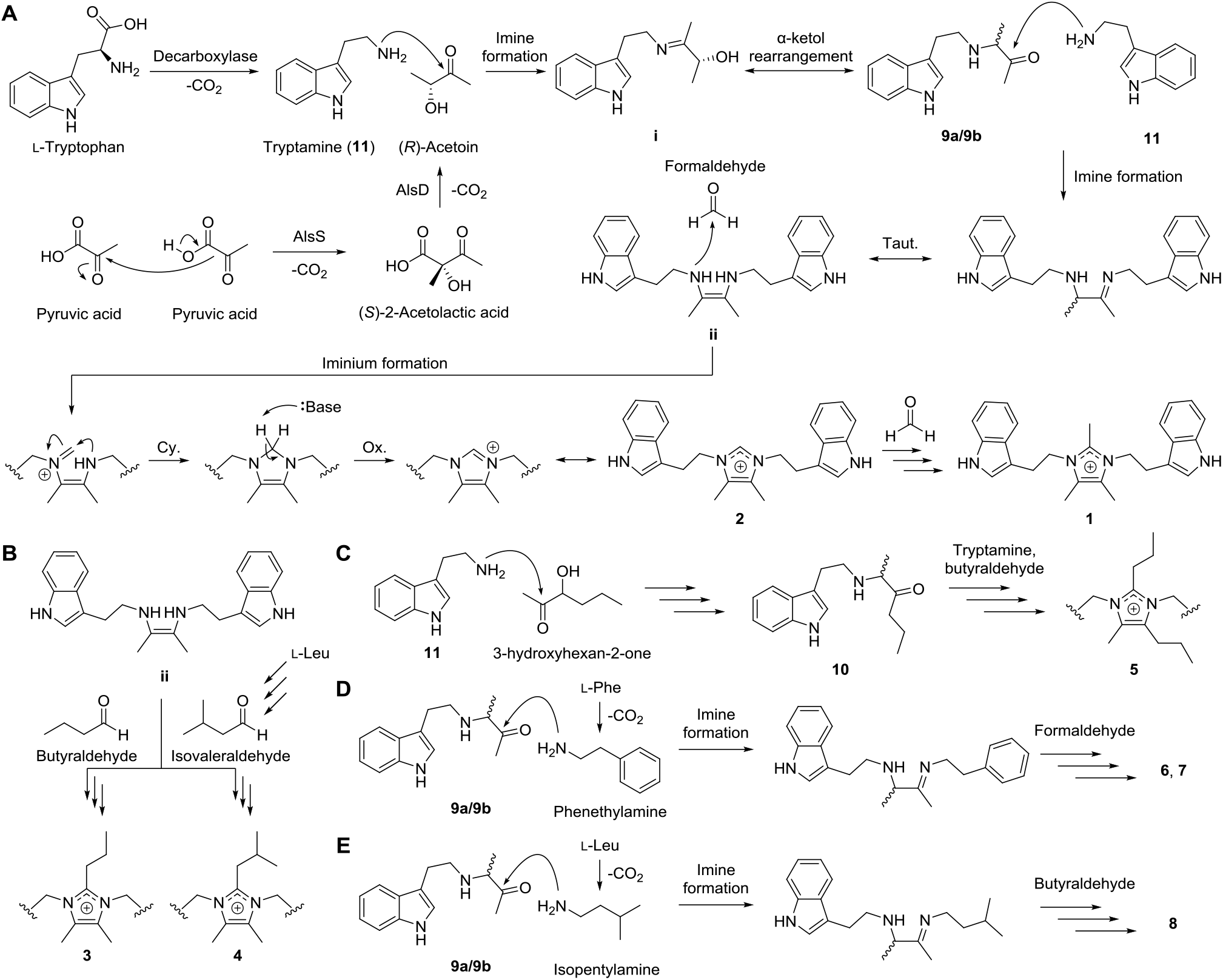
Proposed biosynthetic pathway of **1**–**11** (A: **1, 2, 9a, 9b**, and **11**, B: **3** and **4**, C: **5** and **10**, D: **6** and **7**, E: **8**). Taut., tautomerization. Cy., cyclization. Ox., oxidation.

### Antibacterial Activity of the Metabolites

We then tested all the isolated metabolites **1–13** for their antibacterial activities. As shown in Table 1, the most powerful compound was bacillimidazole G (**3**), a new metabolite discovered in this study, displaying 0.7–1.3 *μ*g/mL (or 1.56–3.13 *μ*M) of MIC values against methicillin-resistant *Staphylococcus aureus* (MRSA) USA300 and other two Gram positive bacteria *L. paracasei* subsp. *paracasei* ATCC25302 and *Brevibacterium epidermidis* ATCC35514. Other imidazolium-containing metabolites **1, 2, 4, 5**, and **8** and the linear indole metabolite **9** (enantiomeric mixture of **9a** and **9b**) also exhibited mild antibacterial activities with MICs ranging 6–22 *μ*g/mL (or 12.5–50 *μ*M). While analyzing the antibacterial data against *L. paracasei* subsp. *paracasei* ATCC25302 we found interesting structure-activity relationships (SARs) among the imidazolium metabolites (Figure 6). First, substitution of *n*-propyl group at C-2′ with hydrogen, methyl group, or isobutyl group led to 32-fold decrease in activity (**3**, 1.56 *μ*M; **1, 2**, and **4**, 50 *μ*M). Second, existence of an indole group at C-10′′ enhanced the activity as deduced from the loss of activity in **6** and **7** having a phenyl group (> 100 *μ*M) and the 16-fold reduced activity in **8** possessing a isopropyl group (25 *μ*M) instead of an indole group in **3** (1.56 *μ*M). Last, we observed 8-fold decrease in the activity when the methyl group at C-4′ in **3** (1.56 *μ*M) was replaced by a *n*-propyl group as in **5** (12.5 *μ*M). The similar SARs against the other two Gram-positive bacteria *B. epidermis* ATCC35514 and MRSA led us to propose *n*-propyl, methyl, and indole groups at C-2′, 4′, and 10′′, respectively, as key functionalities for exhibiting the antibacterial activity against Gram-positive bacteria. However, the isolated metabolites showed no antibacterial activity against Gram-negative bacteria *E. coli* MG1655 and *Acinetobacter baumannii* (MICs >100 *μ*M).

**Table 1.** MIC values (in *μ*g/mL and *μ*M) of compounds **1–5, 8**, and **9** against six bacterial strains.

| Comp. | MIC [$\mu\text{g/mL}$ ($\mu\text{M}$)] | | | | | |
| --- | --- | --- | --- | --- | --- | --- |
|  | Gram-positive bacteria |  |  | Gram-negative bacteria |  |  |
| | *L. paracasei*<br>subsp.<br>*paracasei* | *B.*<br>*epidermis* | MRSA | *E. coli*<br>MG1655 | *A.*<br>*baumannii* | *A.*<br>*baumannii*<br>$\Delta\text{lpxC}$ |
| **1** | 20 (50) | > 100 $\mu\text{M}$ | 20 (50) | > 100 $\mu\text{M}$ | > 100 $\mu\text{M}$ | 40 (100) |
| **2** | 19 (50) | > 100 $\mu\text{M}$ | > 100 $\mu\text{M}$ | > 100 $\mu\text{M}$ | > 100 $\mu\text{M}$ | 38 (100) |
| **3** | 0.7 (1.56) | 1.3 (3.13) | 1.3 (3.13) | > 100 $\mu\text{M}$ | > 100 $\mu\text{M}$ | 2.6 (6.25) |
| **4** | 22 (50) | > 100 $\mu\text{M}$ | > 100 $\mu\text{M}$ | > 100 $\mu\text{M}$ | > 100 $\mu\text{M}$ | 22 (50) |
| **5** | 6 (12.5) | 11 (25) | 11 (25) | > 100 $\mu\text{M}$ | > 100 $\mu\text{M}$ | 11 (25) |
| **8** | 9 (25) | > 100 $\mu\text{M}$ | > 100 $\mu\text{M}$ | > 100 $\mu\text{M}$ | > 100 $\mu\text{M}$ | 9 (25) |
| **9<sup>a</sup>** | 6 (25) | 12 (50) | 12 (50) | > 100 $\mu\text{M}$ | n.t.<sup>b</sup> | n.t. |

<sup>a</sup>Racemic mixture of **9a** and **9b**.<sup>b</sup>not tested **Figure 6.** Structural elements of imidazolium antibiotics that significantly enhance antibacterial activities and structure-activity relationships (SARs) among the isolated metabolites.

**Figure 6.**
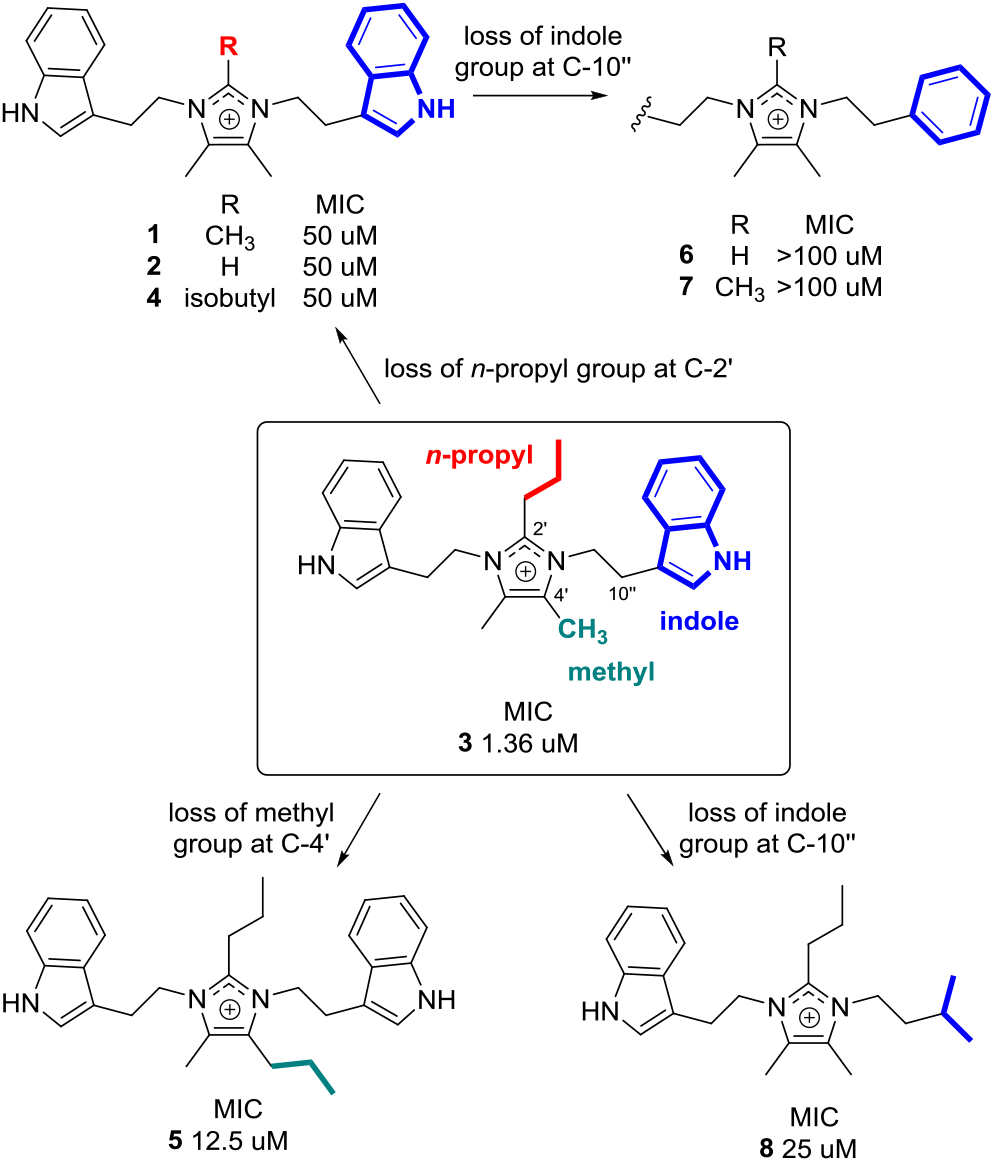
Structural elements of imidazolium antibiotics that significantly enhance antibacterial activities and structure-activity relationships (SARs) among the isolated metabolites.

Gram-negative bacteria have intrinsically antibiotic-resistant, and this is largely due to their outer membrane that prevents the entry of antibiotics inside the cell.^[28]^ As noted, unlike other Gram-negative bacteria, in *A. baumannii* lipopolysaccharide (LPS) deficient mutants which have mutations in the early stage of the lipid A biosynthesis including *lpxC* are still viable.^[29]^ However, their cell permeability is affected significantly, which results in increased susceptibility to most all classes of antibiotics such as azithromycin, rifampicin, and vancomycin.^[30]^ Also, the inhibition of deacetylase, LpxC by a small molecule PF-5081090 demonstrated similar effects showing an increased susceptibility to antibiotics in *A. baumannii*.^[31]^ Therefore, we examined our compounds for their antibacterial potency against *A. baumannii ΔlpxC*. Surprisingly, all compounds that have potency against three Gram-positive bacteria (**1**–**5** and **8**) showed a dramatic improvement in their MICs (up to 16-fold) of the *lpxC* mutant compared to wild type. Importantly these MICs were comparable to those of the tested Gram-positive bacteria *L. paracasei* subsp. *paracasei* ATCC25302 (Table 1). These data suggest that increased cell permeability in the *lpxC* mutant, LPS deficient allows the antibacterials **1**–**5** and **8** to penetrate the bacterial cells as observed with other antibiotics.^[32]^ To further investigate this mechanism of action, we sought to find possible synergism with a membrane-targeting antibiotic, colistin. This is because this colistin has the mechanism of disrupting the outer membrane of Gram-negative bacteria, and therefore, *lpxC* mutant that does not have the cellular target of colistin show strong resistance to it. To this end, we examined our most potent bacillimidazole G (**3**) in the presence of colistin by a checkerboard assay with wild-type Gram-negative bacteria; *E. coli* MG1655 and *A. baumannii*. As shown in Figure 7, the colistin MICs in the presence of bacillimidazole G (**3**) (0.32–84 *μ*g/mL) were improved significantly: for *E. coli*, 8 folds (2.5 *μ*g/mL to 0.313 *μ*g/mL); *A. baumannii*, 4 folds (0.2 *μ*g/mL to 0.05 *μ*g/mL). Colistin which was introduced in early 1970, is still used in the clinic to treat multidrug-resistant *A. baumannii* including carbapenem-resistant *A. baumannii* (CRAB), but its clinical use is limited due to its cytotoxicity and high frequency of resistance to colistin.^[33]^ We note here that our combinatory treatment with a low dose of colistin (well below 4-folds) and our newly discovered compound provides a novel therapeutic option against Gram-negative bacterial infection. Collectively, bacillimidazole G (**3**) could be utilized to treat Gram-positive bacteria including MRSA, and also combination with our compound along with colistin could be treated against Gram-negative bacteria such as CRAB, suggesting possible further development of our compounds to broad-spectrum antibiotics.

**Figure 7.**
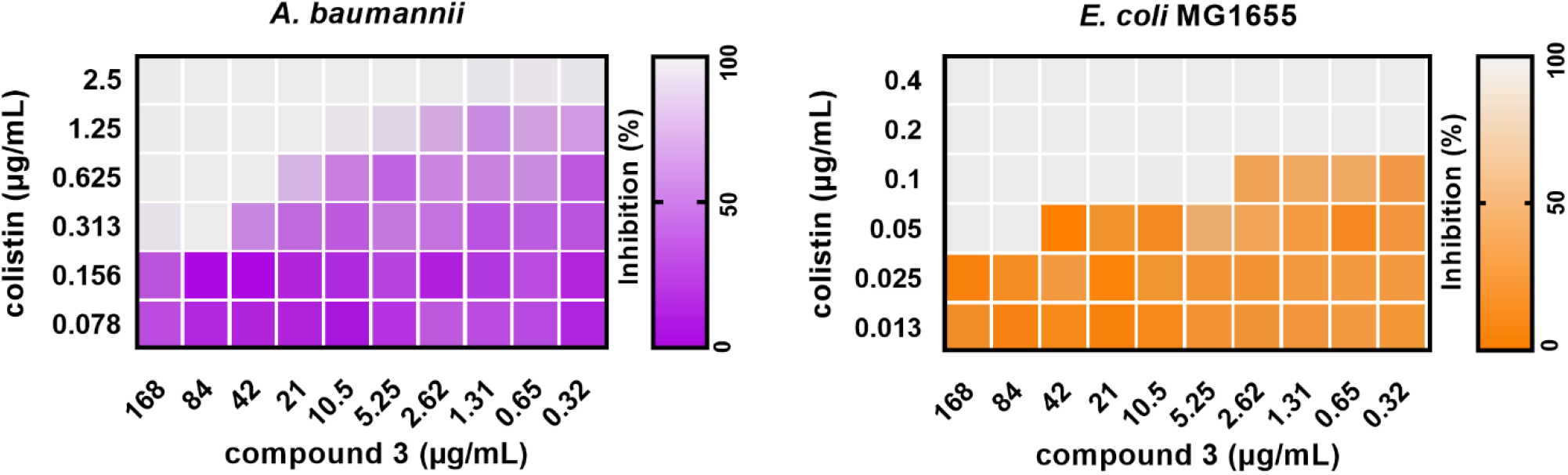
Checkerboard assay results of **3** with colistin against *A. baumannii* (left) and *E. coli* MG1655 (right)

### Cytotoxic and Anti-inflammatory Activity of the Metabolites

There are several structurally similar compounds isolated or synthesized possessing the imidazolium moiety, such as lepidilines A–D, and these compounds showed potent cytotoxicity against human cancer cell lines.^[34-35]^ Therefore, we also evaluated cytotoxic activity of **1–13** against four different human cancer cell lines using HCT-116 (colorectal carcinoma cell line), A549 (lung adenocarcinoma cell line), HepG2 (hepatocellular carcinoma cell line), HT1080 (fibrosarcoma cell line) and normal cell lines such as Wi38 (human fetal lung fibroblast cell line), Chang (human normal liver cell line), and NIH3T3 (mouse fibroblast cell line) cells. As shown in Table S4, all metabolites except compound **5** showed no cytotoxicity. However, compound **5** showed as similar cytotoxicity as for a well-known chemotherapy agent, 5-FU (5-fluorouracil) in HCT-116 and A549 cells (Table S4). It is worth noting that for compound **5** we observed much higher cytotoxicity against HepG2 and HT1080 cells than 5-FU. Interestingly, compound **5** showed relatively little toxicity to normal cells (Wi38, Chang, and NIH3T3 cells). Otherwise, 5-FU showed significant cytotoxicity to normal cells, especially in NIH3T3 fibroblast cells. Thus, these results suggest the possibility that **5** may be developed as a potential anti-cancer agent, because of its selective cytotoxicity. Next, we evaluated the inhibitory effect of metabolites **1**-**13** on endotoxin-mediated nitric oxide (NO) production in RAW 264.7 cells. Except for **4** and **7**, most metabolites were found to inhibit NO production in Figure S5. In particular, **5, 8**, and **12** were observed to suppress NO more strongly. Since NO acts as a major medium in the initial inflammatory response, further research is needed on whether these metabolites can suppress inflammatory diseases and how such effects occur. In addition, the fact that the metabolite **5** shows both anti-cancer and anti-inflammatory activity suggests that **5** may not only inhibit the growth of cancer cells itself but also suppress the inflammatory tumor microenvironment at the same time.

## Conclusion

Although probiotics are considered to show diverse beneficial effects on human and have been widely used to improve human health little is known about underlying mechanisms mediated by specialized antibacterial small molecules. Here, we characterize a family of antibacterial metabolites, some of which have a rare imidazolium scaffold, from a well-known probiotic strain, *B. licheniformis* ATCC14580. The most potent antibacterial metabolite was the previously unreported compound bacillimidazole G (**3**) with MICs of 0.7–1.3 *μ*g/mL against three Gram-positive bacteria, MRSA, *L. paracasei* subsp. *paracasei* ATCC25302 and *B. epidermidis* ATCC35514 and of 2.6 *μ*g/mL against LPS-lacking Gram-negative bacteria *A. baumannii* Δ*lpxC*. Although bacillimidazole G (**3**) was not active against two Gram-negative bacteria, *E. coli* MG1655 and wild-type *A. baumannii*, it lowered MIC values of a Gram-negative antibiotic colistin up to 8-fold. We also propose biosynthetic pathway of the characterized metabolites based on the precursor-feeding studies, chemical biological approach, biomimetic total synthesis, and biosynthetic genes knockout method. Strikingly, we prove that α-hydroxy ketone such as acetoin and 3-hydroxyhexan-2-one is a more biologically relevant precursor for synthesizing the imidazolium compounds (**1–8**) than diketone such as diacetyl, which was the previously proposed precursor by the other groups.^[19, 23]^ In sum, our studies present molecular evidences for the antibiotic property of probiotic bacteria *B. licheniformis* and suggest bacillimidazole G (**3**) as a potential antibacterial agent against Gram-positive and Gram-negative bacteria to overcome the life-threatening infectious disease.

## Acknowledgments

This work was supported by the National Research Foundation of Korea (NRF) grant funded by the Korean government (MSIT) (No. 2021R1C1C1011045, 2022R1A6A1A03054419), by the BK21 FOUR Project, and by the National Supercomputing Center with supercomputing resources including technical support (KSC-2021-CRE-0273).

## Notes

### Competing Interest Statement

The authors have declared no competing interest.

