## Supporting Information for "Discovery and biosynthesis of imidazolium antibiotics from a probiotic *Bacillus licheniformis*"

### Experimental section

**Instruments.** Electronic circular dichroism (ECD) spectra were measured on a JASCO J-1500 CD spectrometer (JASCO, Easton, MD, USA). Nuclear magnetic resonance (NMR) spectra ($^{1}$H, $^{13}$C, $^{1}$H-$^{1}$H COSY, HSQC, and HMBC) were recorded with a Bruker AVANCE III HD 800 NMR spectrometer equipped with a 5 mm TCI CryoProbe operating at 850 MHz ($^{1}$H) and 212.5 MHz ($^{13}$C), with chemical shifts given in ppm (δ) (Bruker, Karlsruhe, Germany). All HRESIMS spectra were obtained using an Agilent G6545B quadrupole time-of-flight mass spectrometer (Agilent Technologies) coupled to an Agilent 1260 Infinity II series furnished a 6545 LC-Q-TOF mass spectrometer (Agilent Technologies) with an Agilent EclipsePlus C18 column (2.1 mm × 50 mm i.d., 1.8 μm; flow rate: 0.3 mL/min). The LC-MS analysis was performed on an Agilent 1260 series HPLC system with a diode array detector and a 6130 Series ESI mass spectrometer equipped with an analytical Kinetex C18 100 Å column (250 mm × 4.6 mm i.d., 5 μm; flow rate: 0.7 mL/min). Semipreparative high performance liquid chromatography (HPLC) was conducted using a Agilent 1260 pump, which was equipped with Luna® C18 100 Å column (250 mm × 10 mm i.d., 5 μm; flow rate: 4.0 mL/min) and Luna 5 μ Phenyl-Hexyl New column (250 mm × 10 mm i.d., 5 μm; flow rate: 4.0 mL/min). Chiral-phase HPLC was carried out using a Gilson 306 pump connected toa Shodex refractive index detector, which was equipped with a Phenomenex Lux® 5 μm Cellulose-1 column (Phenomenex, Torrance, CA, USA).

**Bacterial sources.** *Bacillus licheniformis* ATCC14580 (KCTC 1918), *Lacticaseibacillus paracasei* subsp. *paracasei* ATCC25302 (KCTC 3510), and *Brevibacterium epidermidis* ATCC35514 (KCTC 3090) used in this study were purchased from Korean Collection for Type

Cultures (KCTC). *B. subtilis* HS3 (wild-type and its *ΔalsD* strain) and *B. subtilis* HB1000 (wild-type and its *ΔalsS* strain) were afforded from Dr. Siger Holsappel (Center for Llifesciences, University of Groningen, Netherlands) and Dr. Deepika Awasthi (Lawrence Berkeley National Laboratory, University California Berkeley, California, USA). *Staphylococcus aureus* USA300 and *Acinetobacter baumannii* (wildtype and Δ*lpxC*)[1] were provided from Dr. Wonsik Lee (School of Pharmacy, Sungkyunkwan University, South Korea)

**Cultivation of *B. licheniformis* and metabolites analysis.** *B. licheniformis* KCTC 1918 was grown on Luria-Bertani (LB) agar plates at 37 °C for 1 day. A single colony was inoculated into 5 mL of tryptic soy broth (TSB) broth and incubated at 37 °C for 2 days at 250 rpm. The culture broth was extracted with ethyl acetate, and the ethyl acetate-soluble fraction was evaporated. The dried extract was dissolved in 200 µL of 100% methanol and the metabolites were analyzed using LC-MS equipped with Phenomenex C18 column Luna® 5 µm C18(2) 100 Å (250 × 4.6 mm) column with gradient solvent system (0.7 mL/min, 10-100% acetonitrile, 30 min, 0.01% TFA).

**Isolation of metabolites.** *B. licheniformis* KCTC 1918 was grown in six 5 mL tubes with TSB at 37 °C for 1 day. The cultures were used to inoculate 6 L cultures of TSB in six 4 L Erlenmeyer flasks. After 2 days, the whole culture was extracted with ethyl acetate (6 L, 2 times), and the ethyl acetate-soluble layer was concentrated under reduced pressure to yield a crude extract (5.8 g). The dried extract (5.8 g) was fractionated by semi-preparative HPLC system (Phenomenex, Luna 10 ㎛ C18(2) 250 × 10 mm i.d.) with a gradient elution from 10 to 100% aqueous MeCN with 0.01% trifluoroacetic acid (TFA) over 30 mins (flow rate: 4

mL/min) to afford 30 fractions. Compounds **12** ($t_R$ 8.1 min, 0.7 mg) and **10** ($t_R$ 10.0 min, 0.4 mg) were isolated from the fraction 6 using semi-prep HPLC (Phenomenex, Luna $C_{18}$ 10 µm Phenyl-Hexyl, 250 × 10 mm i.d.) with a gradient system of 10–60% MeCN with 0.01% TFA (flow rate 4 mL/min for 30 min). Fraction 13 was purified by semi-prep HPLC (Phenomenex, Luna $C_{18}$ 10 µm Phenyl-Hexyl, 250 × 10 mm i.d.) with an isocratic solvent of 17 % aqueous MeCN with 0.01% TFA (flow rate 4 mL/min for 30 min) to afford compound **13** ($t_R$ 6.5 min, 0.5 mg). Compounds **11** ($t_R$ 25.7 min, 0.6 mg) and **5** ($t_R$ 10.0 min, 0.4 mg) were purified from fraction 15 through semi-prep HPLC (Phenomenex, Luna $C_{18}$ 10 µm Phenyl-Hexyl, 250 × 10 mm i.d.) eluting with a gradient system of 30–38% MeCN with 0.01% TFA (flow rate 4 mL/min for 30 min). Compound **4** ($t_R$ 8.1 min, 0.5 mg) was isolated from fraction 16 using semi-prep HPLC (Phenomenex, Luna $C_{18}$ 10 µm Phenyl-Hexyl, 250 × 10 mm i.d.) with a gradient system of 40–45% MeCN 0.01% TFA (flow rate 4 mL/min for 30 min). Compounds **6** ($t_R$ 7.7 min, 0.6 mg), **7** ($t_R$ 18.1 min, 0.5 mg), and **1** ($t_R$ 22.2 min, 0.5 mg) were purified from fractions 17 and 18 using semi-prep HPLC (Phenomenex, Luna $C_{18}$ 10 µm Phenyl-Hexyl, 250 × 10 mm i.d.) eluting with a step gradient system of 30–40% MeCN (flow rate 4 mL/min for 15 min) and 40–50% MeCN (flow rate 4 mL/min for another 15 min) with 0.01% TFA. Fractions 19 and 20 were isolated via semi-prep HPLC (Phenomenex, Luna $C_{18}$ 10 µm Phenyl-Hexyl, 250 × 10 mm i.d.) with a gradient system of 40–45% MeCN with 0.01% TFA (flow rate 4 mL/min for 30 min) to yield compounds **8** ($t_R$ 10.4 min, 0.4 mg), **2** ($t_R$ 14.0 min, 0.4 mg), and **3** ($t_R$ 17.8 min, 0.5 mg). Compound **9** ($t_R$ 19.2 min, 0.5 mg) was purified from fraction 25 by conducting semi-prep HPLC (Phenomenex, Luna $C_{18}$ 10 µm Phenyl-Hexyl, 250 × 10 mm i.d.) with a gradient system of 50–60% MeCN with 0.01% TFA (flow rate 4 mL/min for 30 min). ECD analysis revealed that compound **9** was a racemate, which was resolved by chiral-phase semipreparative HPLC eluting with an isocratic mixture of hexanes-*i*PrOH (80:20, 1 mL/min) to afford enantiomers **9a** ($t_R$: 16.0 min, 0.1 mg) and **9b** ($t_R$: 16.8 min, 0.1 mg).

*Bacillimidazole G (3)*: Brownish gum; UV (MeCN/$H_2O$) $\lambda_{max}$ 220, 280 nm; $^1$H MNR (850 MHz) and $^{13}$C NMR (212.5 MHz) data in methanol-$d_4$, see Table 1; HRESIMS (positive-ion mode) *m*/*z* 425.2701 $[M]^+$ (calcd for $C_{28}H_{33}N_4^+$, 425.2700).

*Bacillimidazole H (4)*: Brownish gum; UV (MeCN/$H_2O$) $\lambda_{max}$ 220, 290, 380 nm; $^1$H MNR (850 MHz) and $^{13}$C NMR (212.5 MHz) data in methanol-$d_4$, see Table 1; HRESIMS (positive-ion mode) *m*/*z* 439.2868 $[M]^+$ (calcd for $C_{29}H_{35}N_4^+$, 439.2856).

*Bacillimidazole I (5)*: Brownish gum; UV (MeCN/$H_2O$) $\lambda_{max}$ 220, 280 nm; $^1$H MNR (850 MHz) and $^{13}$C NMR (212.5 MHz) data in methanol-$d_4$, see Table 1; HRESIMS (positive-ion mode) *m*/*z* 453.3072 $[M]^+$ (calcd for $C_{30}H_{37}N_4^+$, 453.3013)

*Bacillimidazole J (8)*: Brownish gum; UV (MeCN/$H_2O$) $\lambda_{max}$ 220, 290 nm; $^1$H MNR (850 MHz) and $^{13}$C NMR (212.5 MHz) data in methanol-$d_4$, see Table 1; HRESIMS (positive-ion mode) *m*/*z* 352.2761 $[M]^+$ (calcd for $C_{23}H_{34}N_3^+$, 352.2747).

*Racemic mixture of bacillindoles A (9a) and B (9b)*: Brownish gum; UV (MeCN/$H_2O$) $\lambda_{max}$ 210, 280 nm; ECD (MeOH) $\lambda_{max}$ ($\Delta\varepsilon$) 265 (-1.11), 274 (-1.16), 297 (0.83), 302 (0.71) nm for **9a**; 237 (-1.07), 254 (0.50), 268 (0.72), 294 (-0.98) nm for **9b**; $^1$H MNR (850 MHz) and $^{13}$C NMR (212.5 MHz) data in methanol-$d_4$, see Table 2; HRESIMS (positive-ion mode) *m*/*z* 231.1490 $[M + H]^+$ (calcd for $C_{14}H_{19}N_2O$, 231.1492).

*Bacillinodle C (10)*: Brownish gum; (MeCN/$H_2O$) $\lambda_{max}$ 220, 280 nm; $^1$H MNR (850 MHz) and $^{13}$C NMR (212.5 MHz) data in methanol-$d_4$, see Table 2; HRESIMS (positive-ion mode) *m*/*z* 259.1804 $[M + H]^+$ (calcd for $C_{16}H_{23}N_2O$, 259.1805).

*Bacillipyrrole B (**12**)*: Brownish gum; UV (MeCN/$H_2O$) $\lambda_{max}$ 220, 310 nm; $^1$H MNR (850 MHz) and $^{13}$C NMR (212.5 MHz) data in methanol-$d_4$, see Table 1; HRESIMS (positive-ion mode) *m*/*z* 281.1660 $[M + H]^+$ (calcd for $C_{18}H_{21}N_2O$, 281.1654).

**Computational analysis.** All conformers of **9** used in this study were found using the Macromodel (version 2019-2, Schrödinger LLC) module with "Mixed torsional/Low-mode sampling" in the MMFF force field. The searches were implemented in the gas phase with a 10 kJ/mol energy window limit and 10,000 maximum number of steps to explore all potential conformers. The Polak−Ribiere Conjugate Gradient (PRCG) method was utilized to minimize conformers with 10,000 iterations and a 0.001 kJ (mol Å)$^{-1}$ convergence threshold on the Root Mean Square (RMS) gradient. All the conformers were subjected to geometry optimization using the Gaussian 16 package (Gaussian Inc.) in the gas phase at B3LYP/6-31G(d) level and proceeded to calculation of excitation energies, oscillator strength, and rotatory strength at B3LYP/6-31G(d) level in the Polarizable Continuum Model (PCM, methanol). The ECD spectra were Boltzmann-averaged based on the calculated Gibbs free energy of each conformer (Table S3) and visualized with SpecDis software (Version 1.71)[2] with a σ/γ value of 0.25 eV.

**MIC assay.** Compounds were prepared in DMSO to a concentration of 10 mM and tested for antibacterial activity at 100, 50, 25, 12.5, 6.25, 3.13, 1.56, 0.78, and 0.39 μM. Bacterial strains were incubated in a 14-mL polystyrene tube containing LB medium for 24 h. After that, the strain cultures were adjusted to $OD_{600}$ = 0.001 in LB medium. LB medium (100 μL) containing compounds at the appropriate concentration was added in the 96-well plate and 100 μL of bacterial culture ($OD_{600}$ = 0.001) was treated to each well. The plate was incubated at 37 °C overnight.

**Checkerboard assay.** To evaluate the synergism of compound **3** and colistin, the checkerboard method was performed. Two 96-well plates were prepared using the serial dilution method with two antibiotics (colistin and compound **3**) in different directions generating 6 x 10 matrix. The 100 μL of bacterial culture in LB medium ($OD_{600}$ = 0.001) was added to each well containing two antibiotics at the appropriate concentration in 100 μL of LB medium. After 24 h incubation at 37 °C, cultures were manually resuspended and $OD_{600}$ value was recorded on a varioskan LUX 3020-80316 to calculate the bacterial growth inhibition percentage of each well.

**Cell cultures.** HCT-116 (human colorectal carcinoma cell line), A549 (human lung adenocarcinoma cell line), HepG2 (human hepatocellular carcinoma cell line), HT1080 (human fibrosarcoma cell line), Wi38 (human fetal lung fibroblast cell line), Chang (human normal liver cell line), NIH3T3 (mouse fibroblast cell line), and Raw264.7 (mouse macrophage cell lines) were purchased from the ATCC and maintained at 37 °C in a humidified 5% $CO_2$ atmosphere. Cells were cultured in Dulbecco's modified Eagle's medium supplemented with 10% fetal bovine serum (FBS) and 1% antibiotics (Invitrogen, Carlsbad, CA).

**Cell cytotoxicity assay.** HCT-116, A549, HepG2, HT1080, Wi38, Chang, and NIH3T3 cells were treated with different concentration of the compounds **1–13** or 5-FU (5-fluorouracil) as indicated. After 24 h, 20 μL of CellTiter 96® Aqueous One solution reagent (Promega, Madison, WI, USA) was added and then read at 490 nm. Dose-response curves were plotted to determine half-maximal inhibitory concentrations ($IC_{50}$) for the compounds with SigmaPlot software.

**Nitric oxide assay.** After pre-incubation of Raw 264.7 cells in a presence of LPS (1 μg/mL) for 24 h with or without pretreatment with the compounds **1**–**13** for 1 h, the quantity of nitrite in the culture medium was measured using Griess reagent and the absorbance was measured at 540 nm. Fresh culture medium was used as a blank in every experiment.

**Structure elucidation of new metabolites**

Bacillimidazole G (**3**) was isolated as a brownish gum and its molecular formula was determined as $C_{28}H_{33}N_4^+$ based on a molecular ion at *m*/*z* 425.2701 $[M]^+$ in the HRESIMS (calcd for $C_{28}H_{33}N_4^+$, 425.2700). The $^1H$ and $^{13}C$ NMR spectra of **3** exhibited characteristic resonances for one indole moiety [$\delta_H$ 7.38 (1H, d, *J* = 8.1 Hz, H-7), 7.31 (1H, d, *J* = 8.1 Hz, H-4), 7.13 (1H, t, *J* = 7.5 Hz, H-6), 7.03 (1H, t, *J* = 7.5 Hz, H-5), and 6.96 (1H, s, H-2); $\delta_C$ 138.1 (C-8), 128.3 (C-9), 124.5 (C-2), 123.0 (C-6), 120.3 (C-5), 118.3 (C-4), 112.8 (C-7), and 110.7 (C-3)], four methylenes [$\delta_H$ 4.21 (2H, t, *J* = 6.8 Hz, H-11), 3.03 (2H, t, *J* = 6.8 Hz, H-10), 1.96 (2H, m, H-8′), and 1.17 (2H, m, H-9′); $\delta_C$ 47.5 (C-11), 26.4 (C-10), 25.3 (C-8′), and 21.6 (C-9′)], and two methyl groups [$\delta_H$ 2.21 (3H, s, H-6′) and 0.62 (3H, t, *J* = 7.3 Hz, H-10′); $\delta_C$ 13.7 (C-10′) and 8.5 (H-6′)], along with two quaternary carbons [$\delta_C$ 146.7 (C-2′) and 127.3 (C-4′)] (Table S2). Detailed inspection of the NMR and the HRESIMS revealed that **3** was a symetrical compound composed of two indole units. Furthermore, $^1H$ and $^{13}C$ NMR quite similar to those of bacillimidazole D (**1**) except for the presence of a propyl group, which indicated that bacillimidazole G (**3**) is an imidazolium-containing alkaloid.[3-4] The structure of **3** was determined on the basis of the 2D NMR analysis, including $^1H$-$^1H$ COSY, HSQC, and HMBC spectra. The linkage of the propyl group was identified to be at C-2′ based on the HMBC correlations of H-8′ ($\delta_H$ 1.96) and H-9′ ($\delta_H$ 1.17) with C-2′ ($\delta_C$ 146.7), respectively (Figure 2B). Thus, the structure of **3** was defined as 1,3-bis(2-(1*H*-indol-3-yl)ethyl)-4,5-dimethyl-2-propyl-1*H*-imidazol-3-ium.

Bacillimidazole H (**4**) was obtained as a brownish gum. The molecular formula was found to be $C_{29}H_{35}N_4^+$ based on a molecular ion at *m*/*z* 439.2868 $[M]^+$ in the HRESIMS (calcd for $C_{29}H_{35}N_4^+$, 439.2856). The $^1H$ and $^{13}C$ NMR spectra of **4** was comparable to those of **3**, with the major difference being the existence of an isobutyl group [$\delta_H$ 1.79 (2H, d, *J* = 7.9 Hz, H-

8′), 1.66 (1H, m, H-9′), and 0.65 (6H, d, $J = 6.6$ Hz, H-10′, 11′); $\delta_C$ 31.7 (C-8′), 29.6 (C-9′), and 22.3 (C-10′, 11′)] instead of the propyl group in **3** (Table S2). The position of the isobutyl group was confirmed to be at C-2′ based on the HMBC cross-peaks of $H_2$-8′ ($\delta_H$ 1.79) and H-9′ ($\delta_H$ 1.66) with C-2′ ($\delta_C$ 146.1), respectively (Figure S1). Thus, the structure of **4** was defined as 1,3-bis(2-(1*H*-indol-3-yl)ethyl)-2-isobutyl-4,5-dimethyl-1*H*-imidazol-3-ium.

Bacillimidazole I (**5**) was isolated as a brownish gum. The molecular formula was assigned as $C_{30}H_{37}N_4^+$ on the basis of a molecular ion at *m*/*z* 453.3072 $[M]^+$ in the HRESIMS (calcd for $C_{30}H_{37}N_4^+$, 453.3013). Analysis of the $^1$H and $^{13}$C NMR data of **5** revealed structural similarities with **3** with the major difference being the presence of an additional propyl group [$\delta_H$ 2.56 (2H, t, $J = 7.8$ Hz, H-8′), 1.56 (2H, q, $J = 7.8$ Hz, H-9′), and 0.94 (3H, t, $J = 7.3$ Hz, H-10′); $\delta_C$ 25.4 (C-8′), 23.8 (C-9′), and 14.0 (C-10′)] (Table S3). The propyl group was assigned to be connected to C-7′ based on the HMBC correlations of H-8′ ($\delta_H$ 2.56) and H-9′ ($\delta_H$ 1.56) with C-7′ ($\delta_C$ 131.3), respectively (Figure S1). Thus, the structure of **5** was defined as 1,3-bis(2-(1*H*-indol-3-yl)ethyl)-4-methyl-2,5-dipropyl-1*H*-imidazol-3-ium.

Bacillimidazole J (**8**) was obtained as a brownish gum and its molecular formula was confirmed as $C_{23}H_{34}N_3^+$ based on a molecular ion at *m*/*z* 352.2761 $[M]^+$ in the HRESIMS (calcd for $C_{23}H_{34}N_3^+$, 352.2747). The $^1$H and $^{13}$C NMR data of **8** similar to those of **3** except for the replacement of an indole moiety by an isopentyl group [$\delta_H$ 3.81 (2H, m, H-4″), 1.58 (1H, m, H-2″), 1.14 (2H, m, H-3″), and 0.94 (6H, d, $J = 6.7$ Hz, H-1″,5″); $\delta_C$ 44.6 (C-4″), 40.1 (C-3″), 27.3 (C-2″), and 22.8 (C-1″,5″)] (Table S3). The HMBC correlations of H-4″ ($\delta_H$ 3.81) with C-2′ ($\delta_C$ 146.6) and C-4′ ($\delta_C$ 127.2) confirmed the location of the isopentyl group (Figure S1). Thus, the structure of **8** was defined as 3-(2-(1*H*-indol-3-yl)ethyl)-1-isopentyl-4,5-dimethyl-2-propyl-1*H*-imidazol-3-ium.

Bacillindoles A and B (**9a** and **9b**), which were obtained as a brownish gum, possessed a molecular formula of $C_{14}H_{18}N_2O$ as assigned by a protonated molecular ion at *m*/*z* 231.1490 $[M + H]^+$ in the HRESIMS data (calcd for $C_{14}H_{19}N_2O$, 231.1492). The $^1H$ and $^{13}C$ NMR spectra of **9**, an enantiomeric mixture of **9a** and **9b**, displayed signals for one indole moiety [$\delta_H$ 7.56 (1H, d, *J* = 7.9 Hz, H-4), 7.37 (1H, d, *J* = 8.1 Hz, H-7), 7.19 (1H, s, H-2), 7.13 (1H, t, *J* = 7.5 Hz, H-6), and 7.05 (1H, t, *J* = 7.4 Hz, H-5); $\delta_C$ 138.3 (C-8), 128.1 (C-9), 124.2 (C-2), 122.8 (C-6), 120.1 (C-5), 118.8 (C-4), 112.6 (C-7), and 110.1 (C-3)], one 1-methyl-2-oxo-propyl group [$\delta_H$ 4.21 (1H, q, *J* = 7.3 Hz, H-3′), 2.26 (3H, s, H-1′), and 1.54 (3H, d, *J* = 7.3 Hz, H-4′); $\delta_C$ 205.1 (C-2′), 62.9 (C-3′), 26.4 (C-1′), and 14.2 (C-4′)], and two methylenes [$\delta_H$ 3.24 (2H, m, H-11) and 3.17 (2H, m, H-10); $\delta_C$ 47.4 (C-11) and 25.5 (C-10)] (Table S5). The planr structure of **9** was verified by the 2D NMR analysis, including $^1H$-$^1H$ COSY, HSQC, and HMBC spectra. The position of the 1-methyl-2-oxo-propyl group was determined to be connected at C-11 through secondary amine (NH) on the basis of the HMBC correlation of H-3′ ($\delta_H$ 4.21) with C-11 ($\delta_C$ 47.4) (Figure 2B). The planar structure of **9** was elucidated by 2D NMR. ECD analysis of **9** indicated that **9** was a racemic mixture, which was isolated into each enantiomer **9a** and **9b** using chiral-phase HPLC. As shown in Figure 2C, the experimental ECD spectrum of **9a** exhibited positive Cotton effects at 242, 297, and 307 nm and negative Cotton effects at 265 and 274 nm, which were in accordance with the calculated ECD spectrum of *R*. In contrast, **9b** displayed negative Cotton effects at 237 and 294 nm and positive Cotton effects at 254 and 268 nm, which well matched with calculated ECD spectrum of *S* (Figure 2C). Thus, the structure of **9a** and **9b** were defined as (*R*)-3-((2-(1*H*-indol-3-yl)ethyl)amino)butan-2-one and (*S*)-3-((2-(1*H*-indol-3-yl)ethyl)amino)butan-2-one, respectively.

Bacillindole C (**10**) was isolated as a brownish gum. Its molecular formula was determined as $C_{16}H_{22}N_2O$ based on a protonated molecular ion at *m*/*z* 259.1804 $[M + H]^+$ in the HRESIMS data (calcd for $C_{16}H_{23}N_2O$, 259.1805). The $^1H$ and $^{13}C$ NMR spectra of **10** was very similar to those of **9**, except for the presence of 1-methyl-2-oxo-pentyl group [$\delta_H$ 4.17 (1H, q, $J$ = 7.3 Hz, H-2′), 2.64 (1H, m, H-4′a), 2.47 (1H, m, H-4′b), 1.63 (2H, m, H-5′), 1.52 (3H, d, $J$ = 7.3 Hz, H-1′), and 0.94 (3H, t, $J$ = 7.4 Hz, H-6′); $\delta_C$ 207.2 (C-3′), 62.4 (C-2′), 41.6 (C-4′), 17.6 (C-5′), 14.3 (C-1′), and 13.8 (C-6′)] rather than the 1-methyl-2-oxo-propyl group in **9** (Table S4). The 3-hexanone moiety was linked at C-11 through secondary amine (NH) based on the HMBC cross-peak from H-2′ ($\delta_H$ 4.17) to C-11 ($\delta_C$ 47.4) (Figure S1). No observed Cotton effect in **10** from ECD analysis (data not shown) indicated that compound **10** was a racemic mixture.

Bacillipyrrole B (**12**) was purified as a brownish gum with a confirmed molecular formula of $C_{18}H_{20}N_2O$ based on a protonated molecular ion at *m/z* 281.1660 $[M + H]^+$ in the HRESIMS data (calcd for $C_{18}H_{21}N_2O$, 281.1654). The $^1H$ and $^{13}C$ NMR spectra of **12** displayed characteristic signals for one indole moiety [$\delta_H$ 7.52 (1H, d, $J$ = 7.9 Hz, H-7), 7.31 (1H, d, $J$ = 8.1 Hz, H-4), 7.07 (1H, t, $J$ = 8.1 Hz, H-6), 6.98 (1H, t, $J$ = 7.6 Hz, H-5), and 6.90 (1H, s, H-2); $\delta_C$ 138.2 (C-8), 128.1 (C-9), 123.9 (C-2), 122.4 (C-6), 119.7 (C-5), 119.4 (C-7), 112.8 (C-3), and 112.1 (C-4)], one pyrrole moiety [$\delta_H$ 6.95 (1H, s, H-3′); $\delta_C$ 139.2 (C-5′), 129.3 (C-2′), 123.8 (C-3′), and 118.3 (C-4′)], two methylene groups [$\delta_H$ 4.51 (2H, t, $J$ = 7.3 Hz, H-11) and 3.05 (2H, t, $J$ = 7.3 Hz, H-10); $\delta_C$ 48.0 (C-11) and 28.1 (C-10)], an acetyl group [$\delta_H$ 2.40 (3H, s, H-9′); $\delta_C$ 188.9 (C-8′) and 27.1 (C-9′)], and two methyl groups [$\delta_H$ 1.95 (3H, s, H-7′) and 1.86 (3H, s, H-6′); $\delta_C$ 11.4 (C-7′) and 9.9 (C-6′)] (Table S6). The structure of **12** was verified based on the 2D NMR analysis, including $^1H$-$^1H$ COSY, HSQC, and HMBC spectra. The connectivity between the indole moiety and the pyrrole moiety was identified to be linked through two methylenes, which was supported both the $^1H$-$^1H$ COSY cross-peak from H-10 ($\delta_H$ 3.05) to H-11 ($\delta_H$ 4.51) and the HMBC correlations of H-10 ($\delta_H$ 3.05) with C-2 ($\delta_C$ 123.9) and C-9 ($\delta_C$ 128.1) and of H-11 ($\delta_H$ 4.51) with C-2′ ($\delta_C$ 129.3) and C-5′ ($\delta_C$ 139.2) (Figure 2B).

The acetyl group was determined to be at C-2′ based on the HMBC cross-peak from H-9′ ($\delta_H$ 2.40) to C-2′ ($\delta_C$ 129.3) (Figure 2B). Furthermore, the position of the two methyl groups were identified to be connected to C-4′ and C-5′ by the HMBC correlations from H-6′ ($\delta_H$ 1.86) to C-5′ ($\delta_C$ 139.2) and from H-7′ ($\delta_H$ 1.95) to C-4′ ($\delta_C$ 118.3), respectively (Figure 2B). Thus, the structure of **12** was defined as 1-(1-(2-(1*H*-indol-3-yl)ethyl)-4,5-dimethyl-1*H*-pyrrol-2-yl)ethan-1-one.

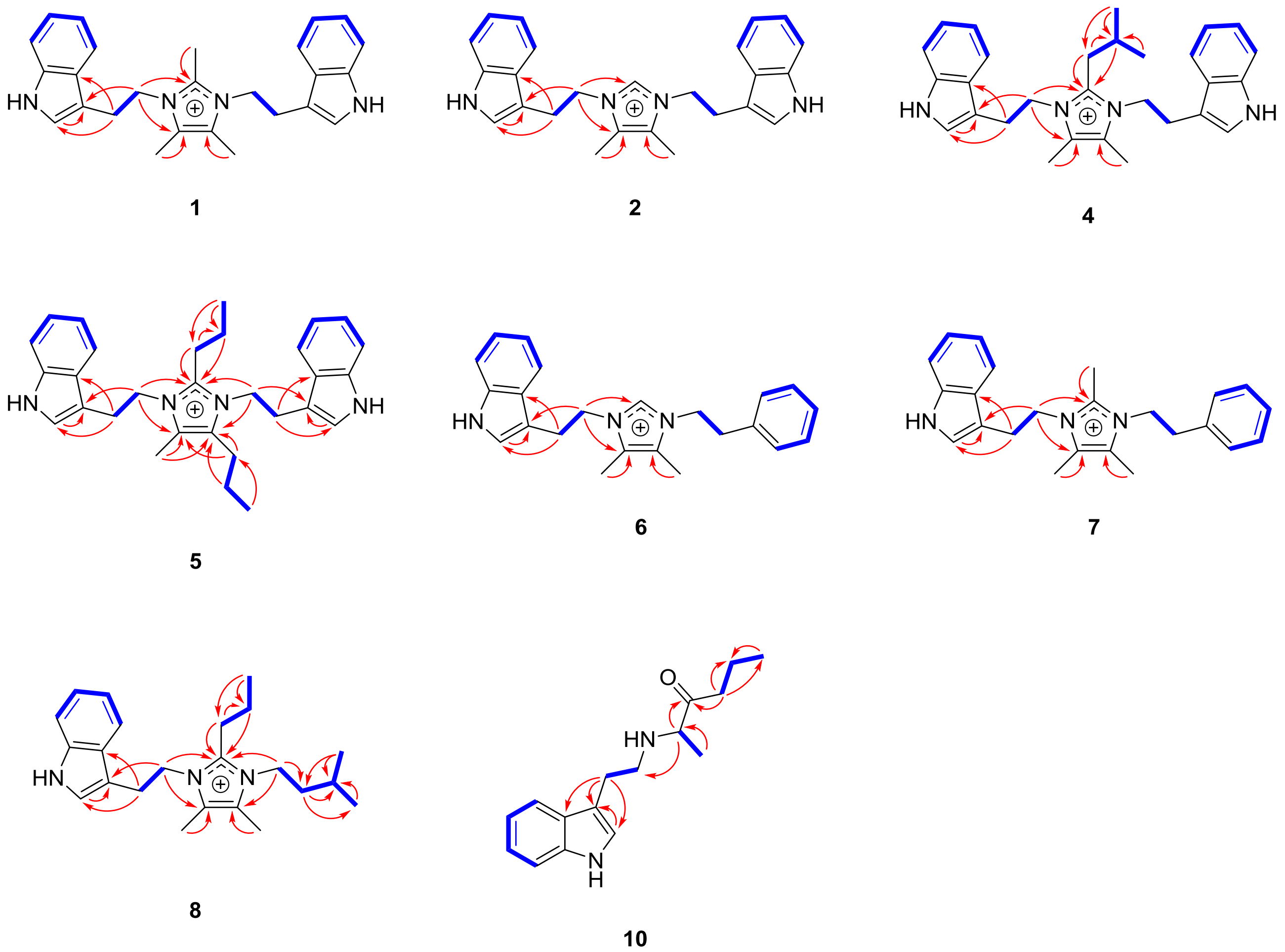

**Figure S1**. 2D NMR correlations of compounds **1**, **2**, **4-8**, and **10**.

**Table S1.** $^{1}$H and $^{13}$C NMR data for compounds **1** and **2** in methanol-$d_4$

| Position | **1** | | **2** | |
|---|---|---|---|---|
| | $\delta_C$ | $\delta_H$ [mult. ($J$ in Hz)] | $\delta_C$ | $\delta_H$ [mult. ($J$ in Hz)] |
| 2/2″ | 124.2 | 6.95, s | 124.4 | 6.93, s |
| 3/3″ | 110.8 | | 110.5 | |
| 4/4″ | 118.2 | 7.21, d (7.9) | 118.4 | 7.29, d (7.9) |
| 5/5″ | 120.2 | 7.00, t (7.4) | 120.1 | 7.00, t (7.0) |
| 6/6″ | 122.9 | 7.11, t (7.5) | 122.8 | 7.12, t (7.0) |
| 7/7″ | 112.7 | 7.36, d (8.1) | 112.6 | 7.36, d (8.1) |
| 8/8″ | 138.0 | | 138.1 | |
| 9/9″ | 128.4 | | 128.2 | |
| 10/10″ | 26.0 | 3.00, t (6.7) | 26.8 | 3.04, t (6.8) |
| 11/11″ | 47.5 | 4.16, t (6.7) | 49.3 | 4.22, t (7.9) |
| 2′ | 143.9 | | 135.4 | 8.10, s |
| 4′/5′ | 127.0 | | 128.3 | |
| 6′/7′ | 8.3 | 2.15, s | 7.9 | 2.06, s |
| 8′ | 9.3 | 1.61, s | | |

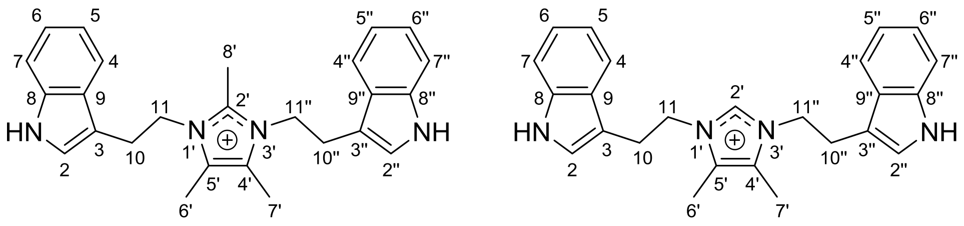

**1** **2**

**Table S2.** $^1$H and $^{13}$C NMR data for compounds **3** and **4** in methanol-$d_4$

| Position | **3** | | **4** | |
|---|---|---|---|---|
| | $\delta_C$ | $\delta_H$ [mult. ($J$ in Hz)] | $\delta_C$ | $\delta_H$ [mult. ($J$ in Hz)] |
| 2/2″ | 124.5 | 6.96, s | 124.2 | 6.96, s |
| 3/3″ | 110.7 | | 110.7 | |
| 4/4″ | 118.3 | 7.31, d (8.0) | 118.4 | 7.30, d (7.8) |
| 5/5″ | 120.3 | 7.03, t (7.4) | 120.4 | 7.05, t (7.4) |
| 6/6″ | 123.0 | 7.13, t (7.5) | 123.0 | 7.14, t (7.2) |
| 7/7″ | 112.8 | 7.38, d (8.1) | 112.9 | 7.39, d (8.2) |
| 8/8″ | 138.1 | | 138.1 | |
| 9/9″ | 128.3 | | 128.4 | |
| 10/10″ | 26.4 | 3.03, t (7.8) | 26.5 | 3.03, t (6.8) |
| 11/11″ | 47.5 | 4.21, t (6.8) | 47.4 | 4.21, t (6.8) |
| 2′ | 146.7 | | 146.1 | |
| 4′/5′ | 127.3 | | 127.3 | |
| 6′/7′ | 8.5 | 2.21, s | 8.5 | 2.27, s |
| 8′ | 25.3 | 1.96, m | 31.7 | 1.79, d (7.9) |
| 9′ | 21.6 | 1.17, m | 29.6 | 1.66, m |
| 10′ | 13.7 | 0.62, t (7.3) | 22.3 | 0.65, d (6.6) |
| 11′ | | | 22.3 | 0.65, d (6.6) |

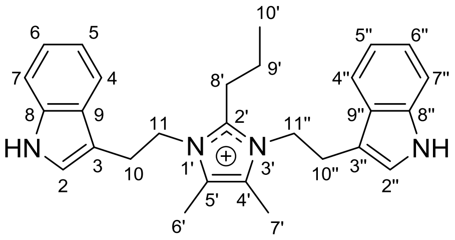

**3**

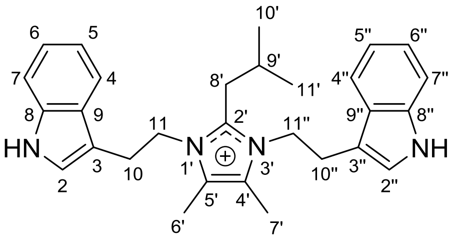

**4**

**Table S3.** $^{1}$H and $^{13}$C NMR data for compounds **5** and **8** in methanol-$d_4$

| Position | **5** | | **8** | |
|---|---|---|---|---|
| | $\delta_C$ | $\delta_H$ [mult. ($J$ in Hz)] | $\delta_C$ | $\delta_H$ [mult. ($J$ in Hz)] |
| 2 | 124.5 | 7.00, s | 124.7 | 7.05, s |
| 3 | 110.8 | | 111.2 | |
| 4 | 118.2 | 7.26, d (8.0) | 118.0 | 7.10, d (8.4) |
| 5 | 120.5 | 7.01, t (7.4) | 120.4 | 6.95, t (7.5) |
| 6 | 123.0 | 7.12, t (7.0) | 123.2 | 7.10, t (7.5) |
| 7 | 129.4 | 7.28, d (8.0) | 112.7 | 7.36, d (8.4) |
| 8 | 138.2 | | 138.2 | |
| 9 | 128.4 | | 129.0 | |
| 10 | 26.3 | 3.08, t (6.9) | 26.5 | 3.23, t (6.1) |
| 11 | 47.6 | 4.24, t (6.6) | 48.2 | 4.36, m |
| 2′ | 147.0 | | 146.5 | |
| 4′ | 131.3 | | 127.2 | |
| 5′ | 128.1 | | 127.5 | |
| 6′ | 8.7 | 2.20, s | 8.4 | 2.24, s |
| 7′ | 25.4 | 2.56, t (7.8) | 8.2 | 2.29, s |
| 8′ | 23.8 | 1.56, q (7.5) | 25.5 | 2.13, t (8.2) |
| 9′ | 25.4 | 0.94, t (7.3) | 22.0 | 1.37, m |
| 10′ | 25.4 | 2.03, m | 14.0 | 0.81, t (7.3) |
| 11′ | 21.8 | 1.24, m | | |
| 12′ | 14.0 | 0.67, t (7.3) | | |

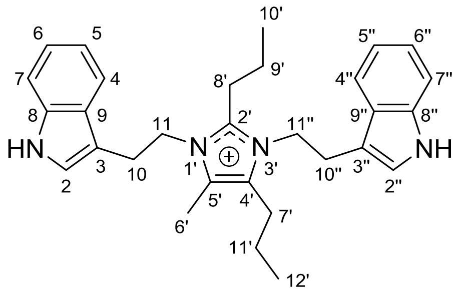

**5**

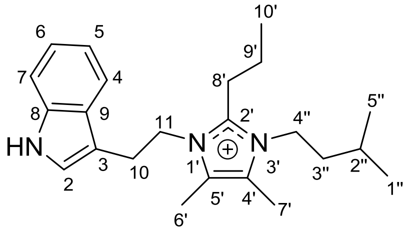

**8**

**Table S4.** $^{1}H$ and $^{13}C$ NMR data for compounds **6** and **7** in methanol-$d_4$

| Position | **6** | | **7** | |
|---|---|---|---|---|
| | $\delta_C$ | $\delta_H$ [mult. ($J$ in Hz)] | $\delta_C$ | $\delta_H$ [mult. ($J$ in Hz)] |
| 2 | 124.5 | 6.98, s | 124.6 | 6.98, s |
| 3 | 110.4 | | 110.5 | |
| 4 | 118.4 | 7.29, d (7.9) | 118.2 | 7.21, d (7.4) |
| 5 | 120.2 | 7.00, t (7.5) | 120.3 | 6.98, t (7.4) |
| 6 | 122.9 | 7.10, t (7.5) | 122.9 | 7.09, t (7.6) |
| 7 | 112.7 | 7.35, d (8.3) | 112.7 | 7.35, d (8.2) |
| 8 | 138.1 | | 138.0 | |
| 9 | 128.4 | | 128.5 | |
| 10 | 26.8 | 3.18, t (6.5) | 26.0 | 3.15, t (6.4) |
| 11 | 48.8 | 4.33, t (6.5) | 47.8 | 4.29, t (6.4) |
| 2′ | 135.4 | 8.21, s | 144.0 | |
| 4′ | 128.2 | | 126.96 | |
| 5′ | 128.4 | | 127.01 | |
| 6′ | 7.9 | 1.99, s | 8.2 | 2.05, s |
| 7′ | 8.0 | 2.07, s | 8.3 | 2.13, s |
| 8′ | | | 9.4 | 1.75, s |
| 1″ | 137.8 | | 138.1 | |
| 2″/6″ | 129.9 | 6.99, d (8.0) | 130.0 | 7.01, m |
| 3″/5″ | 130.0 | 7.24, t (8.0) | 130.1 | 7.27, q (6.4) |
| 4″ | 128.3 | 7.22, t (7.1) | 129.5 | 7.24, d (7.7) |
| 7″ | 37.0 | 2.78, t (7.2) | 36.3 | 2.75,t (7.1) |
| 8″ | 49.0 | 4.13, t (7.2) | 47.6 | 4.09, t (7.1) |

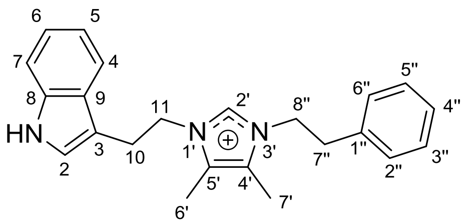

**6**

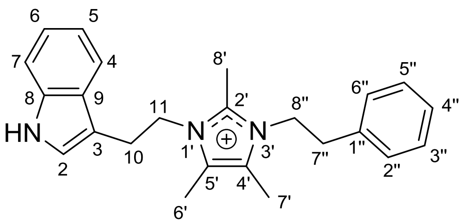

**7**

**Table S5.** $^{1}H$ and $^{13}C$ NMR data for compounds **9** and **10** in methanol-$d_4$

| Position | **9** | | **10** | |
|---|---|---|---|---|
| | $\delta_C$ | $\delta_H$ [mult. ($J$ in | $\delta_C$ | $\delta_H$ [mult. ($J$ in |
| 2 | 124.2 | 7.19, s | 124.2 | 7.19, s |
| 3 | 110.1 | | 110.1 | |
| 4 | 118.8 | 7.56, d (7.9) | 118.8 | 7.56, d (7.9) |
| 5 | 120.1 | 7.05, t (7.4) | 120.1 | 7.05, t (7.4) |
| 6 | 122.8 | 7.13, t (7.5) | 122.8 | 7.13, t (7.5) |
| 7 | 112.6 | 7.37, d (8.1) | 112.6 | 7.37, d (8.1) |
| 8 | 138.3 | | 138.3 | |
| 9 | 128.1 | | 128.1 | |
| 10 | 25.5 | 3.17, m | 23.5 | 3.18, m |
| 11 | 47.4 | 3.24, m | 47.4 | 3.23, m |
| 1′ | 26.4 | 1.54, d (7.3) | 14.3 | 1.52, d (7.3) |
| 2′ | 62.9 | 4.21, q (7.3) | 62.4 | 4.17, q (7.3) |
| 3′ | 205.1 | | 207.2 | |
| 4′a | 26.4 | 2.26, s | 41.6 | 2.64, m |
| 4′b | | | 41.6 | 2.47, m |
| 5′ | | | 17.6 | 1.63, m |
| 6′ | | | 13.8 | 0.94, t (7.4) |

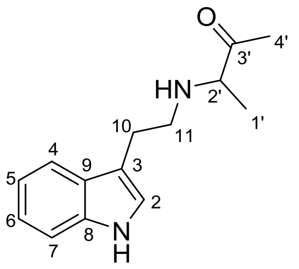

**9**

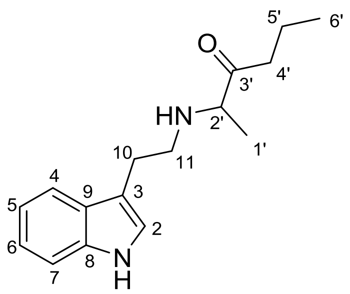

**10**

**Table S6.** $^{1}$H and $^{13}$C NMR data for compound **12** in methanol-$d_4$

| Position | **12** | |
|---|---|---|
| | $\delta_C$ | $\delta_H$ [mult. ($J$ in Hz)] |
| 2 | 123.9 | 6.90, s |
| 3 | 112.8 | |
| 4 | 112.1 | 7.31, d (8.1) |
| 5 | 119.7 | 6.98, t (7.6) |
| 6 | 122.4 | 7.07, t (8.1) |
| 7 | 119.4 | 7.52, d (7.9) |
| 8 | 138.2 | |
| 9 | 128.1 | |
| 10 | 28.1 | 3.05, t (7.4) |
| 11 | 48.0 | 4.51, t (7.3) |
| 2′ | 129.3 | |
| 4′ | 118.3 | |
| 5′ | 139.2 | |
| 6′ | 9.9 | 1.86, s |
| 7′ | 11.4 | 1.95, s |
| 8′ | 188.9 | |
| 9′ | 27.1 | 2.40, s |

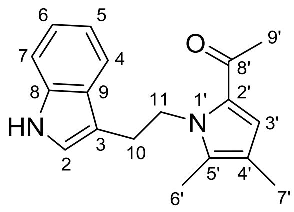

**12**

**Table S7.** Gibbs free energies and Boltzmann distribution of conformers **9*R***

| Conformers | B3LYP/6-31G(d) Gibbs free energy (298.15 K) | | |
|---|---|---|---|
| | G (Hartree) | ΔG (kcal/mol) | Boltmann distribution (%) |
| 9*R*-1 | -729.0559552 | 0.00 | 16.7 |
| 9*R*-2 | -729.0560987 | -0.09 | 19.5 |
| 9*R*-3 | -729.0554104 | 0.34 | 9.4 |
| 9*R*-4 | -729.0541058 | 1.16 | 2.4 |
| 9*R*-5 | -729.0539888 | 1.23 | 2.1 |
| 9*R*-6 | -729.0561272 | -0.11 | 20.1 |
| 9*R*-7 | -729.0538439 | 1.32 | 1.8 |
| 9*R*-8 | -729.0550258 | 0.58 | 6.3 |
| 9*R*-9 | -729.0537172 | 1.40 | 1.6 |
| 9*R*-10 | -729.0538007 | 1.35 | 1.7 |
| 9*R*-11 | -729.0549249 | 0.65 | 5.6 |
| 9*R*-12 | -729.0525684 | 2.13 | 0.5 |
| 9*R*-13 | -729.0556336 | 0.20 | 11.9 |
| 9*R*-14 | -729.0525864 | 2.11 | 0.5 |

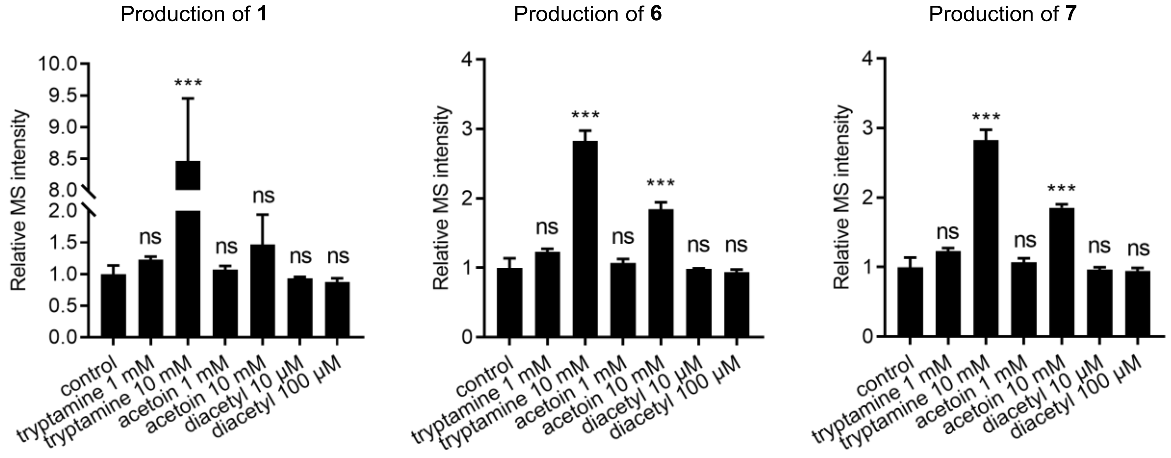

**Figure S2.** Individual tryptamine, acetoin, and diacetyl feeding experiments results indicating tryptamine and acetoin as dominant precursor of **1**, **6**, and **7**.

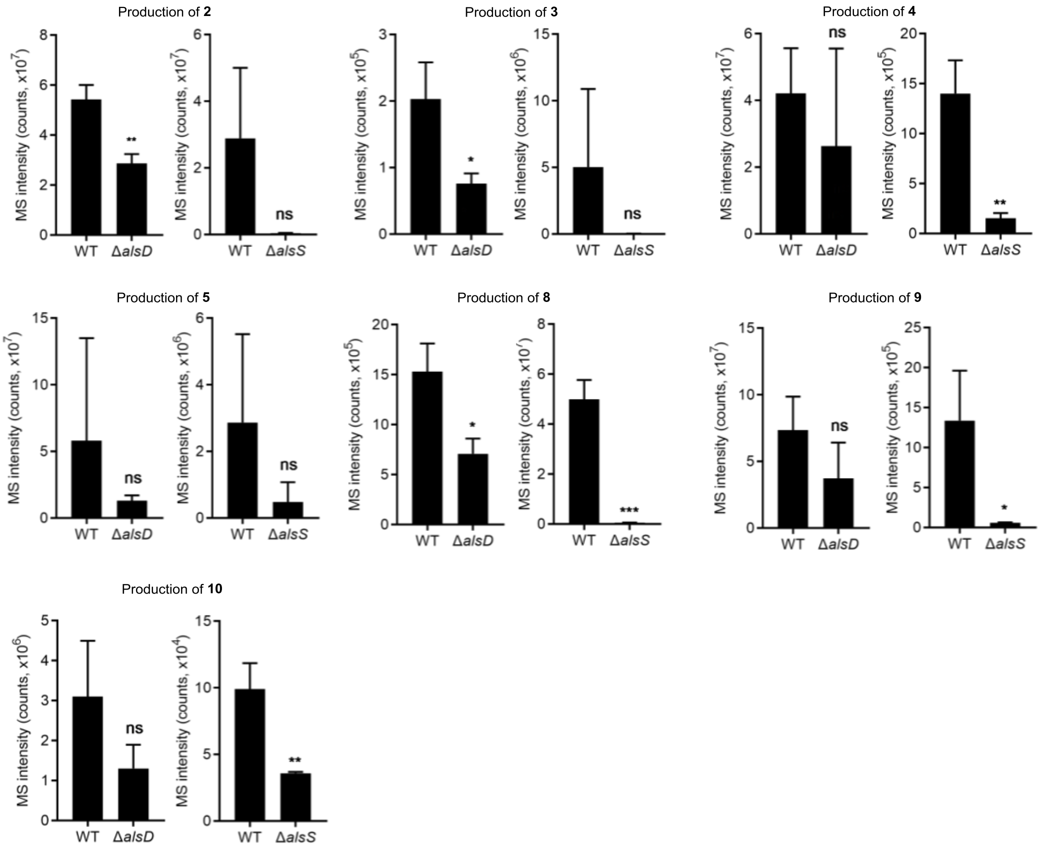

**Figure S3.** Comparison of production levels of **2–5** and **8–10** between WT *B. subtilis* HS3 and its *alsD* mutant strain (left) and between WT *B. subtilis* HB1000 and its *alsS* mutant strain (right).

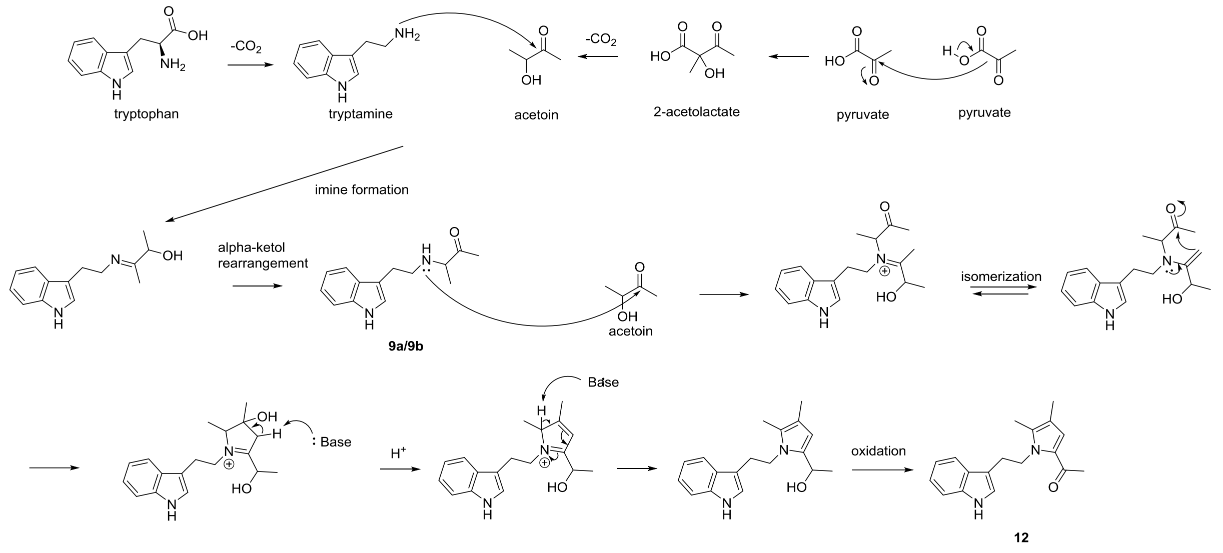

**Figure S4**. Plausible biosynthetic pathway of bacillipyrrole B (**12**).

**Table S8.** Cytotoxicity of compounds **1–13** against cancer and normal cell lines.

| Comp. | $IC_{50}$ ($\mu$M)[a] | | | | | | |
|---|---|---|---|---|---|---|---|
| | Cancer cell lines | | | | Normal cell lines | | |
| | HCT-116 | A549 | HepG2 | HT1080 | Wi38 | Chang | NIH-3T3 |
| 5 | 46.53 ± 0.80 | 69.09 ± 1.98 | 68.84 ± 1.22 | 29.22 ± 0.75 | > 100 | 59.37 ± 1.27 | 87.88 ± 3.27 |
| 1–4, 6–13 | > 100 | > 100 | > 100 | > 100 | > 100 | > 100 | > 100 |
| 5-FU[b] | 39.82 ± 2.78 | 46.9 ± 1.90 | > 100 | > 100 | > 100 | 34.93 ± 3.75 | 0.18 ± 0.013 |

[a] Average IC50 values are shown. Each compound was tested at six different concentrations, and each drug dilution was repeated three times. Cells treated with DMSO (equivalent volume) were used as a vehicle control. Except the $IC_{50}$ of **5**, them of other metabolites were exceeded 100 μM ( > 100).

[b] 5-FU (5-fluorouracil) was used as a positive control

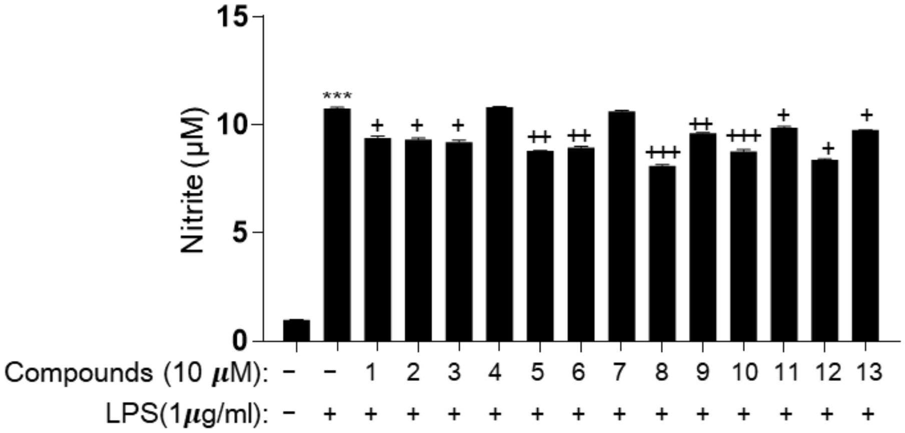

**Figure S5**. Effect of compound **1–13** on NO production in RAW 264.7 macrophage cells.

RAW 264.7 cells were treated with LPS (1 μg/mL) alone or LPS + compounds (10 μM) for 24 h. Griess reagent was used to quantify the NO amount, and values are presented as the mean ± SD ($n$ = 5). *** $p < 0.001$, vehicle vs. LPS-treated group; + $p < 0.05$; ++ $p < 0.01$; +++ $p <$ 0.001, LPS vs. LPS + compound-treated group.

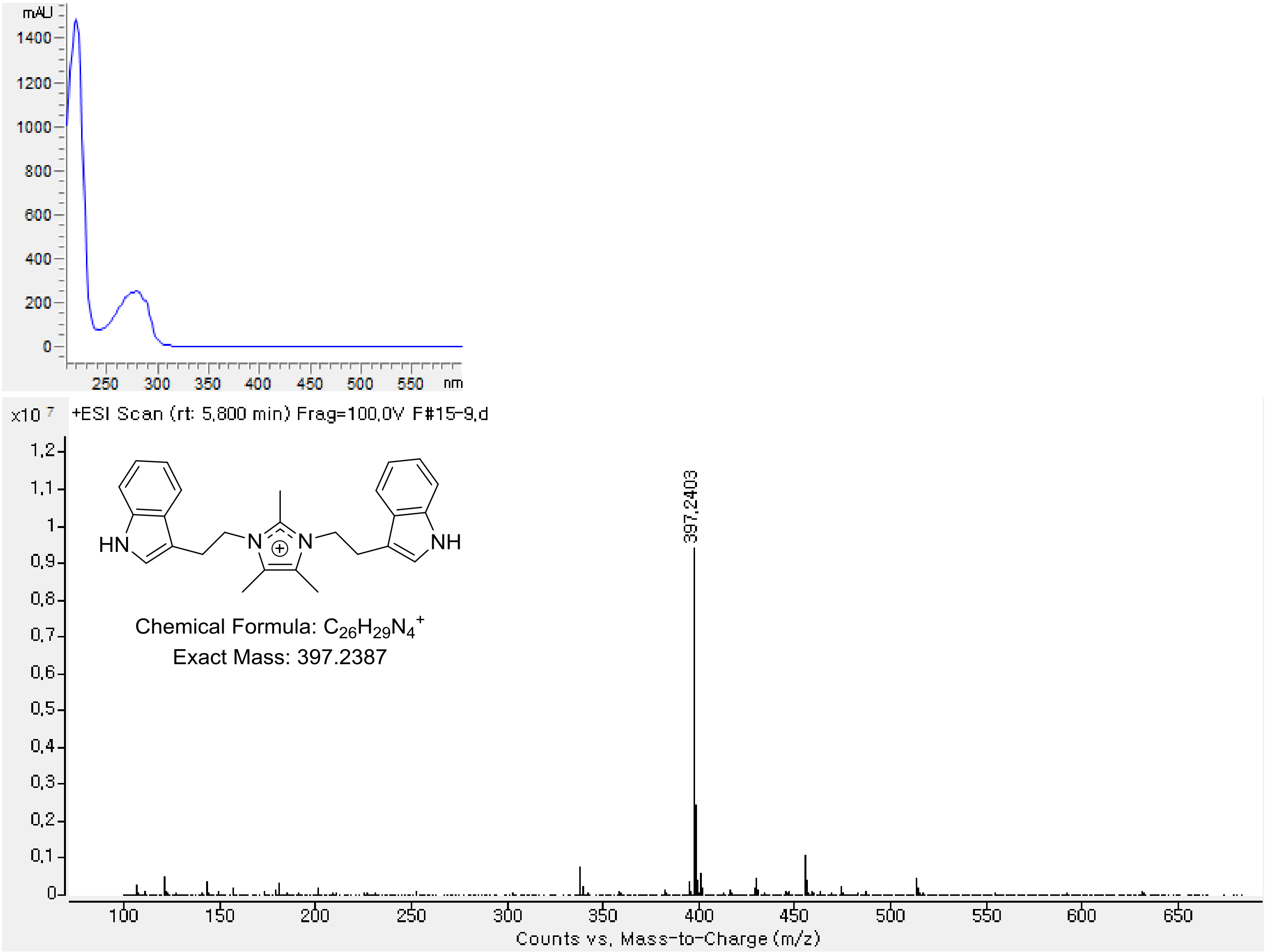

**Figure S5**. UV-Vis and HRESIMS of **1**.

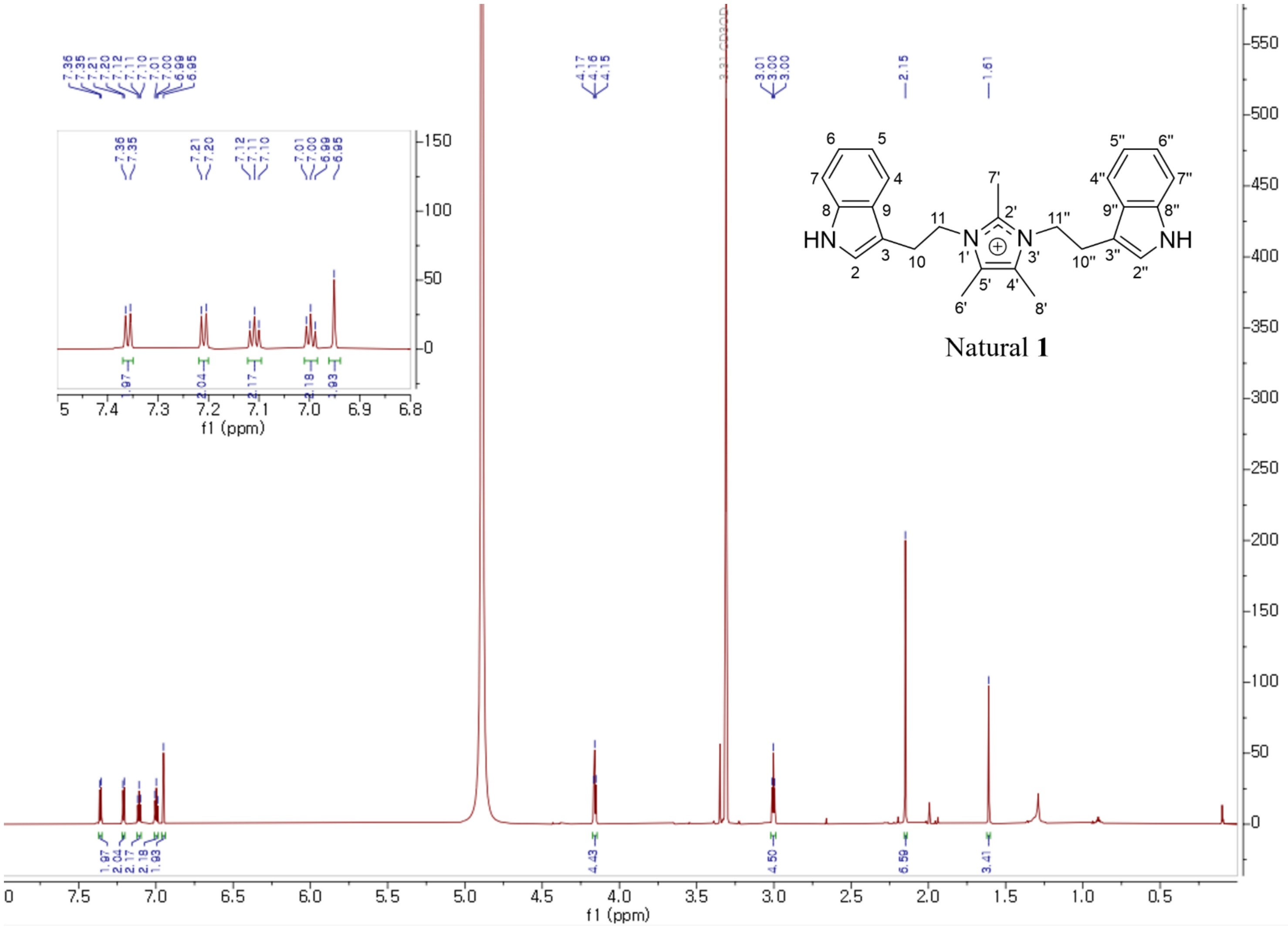

**Figure S6**. The $^{1}$H NMR spectrum of synthetic **1** in methanol-$d_4$.

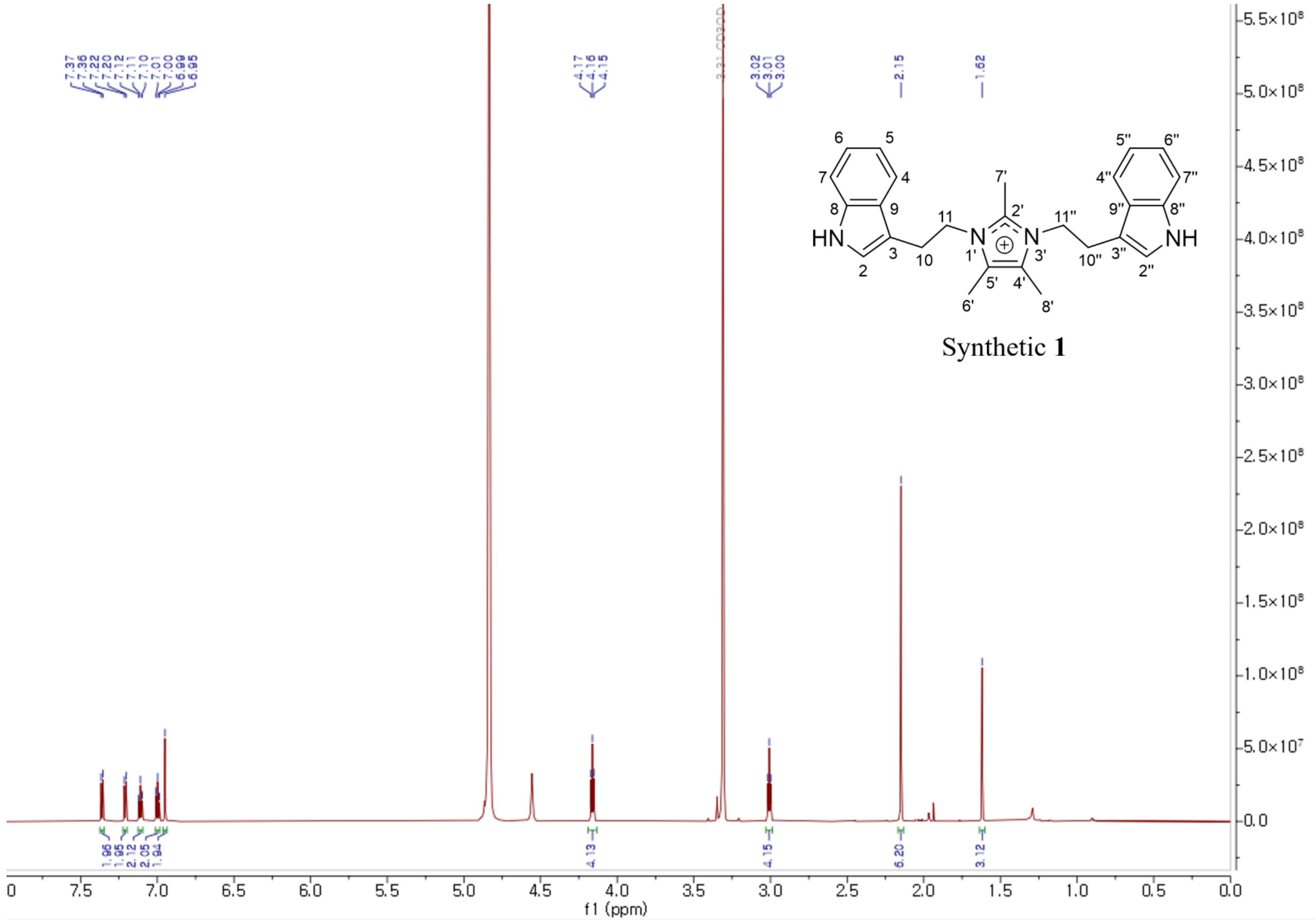

**Figure S6**. The $^{1}$H NMR spectrum of synthetic **1** in methanol-$d_4$.

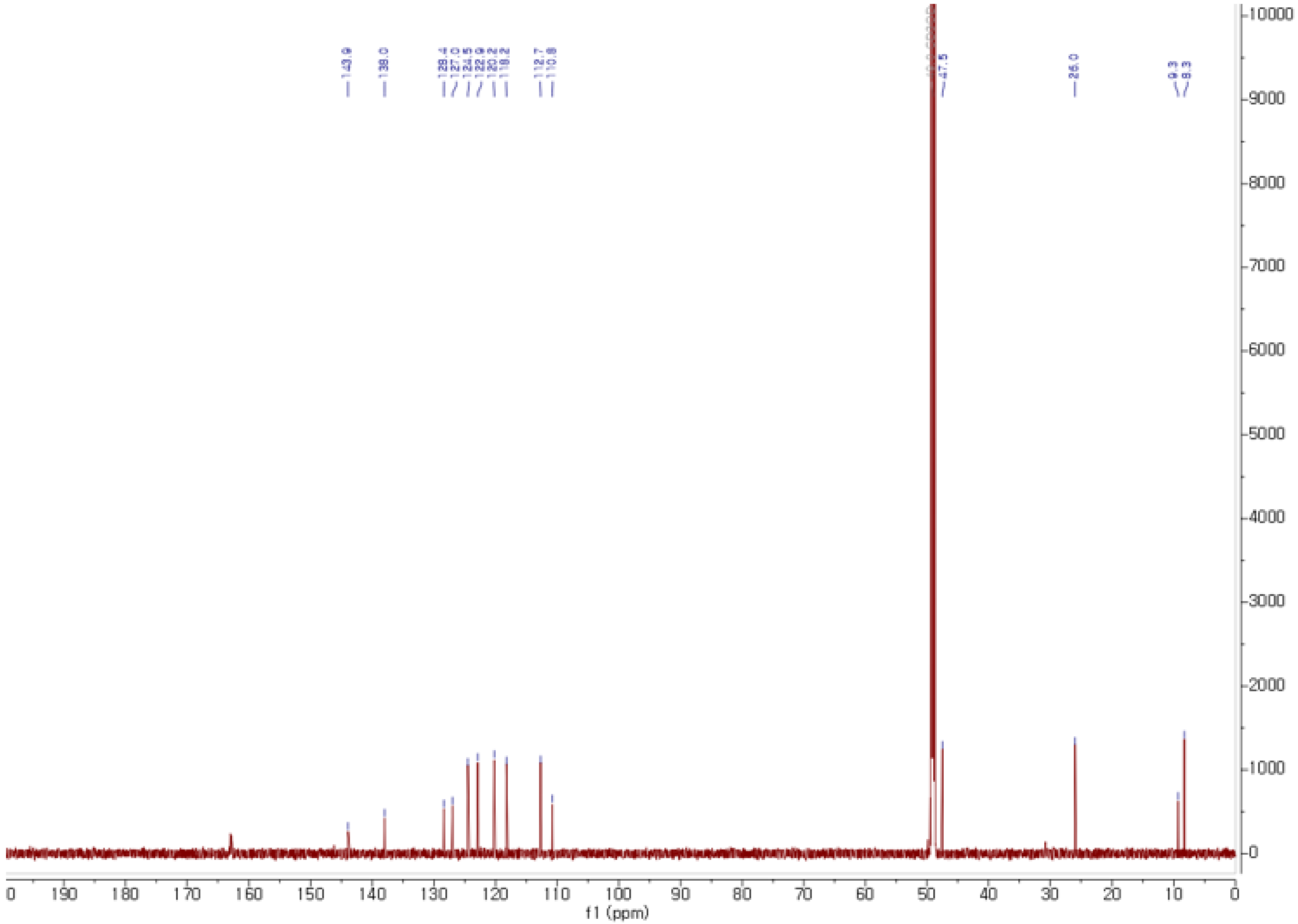

**Figure S7**. The $^{13}C$ NMR spectrum of **1** in methanol-$d_4$.

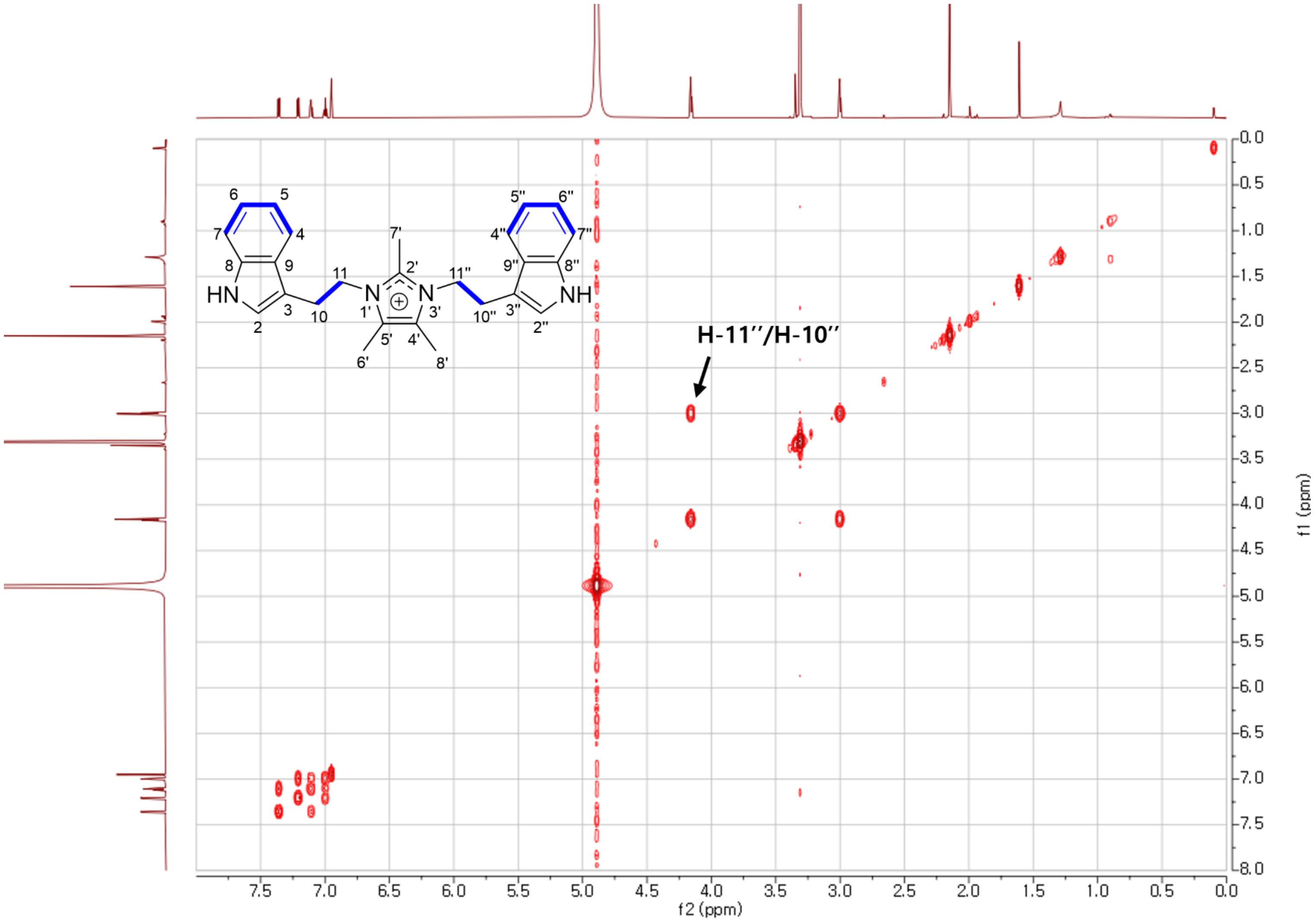

**Figure S8**. The COSY spectrum of **1** in methanol-$d_4$.

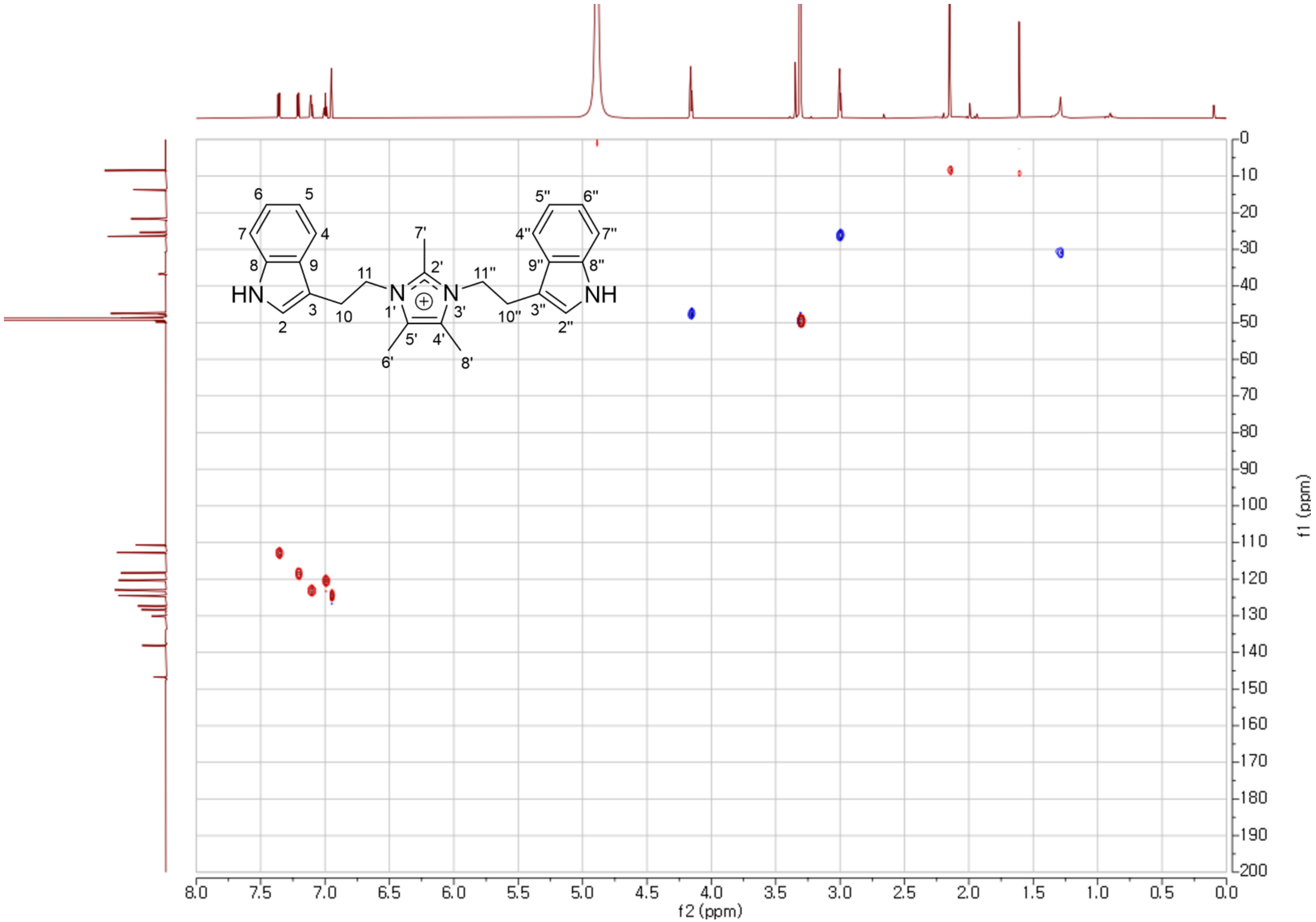

**Figure S9**. The HSQC spectrum of **1** in methanol-$d_4$.

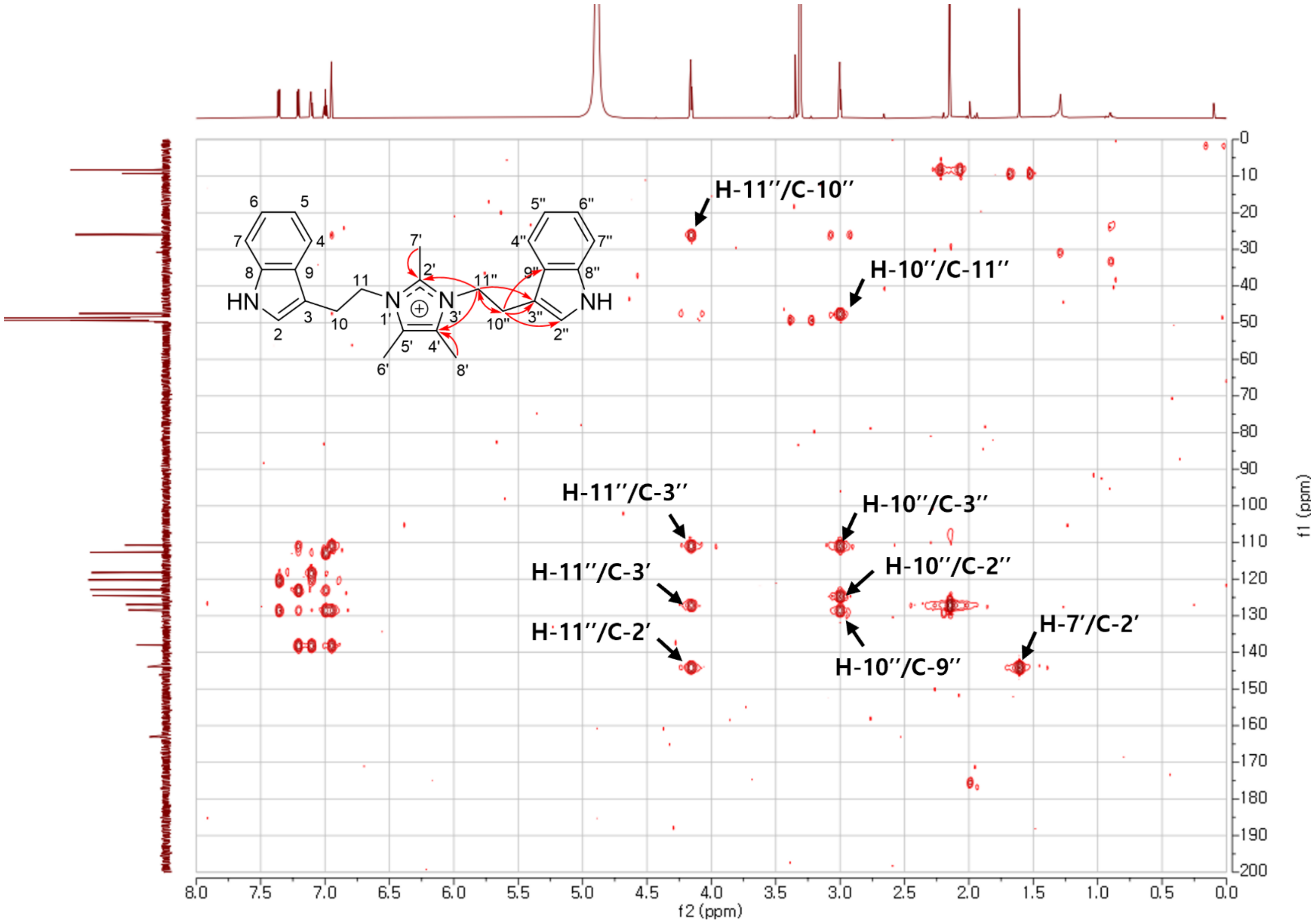

**Figure S10**. The HMBC spectrum of **1** in methanol-$d_4$.

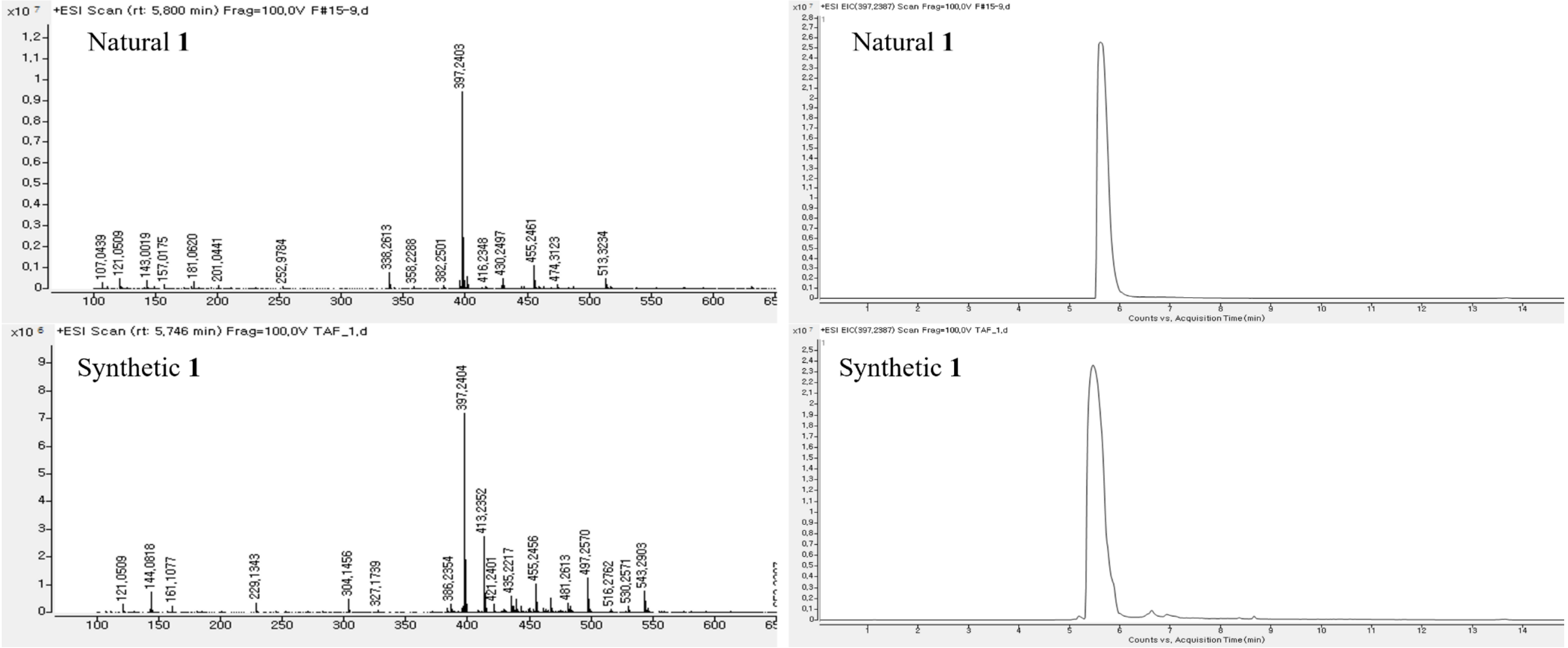

**Figure S11**. HRESIMS and HPLC chromatogram of **1**.

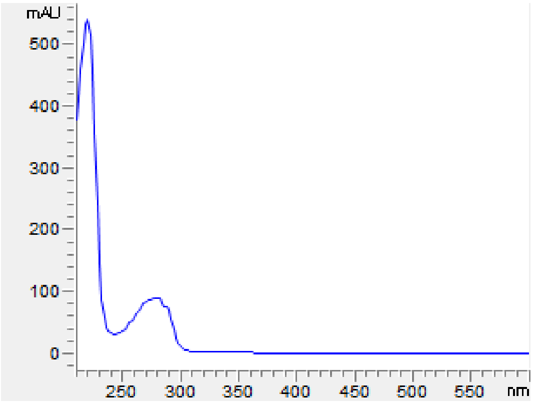

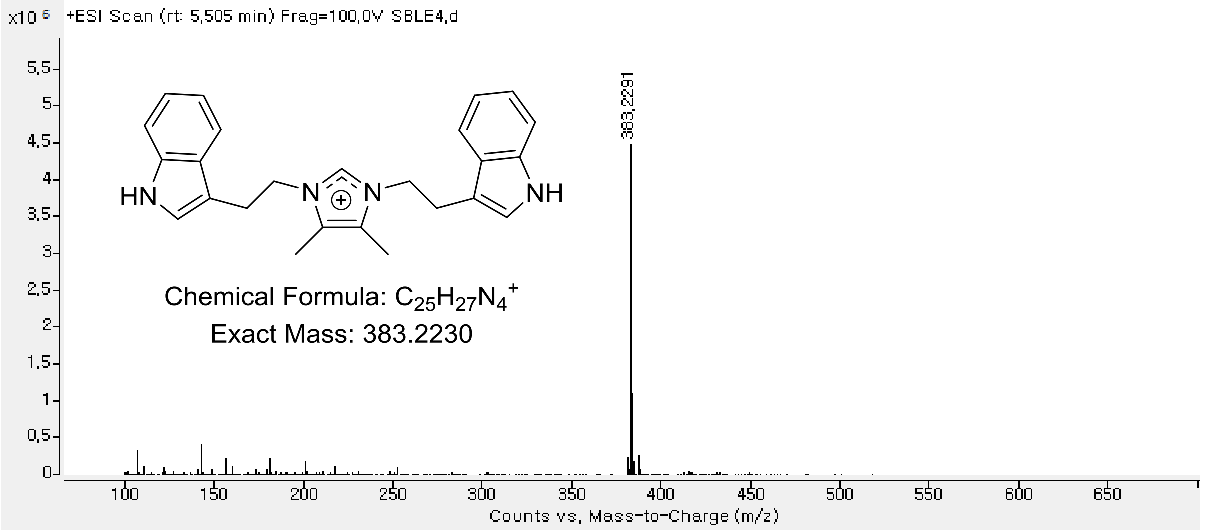

**Figure S12**. UV-Vis and HRESIMS spectrum of **2**.

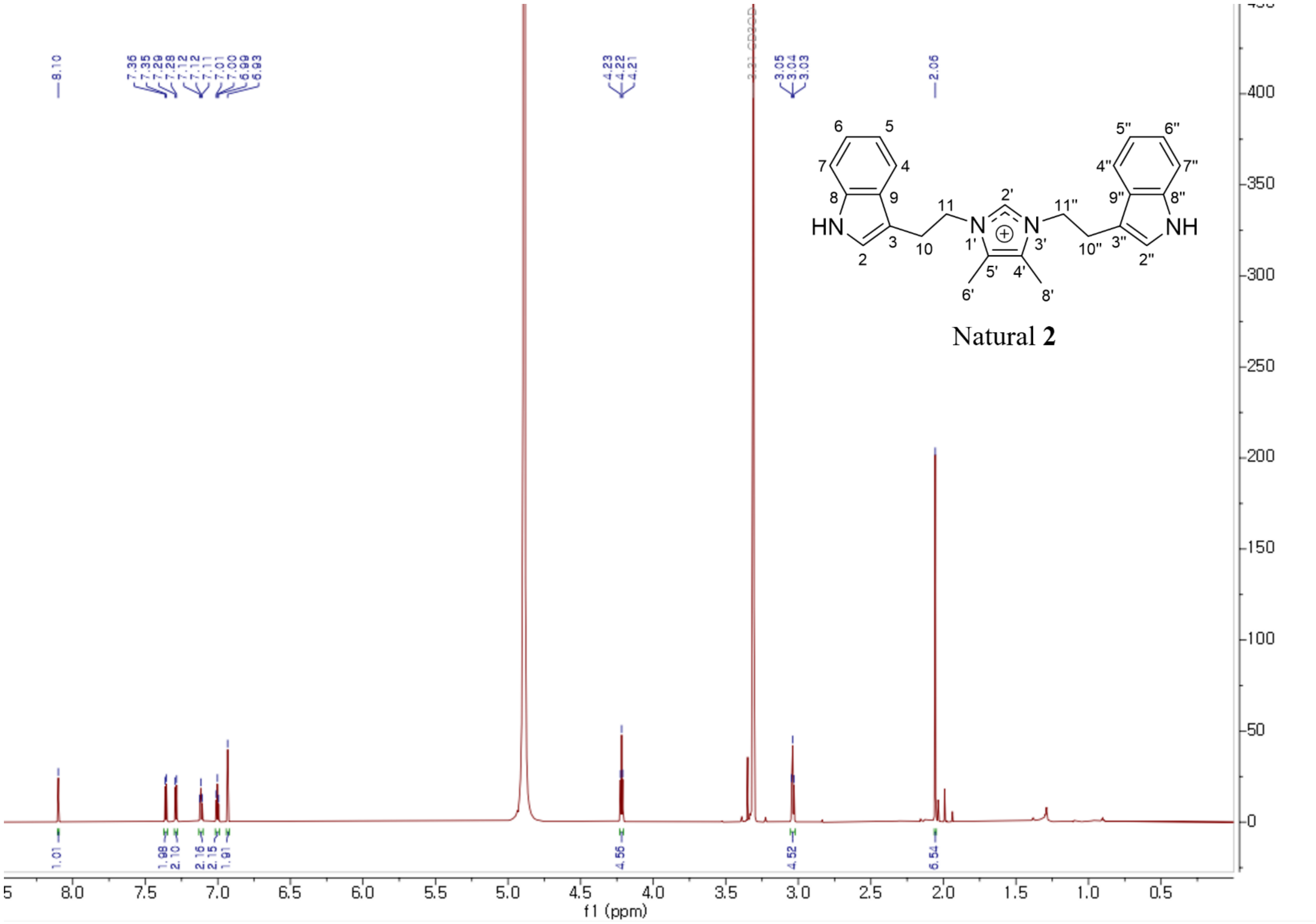

**Figure S13**. The $^{1}$H NMR spectrum of natural **2** in methanol-$d_4$.

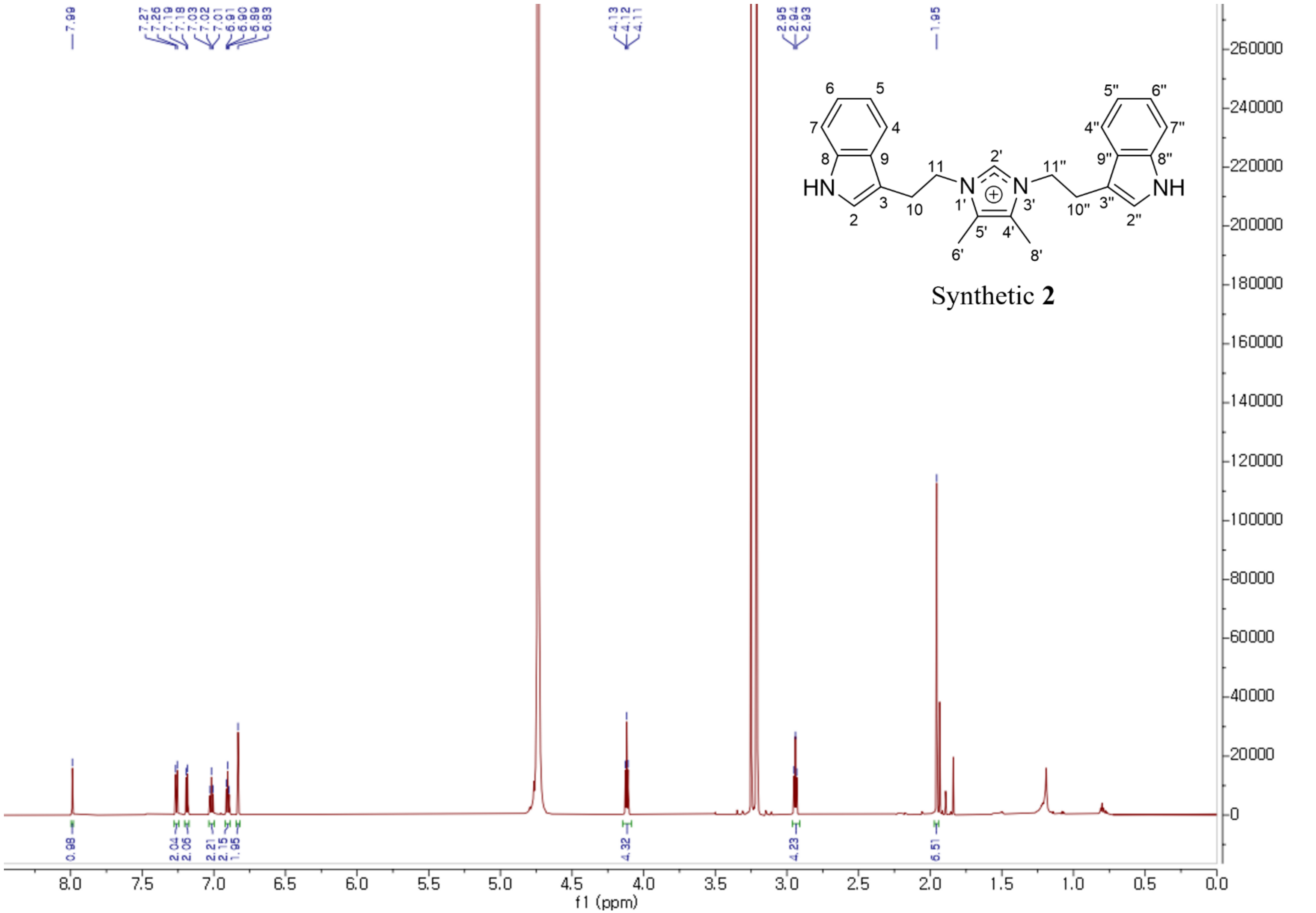

**Figure S14**. The [1]H NMR spectrum of synthetic **2** in methanol-$d_4$.

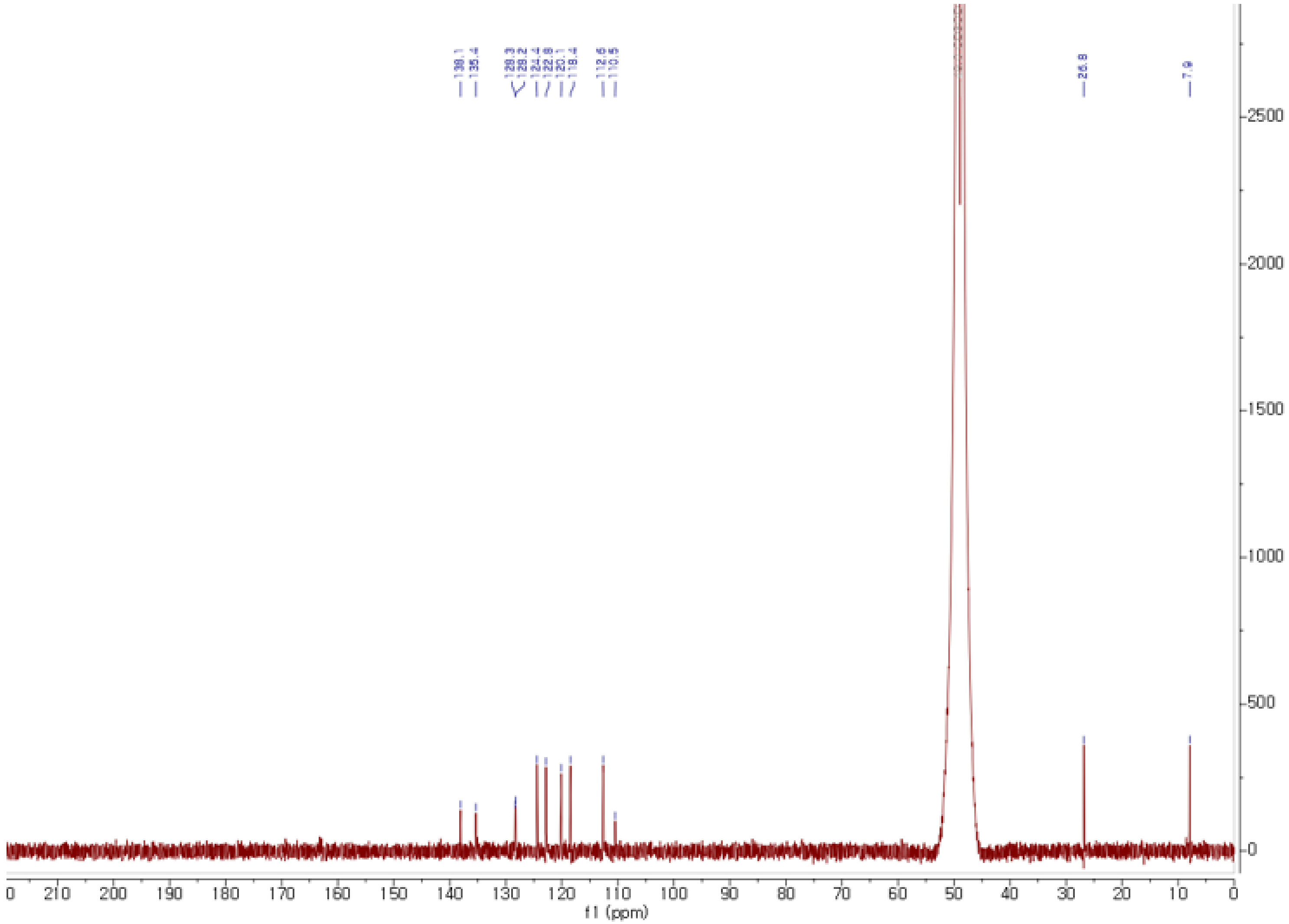

**Figure S15**. The $^{13}$C NMR spectrum of **2** in methanol-$d_4$.

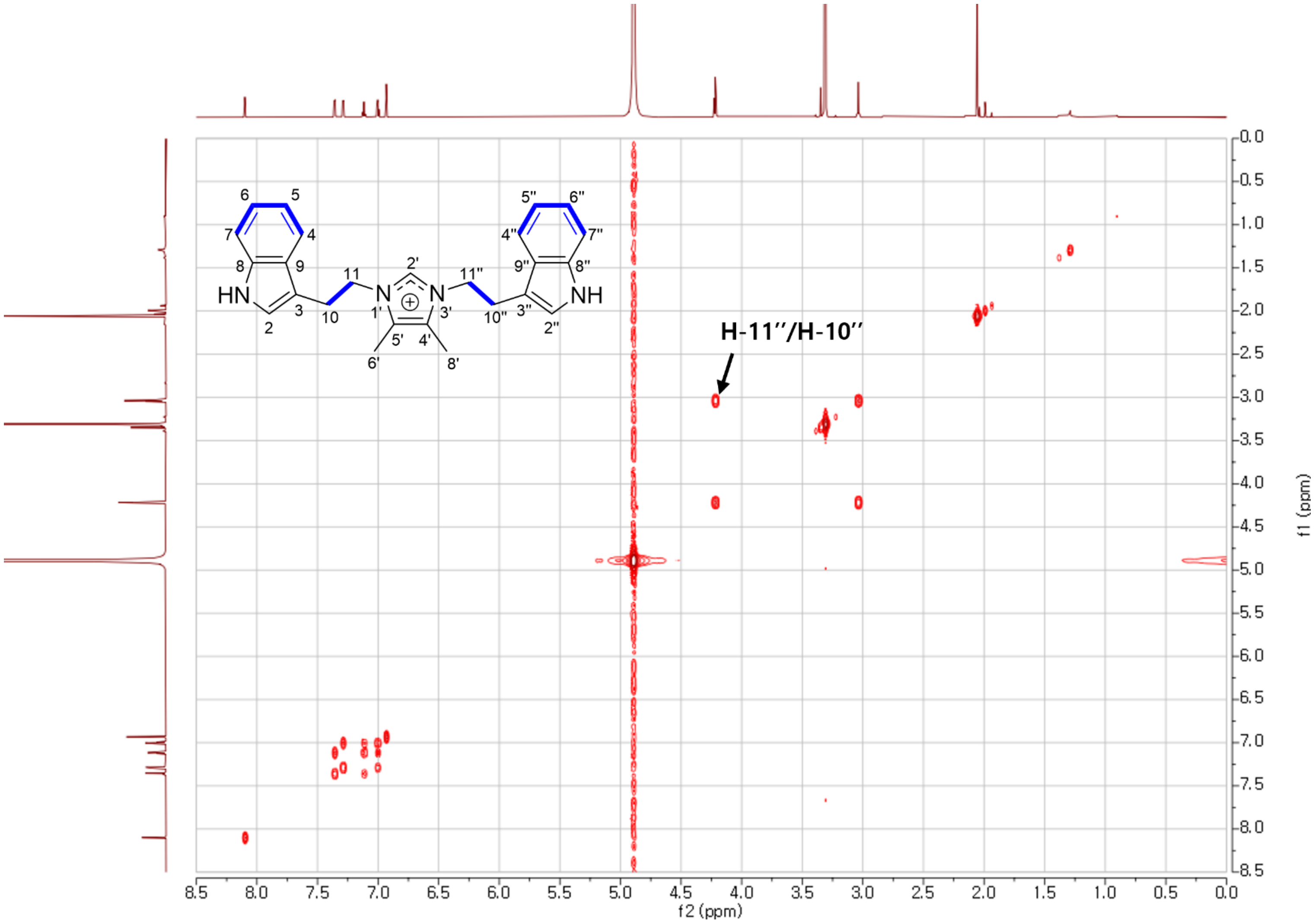

**Figure S16**. The COSY spectrum of **2** in methanol-$d_4$.

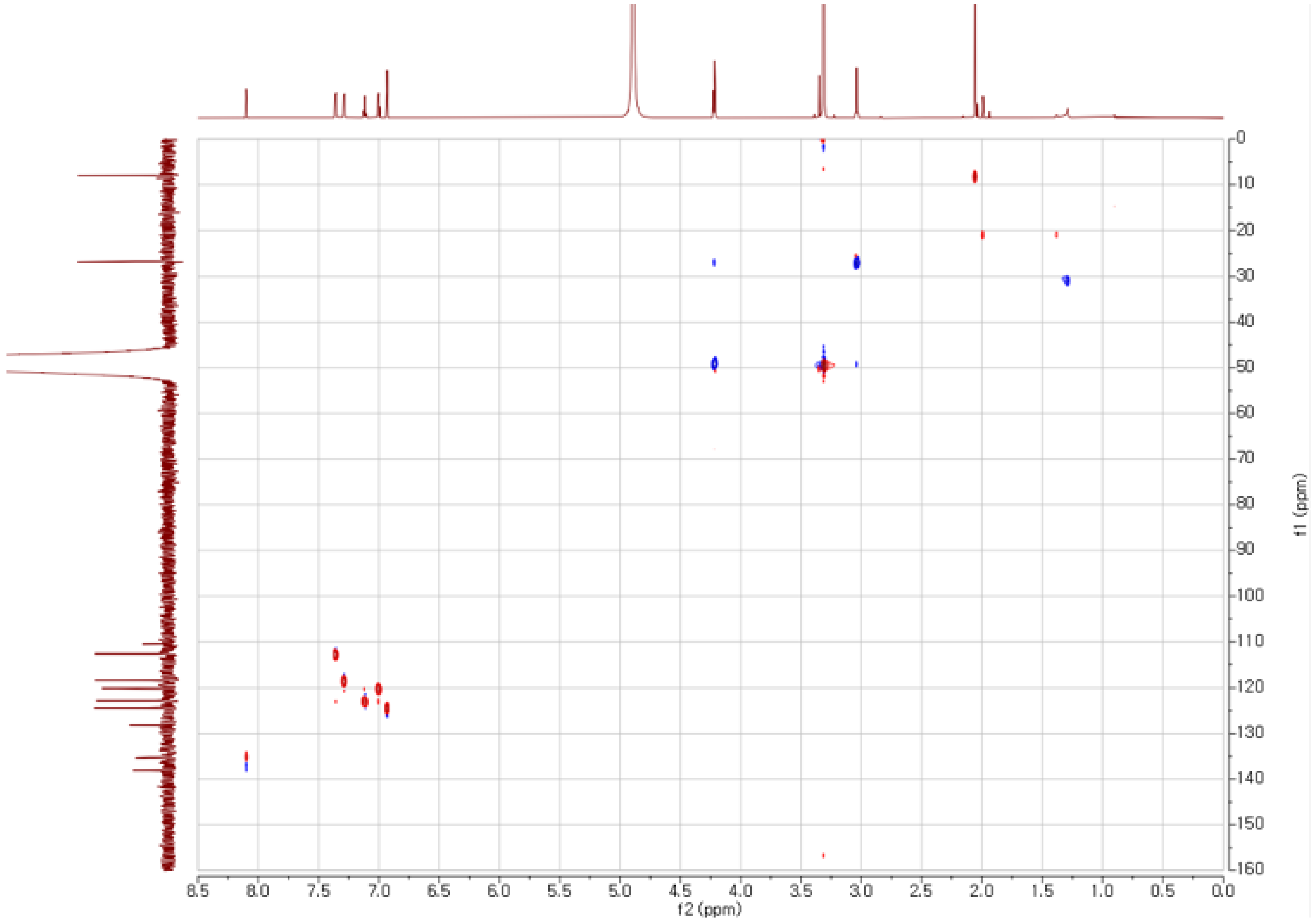

**Figure S17**. The HSQC spectrum of **2** in methanol-$d_4$.

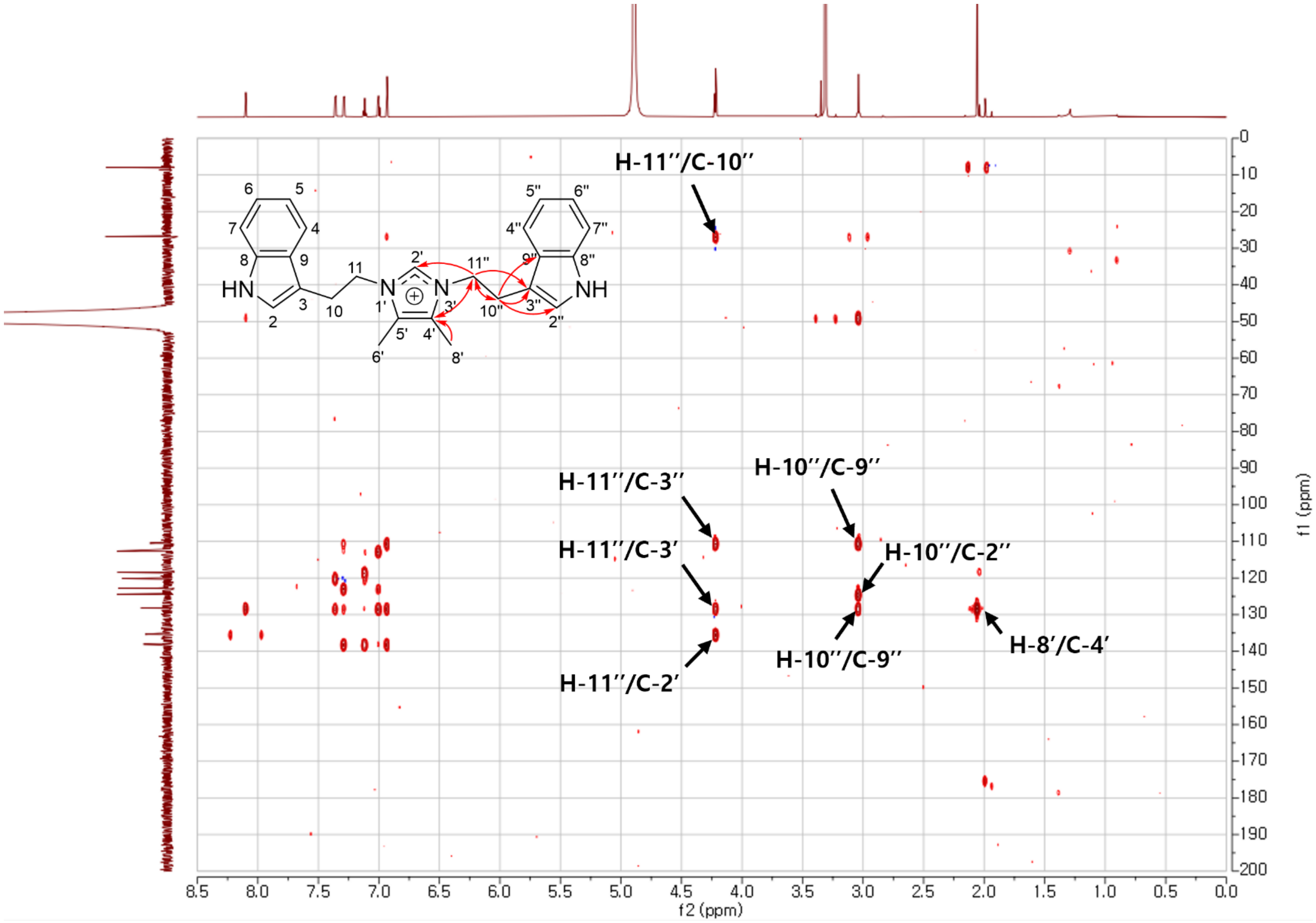

**Figure S18**. The HMBC spectrum of **2** in methanol-$d_4$.

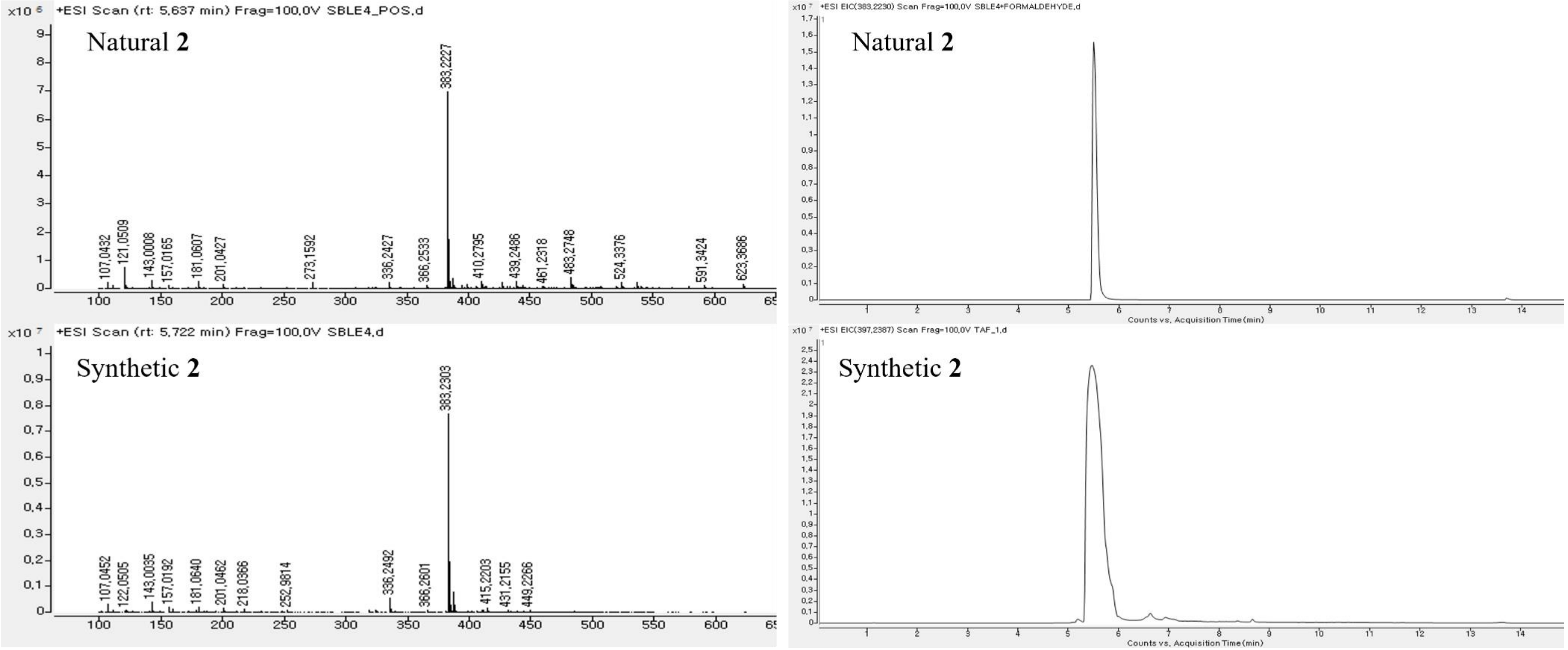

**Figure S19**. HRESIMS and HPLC chromatogram of **2**.

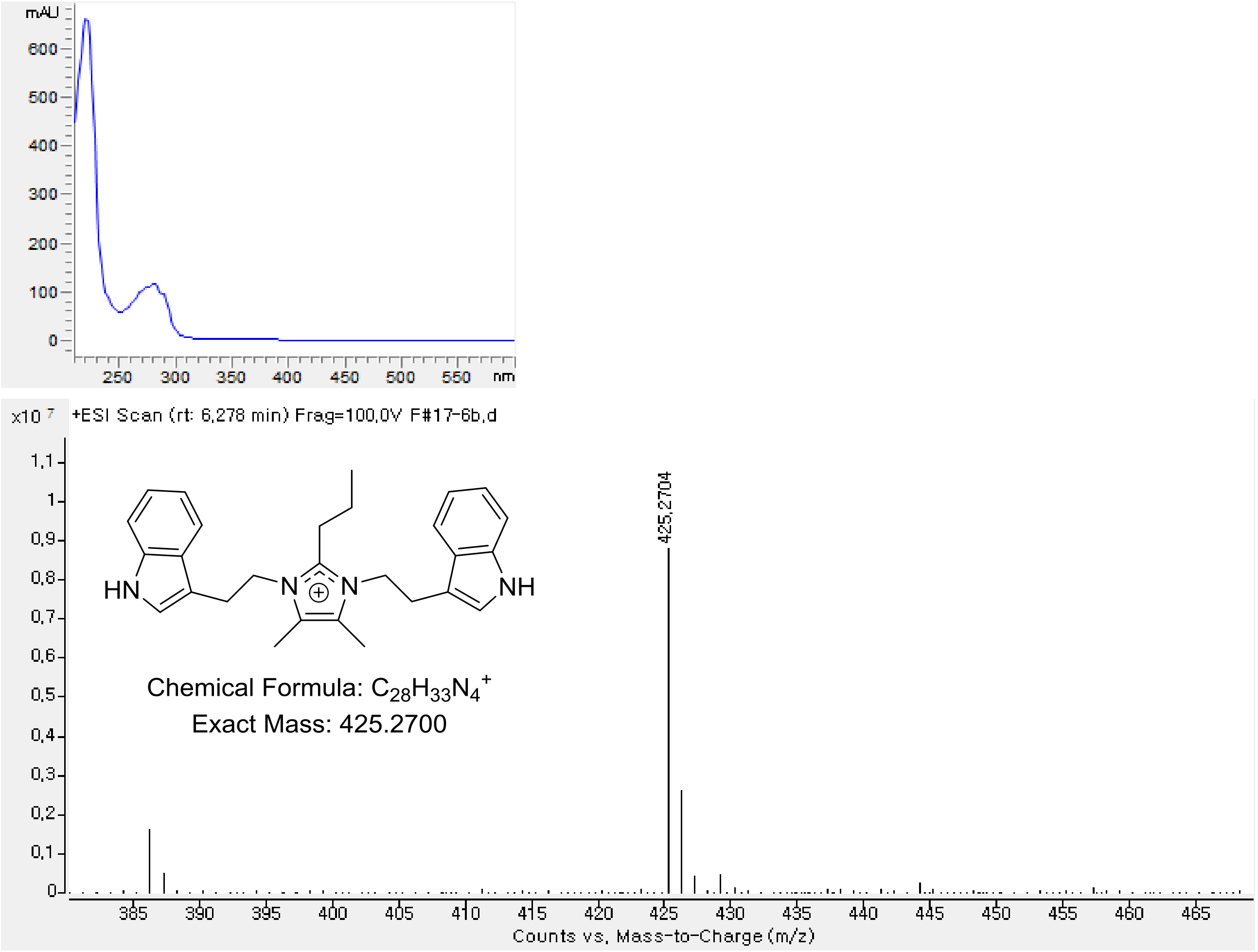

**Figure S20**. UV-Vis and HRESIMS spectrum of **3**.

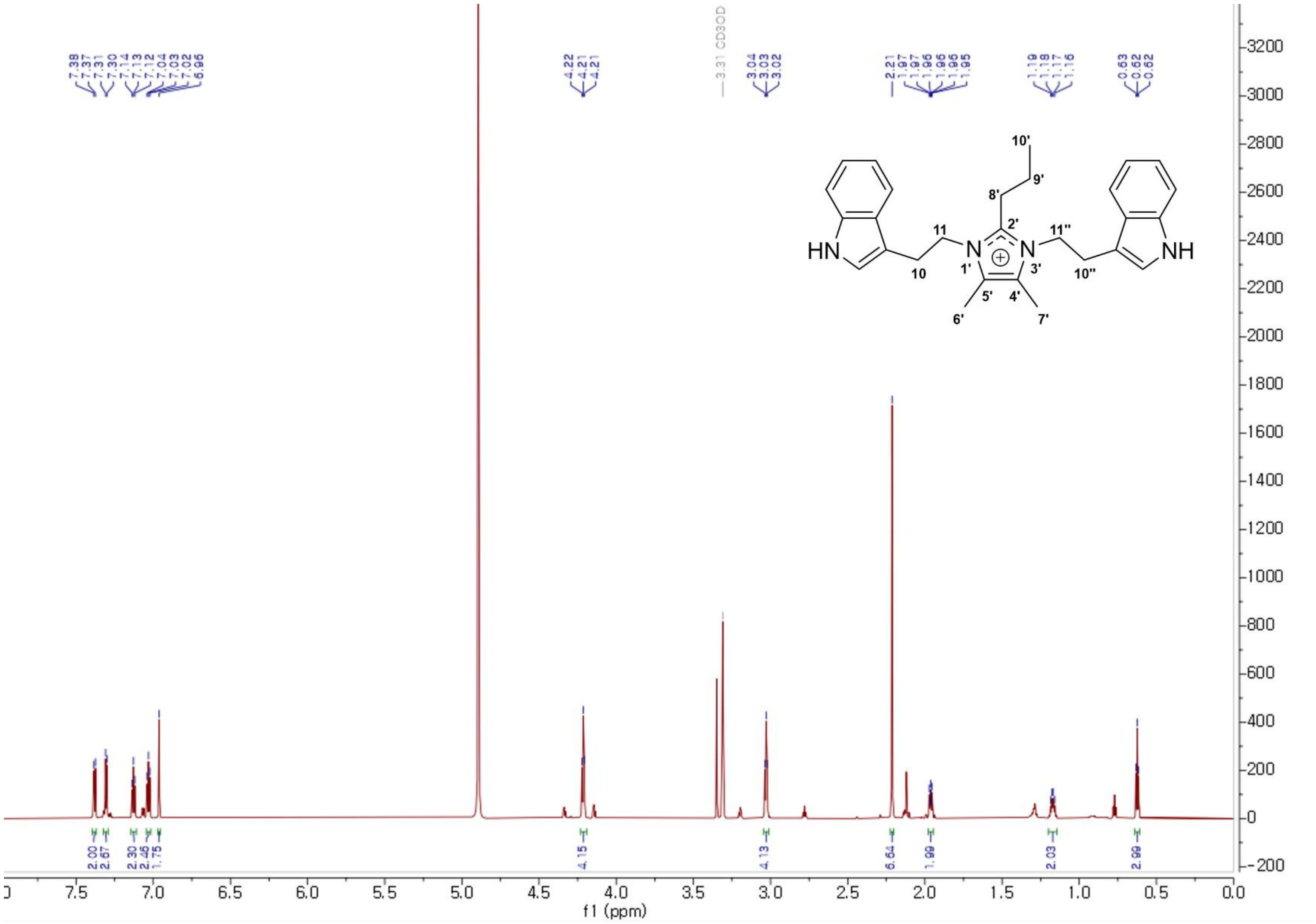

**Figure S21**. The $^{1}$H NMR spectrum of **3** in methanol-$d_4$.

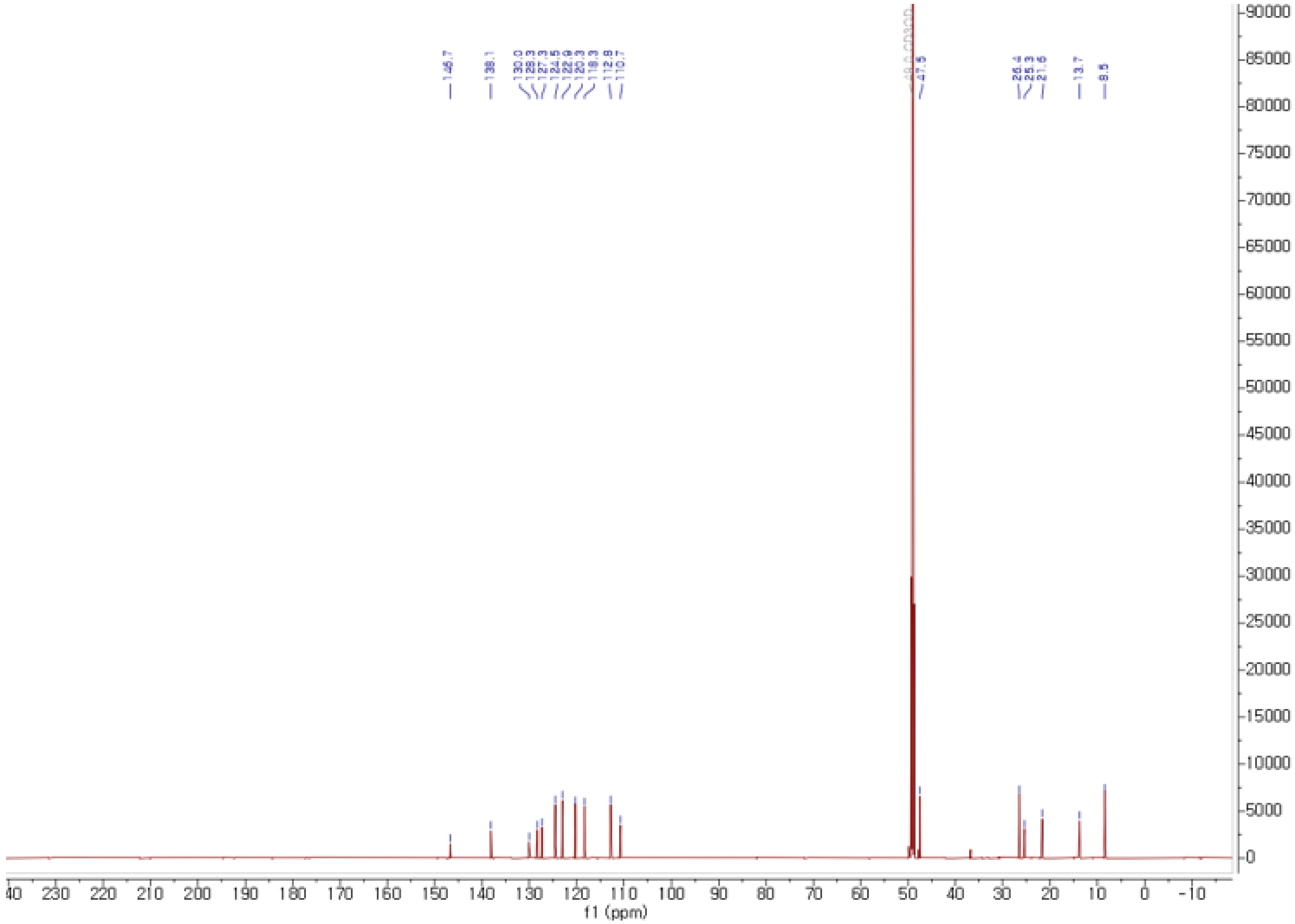

**Figure S22**. The $^{13}C$ NMR spectrum of **3** in methanol-$d_4$.

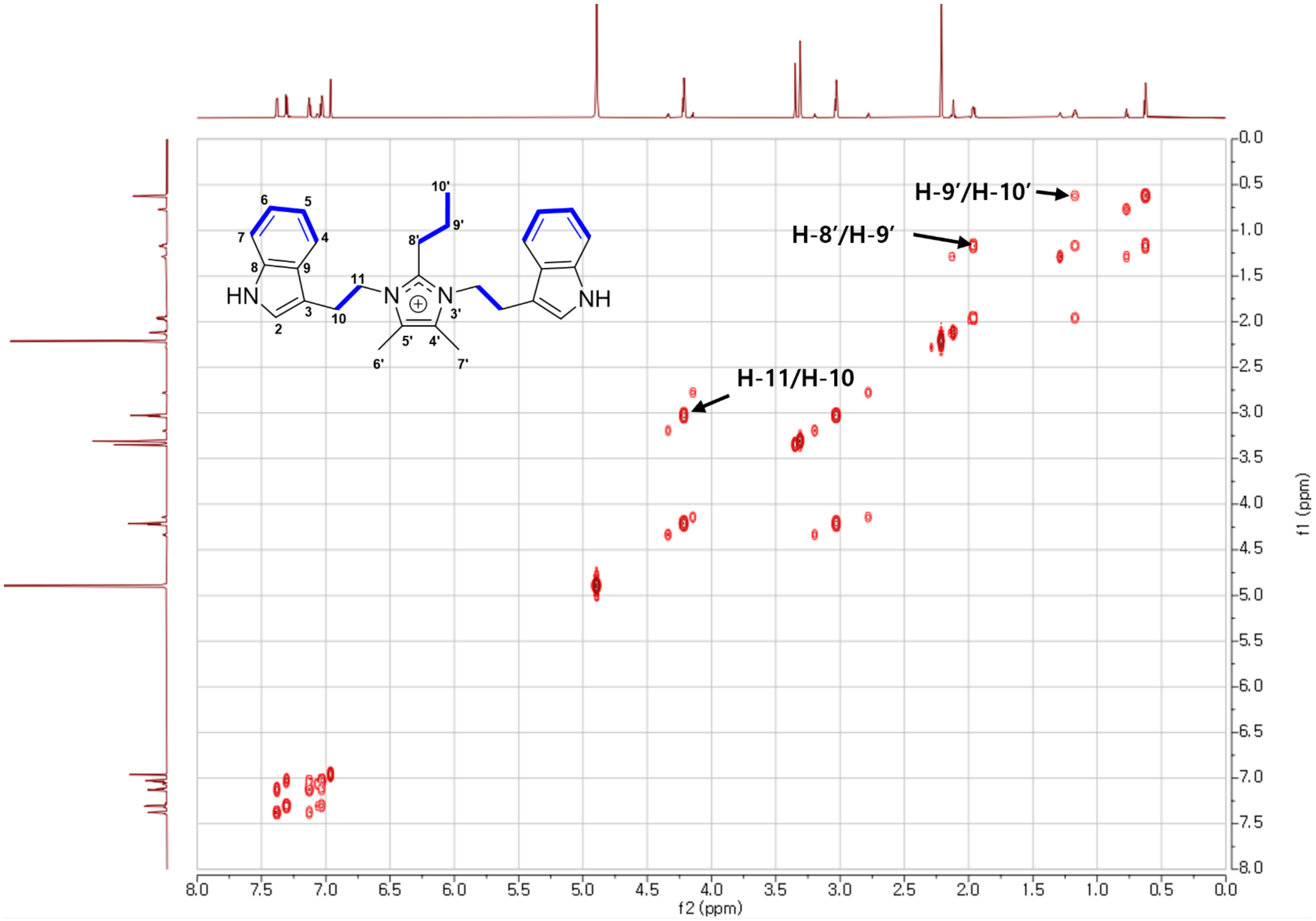

**Figure S23**. The COSY spectrum of **3** in methanol-$d_4$.

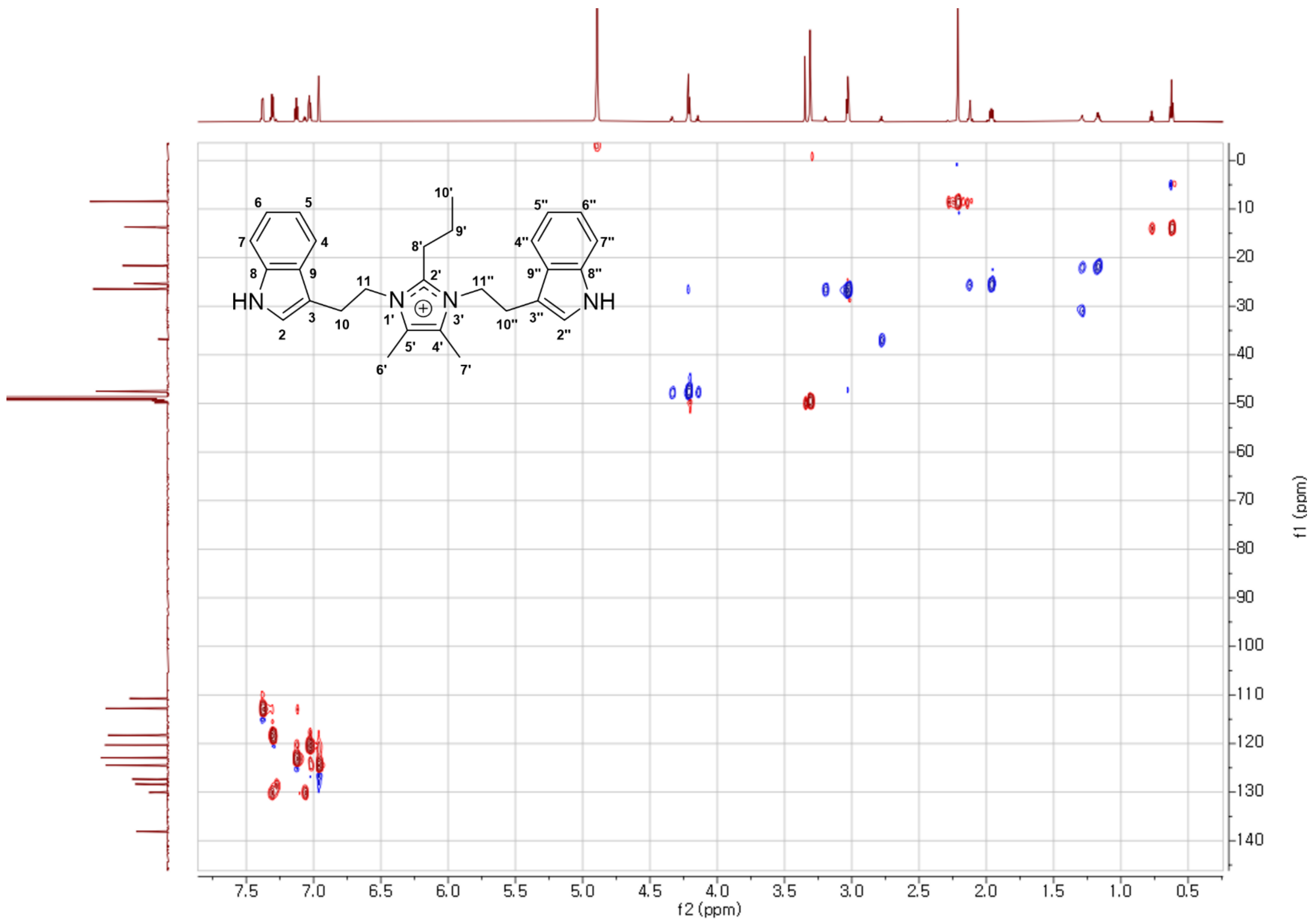

**Figure S24**. The HSQC spectrum of **3** in methanol-$d_4$.

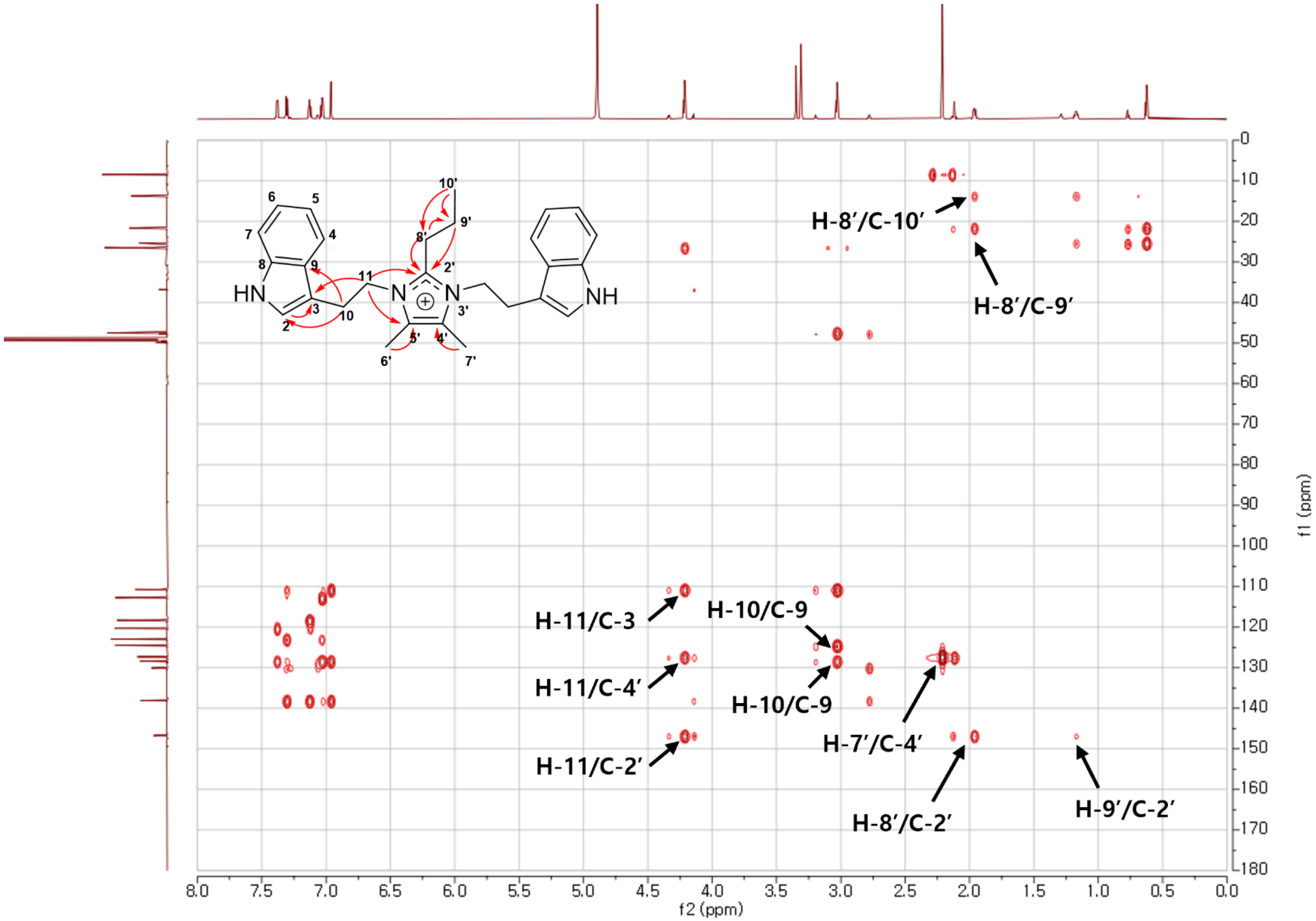

**Figure S25**. The HMBC spectrum of **3** in methanol-$d_4$.

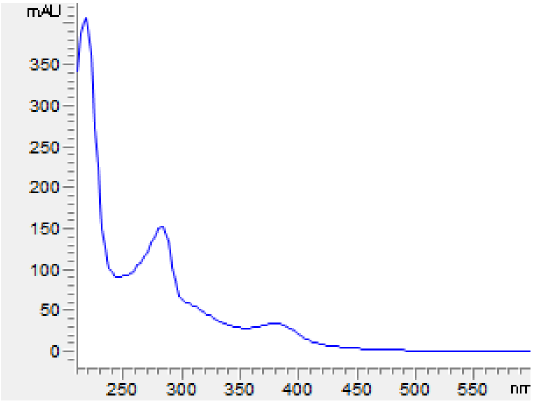

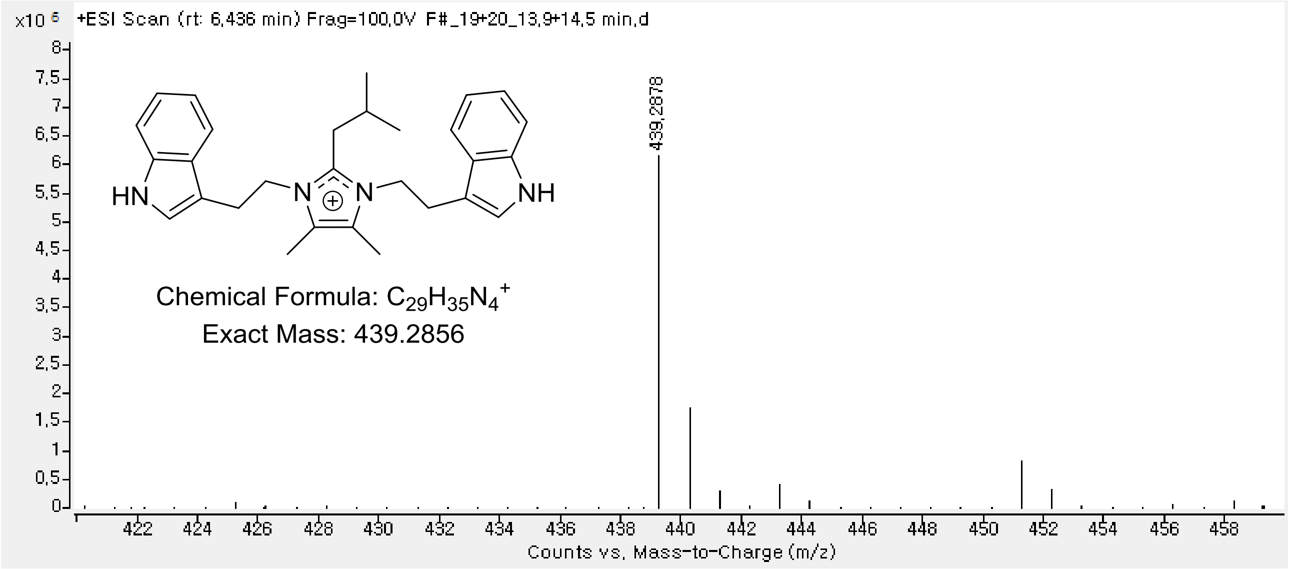

**Figure S26**. UV-Vis and HRESIMS spectrum of **4**.

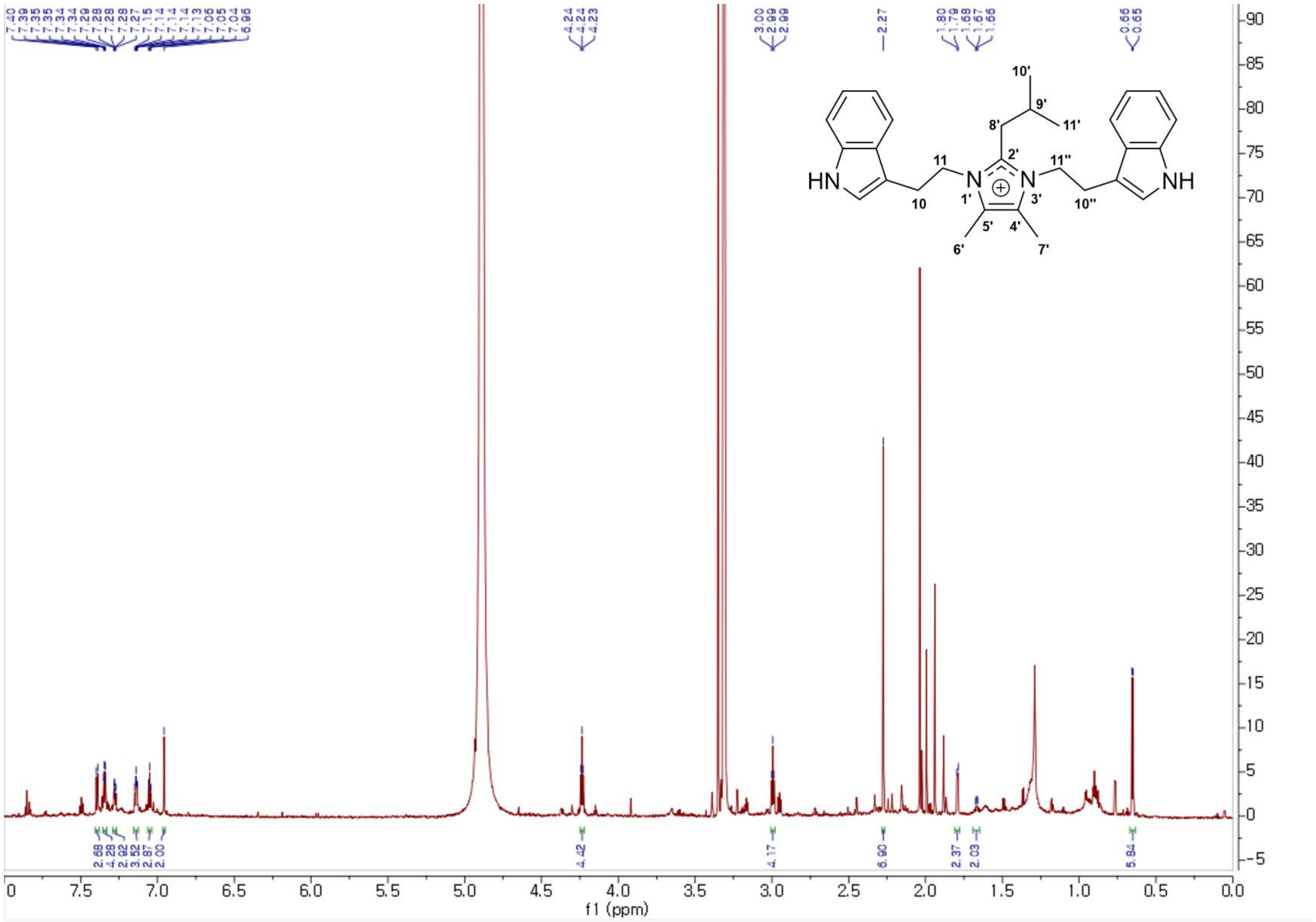

**Figure S27**. The $^{1}H$ NMR spectrum of **4** in methanol-$d_4$.

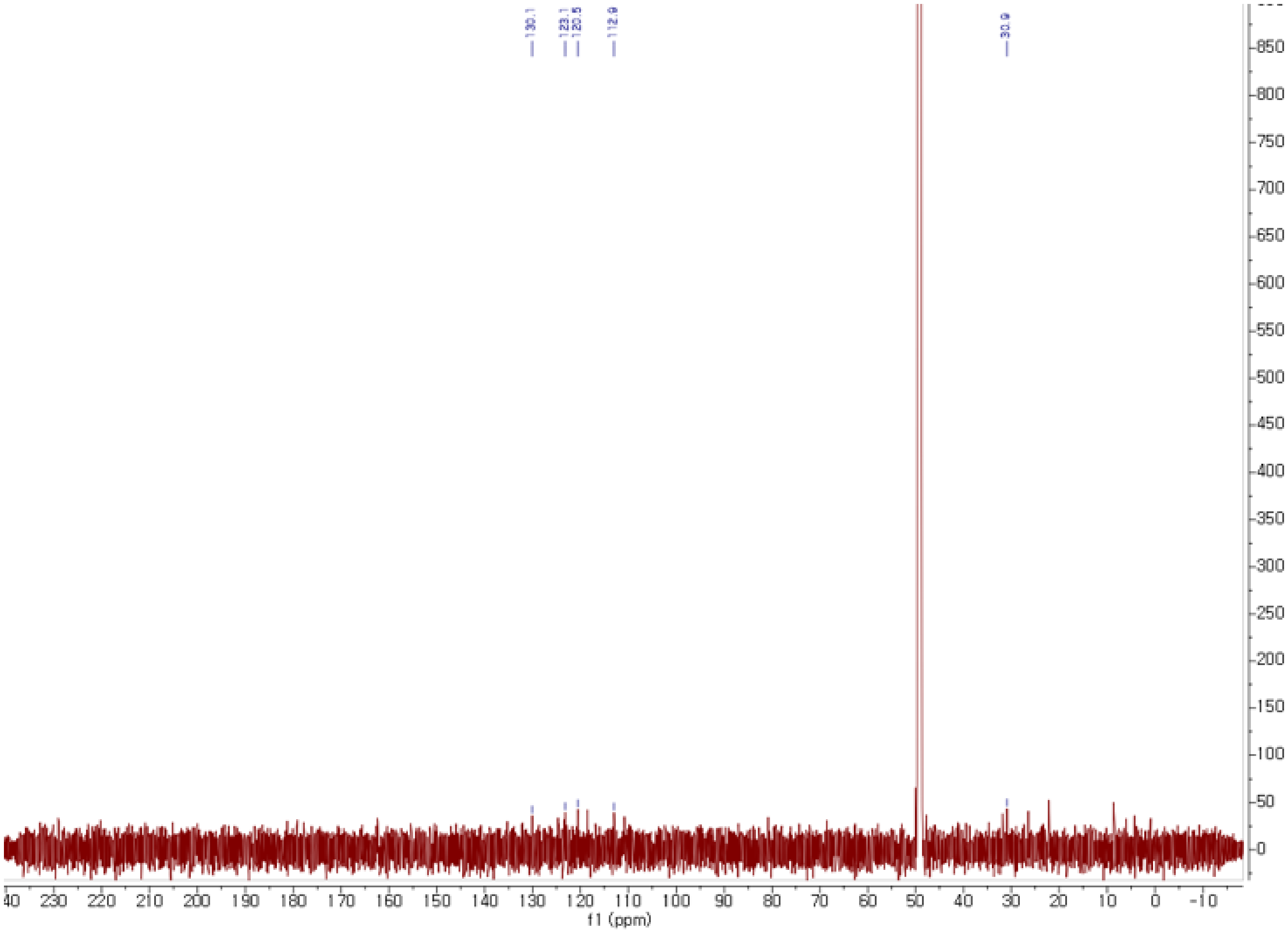

**Figure S28**. The $^{13}$C NMR spectrum of **4** in methanol-$d_4$.

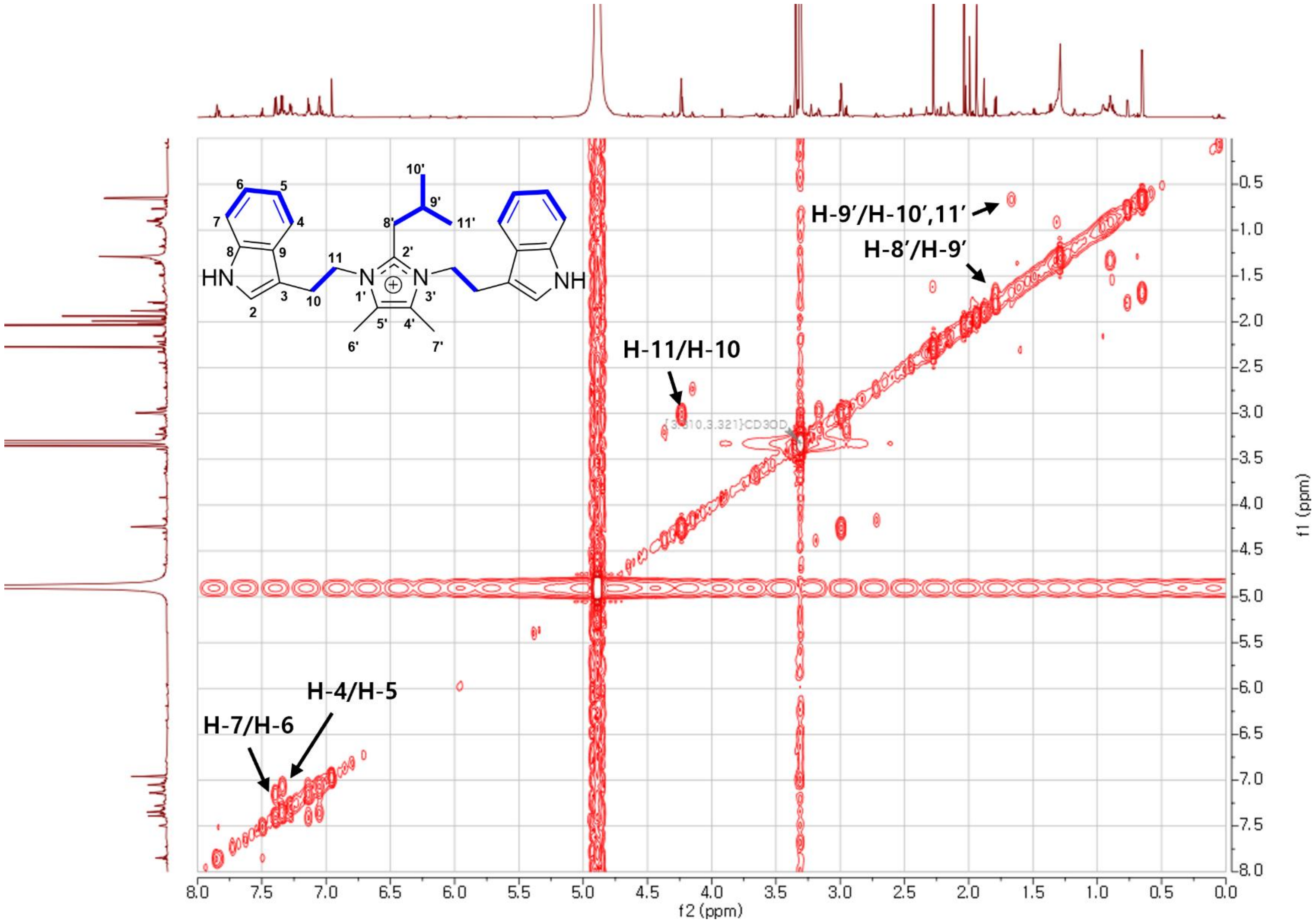

**Figure S29**. The COSY spectrum of **4** in methanol-$d_4$.

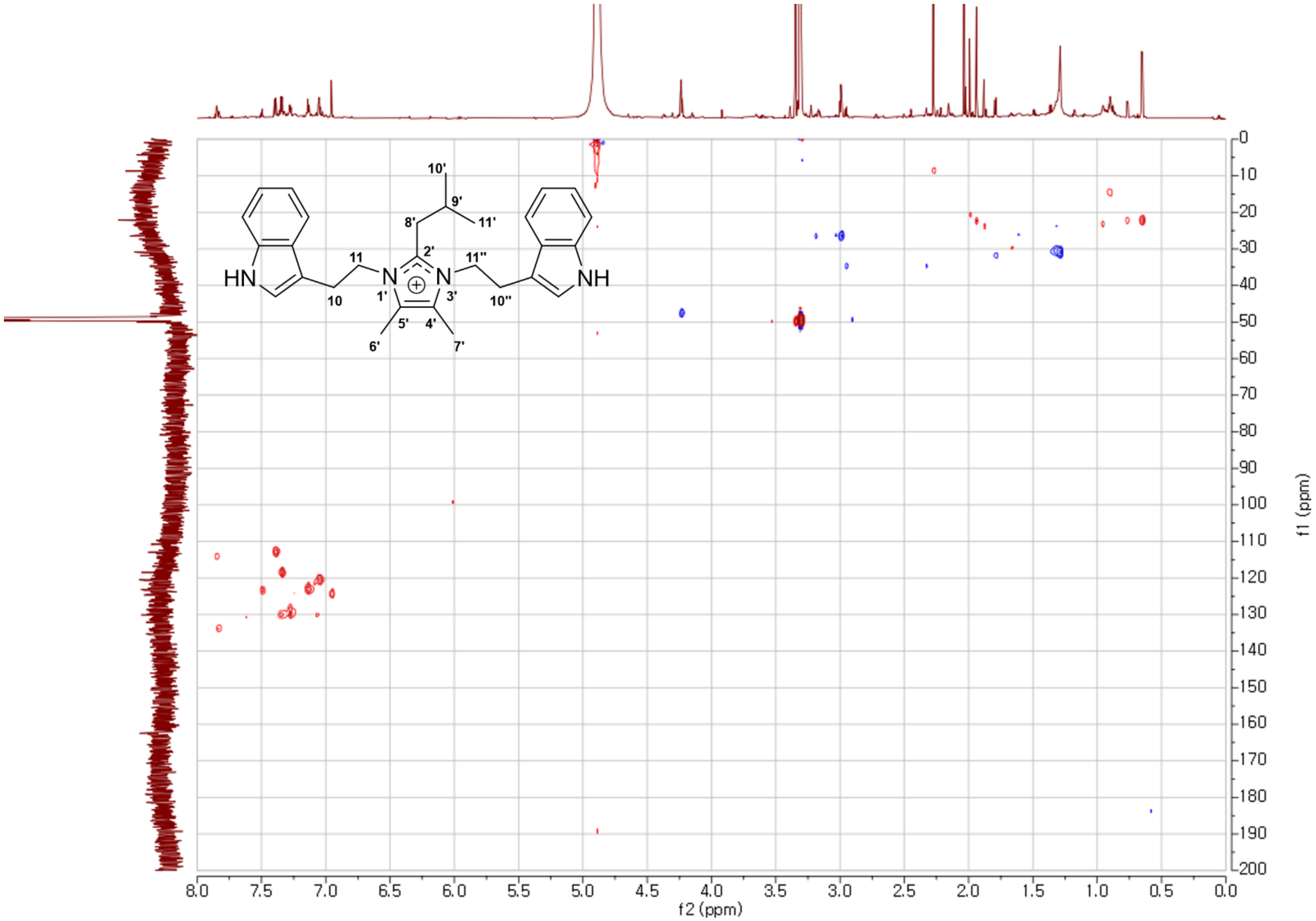

**Figure S30**. The HSQC spectrum of **4** in methanol-$d_4$.

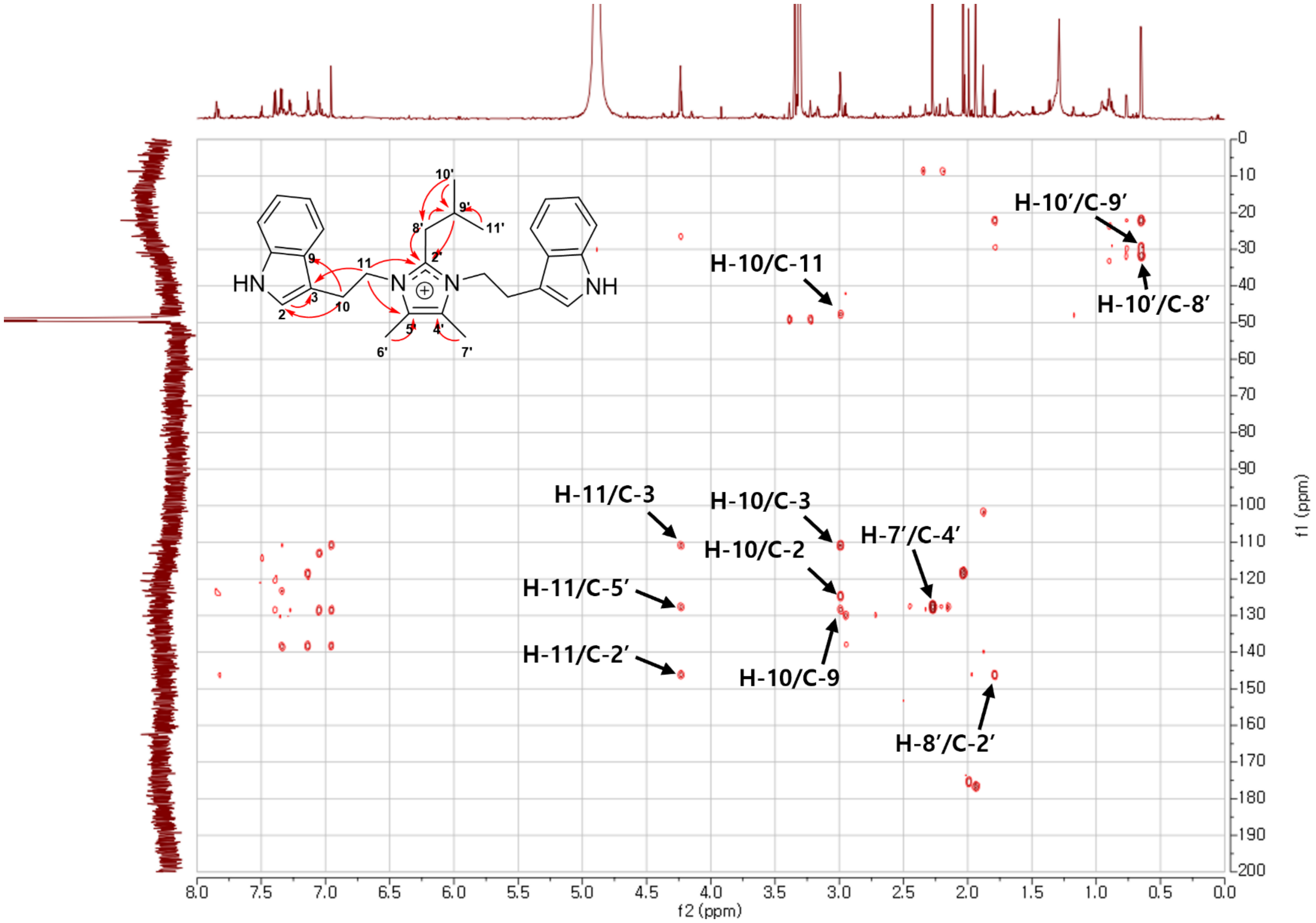

**Figure S31**. The HMBC spectrum of **4** in methanol-$d_4$.

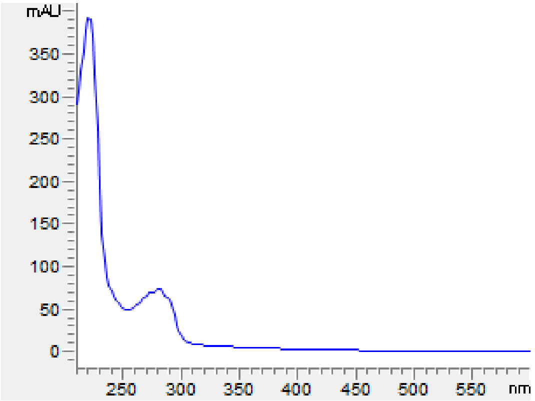

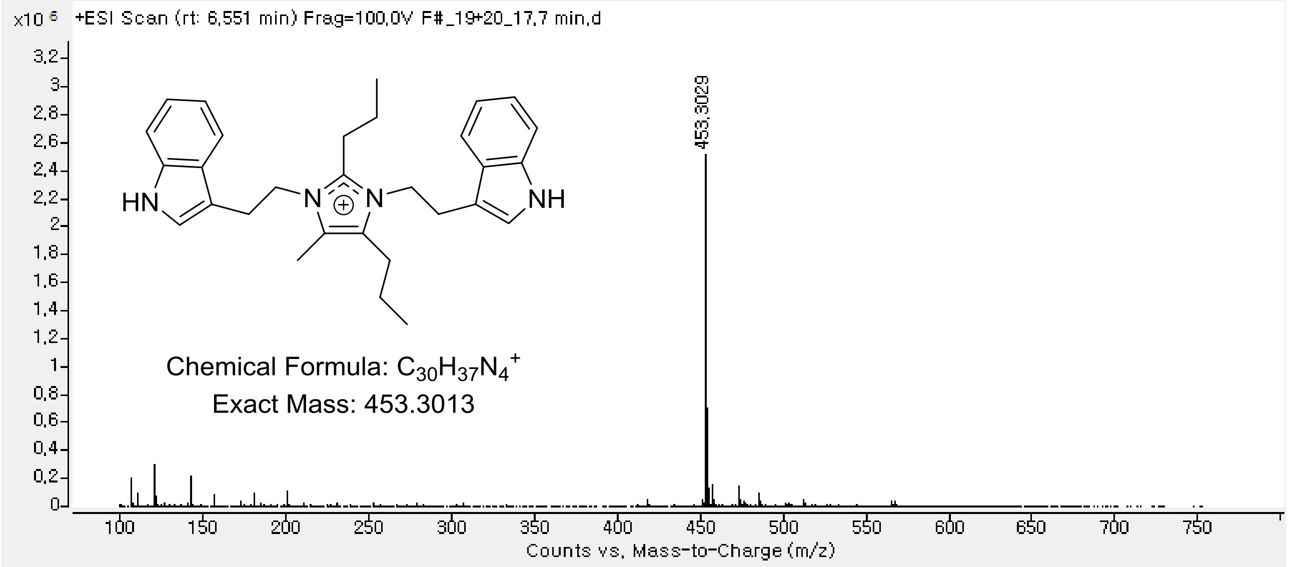

**Figure S32**. UV-Vis and HRESIMS spectrum of **5**.

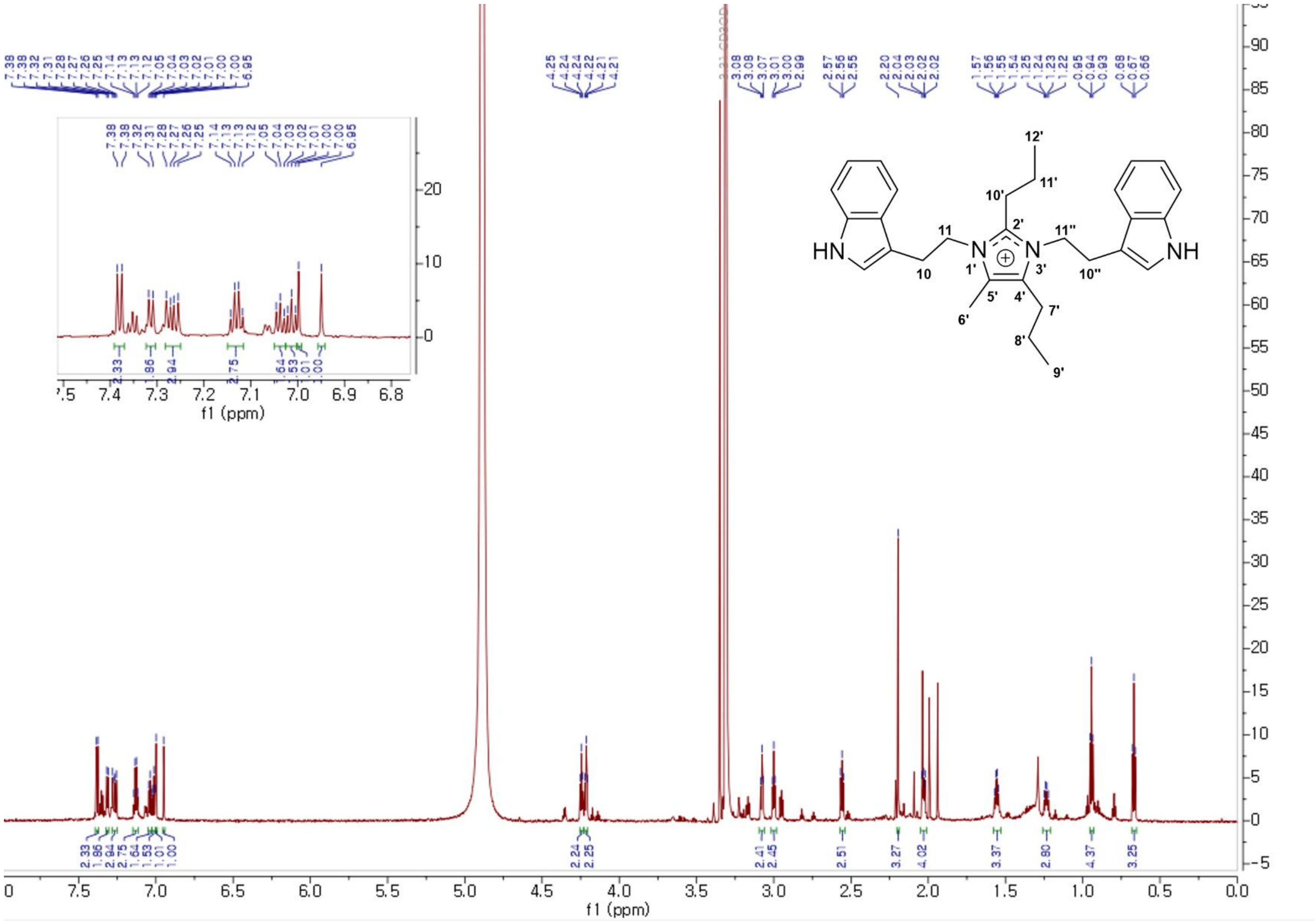

**Figure S33**. The $^{1}H$ NMR spectrum of **5** in methanol-$d_4$.

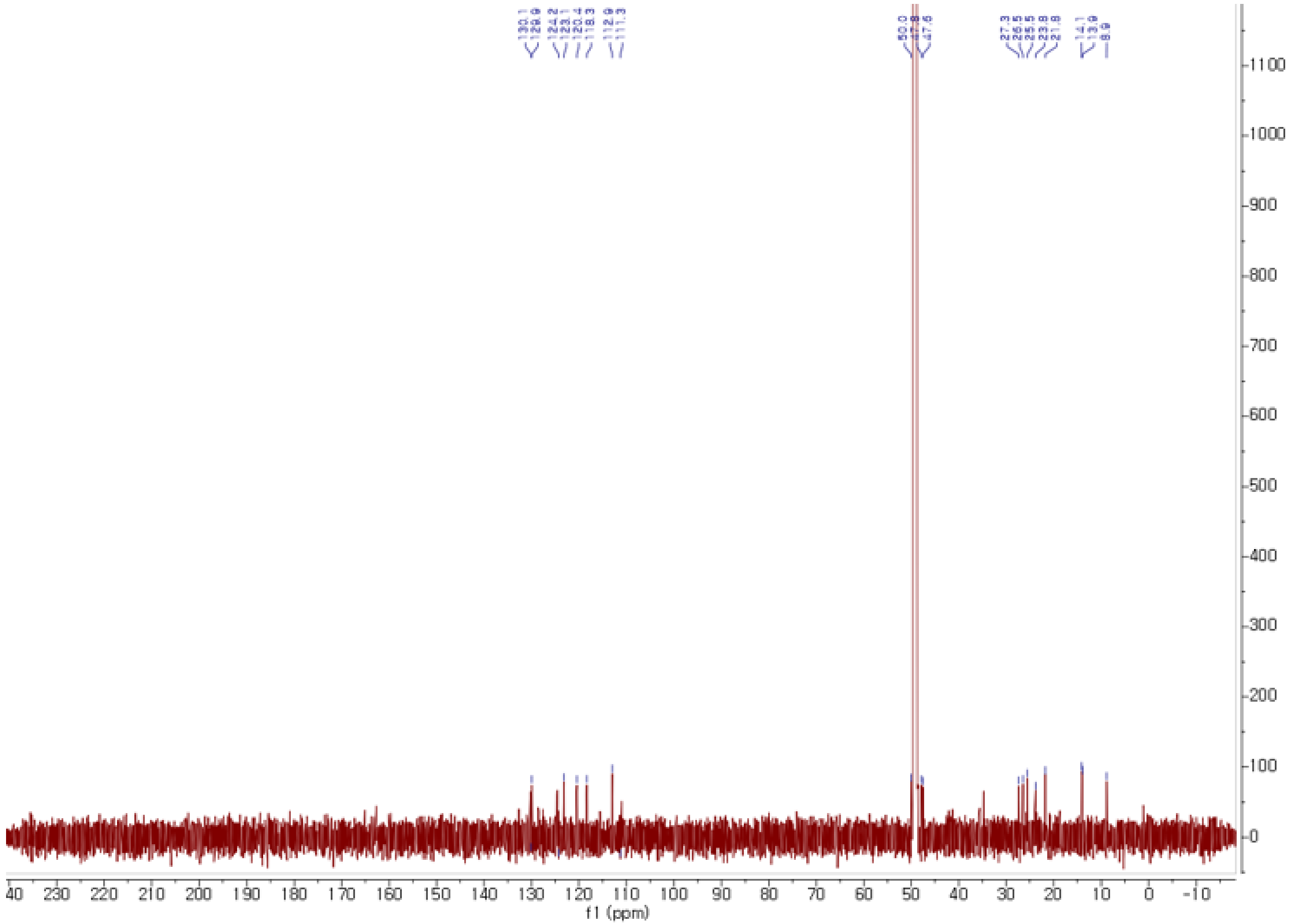

**Figure S34**. The $^{13}C$ NMR spectrum of **5** in methanol-$d_4$.

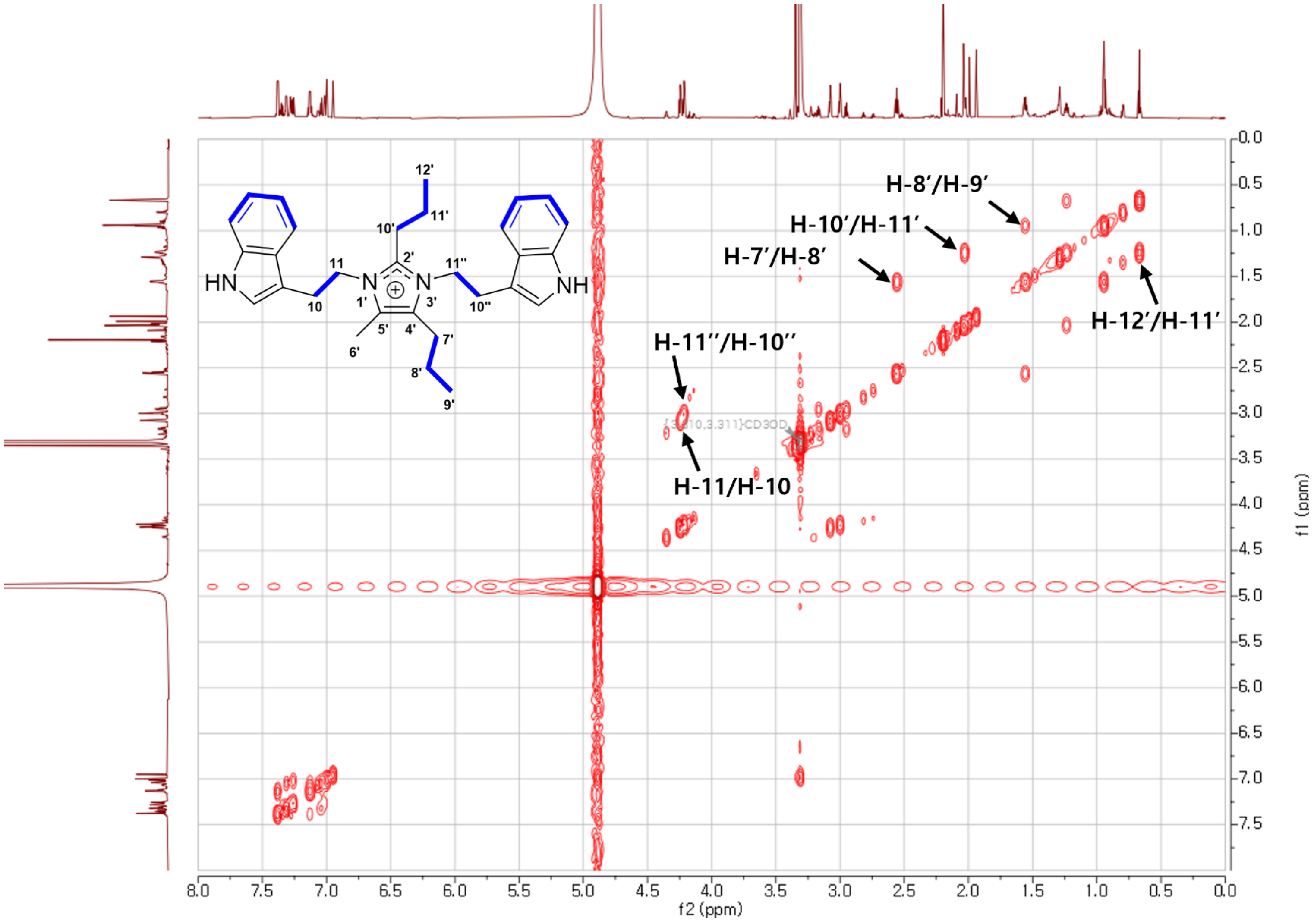

**Figure S35**. The COSY spectrum of **5** in methanol-$d_4$.

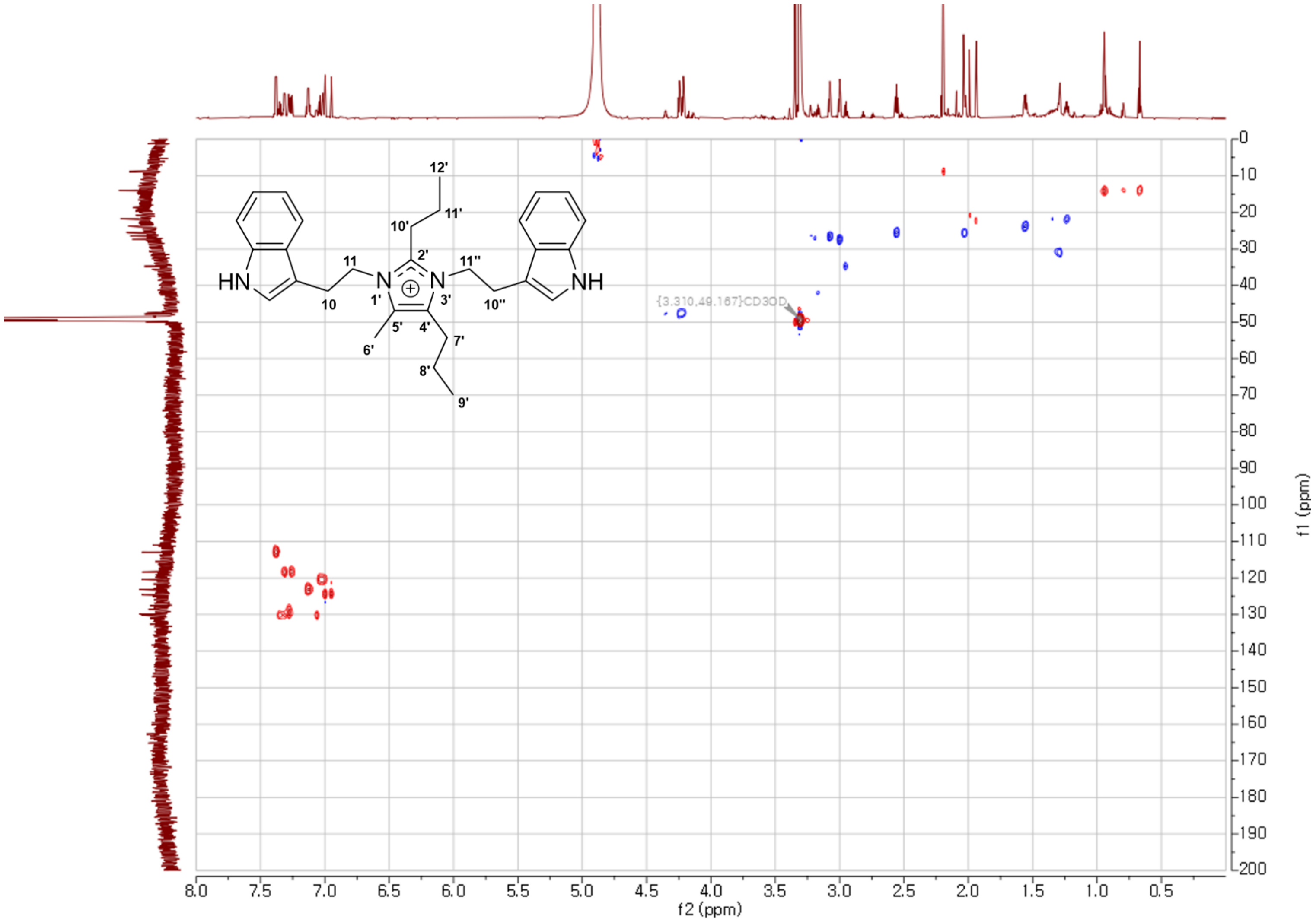

**Figure S36**. The HSQC spectrum of **5** in methanol-*d*$_4$.

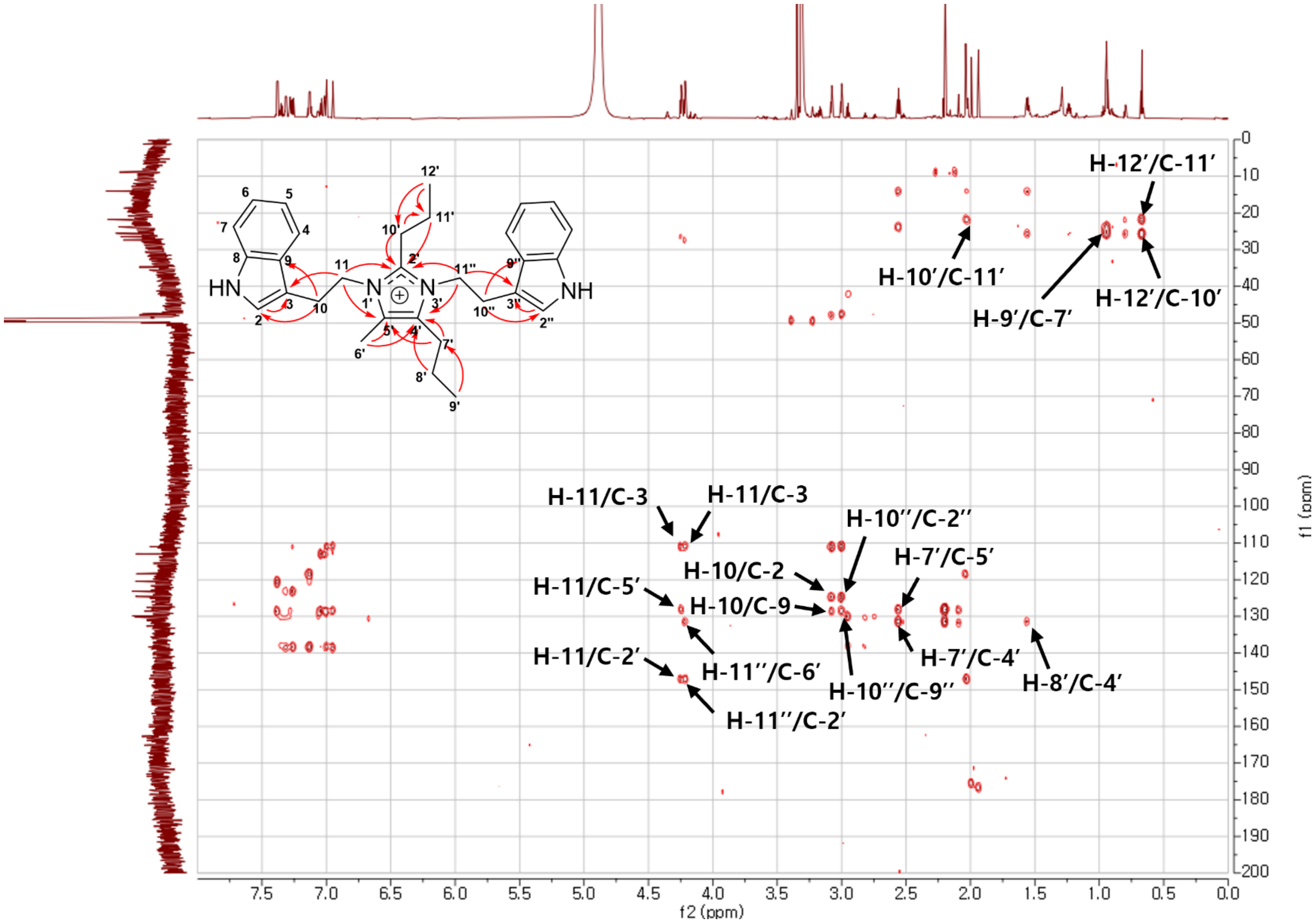

**Figure S37**. The HMBC spectrum of **5** in methanol-$d_4$.

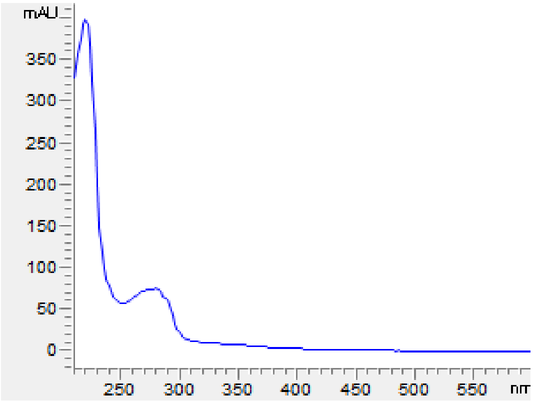

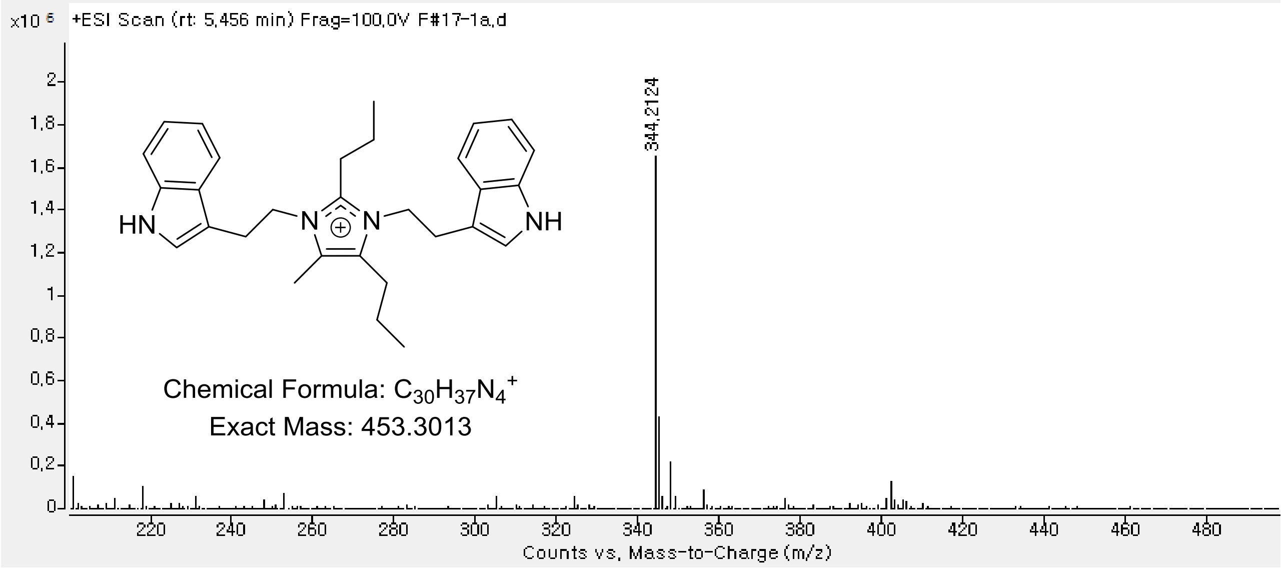

**Figure S38**. UV-Vis and HRESIMS spectrum of **6**.

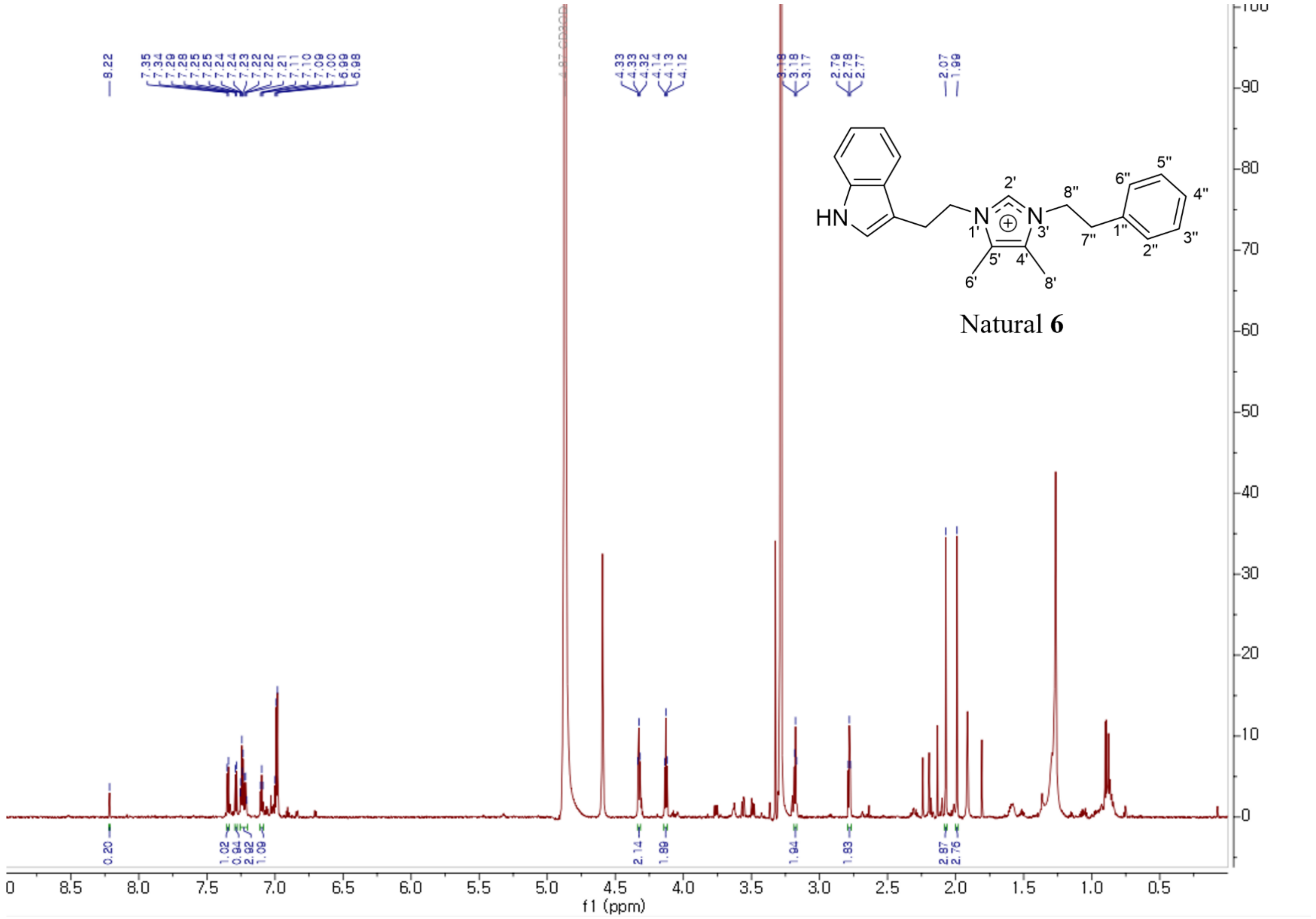

**Figure S39**. The $^{1}$H NMR spectrum of **6** in methanol-$d_4$.

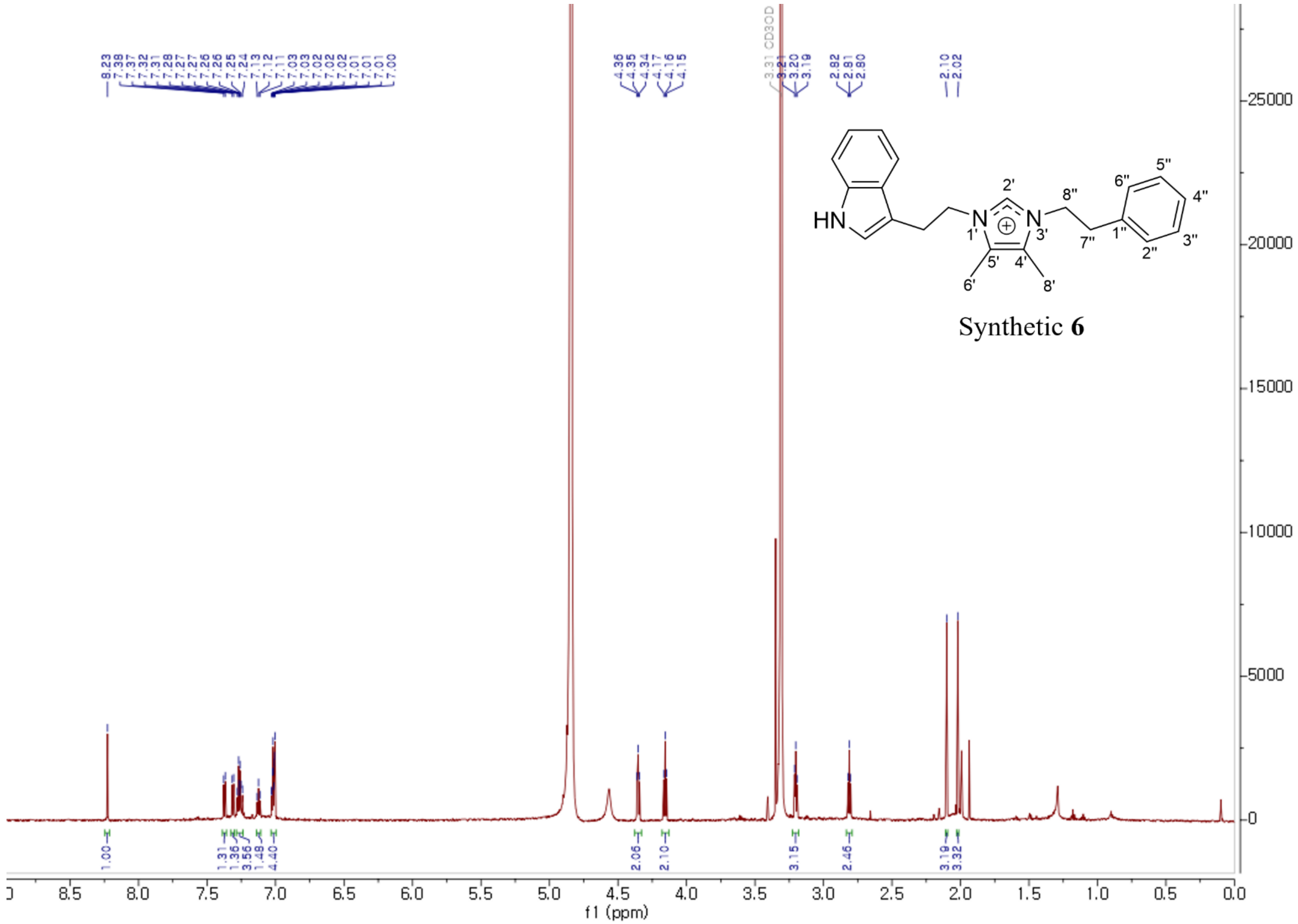

**Figure S40**. The $^{1}H$ NMR spectrum of synthetic **6** in methanol-$d_4$.

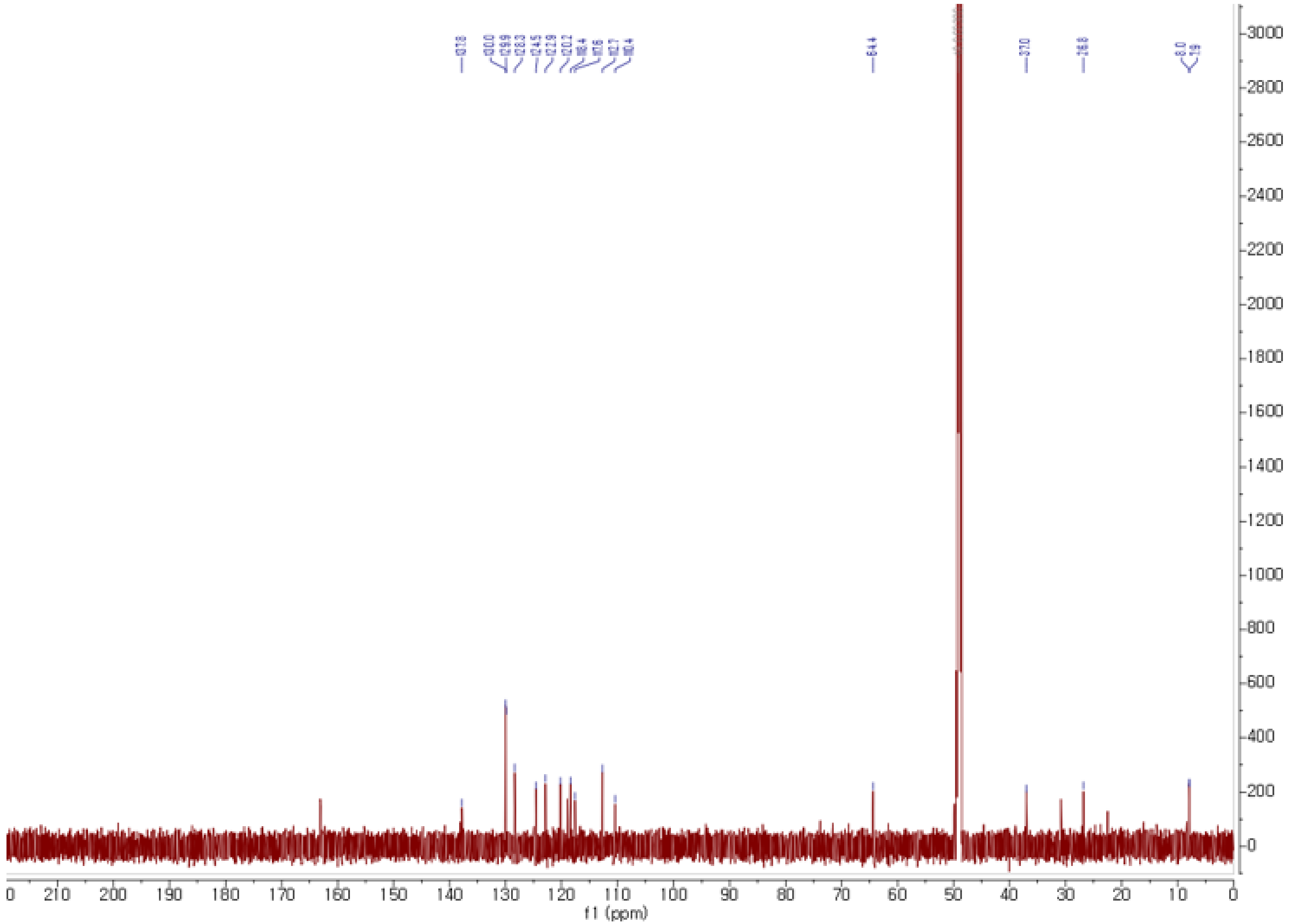

**Figure S41**. The $^{13}C$ NMR spectrum of **6** in methanol-$d_4$.

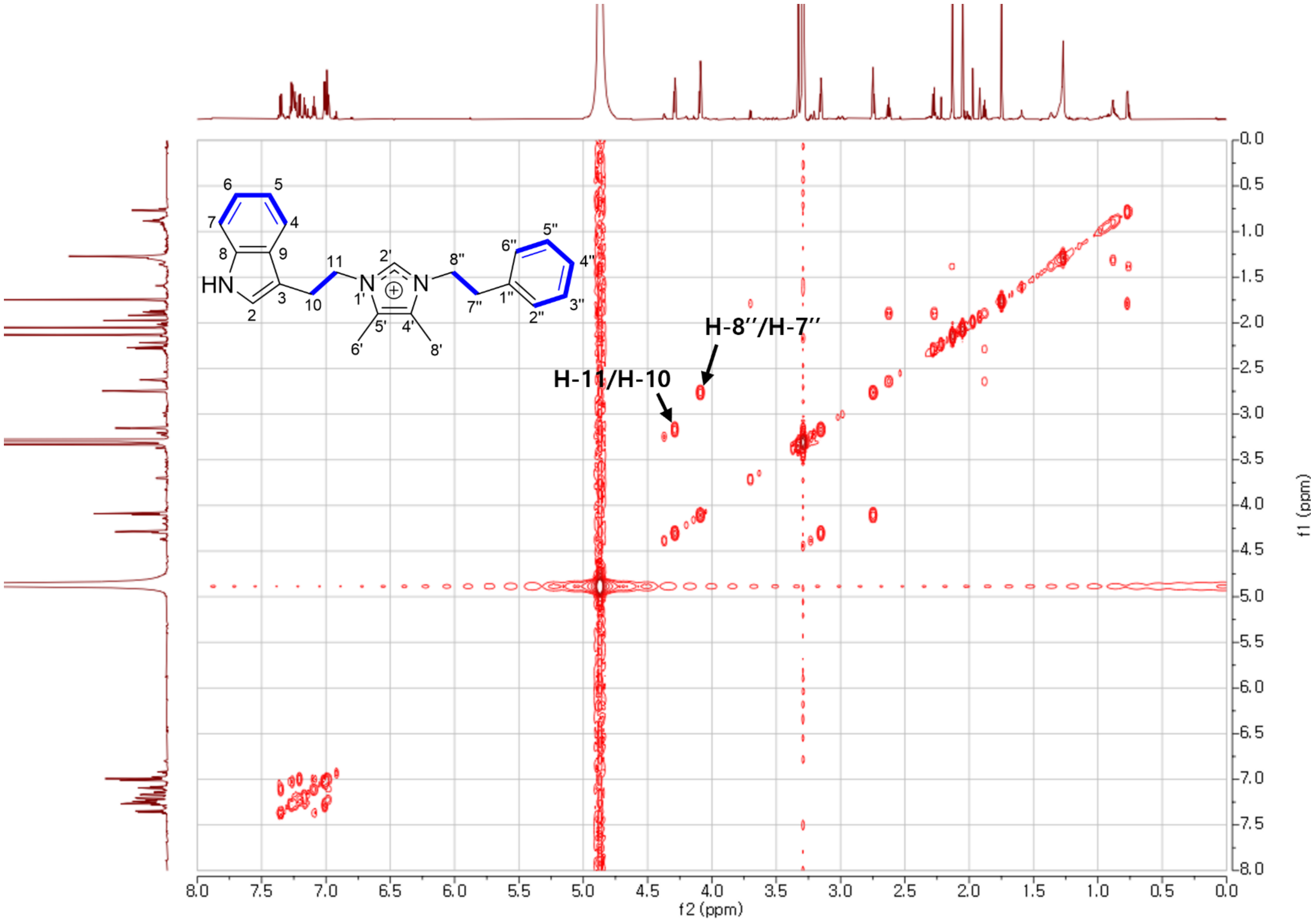

**Figure S42**. The COSY spectrum of **6** in methanol-$d_4$.

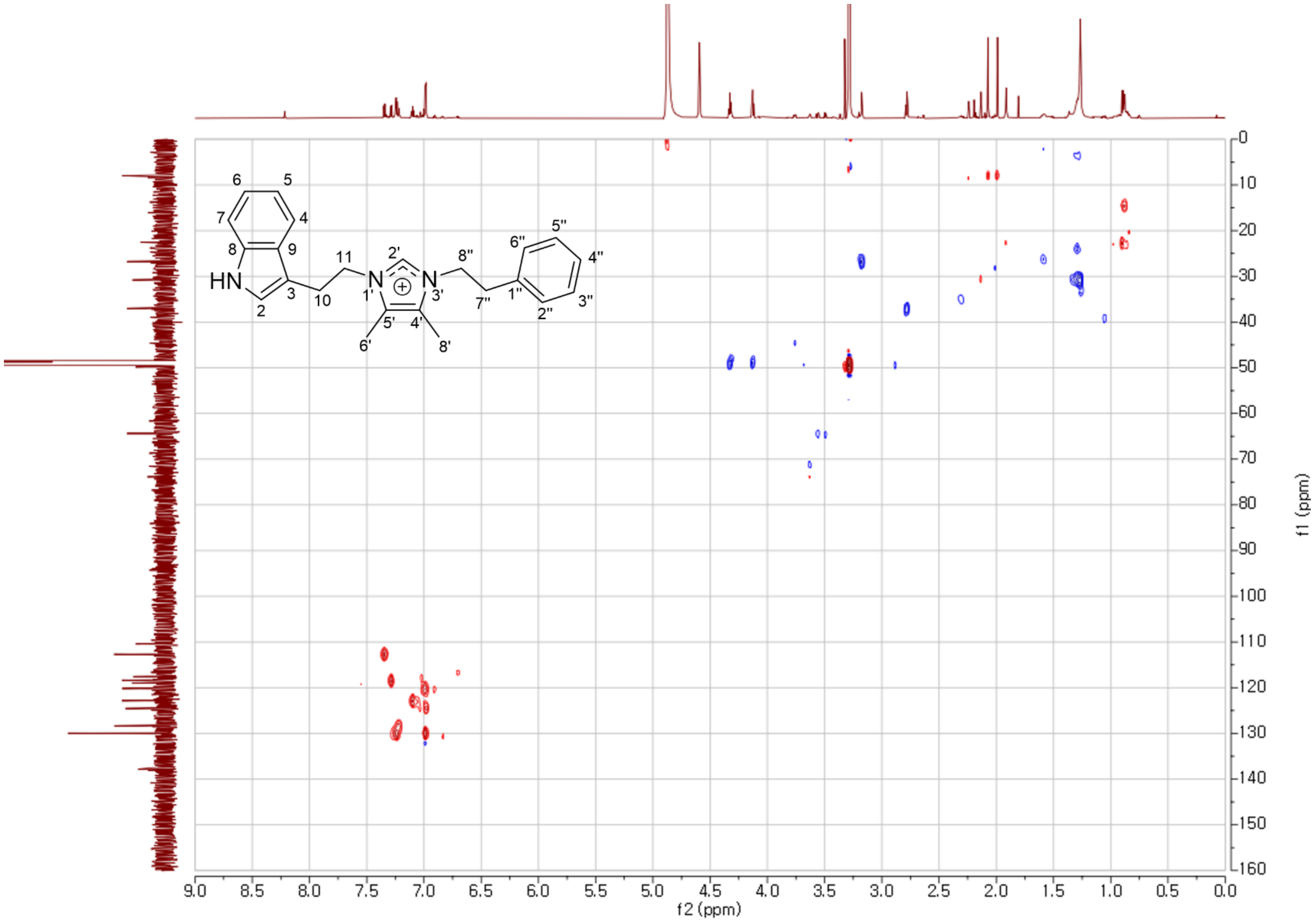

**Figure S43**. The HSQC spectrum of **6** in methanol-$d_4$.

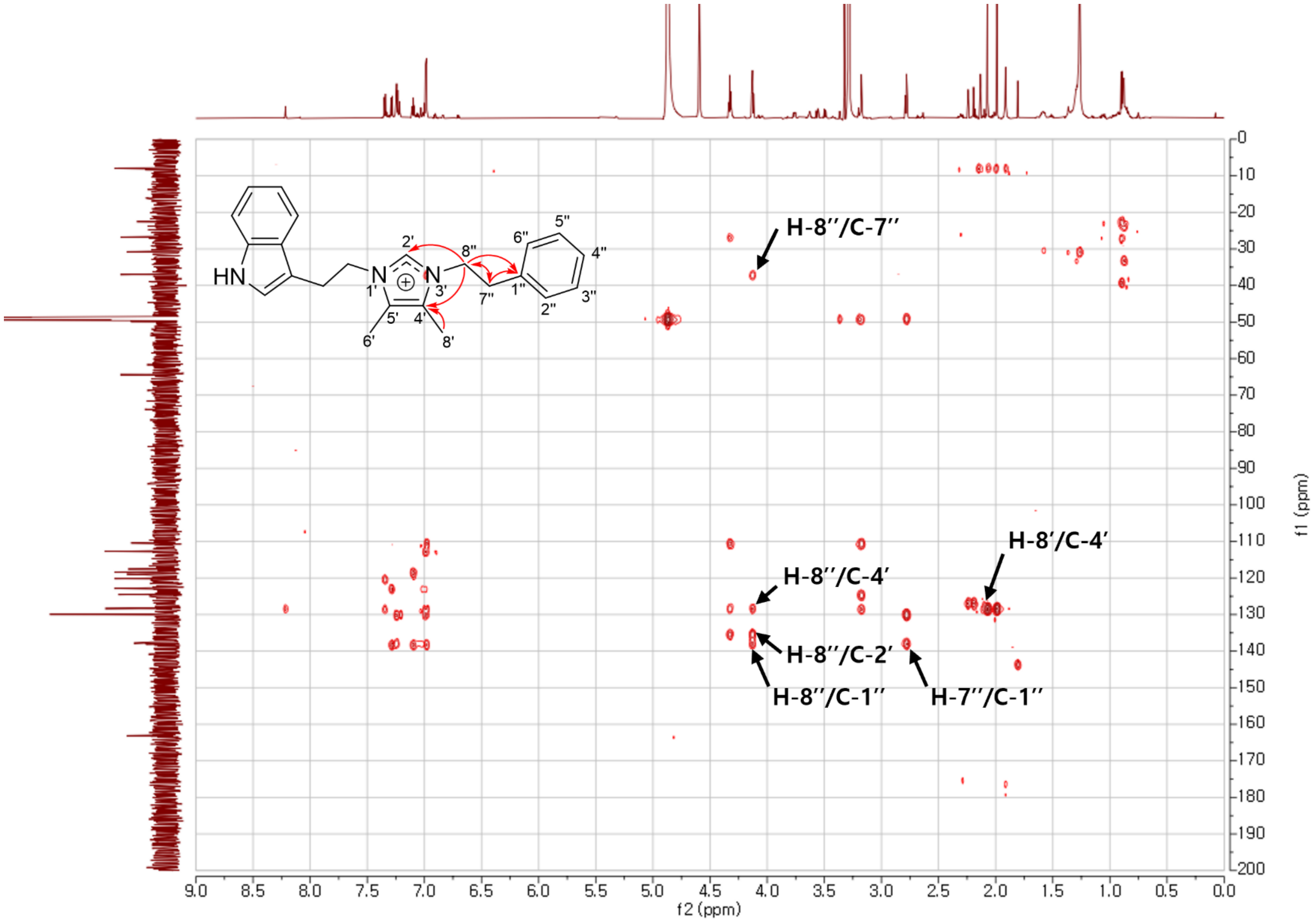

**Figure S44**. The HMBC spectrum of **6** in methanol-$d_4$.

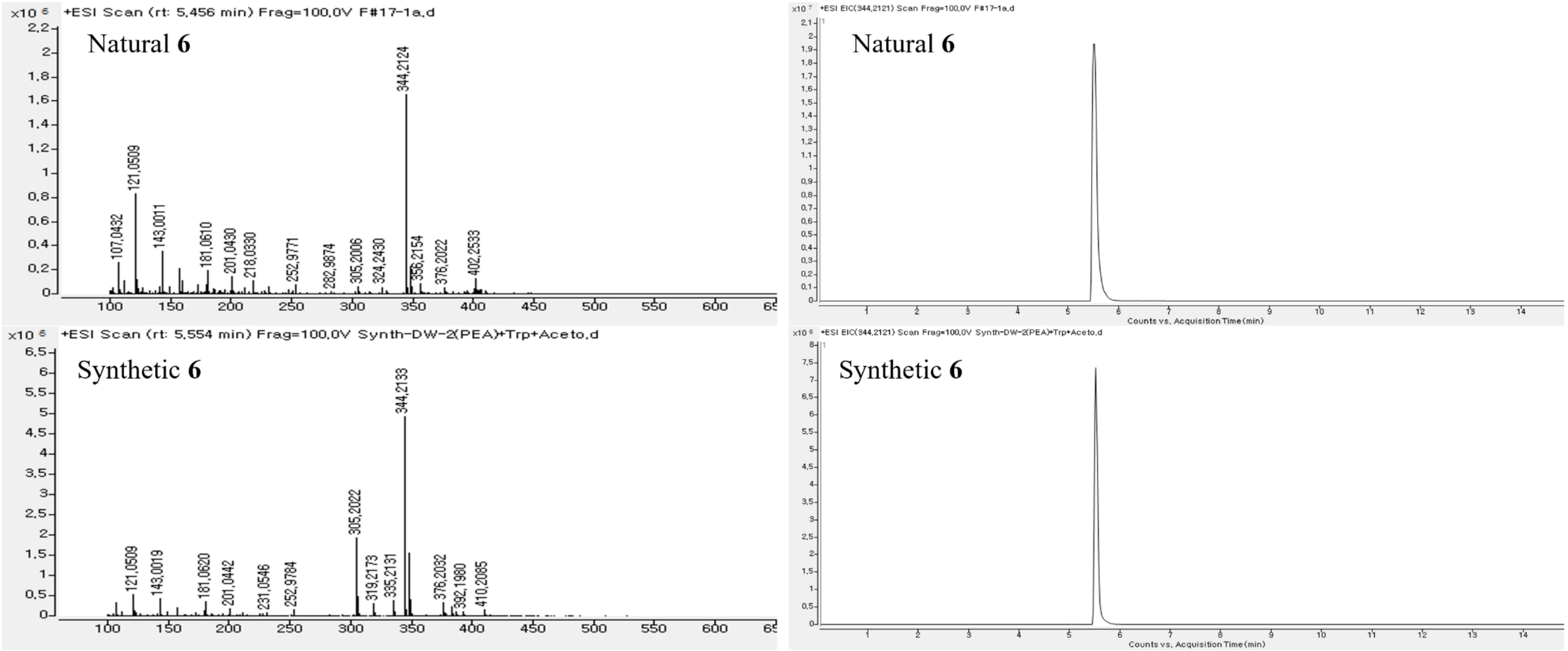

**Figure S45**. HRESIMS and HPLC chromatogram of **6**.

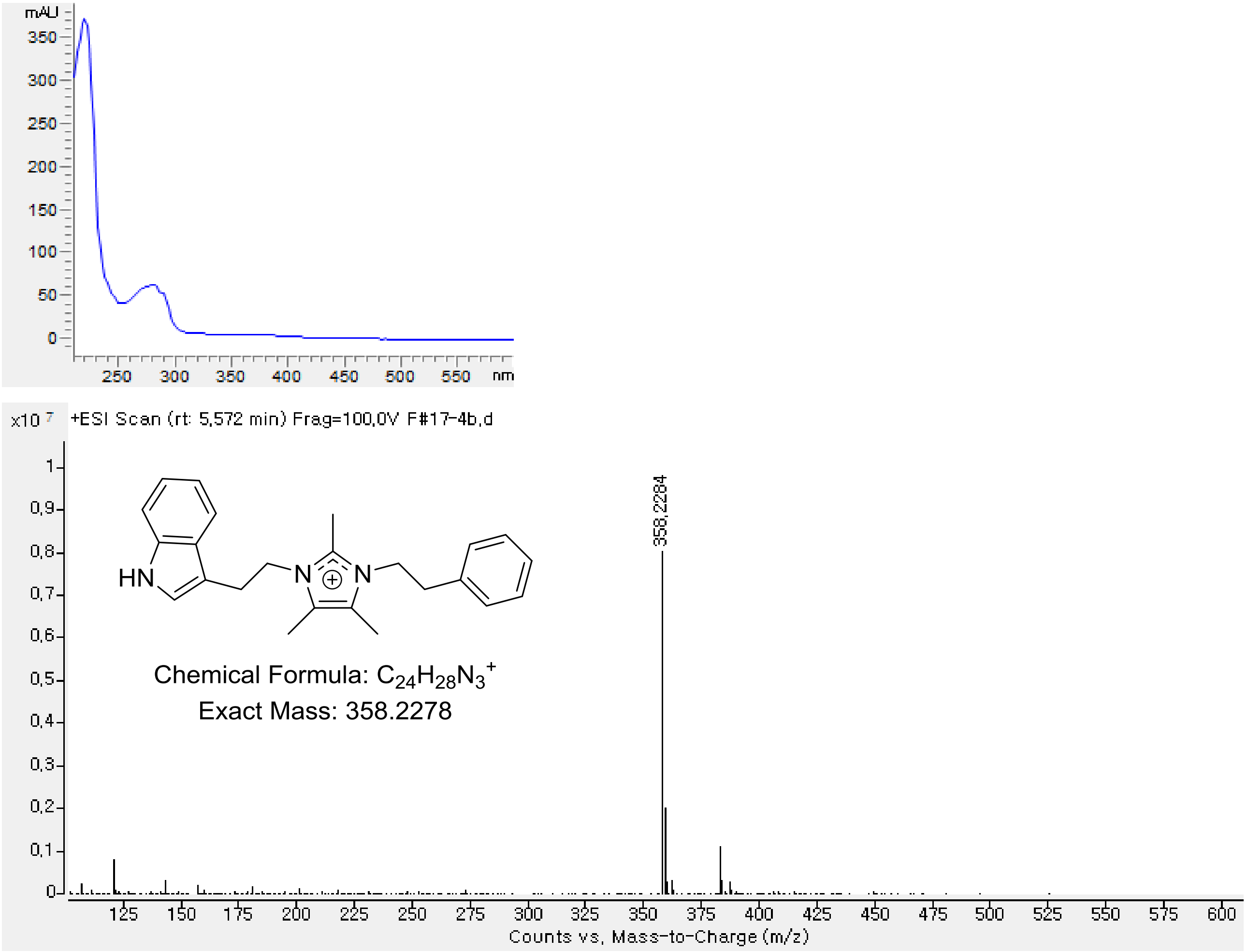

**Figure S46**. UV-Vis and HRESIMS spectrum of **7**.

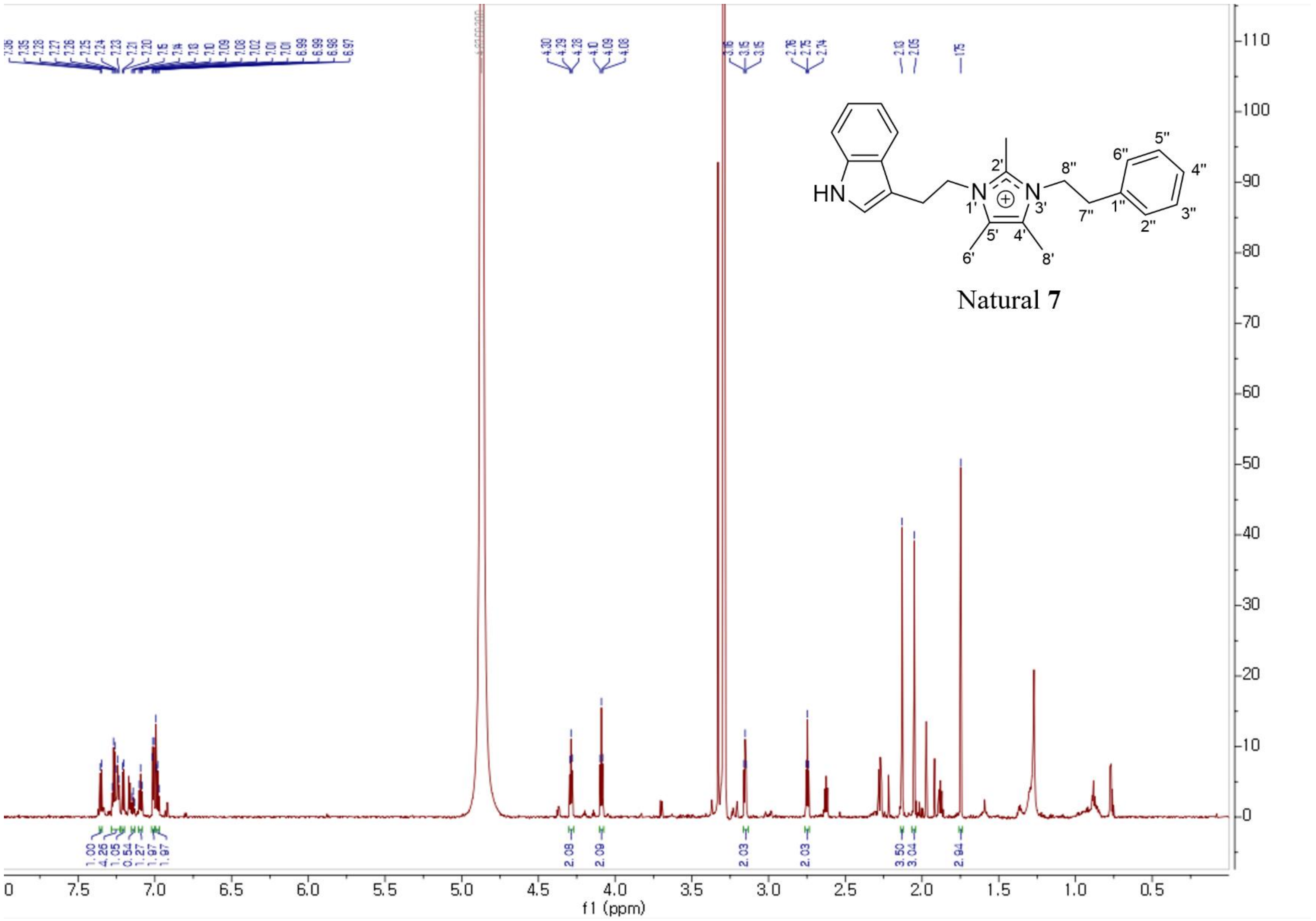

**Figure S47**. The [1]H NMR spectrum of natural **7** in methanol-$d_4$.

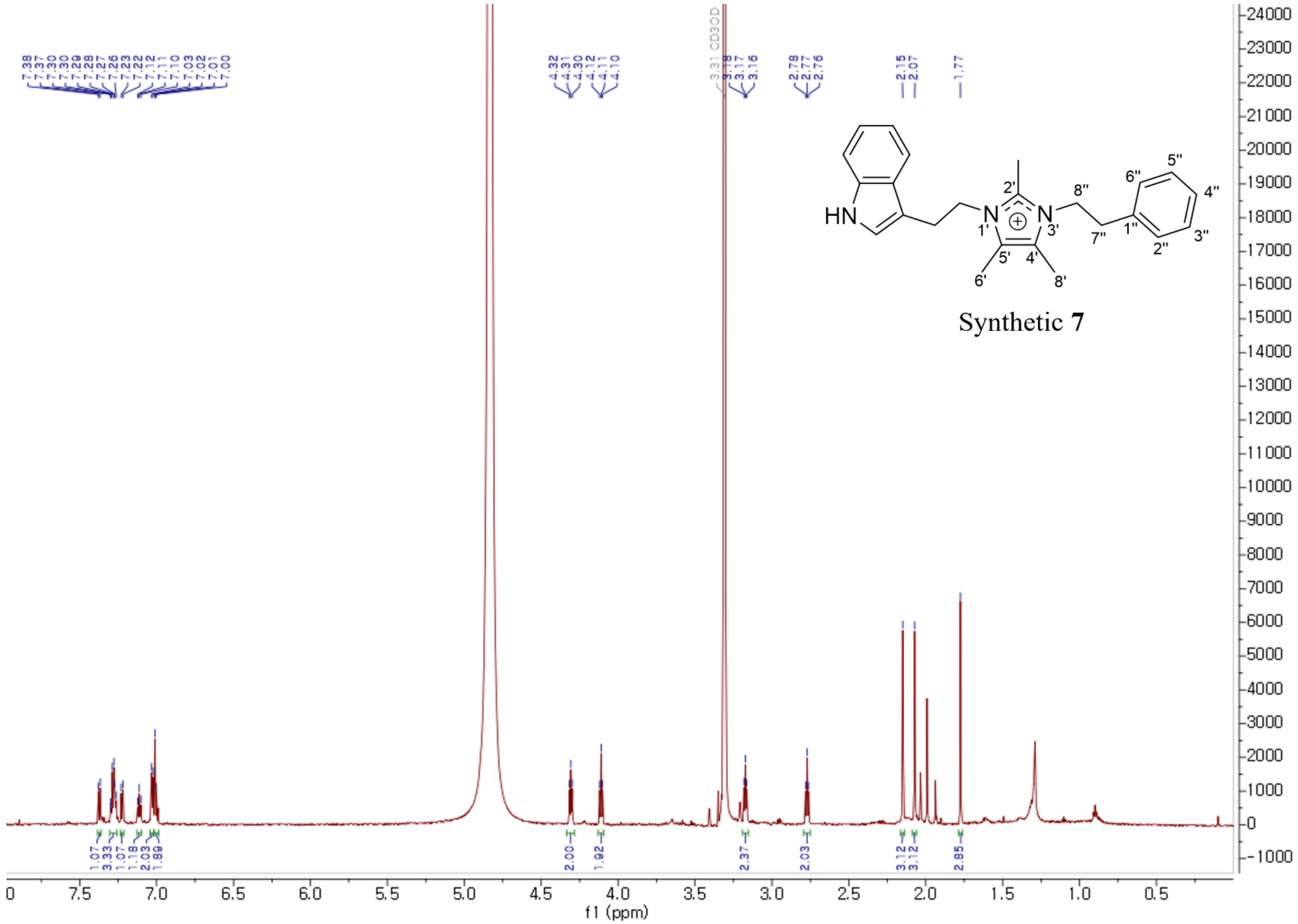

**Figure S48**. The $^{1}$H NMR spectrum of synthetic **7** in methanol-$d_4$.

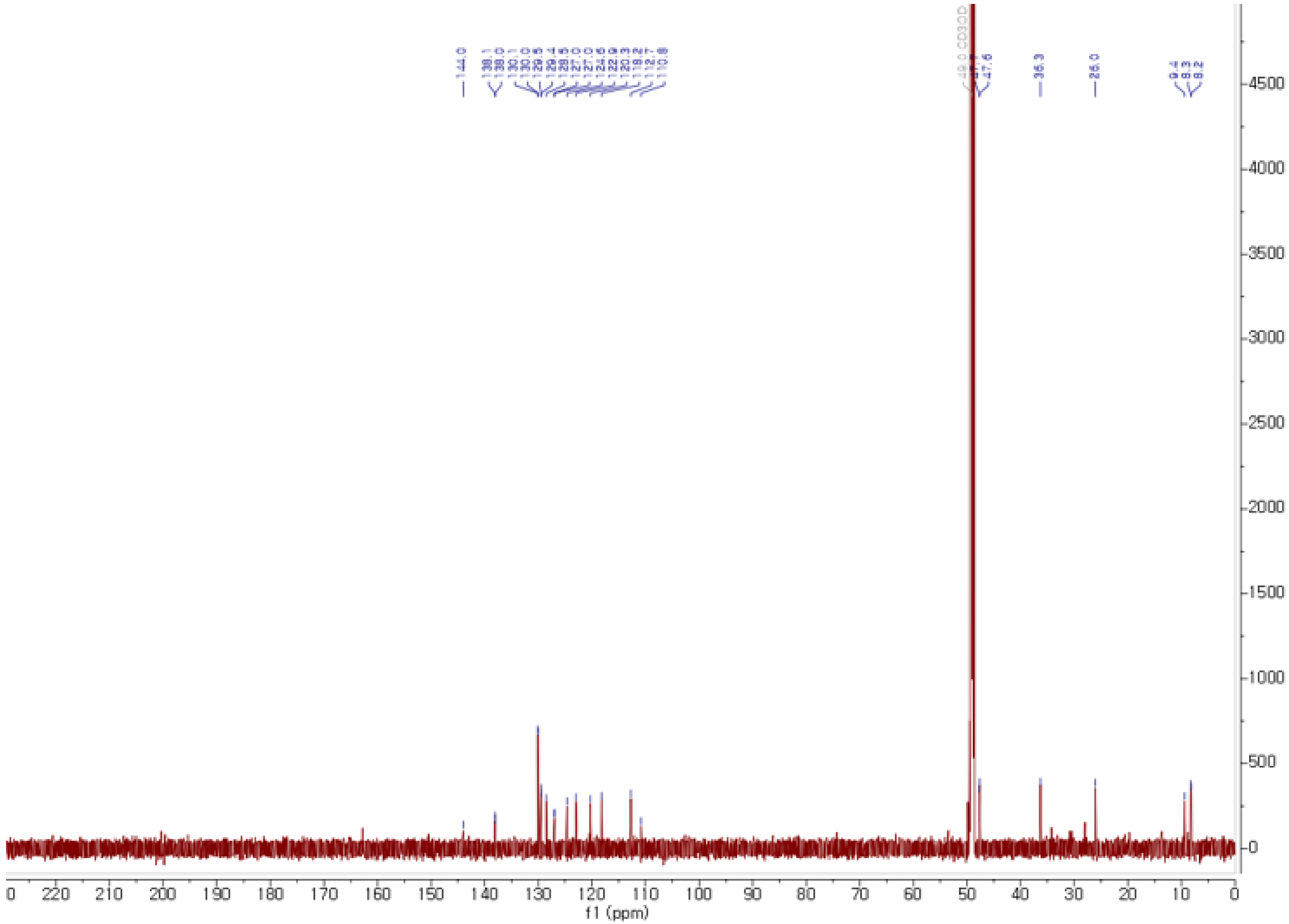

**Figure S49**. The $^{13}C$ NMR spectrum of **7** in methanol-$d_4$.

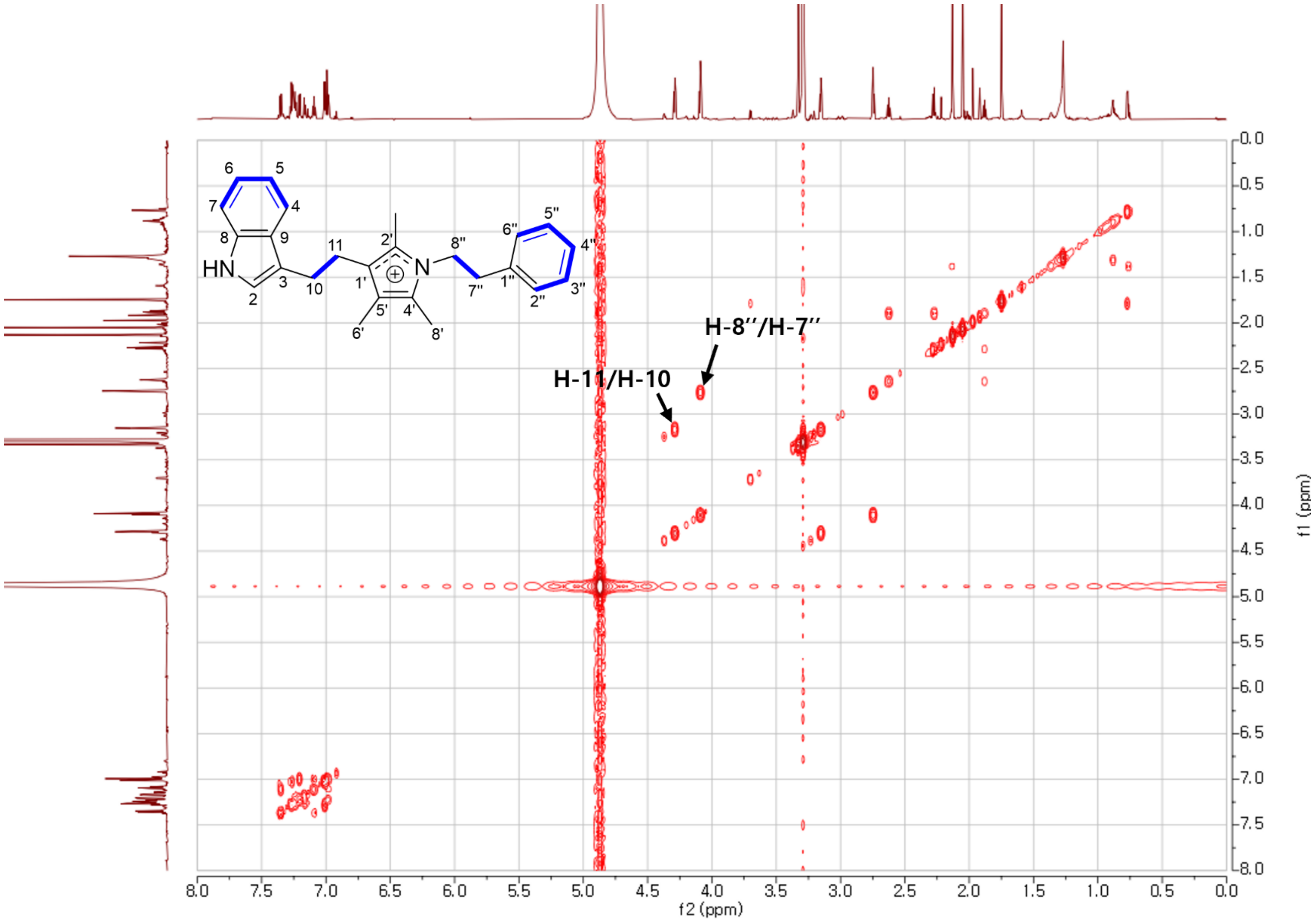

**Figure S50**. The COSY spectrum of **7** in methanol-$d_4$.

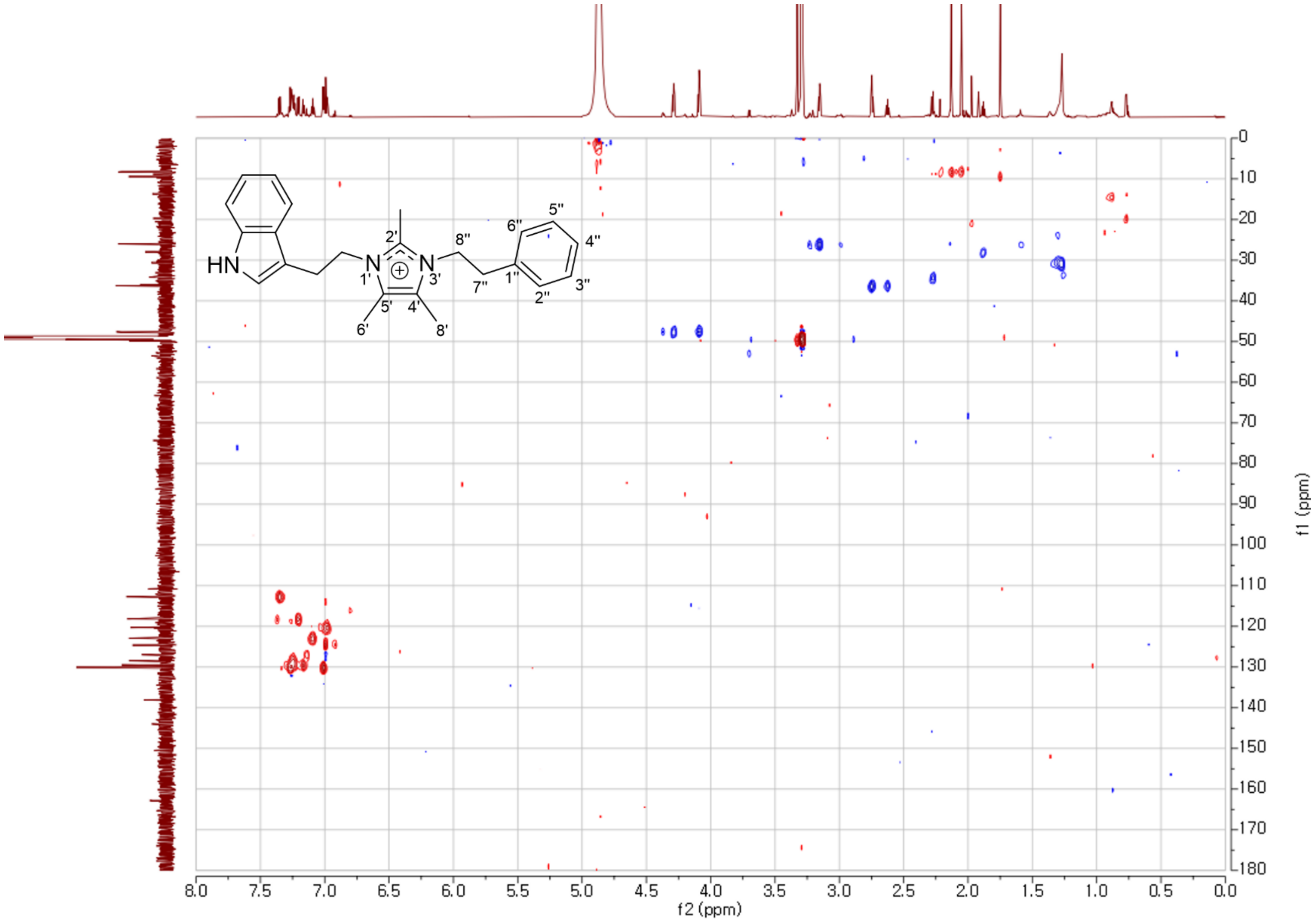

**Figure S51**. The HSQC spectrum of **7** in methanol-$d_4$.

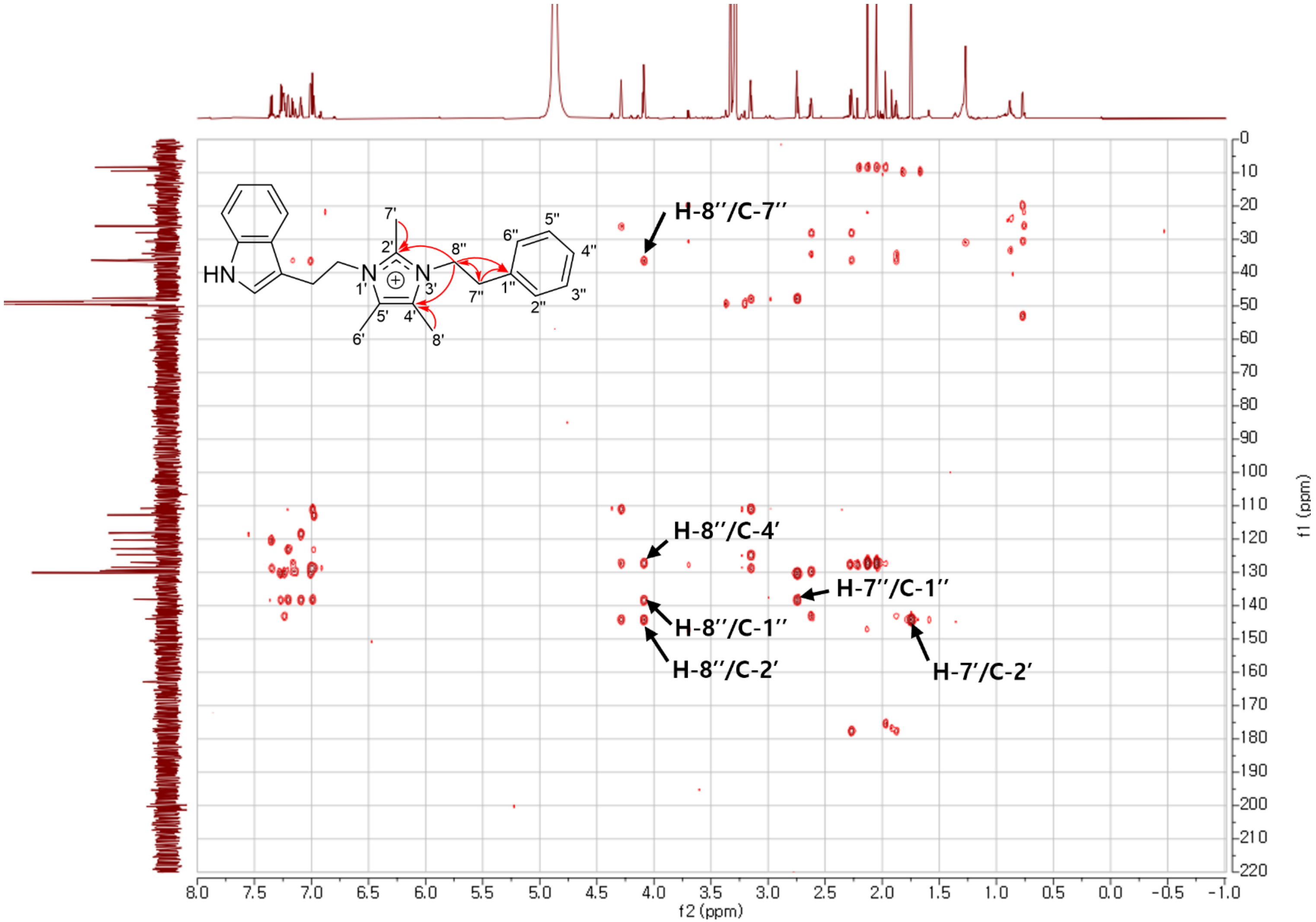

**Figure S52**. The HMBC spectrum of **7** in methanol-$d_4$.

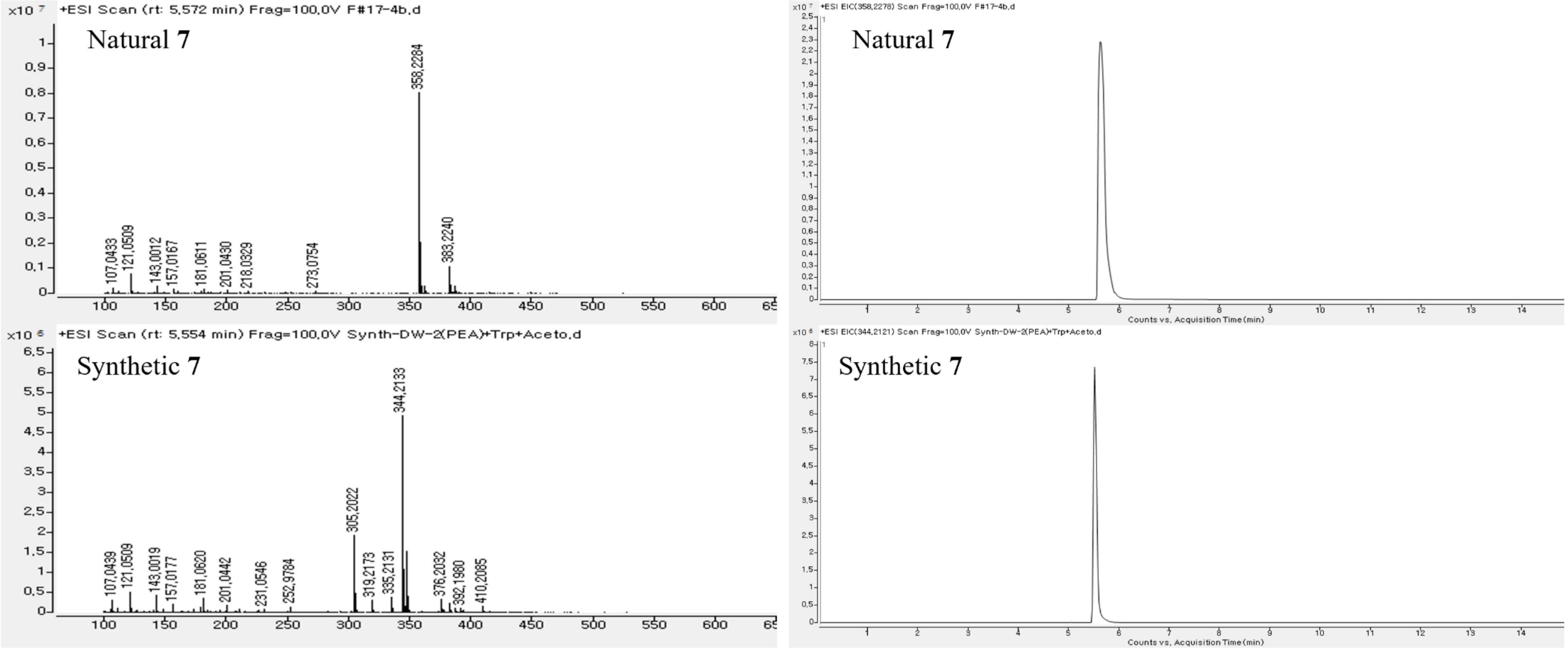

**Figure S53**. HRESIMS and HPLC chromatogram of **7**.

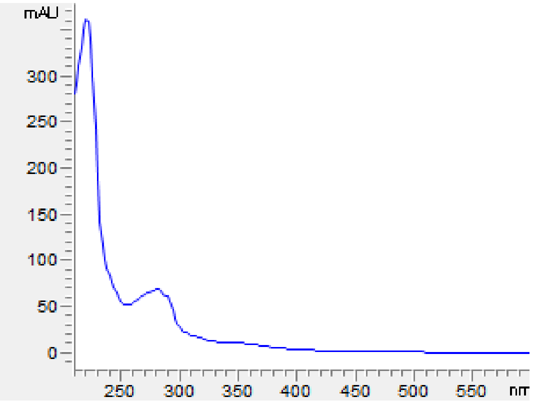

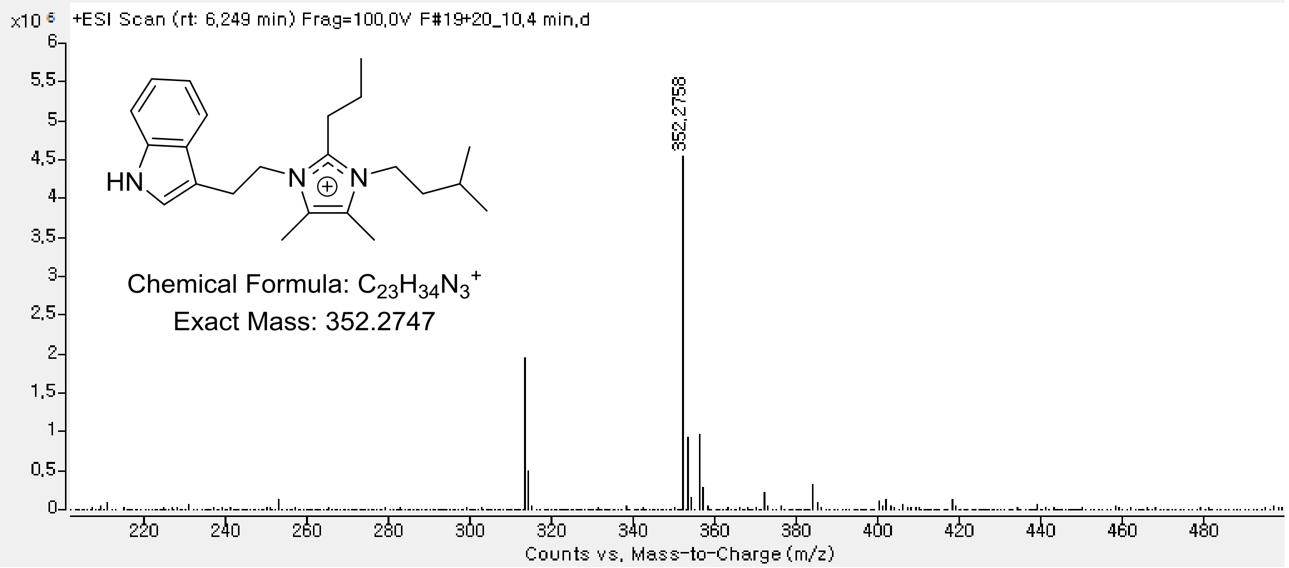

**Figure S54**. UV-Vis and HRESIMS spectrum of **8**.

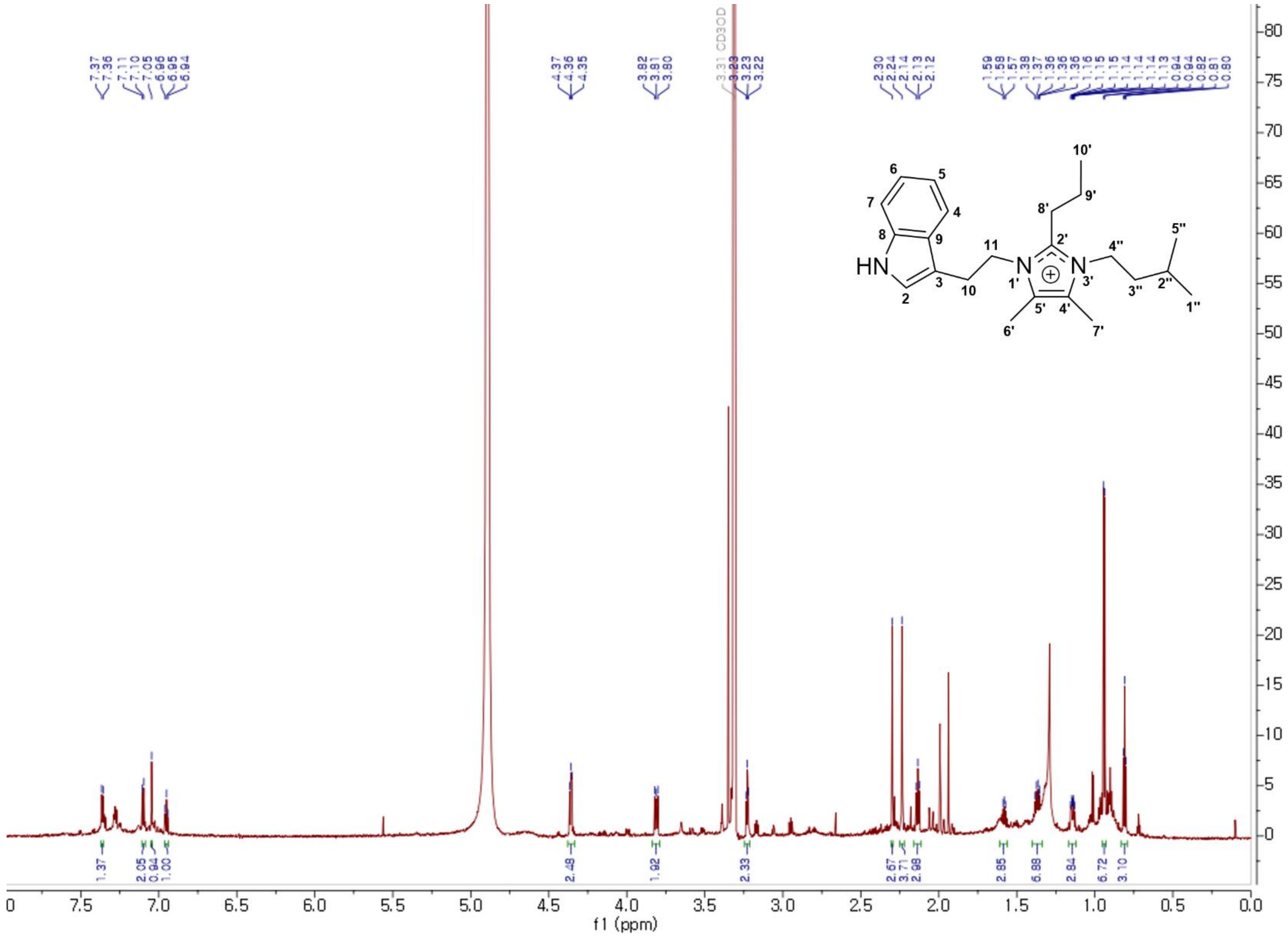

**Figure S55**. The $^{1}$H NMR spectrum of **8** in methanol-$d_4$.

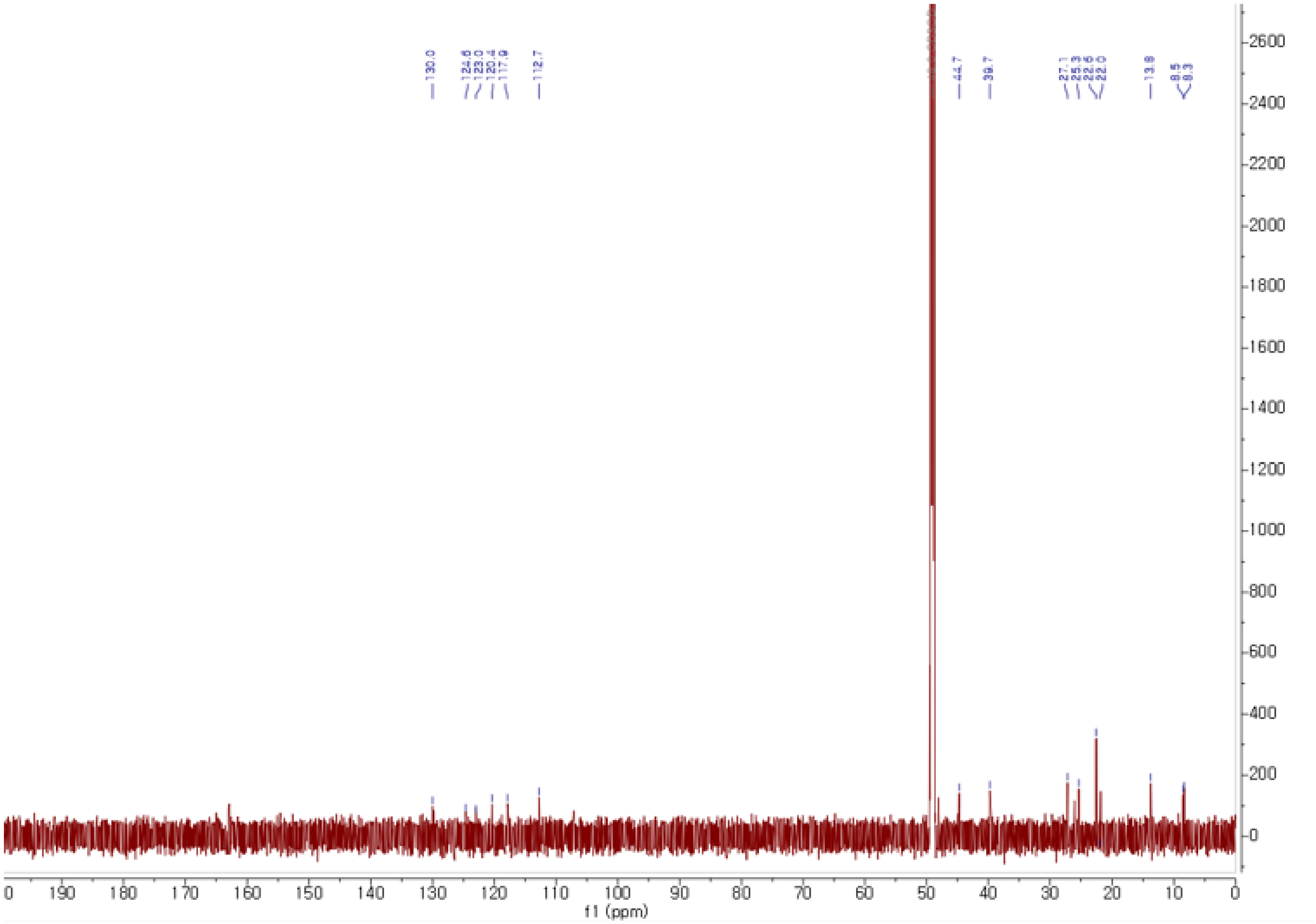

**Figure S56**. The $^{13}$C NMR spectrum of **8** in methanol-$d_4$.

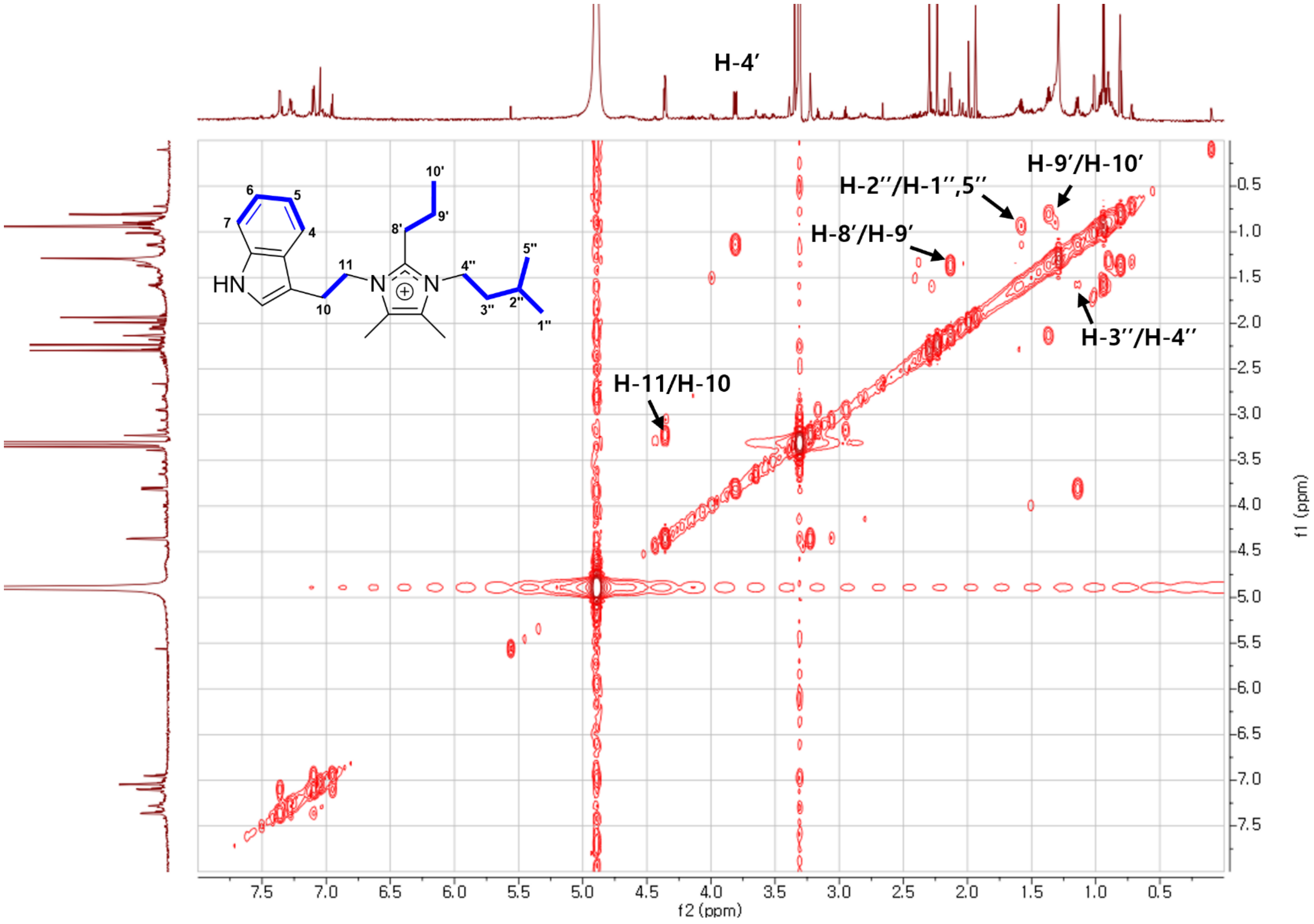

**Figure S57**. The COSY spectrum of **8** in methanol-$d_4$.

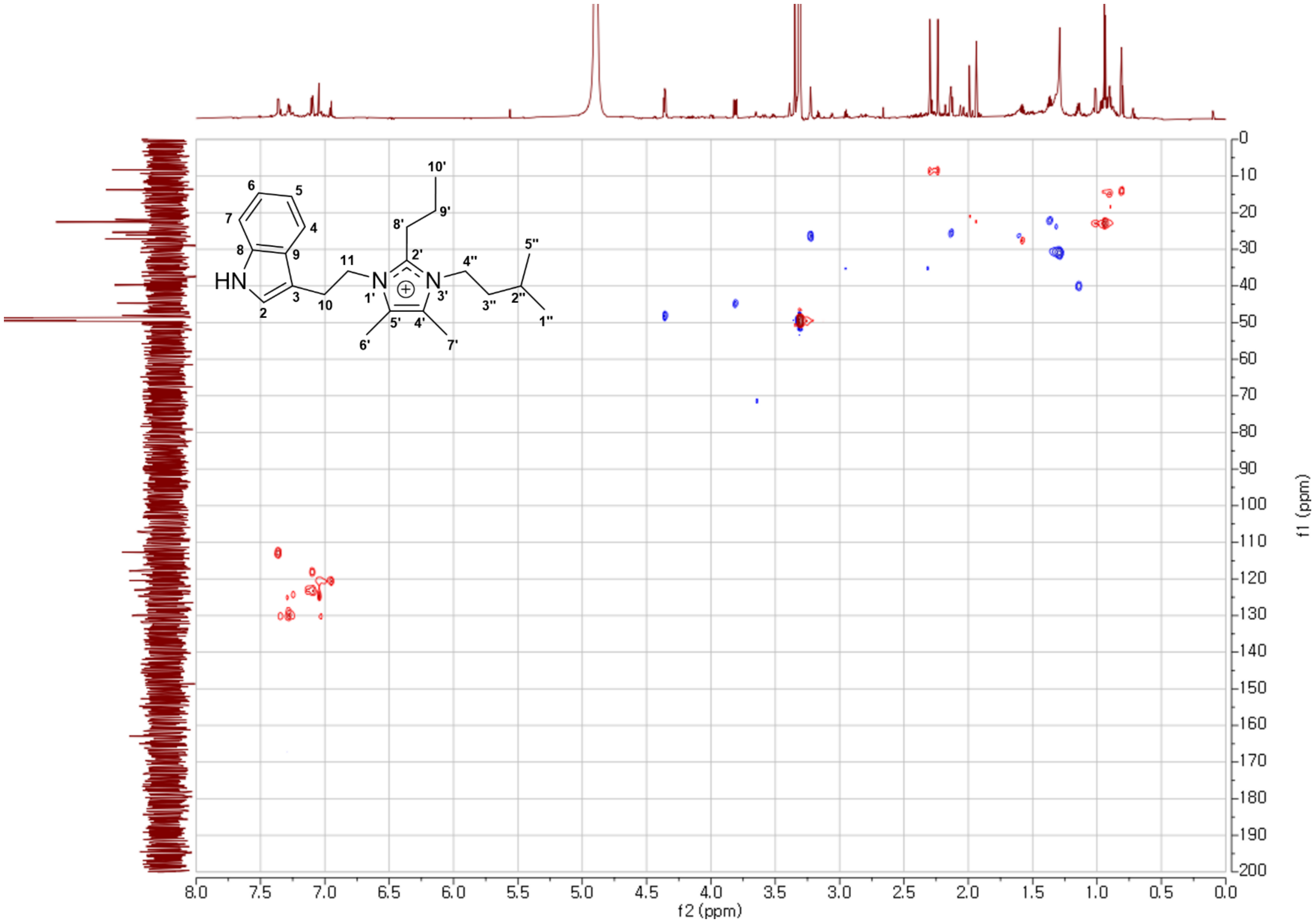

**Figure S58**. The HSQC spectrum of **8** in methanol-$d_4$.

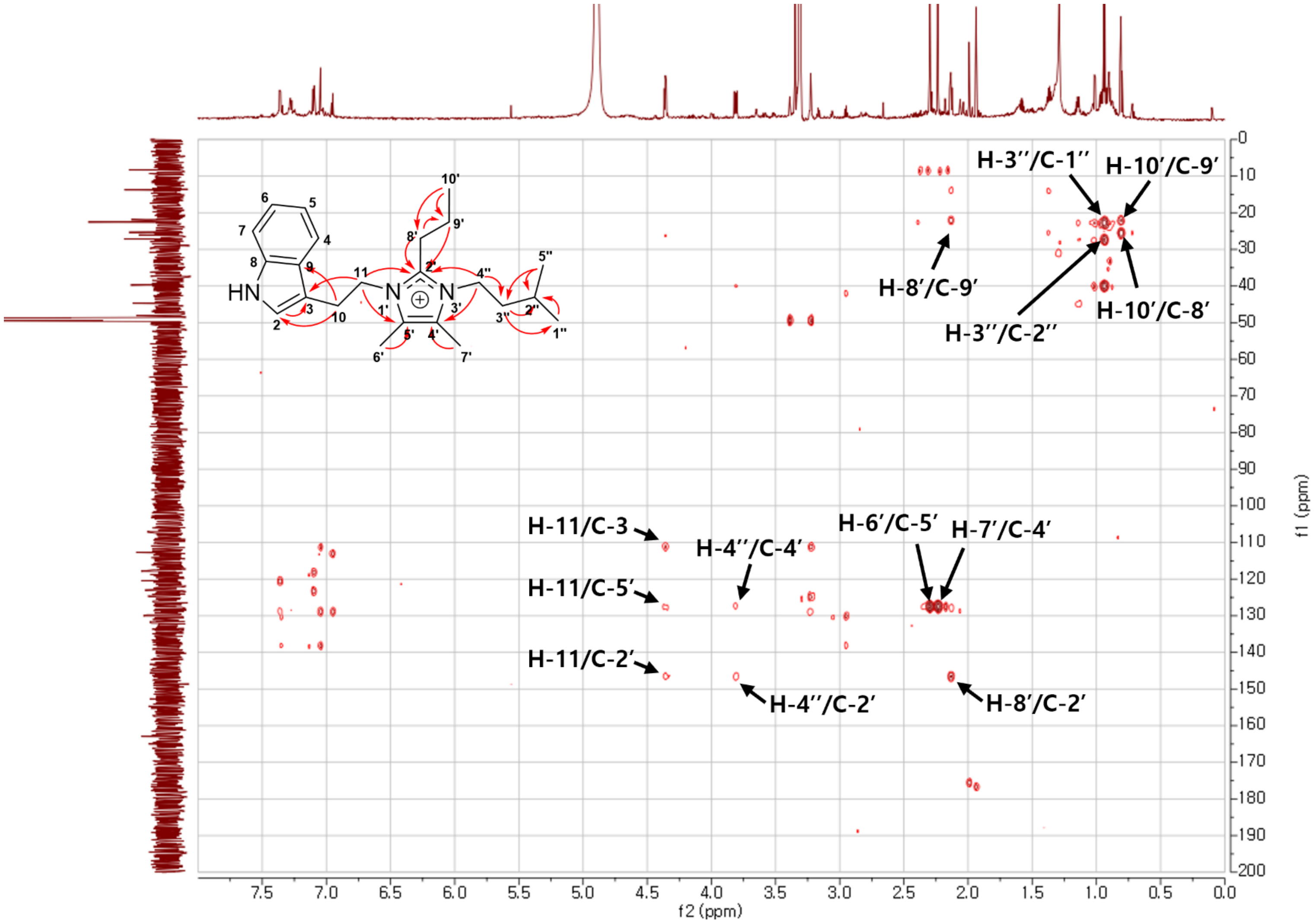

**Figure S59**. The HMBC spectrum of **8** in methanol-$d_4$.

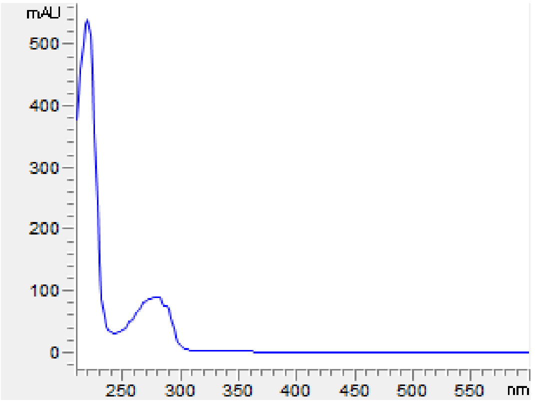

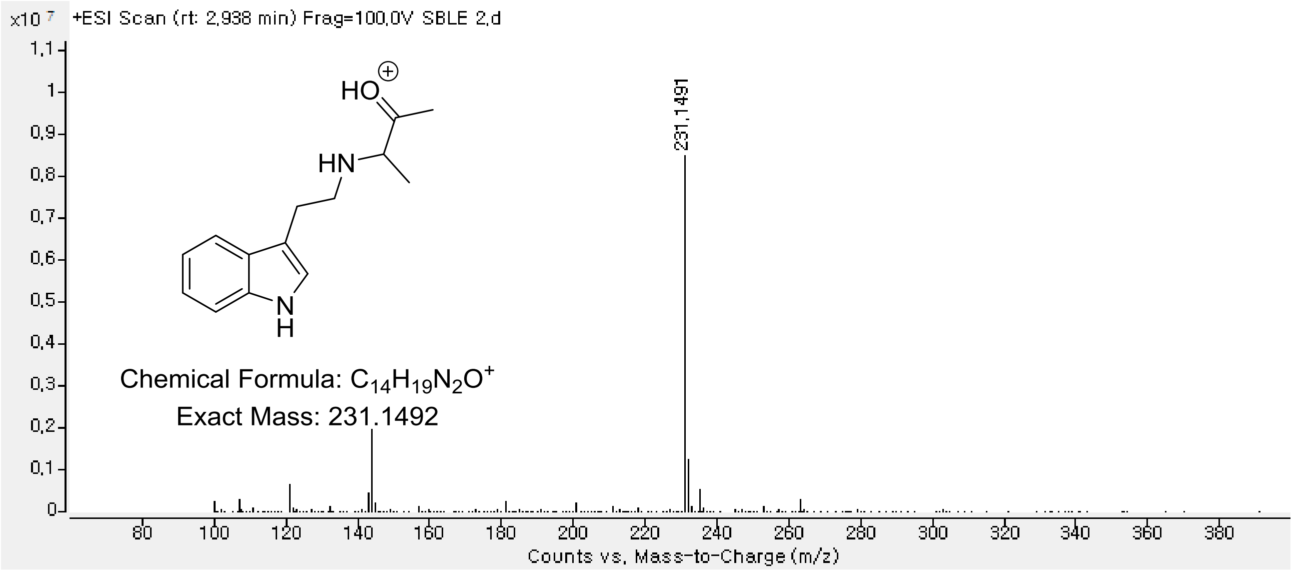

**Figure S60**. The HRESIMS spectrum of **9**.

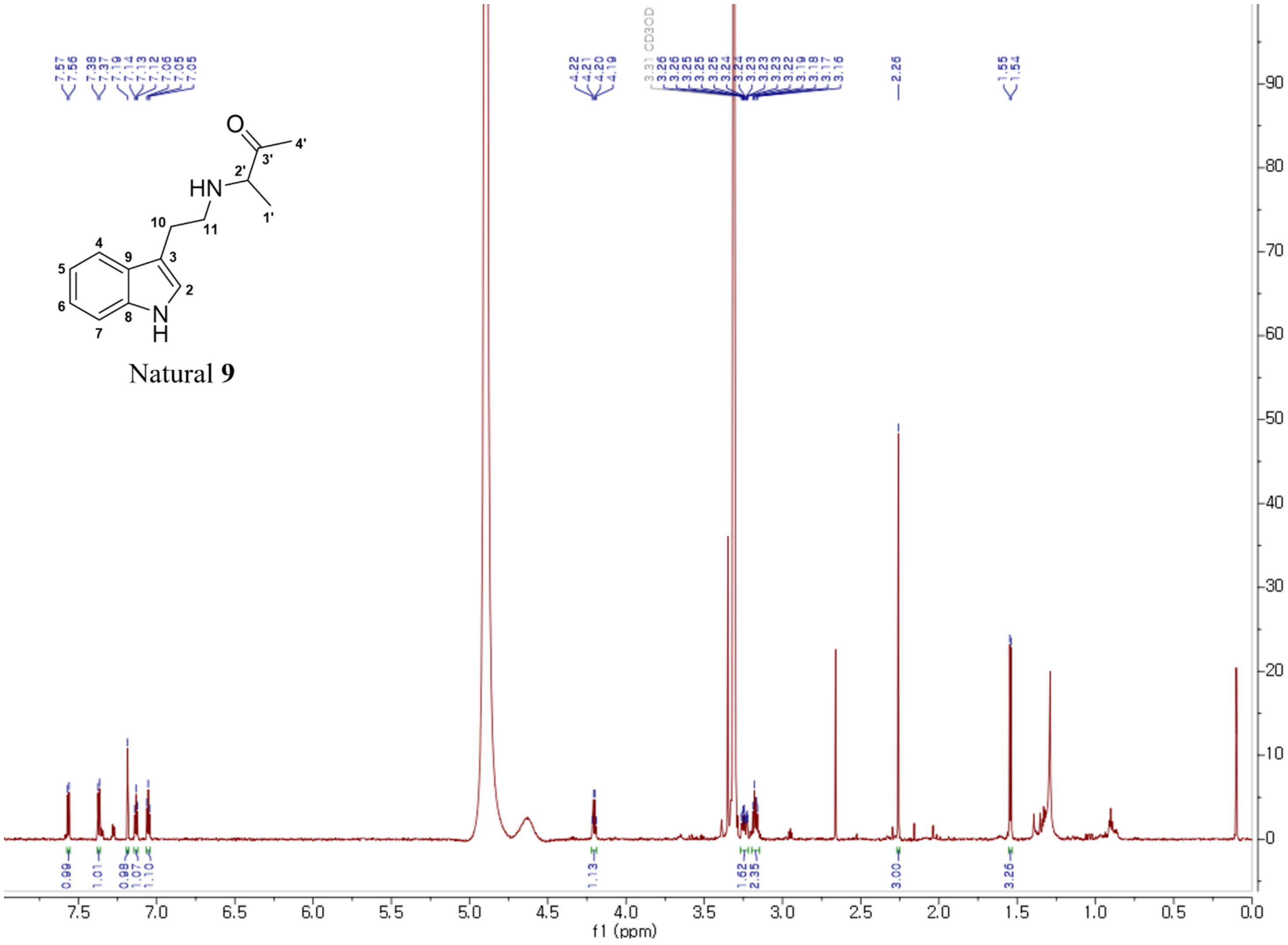

**Figure S61**. The $^{1}$H NMR spectrum of natural **9** in methanol-$d_4$.

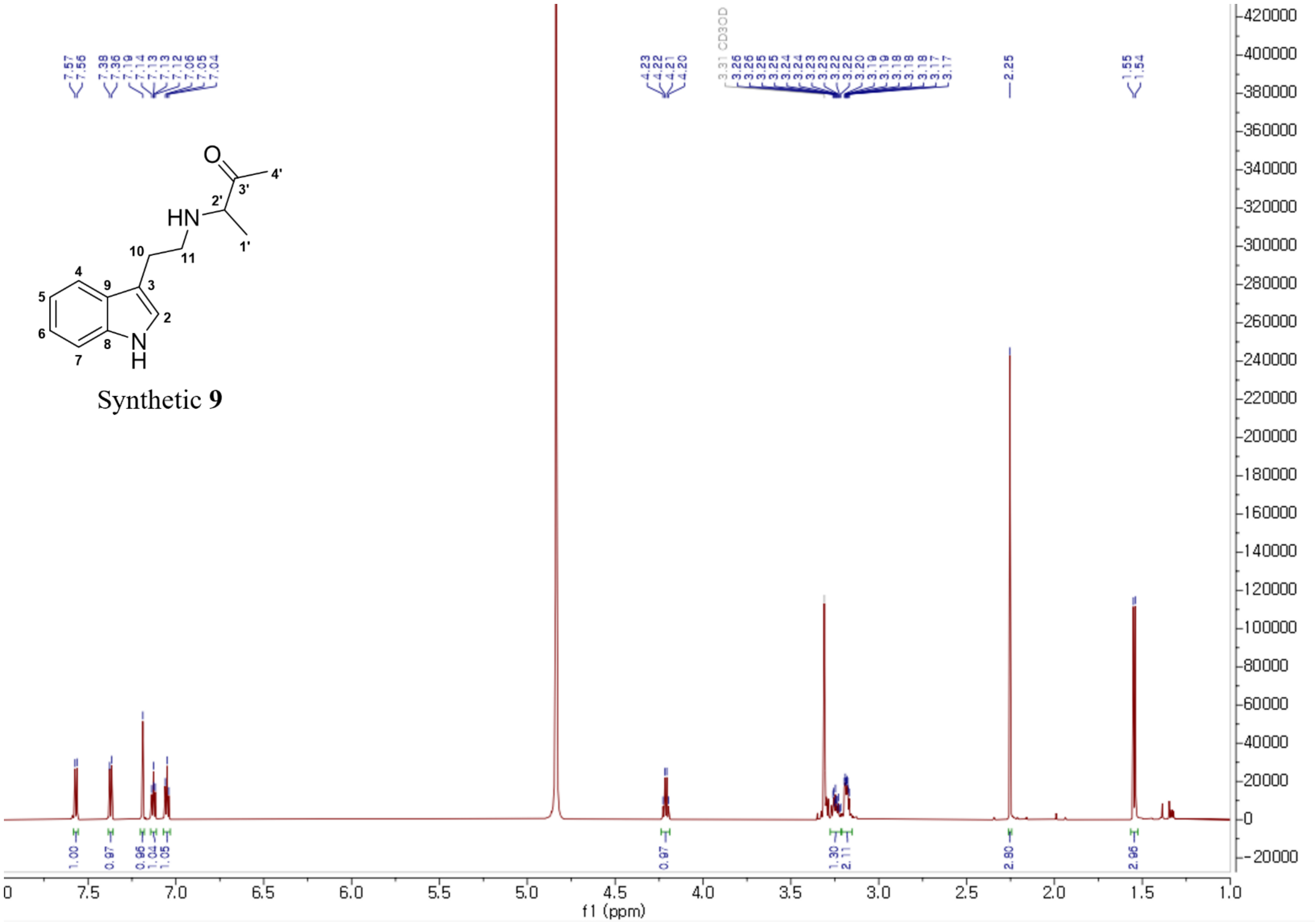

**Figure S62**. The $^{1}$H NMR spectrum of synthetic **9** in methanol-$d_4$.

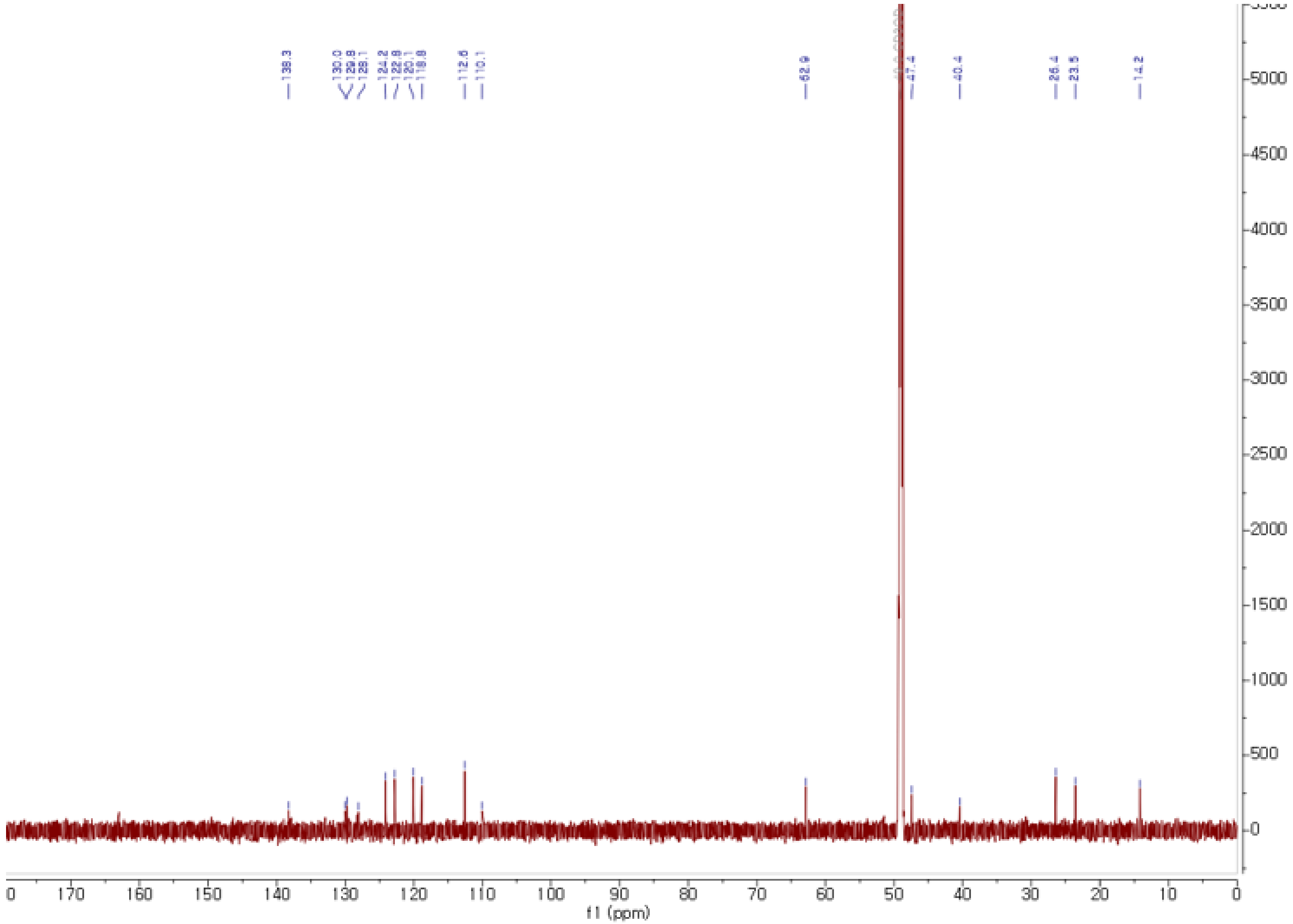

**Figure S63**. The [13]C NMR spectrum of **9** in methanol-$d_4$.

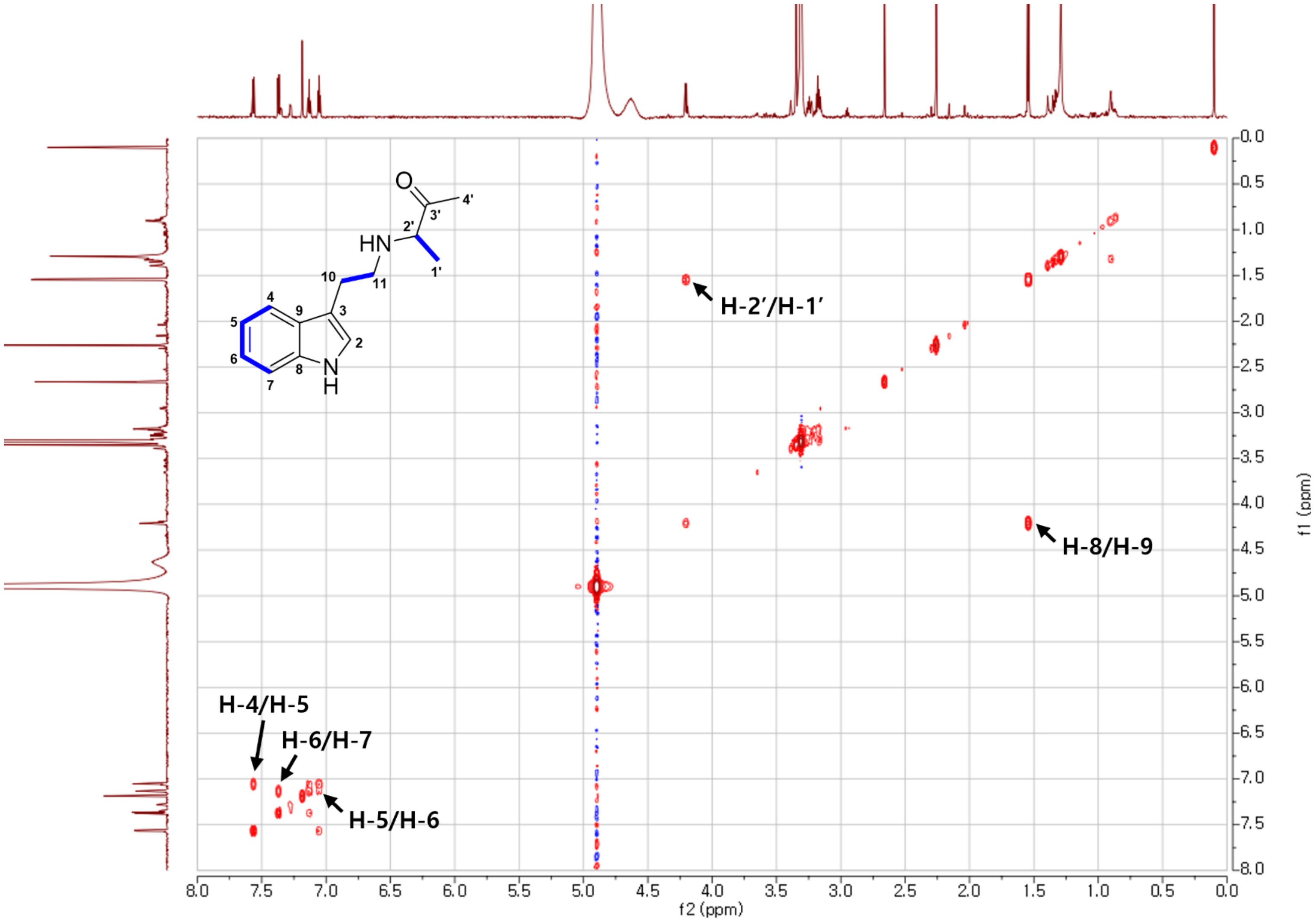

**Figure S64**. The COSY spectrum of **9** in methanol-$d_4$.

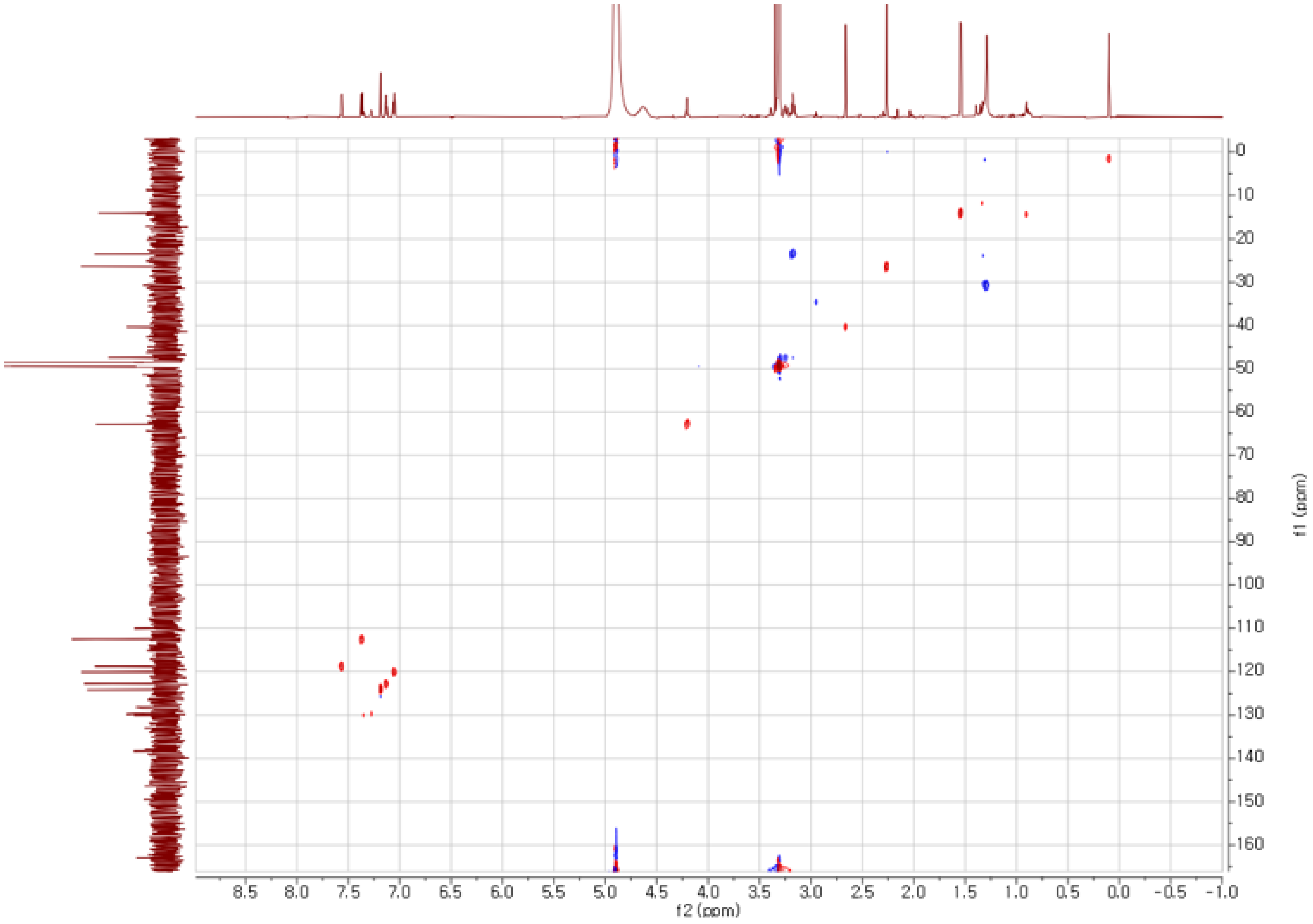

**Figure S65**. The HSQC spectrum of **9** in methanol-$d_4$.

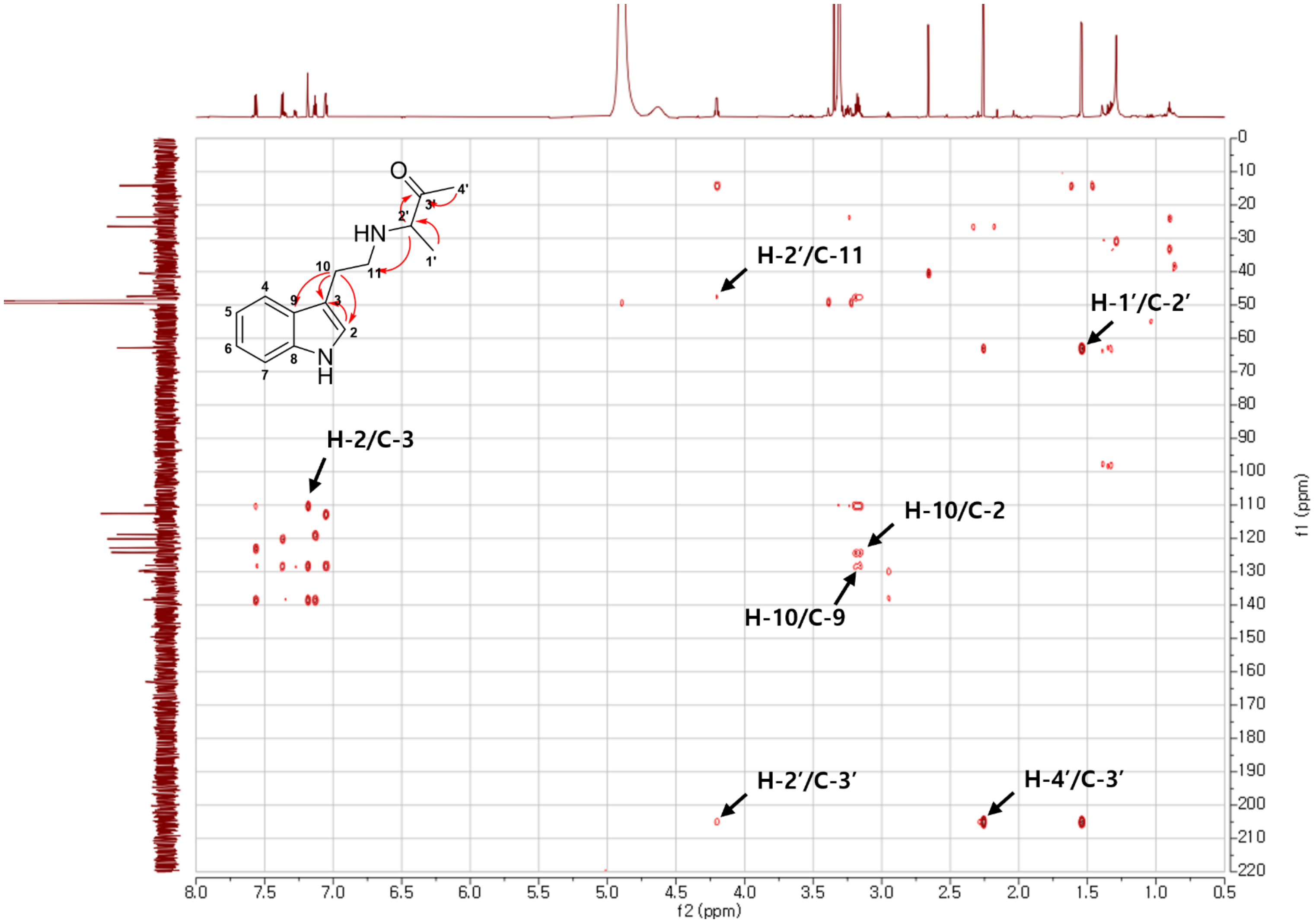

**Figure S66**. The HMBC spectrum of **9** in methanol-$d_4$.

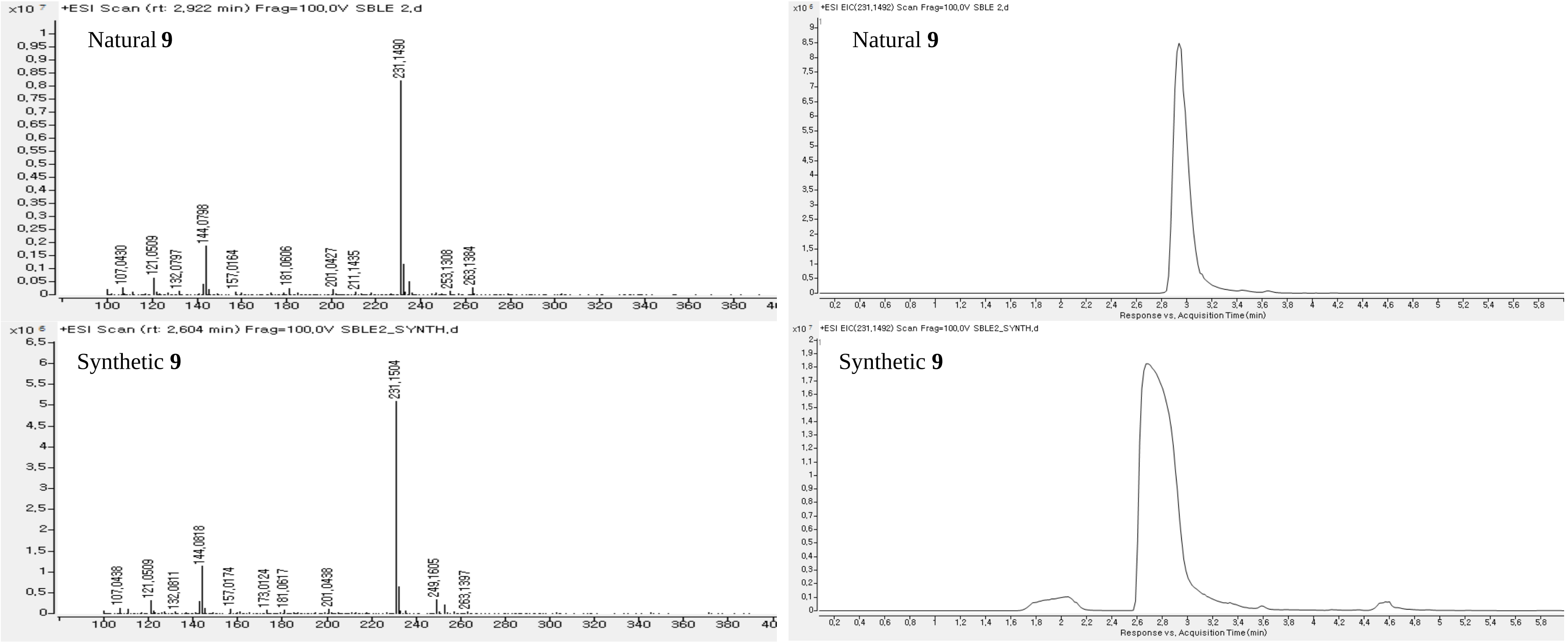

**Figure S67.** HRESIMS and HPLC chromatogram of **9**.

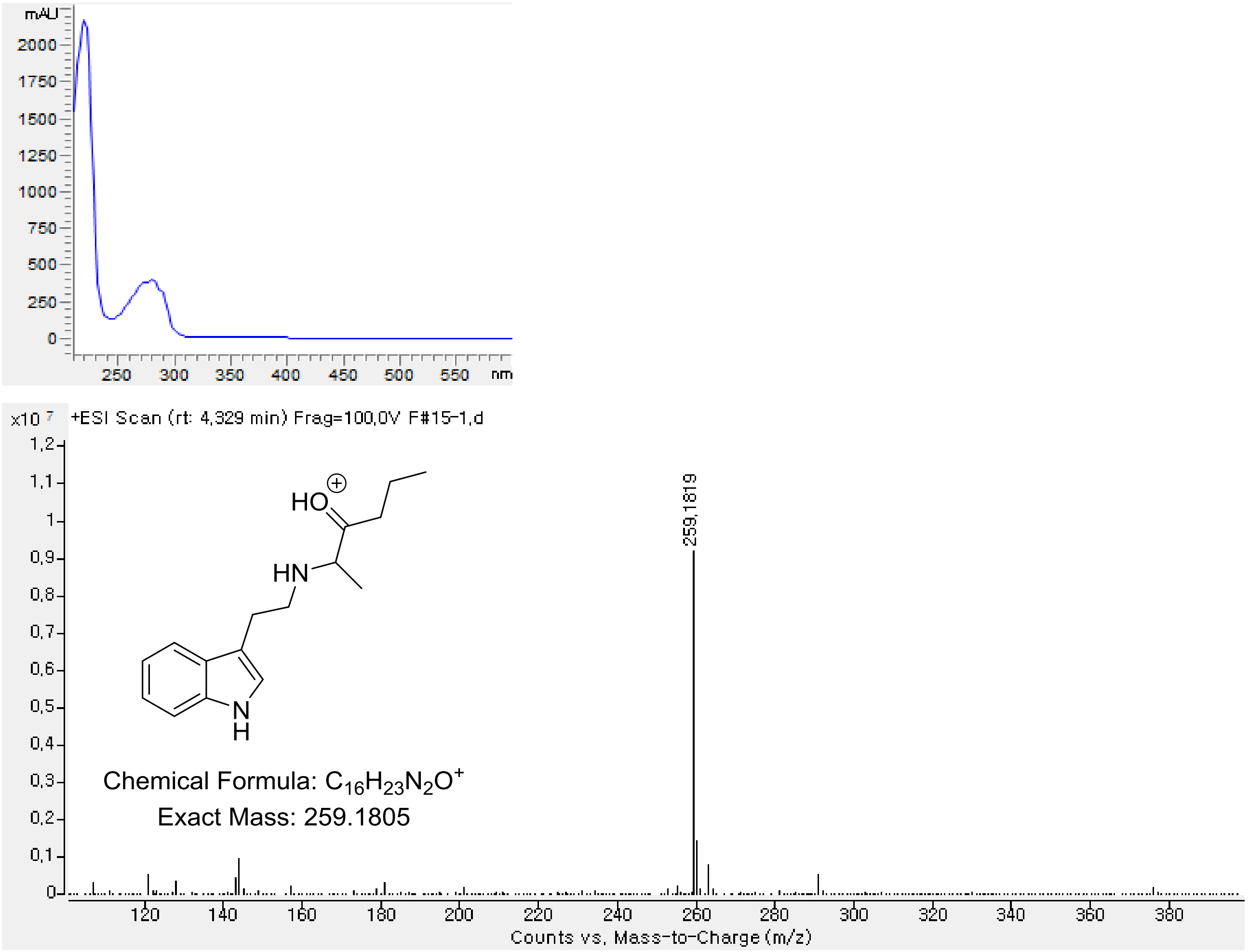

**Figure S68**. The HRESIMS spectrum of **10**.

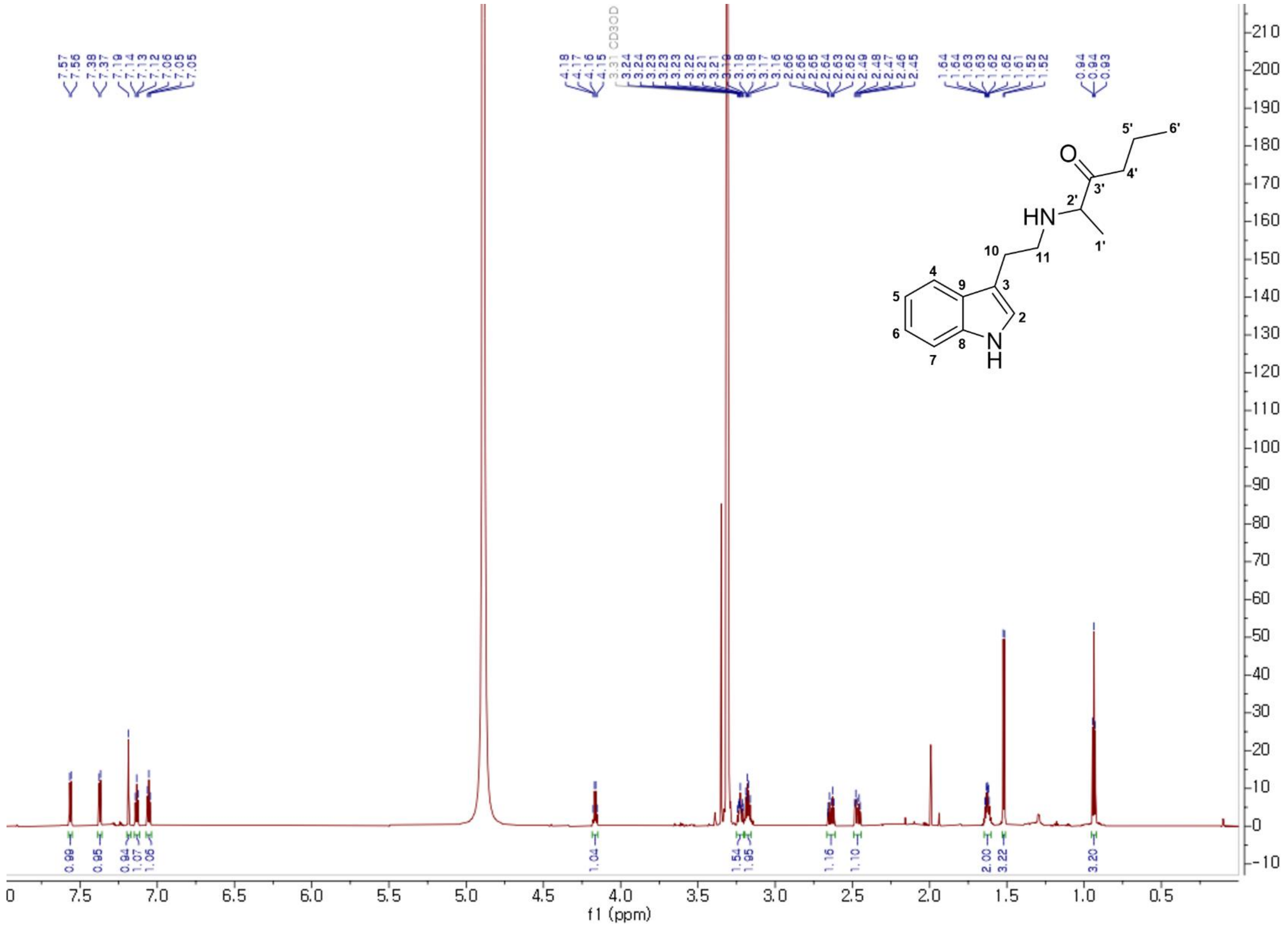

**Figure S69**. The $^{1}H$ NMR spectrum of **10** in methanol-$d_4$.

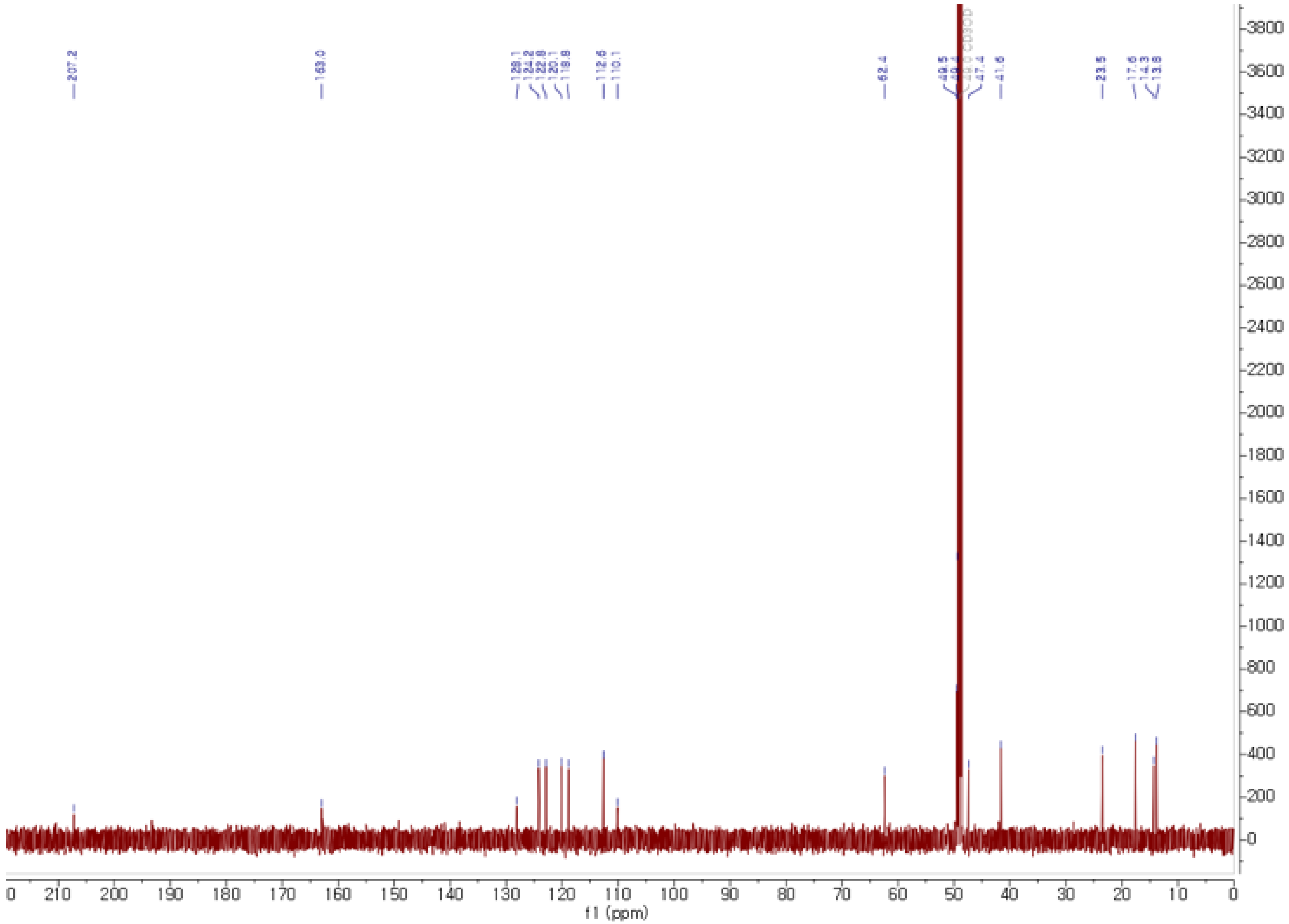

**Figure S70**. The $^{13}$C NMR spectrum of **10** in methanol-$d_4$.

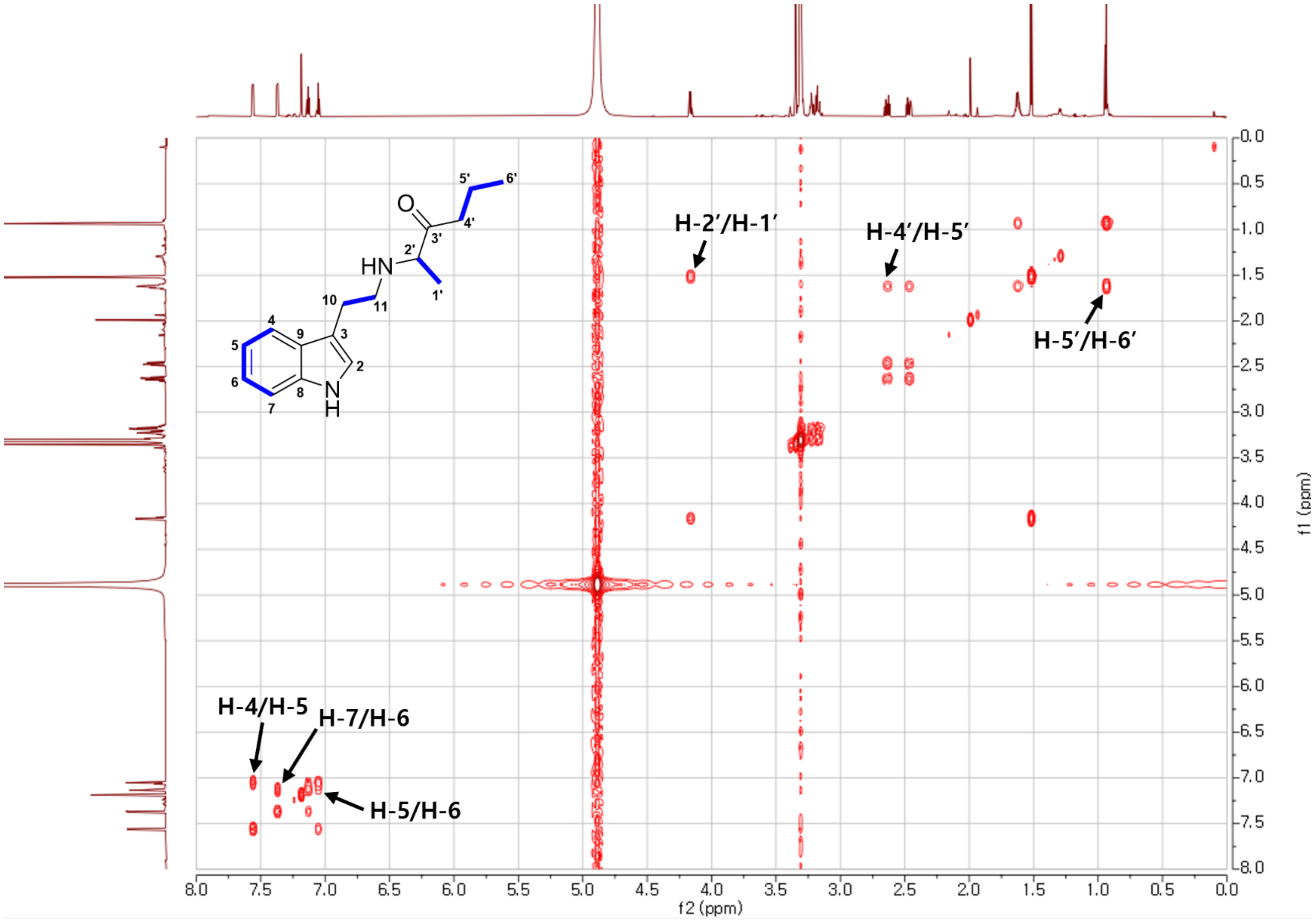

**Figure S71**. The COSY spectrum of **10** in methanol-$d_4$.

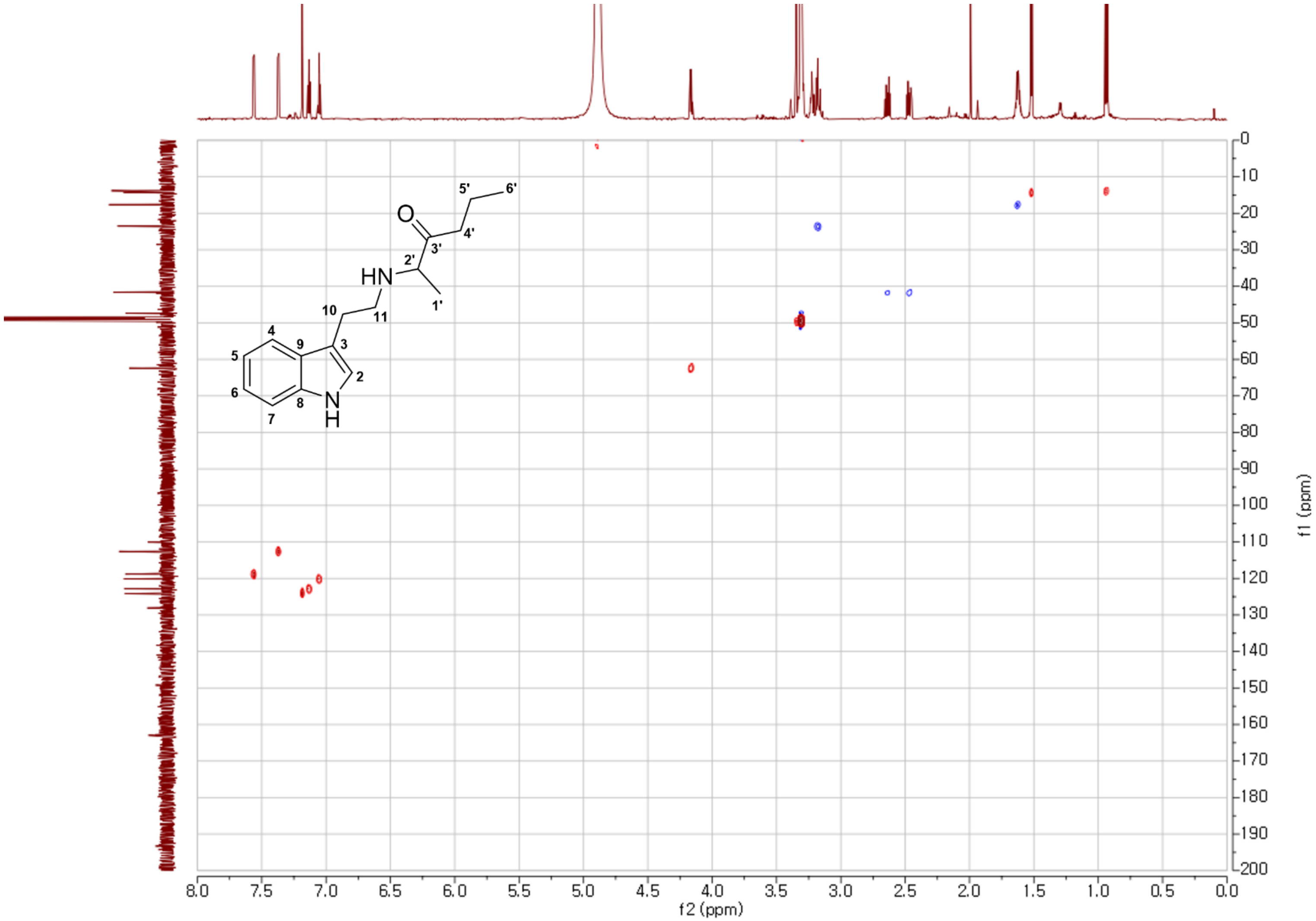

**Figure S72**. The HSQC spectrum of **10** in methanol-$d_4$.

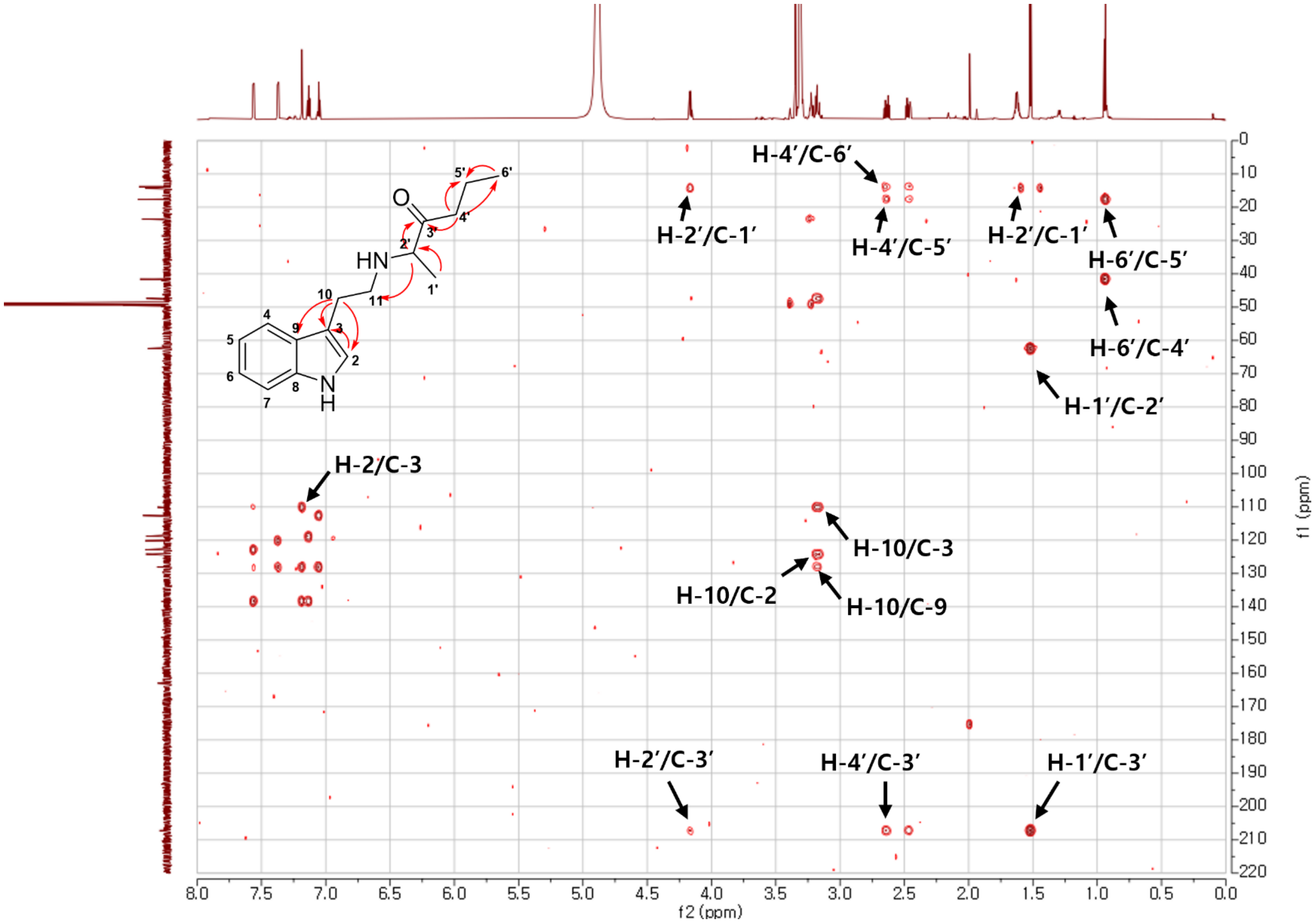

**Figure S73**. The HMBC spectrum of **10** in methanol-$d_4$.

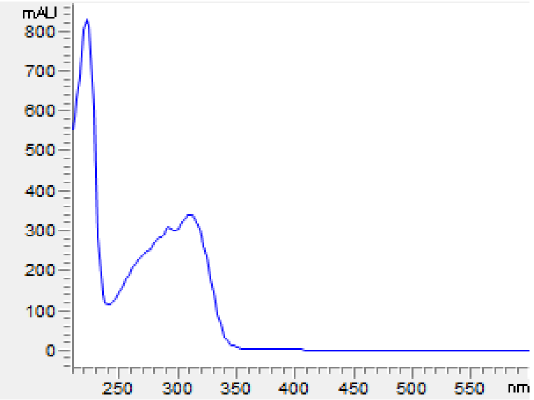

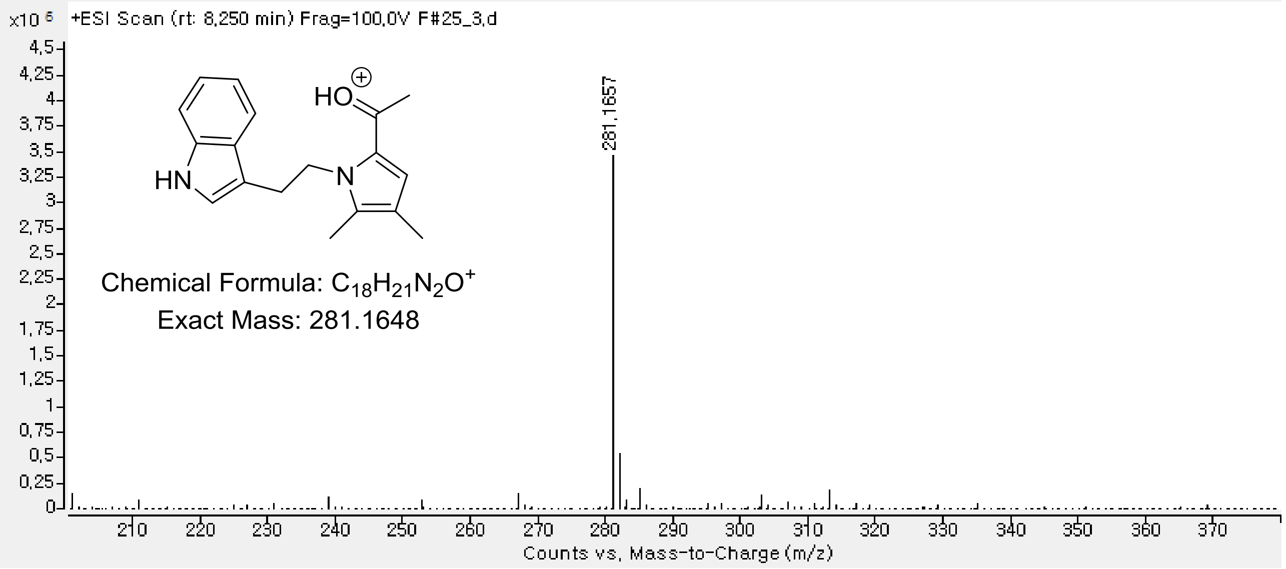

**Figure S74**. The HRESIMS spectrum of **12**.

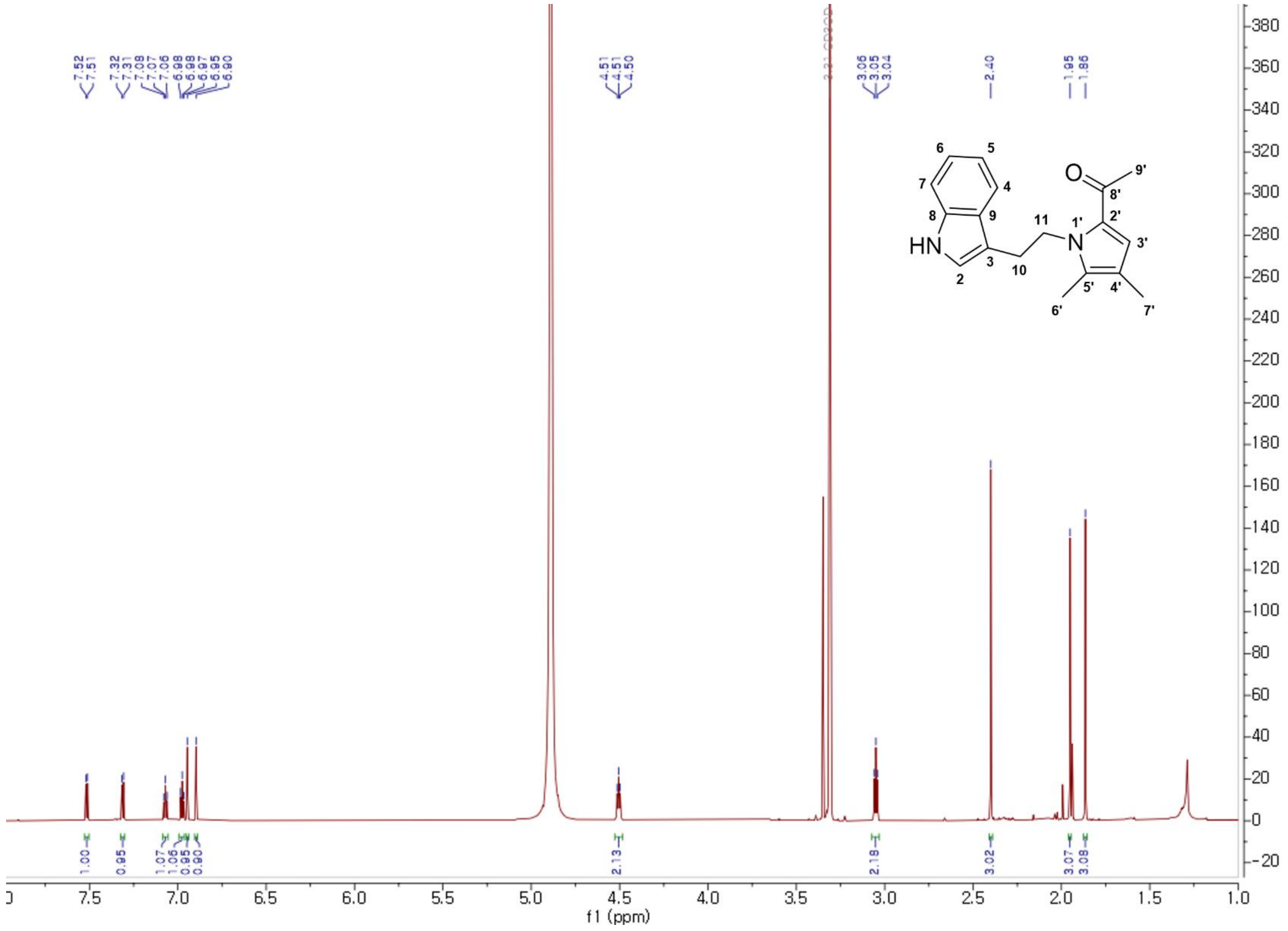

**Figure S75**. The $^1$H NMR spectrum of **12** in methanol-$d_4$.

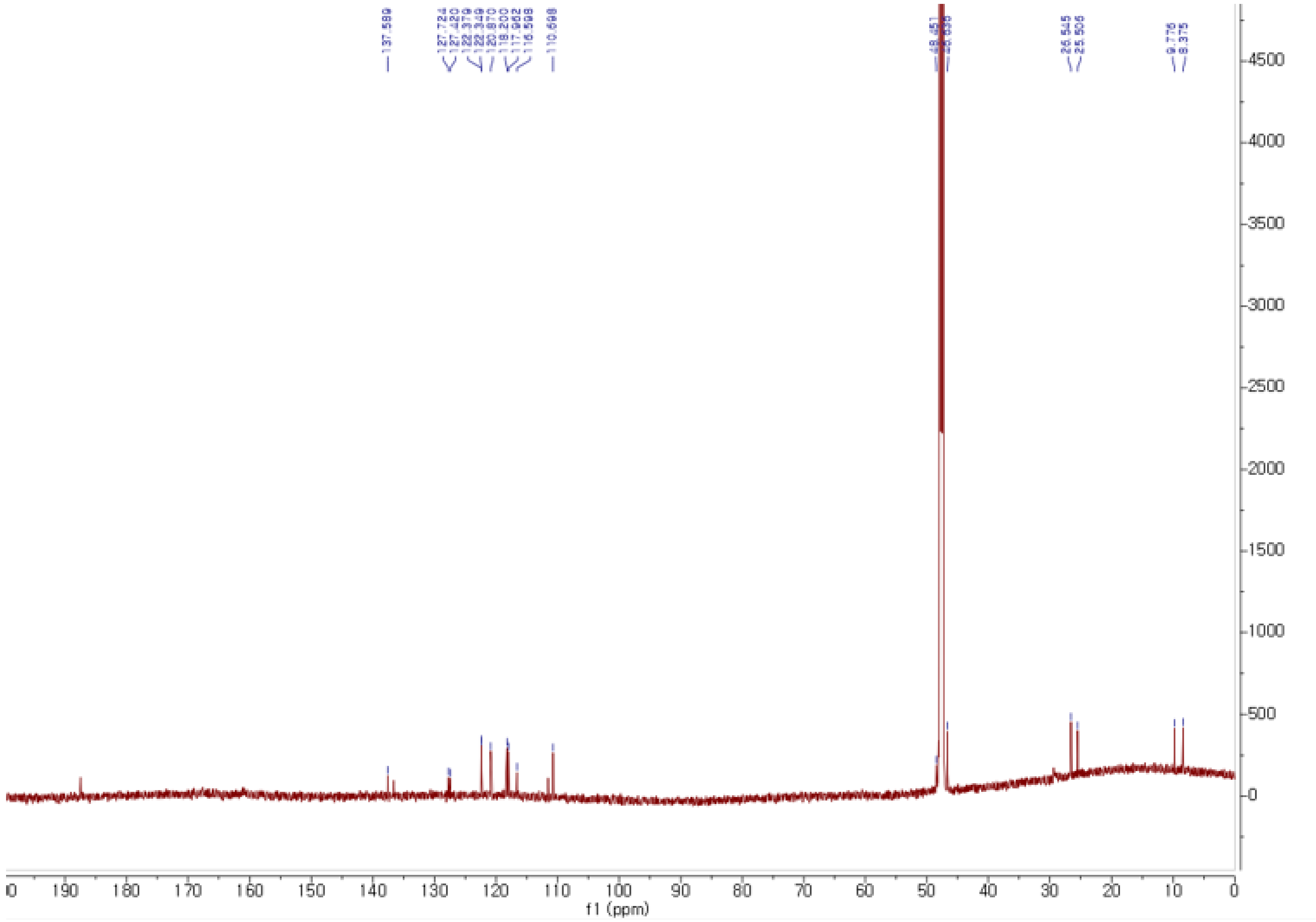

**Figure S76**. The $^{13}$C NMR spectrum of **12** in methanol-$d_4$.

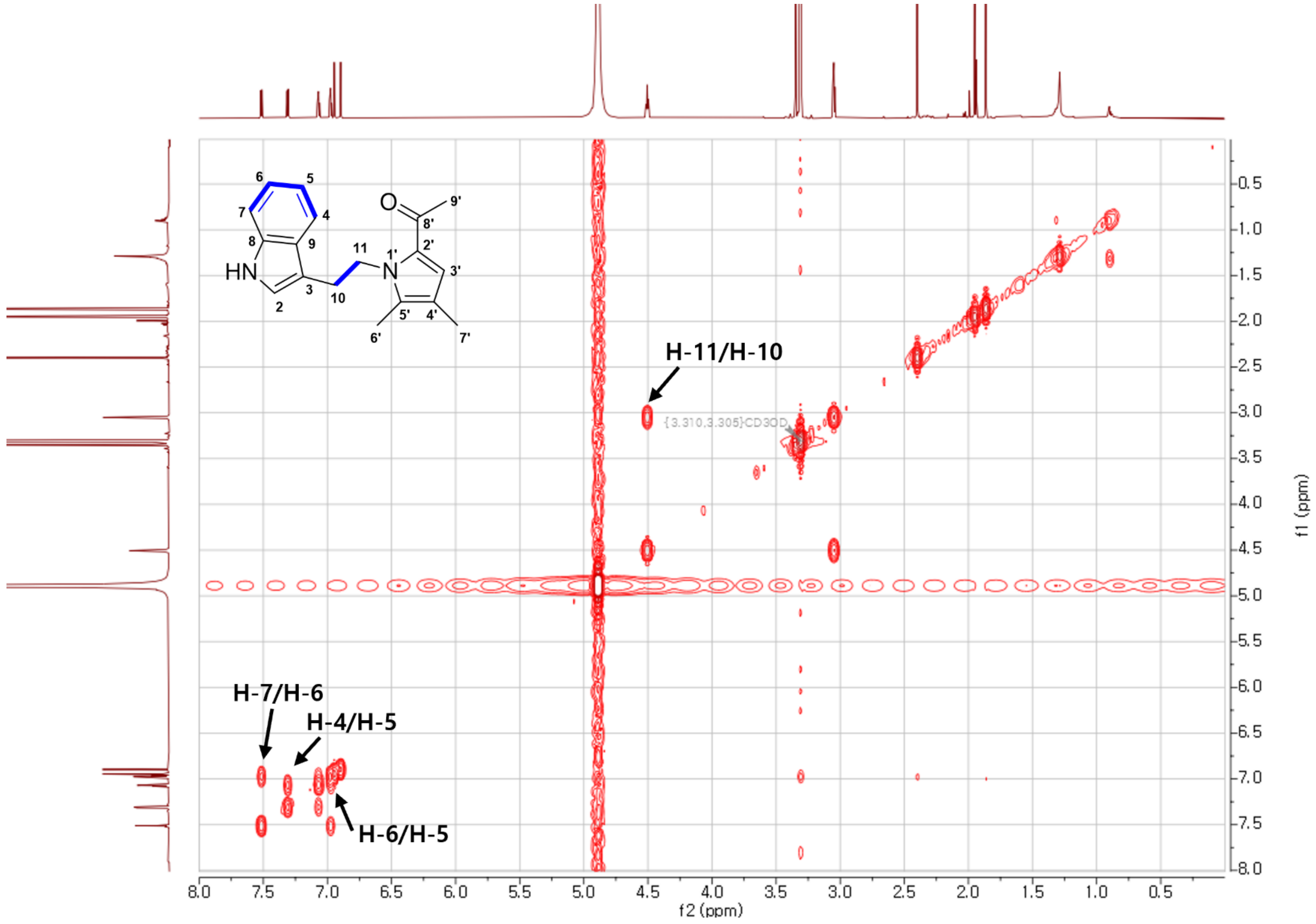

**Figure S77**. The COSY spectrum of **9** in methanol-$d_4$.

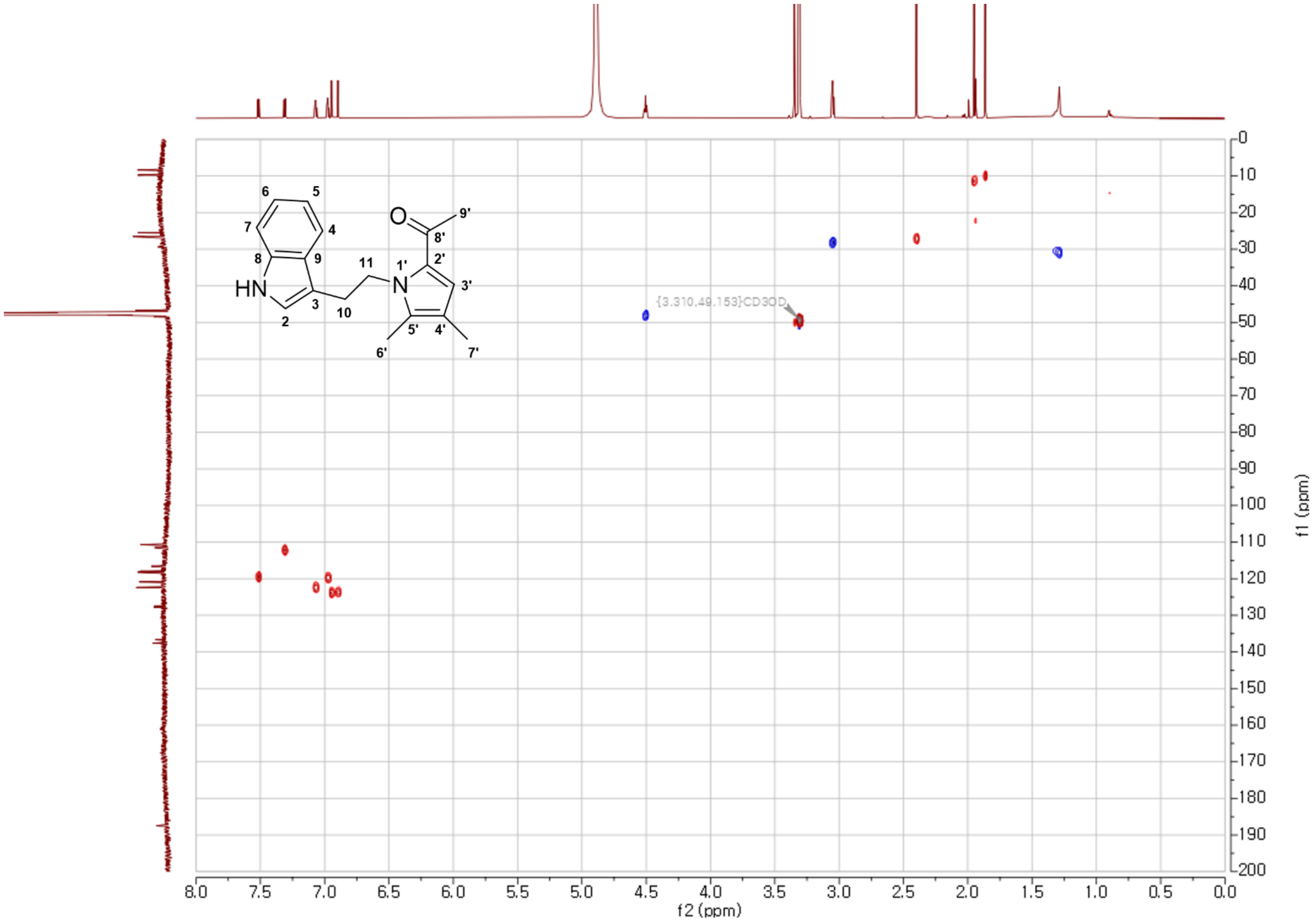

**Figure S78**. The HSQC spectrum of **12** in methanol-$d_4$.

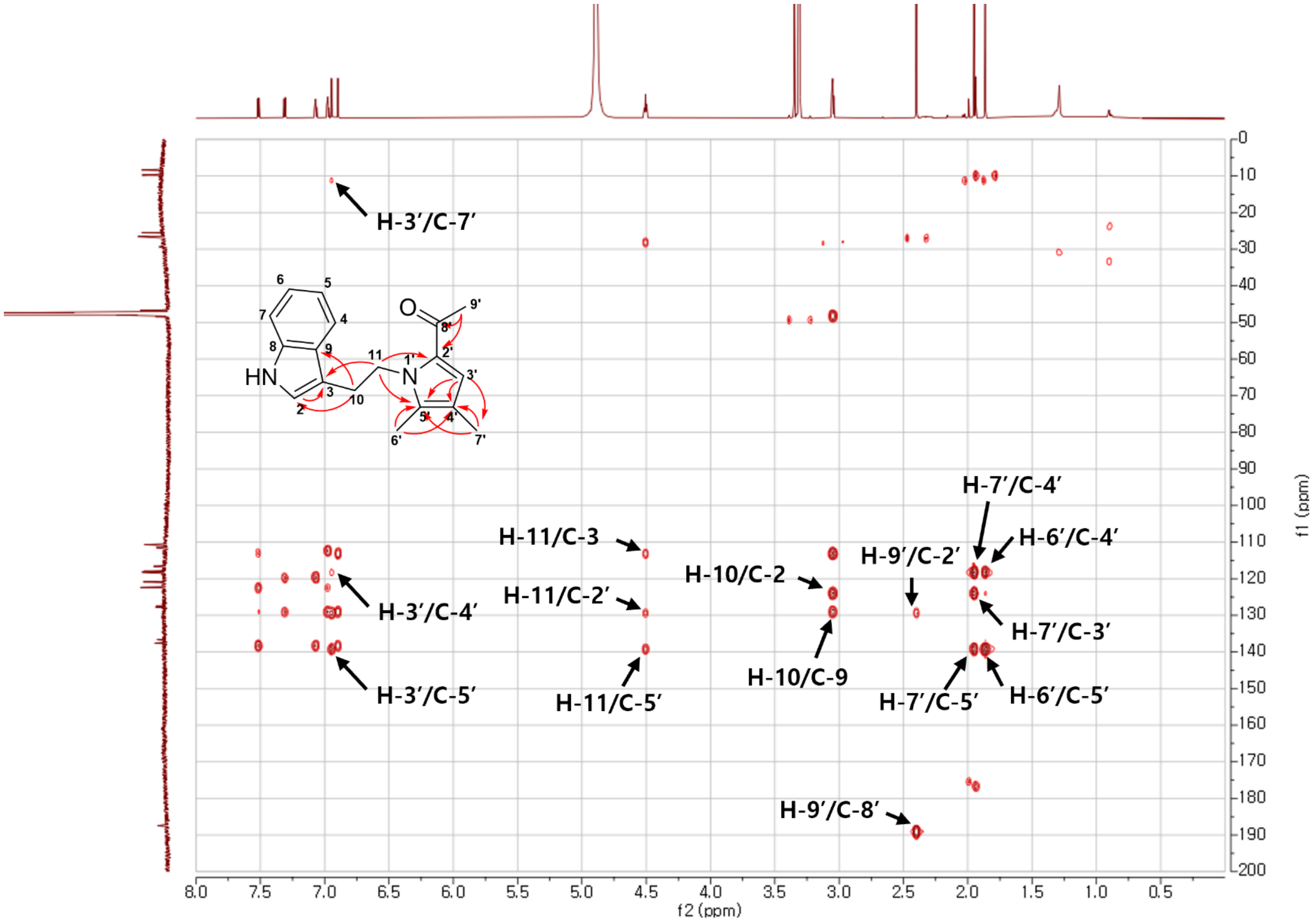

**Figure S79**. The HMBC spectrum of **12** in methanol-$d_4$.

### Coordinates of the conformers

### Conformer 9*R*-1

---------------------------------------------------------------------

| Center Number | Atomic Number | Atomic Type | Coordinates (Angstroms) X | Y | Z |
|---|---|---|---|---|---|
| 1 | 6 | 0 | 4.150759 | 1.822410 | 1.319085 |
| 2 | 6 | 0 | 3.540276 | 1.039628 | 0.171919 |
| 3 | 6 | 0 | 3.481068 | -0.499150 | 0.294780 |
| 4 | 6 | 0 | 4.827489 | -1.081329 | -0.177595 |
| 5 | 7 | 0 | 2.392551 | -1.088466 | -0.475150 |
| 6 | 6 | 0 | 1.061816 | -0.835178 | 0.087281 |
| 7 | 6 | 0 | -0.024023 | -1.453033 | -0.808991 |
| 8 | 6 | 0 | -1.416400 | -1.258434 | -0.280261 |
| 9 | 6 | 0 | -2.211579 | -2.212424 | 0.303130 |
| 10 | 7 | 0 | -3.419988 | -1.657968 | 0.687691 |
| 11 | 6 | 0 | -3.432202 | -0.320498 | 0.349239 |
| 12 | 6 | 0 | -4.423479 | 0.651603 | 0.520810 |
| 13 | 6 | 0 | -4.148145 | 1.936516 | 0.067295 |
| 14 | 6 | 0 | -2.915747 | 2.246565 | -0.543996 |
| 15 | 6 | 0 | -1.933991 | 1.277949 | -0.711151 |
| 16 | 6 | 0 | -2.180228 | -0.031978 | -0.262058 |

| | | | | | |
|---|---|---|---|---|---|
| 17 | 8 | 0 | 3.113612 | 1.589531 | -0.827657 |
| 18 | 1 | 0 | 3.337979 | -0.764331 | 1.351254 |
| 19 | 1 | 0 | 3.440245 | 1.840532 | 2.156591 |
| 20 | 1 | 0 | 5.067556 | 1.350121 | 1.689896 |
| 21 | 1 | 0 | 4.356045 | 2.848997 | 1.007854 |
| 22 | 1 | 0 | 5.671601 | -0.692289 | 0.402909 |
| 23 | 1 | 0 | 4.997660 | -0.833777 | -1.231743 |
| 24 | 1 | 0 | 4.804159 | -2.170083 | -0.080164 |
| 25 | 1 | 0 | 2.427679 | -0.665465 | -1.404502 |
| 26 | 1 | 0 | 1.015893 | -1.305021 | 1.078432 |
| 27 | 1 | 0 | 0.847747 | 0.237987 | 0.231888 |
| 28 | 1 | 0 | 0.192767 | -2.521386 | -0.925925 |
| 29 | 1 | 0 | 0.056846 | -1.009371 | -1.812747 |
| 30 | 1 | 0 | -2.011202 | -3.261689 | 0.475163 |
| 31 | 1 | 0 | -4.181822 | -2.160402 | 1.114864 |
| 32 | 1 | 0 | -5.374552 | 0.412463 | 0.990209 |
| 33 | 1 | 0 | -4.896954 | 2.715149 | 0.185190 |
| 34 | 1 | 0 | -2.733896 | 3.260942 | -0.888558 |
| 35 | 1 | 0 | -0.988278 | 1.530515 | -1.184052 |

---------------------------------------------------------------------

**Conformer 9*R*-2**

---------------------------------------------------------------------

| Center Number | Atomic Number | Atomic Type | Coordinates (Angstroms) X | Y | Z |
|---|---|---|---|---|---|
| 1 | 6 | 0 | 4.559350 | -0.002521 | 2.059195 |
| 2 | 6 | 0 | 3.951479 | 0.352107 | 0.715182 |
| 3 | 6 | 0 | 3.331330 | -0.787849 | -0.122403 |
| 4 | 6 | 0 | 4.450744 | -1.458081 | -0.942683 |
| 5 | 7 | 0 | 2.274946 | -0.328011 | -1.015369 |
| 6 | 6 | 0 | 1.035222 | 0.048913 | -0.328944 |
| 7 | 6 | 0 | -0.014346 | 0.534029 | -1.343273 |
| 8 | 6 | 0 | -1.345402 | 0.840058 | -0.718276 |
| 9 | 6 | 0 | -1.852770 | 2.081101 | -0.426227 |
| 10 | 7 | 0 | -3.093182 | 1.961160 | 0.175778 |
| 11 | 6 | 0 | -3.418092 | 0.624569 | 0.280575 |
| 12 | 6 | 0 | -4.557640 | 0.003657 | 0.802858 |
| 13 | 6 | 0 | -4.606226 | -1.385234 | 0.766678 |
| 14 | 6 | 0 | -3.546581 | -2.138799 | 0.220922 |
| 15 | 6 | 0 | -2.416831 | -1.517270 | -0.296296 |
| 16 | 6 | 0 | -2.334709 | -0.113620 | -0.272489 |
| 17 | 8 | 0 | 3.952717 | 1.493108 | 0.288811 |
| 18 | 1 | 0 | 2.903374 | -1.535393 | 0.559403 |
| 19 | 1 | 0 | 3.751022 | -0.201499 | 2.775820 |

| | | | | | |
|---|---|---|---|---|---|
| 20 | 1 | 0 | 5.164499 | -0.914484 | 2.001889 |
| 21 | 1 | 0 | 5.164840 | 0.827401 | 2.430037 |
| 22 | 1 | 0 | 5.249073 | -1.853874 | -0.304791 |
| 23 | 1 | 0 | 4.896741 | -0.735140 | -1.635448 |
| 24 | 1 | 0 | 4.027353 | -2.278630 | -1.528189 |
| 25 | 1 | 0 | 2.634609 | 0.498437 | -1.496356 |
| 26 | 1 | 0 | 0.653705 | -0.839088 | 0.191344 |
| 27 | 1 | 0 | 1.178459 | 0.831516 | 0.436787 |
| 28 | 1 | 0 | -0.119760 | -0.235591 | -2.118872 |
| 29 | 1 | 0 | 0.369426 | 1.431583 | -1.847430 |
| 30 | 1 | 0 | -1.422673 | 3.057954 | -0.603587 |
| 31 | 1 | 0 | -3.677321 | 2.729929 | 0.464120 |
| 32 | 1 | 0 | -5.376169 | 0.584217 | 1.221149 |
| 33 | 1 | 0 | -5.477537 | -1.898578 | 1.164331 |
| 34 | 1 | 0 | -3.618776 | -3.222982 | 0.205504 |
| 35 | 1 | 0 | -1.608119 | -2.109053 | -0.717963 |

---------------------------------------------------------------------

**Conformer 9*R*-3**

---------------------------------------------------------------------

| Center Number | Atomic Number | Atomic Type | Coordinates (Angstroms) X | Y | Z |
|---|---|---|---|---|---|
| 1 | 6 | 0 | 3.864331 | 0.589034 | 1.416817 |

| | | | | | |
|---|---|---|---|---|---|
| 2 | 6 | 0 | 3.926259 | 0.162570 | -0.038041 |
| 3 | 6 | 0 | 2.629781 | 0.225690 | -0.866714 |
| 4 | 6 | 0 | 2.497658 | 1.641403 | -1.452699 |
| 5 | 7 | 0 | 1.452273 | -0.097215 | -0.064033 |
| 6 | 6 | 0 | 1.260794 | -1.530200 | 0.158484 |
| 7 | 6 | 0 | 0.013031 | -1.815386 | 1.013786 |
| 8 | 6 | 0 | -1.290606 | -1.410506 | 0.379070 |
| 9 | 6 | 0 | -2.135130 | -2.226397 | -0.333277 |
| 10 | 7 | 0 | -3.232741 | -1.510726 | -0.775026 |
| 11 | 6 | 0 | -3.126352 | -0.205015 | -0.343776 |
| 12 | 6 | 0 | -3.986015 | 0.883866 | -0.524007 |
| 13 | 6 | 0 | -3.613030 | 2.097068 | 0.042427 |
| 14 | 6 | 0 | -2.411568 | 2.224236 | 0.769936 |
| 15 | 6 | 0 | -1.559380 | 1.141280 | 0.945516 |
| 16 | 6 | 0 | -1.908321 | -0.102614 | 0.385612 |
| 17 | 8 | 0 | 4.967787 | -0.179863 | -0.566416 |
| 18 | 1 | 0 | 2.780907 | -0.478954 | -1.703535 |
| 19 | 1 | 0 | 4.879599 | 0.679909 | 1.808690 |
| 20 | 1 | 0 | 3.323933 | 1.534956 | 1.529327 |
| 21 | 1 | 0 | 3.309049 | -0.150748 | 2.004324 |
| 22 | 1 | 0 | 2.351342 | 2.382965 | -0.659696 |
| 23 | 1 | 0 | 1.634824 | 1.692883 | -2.127447 |
| 24 | 1 | 0 | 3.392898 | 1.905045 | -2.023279 |
| 25 | 1 | 0 | 0.622092 | 0.276454 | -0.520582 |
| 26 | 1 | 0 | 1.189459 | -2.096576 | -0.789382 |

| | | | | | |
|---|---|---|---|---|---|
| 27 | 1 | 0 | 2.143483 | -1.916399 | 0.686009 |
| 28 | 1 | 0 | 0.131673 | -1.305927 | 1.978476 |
| 29 | 1 | 0 | -0.007159 | -2.891802 | 1.226208 |
| 30 | 1 | 0 | -2.043830 | -3.280915 | -0.557860 |
| 31 | 1 | 0 | -4.006475 | -1.893736 | -1.295030 |
| 32 | 1 | 0 | -4.912057 | 0.785881 | -1.084952 |
| 33 | 1 | 0 | -4.259029 | 2.962415 | -0.078078 |
| 34 | 1 | 0 | -2.150415 | 3.187881 | 1.198864 |
| 35 | 1 | 0 | -0.628033 | 1.251002 | 1.493880 |

---------------------------------------------------------------------

**Conformer 9*R*-4**

---------------------------------------------------------------------

| Center Number | Atomic Number | Atomic Type | Coordinates (Angstroms) X | Y | Z |
|---|---|---|---|---|---|
| 1 | 6 | 0 | -3.601363 | 2.289507 | -0.145507 |
| 2 | 6 | 0 | -3.475886 | 0.911234 | 0.476175 |
| 3 | 6 | 0 | -3.110984 | -0.271026 | -0.449128 |
| 4 | 6 | 0 | -4.399071 | -0.786992 | -1.120718 |
| 5 | 7 | 0 | -2.457307 | -1.360149 | 0.268673 |
| 6 | 6 | 0 | -1.046570 | -1.112210 | 0.600783 |
| 7 | 6 | 0 | -0.112810 | -1.483184 | -0.568447 |
| 8 | 6 | 0 | 1.340591 | -1.257190 | -0.261786 |

| | | | | | |
|---|---|---|---|---|---|
| 9 | 6 | 0 | 2.265101 | -2.214352 | 0.073846 |
| 10 | 7 | 0 | 3.495381 | -1.628354 | 0.315206 |
| 11 | 6 | 0 | 3.390884 | -0.265281 | 0.132759 |
| 12 | 6 | 0 | 4.352604 | 0.743936 | 0.248599 |
| 13 | 6 | 0 | 3.948746 | 2.049309 | -0.006776 |
| 14 | 6 | 0 | 2.618628 | 2.343468 | -0.371051 |
| 15 | 6 | 0 | 1.667662 | 1.337158 | -0.483592 |
| 16 | 6 | 0 | 2.041798 | 0.005561 | -0.230214 |
| 17 | 8 | 0 | -3.665855 | 0.722301 | 1.664086 |
| 18 | 1 | 0 | -2.435402 | 0.091953 | -1.234156 |
| 19 | 1 | 0 | -2.595277 | 2.681002 | -0.348943 |
| 20 | 1 | 0 | -4.130254 | 2.257129 | -1.104599 |
| 21 | 1 | 0 | -4.110323 | 2.968380 | 0.542268 |
| 22 | 1 | 0 | -4.911562 | -0.003911 | -1.690962 |
| 23 | 1 | 0 | -5.093791 | -1.166265 | -0.362563 |
| 24 | 1 | 0 | -4.148566 | -1.609131 | -1.796462 |
| 25 | 1 | 0 | -2.973368 | -1.478552 | 1.140296 |
| 26 | 1 | 0 | -0.853438 | -0.072280 | 0.917458 |
| 27 | 1 | 0 | -0.791684 | -1.744417 | 1.458326 |
| 28 | 1 | 0 | -0.395833 | -0.904427 | -1.459200 |
| 29 | 1 | 0 | -0.290516 | -2.535001 | -0.820886 |
| 30 | 1 | 0 | 2.143774 | -3.286207 | 0.157968 |

| | | | | | |
|---|---|---|---|---|---|
| 31 | 1 | 0 | 4.338304 | -2.123096 | 0.560370 |
| 32 | 1 | 0 | 5.378638 | 0.517389 | 0.527647 |
| 33 | 1 | 0 | 4.671356 | 2.856692 | 0.074905 |
| 34 | 1 | 0 | 2.336919 | 3.374895 | -0.565192 |
| 35 | 1 | 0 | 0.645377 | 1.577326 | -0.765763 |

---------------------------------------------------------------------

**Conformer 9*R*-5**

---------------------------------------------------------------------

| Center Number | Atomic Number | Atomic Type | Coordinates (Angstroms) X | Y | Z |
|---|---|---|---|---|---|
| 1 | 6 | 0 | -4.709122 | 1.587976 | 0.668038 |
| 2 | 6 | 0 | -3.974594 | 0.264137 | 0.772653 |
| 3 | 6 | 0 | -3.326308 | -0.307818 | -0.507804 |
| 4 | 6 | 0 | -4.402846 | -1.067474 | -1.308139 |
| 5 | 7 | 0 | -2.207131 | -1.197607 | -0.219098 |
| 6 | 6 | 0 | -0.968647 | -0.515606 | 0.182577 |
| 7 | 6 | 0 | -0.155730 | -0.034371 | -1.036245 |
| 8 | 6 | 0 | 1.166694 | 0.570746 | -0.659343 |
| 9 | 6 | 0 | 1.483710 | 1.905332 | -0.619778 |
| 10 | 7 | 0 | 2.789195 | 2.075196 | -0.192410 |
| 11 | 6 | 0 | 3.350085 | 0.838360 | 0.050113 |

| | | | | | |
|---|---|---|---|---|---|
| 12 | 6 | 0 | 4.634142 | 0.491359 | 0.483415 |
| 13 | 6 | 0 | 4.917365 | -0.861211 | 0.634260 |
| 14 | 6 | 0 | 3.946457 | -1.845806 | 0.357731 |
| 15 | 6 | 0 | 2.672636 | -1.495772 | -0.072054 |
| 16 | 6 | 0 | 2.352451 | -0.135808 | -0.232060 |
| 17 | 8 | 0 | -3.900331 | -0.347770 | 1.822768 |
| 18 | 1 | 0 | -2.966077 | 0.527039 | -1.122216 |
| 19 | 1 | 0 | -5.330042 | 1.638261 | -0.233454 |
| 20 | 1 | 0 | -5.323293 | 1.749188 | 1.556708 |
| 21 | 1 | 0 | -3.972759 | 2.399722 | 0.593817 |
| 22 | 1 | 0 | -5.254187 | -0.428353 | -1.568495 |
| 23 | 1 | 0 | -4.779461 | -1.914849 | -0.723831 |
| 24 | 1 | 0 | -3.959407 | -1.457170 | -2.228334 |
| 25 | 1 | 0 | -2.506222 | -1.794786 | 0.551535 |
| 26 | 1 | 0 | -1.142015 | 0.336257 | 0.863893 |
| 27 | 1 | 0 | -0.364868 | -1.236707 | 0.743806 |
| 28 | 1 | 0 | -0.742209 | 0.699210 | -1.605071 |
| 29 | 1 | 0 | -0.011504 | -0.894019 | -1.703066 |
| 30 | 1 | 0 | 0.874893 | 2.764269 | -0.869712 |
| 31 | 1 | 0 | 3.261793 | 2.960291 | -0.099861 |
| 32 | 1 | 0 | 5.383936 | 1.249877 | 0.694221 |
| 33 | 1 | 0 | 5.905398 | -1.164625 | 0.969704 |

| | | | | | |
|---|---|---|---|---|---|
| 34 | 1 | 0 | 4.202410 | -2.894310 | 0.483823 |
| 35 | 1 | 0 | 1.934051 | -2.264525 | -0.285235 |

---------------------------------------------------------------------

**Conformer 9*R*-6**

---------------------------------------------------------------------

| Center Number | Atomic Number | Atomic Type | Coordinates (Angstroms) X | Y | Z |
|---|---|---|---|---|---|
| 1 | 6 | 0 | -4.754990 | -0.189606 | -1.111316 |
| 2 | 6 | 0 | -3.270321 | 0.032838 | -0.883363 |
| 3 | 6 | 0 | -2.825299 | 0.544634 | 0.504284 |
| 4 | 6 | 0 | -2.921118 | 2.082484 | 0.511364 |
| 5 | 7 | 0 | -1.474139 | 0.138508 | 0.867326 |
| 6 | 6 | 0 | -1.321166 | -1.294712 | 1.127056 |
| 7 | 6 | 0 | 0.095797 | -1.626576 | 1.630921 |
| 8 | 6 | 0 | 1.198943 | -1.350175 | 0.645218 |
| 9 | 6 | 0 | 1.773032 | -2.258919 | -0.209148 |
| 10 | 7 | 0 | 2.754250 | -1.648490 | -0.968808 |
| 11 | 6 | 0 | 2.838782 | -0.318116 | -0.615645 |
| 12 | 6 | 0 | 3.678079 | 0.695816 | -1.089559 |
| 13 | 6 | 0 | 3.538616 | 1.959537 | -0.527912 |
| 14 | 6 | 0 | 2.584520 | 2.208653 | 0.480368 |

| | | | | | |
|---|---|---|---|---|---|
| 15 | 6 | 0 | 1.751175 | 1.199787 | 0.946936 |
| 16 | 6 | 0 | 1.868485 | -0.092061 | 0.400684 |
| 17 | 8 | 0 | -2.447809 | -0.172288 | -1.756945 |
| 18 | 1 | 0 | -3.516519 | 0.149164 | 1.261821 |
| 19 | 1 | 0 | -5.068221 | -1.097915 | -0.578922 |
| 20 | 1 | 0 | -5.354376 | 0.635912 | -0.710553 |
| 21 | 1 | 0 | -4.956825 | -0.316756 | -2.177145 |
| 22 | 1 | 0 | -3.934166 | 2.432679 | 0.283668 |
| 23 | 1 | 0 | -2.239002 | 2.506787 | -0.234150 |
| 24 | 1 | 0 | -2.631205 | 2.458295 | 1.496460 |
| 25 | 1 | 0 | -0.856694 | 0.389966 | 0.094438 |
| 26 | 1 | 0 | -2.041579 | -1.570131 | 1.909772 |
| 27 | 1 | 0 | -1.543021 | -1.918743 | 0.243562 |
| 28 | 1 | 0 | 0.112517 | -2.689942 | 1.903439 |
| 29 | 1 | 0 | 0.271822 | -1.060313 | 2.554173 |
| 30 | 1 | 0 | 1.562611 | -3.312944 | -0.333643 |
| 31 | 1 | 0 | 3.317482 | -2.105116 | -1.668583 |
| 32 | 1 | 0 | 4.412158 | 0.503849 | -1.868194 |
| 33 | 1 | 0 | 4.175256 | 2.769569 | -0.873588 |
| 34 | 1 | 0 | 2.500379 | 3.209472 | 0.895548 |
| 35 | 1 | 0 | 1.003863 | 1.402850 | 1.708507 |

---------------------------------------------------------------------

**Conformer 9*R*-7**

| Center Number | Atomic Number | Atomic Type | Coordinates (Angstroms) X | Y | Z |
|---|---|---|---|---|---|
| 1 | 6 | 0 | 4.333354 | 1.942226 | -0.703640 |
| 2 | 6 | 0 | 4.506320 | 0.566952 | -0.090502 |
| 3 | 6 | 0 | 3.289804 | -0.002863 | 0.669754 |
| 4 | 6 | 0 | 3.689473 | -1.153167 | 1.599018 |
| 5 | 7 | 0 | 2.292385 | -0.335104 | -0.360642 |
| 6 | 6 | 0 | 0.925541 | -0.538552 | 0.122949 |
| 7 | 6 | 0 | -0.027005 | -0.815041 | -1.053784 |
| 8 | 6 | 0 | -1.463163 | -0.944092 | -0.633818 |
| 9 | 6 | 0 | -2.183643 | -2.105130 | -0.506121 |
| 10 | 7 | 0 | -3.465125 | -1.826145 | -0.065754 |
| 11 | 6 | 0 | -3.601643 | -0.462781 | 0.095942 |
| 12 | 6 | 0 | -4.700362 | 0.298395 | 0.508624 |
| 13 | 6 | 0 | -4.539065 | 1.677666 | 0.571068 |
| 14 | 6 | 0 | -3.313851 | 2.285835 | 0.228204 |
| 15 | 6 | 0 | -2.225639 | 1.525284 | -0.180614 |
| 16 | 6 | 0 | -2.354480 | 0.126710 | -0.251838 |
| 17 | 8 | 0 | 5.532861 | -0.075164 | -0.209739 |

| | | | | | |
|---|---|---|---|---|---|
| 18 | 1 | 0 | 2.856321 | 0.817342 | 1.259845 |
| 19 | 1 | 0 | 4.321782 | 2.700100 | 0.091462 |
| 20 | 1 | 0 | 5.160117 | 2.155689 | -1.384413 |
| 21 | 1 | 0 | 3.375476 | 2.001108 | -1.228963 |
| 22 | 1 | 0 | 2.812305 | -1.568896 | 2.105272 |
| 23 | 1 | 0 | 4.395299 | -0.813300 | 2.364926 |
| 24 | 1 | 0 | 4.183983 | -1.947459 | 1.030923 |
| 25 | 1 | 0 | 2.605585 | -1.169911 | -0.856781 |
| 26 | 1 | 0 | 0.612615 | 0.378810 | 0.636907 |
| 27 | 1 | 0 | 0.830386 | -1.357414 | 0.857674 |
| 28 | 1 | 0 | 0.291135 | -1.737115 | -1.560160 |
| 29 | 1 | 0 | 0.091262 | -0.005104 | -1.785101 |
| 30 | 1 | 0 | -1.882128 | -3.125970 | -0.700375 |
| 31 | 1 | 0 | -4.191385 | -2.508708 | 0.082998 |
| 32 | 1 | 0 | -5.646078 | -0.169604 | 0.770265 |
| 33 | 1 | 0 | -5.373068 | 2.297951 | 0.887872 |
| 34 | 1 | 0 | -3.223469 | 3.367194 | 0.285151 |
| 35 | 1 | 0 | -1.287694 | 2.006670 | -0.446273 |

---------------------------------------------------------------------

**Conformer 9*R*-8**

---------------------------------------------------------------------

Center Atomic Atomic Coordinates (Angstroms)

| Number | Number | Type | X | Y | Z |
|---|---|---|---|---|---|
| 1 | 6 | 0 | 5.826691 | 0.423325 | -0.730272 |
| 2 | 6 | 0 | 4.604455 | -0.221627 | -0.104071 |
| 3 | 6 | 0 | 3.238300 | 0.438695 | -0.343939 |
| 4 | 6 | 0 | 3.272577 | 1.963588 | -0.132146 |
| 5 | 7 | 0 | 2.237790 | -0.185417 | 0.505398 |
| 6 | 6 | 0 | 0.911446 | -0.299849 | -0.089666 |
| 7 | 6 | 0 | -0.066076 | -0.971841 | 0.889878 |
| 8 | 6 | 0 | -1.469019 | -1.063779 | 0.361813 |
| 9 | 6 | 0 | -2.088656 | -2.180081 | -0.141011 |
| 10 | 7 | 0 | -3.370088 | -1.868073 | -0.560662 |
| 11 | 6 | 0 | -3.608926 | -0.529452 | -0.328067 |
| 12 | 6 | 0 | -4.747618 | 0.246516 | -0.569261 |
| 13 | 6 | 0 | -4.695136 | 1.590723 | -0.218692 |
| 14 | 6 | 0 | -3.537334 | 2.149309 | 0.361006 |
| 15 | 6 | 0 | -2.408888 | 1.374395 | 0.597947 |
| 16 | 6 | 0 | -2.427771 | 0.011415 | 0.252203 |
| 17 | 8 | 0 | 4.696074 | -1.247676 | 0.546305 |
| 18 | 1 | 0 | 3.036028 | 0.255503 | -1.422329 |
| 19 | 1 | 0 | 5.633044 | 0.743416 | -1.760160 |
| 20 | 1 | 0 | 6.109144 | 1.318180 | -0.161720 |

| | | | | | |
|---|---|---|---|---|---|
| 21 | 1 | 0 | 6.658226 | -0.283881 | -0.707540 |
| 22 | 1 | 0 | 3.954383 | 2.463301 | -0.828586 |
| 23 | 1 | 0 | 3.571093 | 2.198084 | 0.894823 |
| 24 | 1 | 0 | 2.272339 | 2.378579 | -0.290754 |
| 25 | 1 | 0 | 2.589791 | -1.101482 | 0.782109 |
| 26 | 1 | 0 | 0.542791 | 0.705497 | -0.329741 |
| 27 | 1 | 0 | 0.912288 | -0.863534 | -1.043610 |
| 28 | 1 | 0 | 0.305344 | -1.979007 | 1.124619 |
| 29 | 1 | 0 | -0.043939 | -0.408532 | 1.831636 |
| 30 | 1 | 0 | -1.714029 | -3.191027 | -0.232636 |
| 31 | 1 | 0 | -4.033285 | -2.522979 | -0.943575 |
| 32 | 1 | 0 | -5.641592 | -0.184308 | -1.013332 |
| 33 | 1 | 0 | -5.562914 | 2.221184 | -0.392746 |
| 34 | 1 | 0 | -3.531714 | 3.203096 | 0.626281 |
| 35 | 1 | 0 | -1.523769 | 1.815727 | 1.049346 |

---------------------------------------------------------------------

**Conformer 9*R*-9**

---------------------------------------------------------------------

| Center Number | Atomic Number | Atomic Type | Coordinates (Angstroms) X | Y | Z |
|---|---|---|---|---|---|
| 1 | 6 | 0 | 3.263174 | 1.839131 | -0.802964 |

| | | | | | |
|---|---|---|---|---|---|
| 2 | 6 | 0 | 3.604685 | 1.049716 | 0.446562 |
| 3 | 6 | 0 | 3.493493 | -0.485644 | 0.388286 |
| 4 | 6 | 0 | 4.840472 | -1.044939 | -0.099359 |
| 5 | 7 | 0 | 2.417202 | -0.924308 | -0.497610 |
| 6 | 6 | 0 | 1.081888 | -0.847266 | 0.098322 |
| 7 | 6 | 0 | 0.013868 | -1.352703 | -0.886166 |
| 8 | 6 | 0 | -1.387574 | -1.226547 | -0.361106 |
| 9 | 6 | 0 | -2.167567 | -2.230821 | 0.155117 |
| 10 | 7 | 0 | -3.387890 | -1.724689 | 0.565091 |
| 11 | 6 | 0 | -3.422978 | -0.368350 | 0.314609 |
| 12 | 6 | 0 | -4.432980 | 0.572341 | 0.542319 |
| 13 | 6 | 0 | -4.179459 | 1.888565 | 0.174369 |
| 14 | 6 | 0 | -2.950623 | 2.259903 | -0.409246 |
| 15 | 6 | 0 | -1.950951 | 1.321764 | -0.632753 |
| 16 | 6 | 0 | -2.173924 | -0.018269 | -0.269294 |
| 17 | 8 | 0 | 4.002954 | 1.587744 | 1.462639 |
| 18 | 1 | 0 | 3.345396 | -0.810464 | 1.433294 |
| 19 | 1 | 0 | 3.723486 | 1.394680 | -1.691564 |
| 20 | 1 | 0 | 2.181344 | 1.818921 | -0.977617 |
| 21 | 1 | 0 | 3.591472 | 2.872900 | -0.676097 |
| 22 | 1 | 0 | 5.655444 | -0.689062 | 0.537272 |
| 23 | 1 | 0 | 5.040344 | -0.742288 | -1.133266 |

| | | | | | |
|---|---|---|---|---|---|
| 24 | 1 | 0 | 4.832615 | -2.141241 | -0.061517 |
| 25 | 1 | 0 | 2.594568 | -1.886871 | -0.781921 |
| 26 | 1 | 0 | 1.003611 | -1.412828 | 1.045452 |
| 27 | 1 | 0 | 0.874930 | 0.199981 | 0.352302 |
| 28 | 1 | 0 | 0.222321 | -2.405310 | -1.125341 |
| 29 | 1 | 0 | 0.124587 | -0.796460 | -1.826117 |
| 30 | 1 | 0 | -1.948916 | -3.285097 | 0.262227 |
| 31 | 1 | 0 | -4.139366 | -2.266044 | 0.962163 |
| 32 | 1 | 0 | -5.381428 | 0.286251 | 0.990010 |
| 33 | 1 | 0 | -4.942748 | 2.644287 | 0.338366 |
| 34 | 1 | 0 | -2.786450 | 3.297489 | -0.686866 |
| 35 | 1 | 0 | -1.009175 | 1.620900 | -1.086433 |

---------------------------------------------------------------------

**Conformer 9*R*-10**

---------------------------------------------------------------------

| Center Number | Atomic Number | Atomic Type | Coordinates (Angstroms) X | Y | Z |
|---|---|---|---|---|---|
| 1 | 6 | 0 | 4.182631 | 1.602185 | 0.153917 |
| 2 | 6 | 0 | 4.005023 | 0.264915 | 0.848068 |
| 3 | 6 | 0 | 3.336213 | -0.877552 | 0.060486 |
| 4 | 6 | 0 | 4.431494 | -1.636715 | -0.706913 |

| | | | | | |
|---|---|---|---|---|---|
| 5 | 7 | 0 | 2.323428 | -0.380544 | -0.868540 |
| 6 | 6 | 0 | 1.040436 | -0.060700 | -0.238697 |
| 7 | 6 | 0 | 0.034033 | 0.453104 | -1.281581 |
| 8 | 6 | 0 | -1.305274 | 0.803358 | -0.698787 |
| 9 | 6 | 0 | -1.783750 | 2.065111 | -0.448436 |
| 10 | 7 | 0 | -3.045012 | 1.994422 | 0.114785 |
| 11 | 6 | 0 | -3.413401 | 0.670496 | 0.236324 |
| 12 | 6 | 0 | -4.587643 | 0.095916 | 0.734186 |
| 13 | 6 | 0 | -4.678607 | -1.291043 | 0.723841 |
| 14 | 6 | 0 | -3.626284 | -2.087888 | 0.227691 |
| 15 | 6 | 0 | -2.461945 | -1.512089 | -0.264473 |
| 16 | 6 | 0 | -2.336407 | -0.111395 | -0.266562 |
| 17 | 8 | 0 | 4.412763 | 0.063448 | 1.976654 |
| 18 | 1 | 0 | 2.918383 | -1.557290 | 0.823861 |
| 19 | 1 | 0 | 4.846547 | 2.231501 | 0.750411 |
| 20 | 1 | 0 | 4.580450 | 1.473728 | -0.858203 |
| 21 | 1 | 0 | 3.212631 | 2.099147 | 0.039197 |
| 22 | 1 | 0 | 4.885971 | -1.000904 | -1.474850 |
| 23 | 1 | 0 | 4.006210 | -2.517786 | -1.203016 |
| 24 | 1 | 0 | 5.212934 | -1.977175 | -0.021642 |
| 25 | 1 | 0 | 2.167751 | -1.079282 | -1.593972 |
| 26 | 1 | 0 | 0.609670 | -0.919772 | 0.307689 |

| | | | | | |
|---|---|---|---|---|---|
| 27 | 1 | 0 | 1.206497 | 0.723373 | 0.511019 |
| 28 | 1 | 0 | -0.089927 | -0.315783 | -2.059711 |
| 29 | 1 | 0 | 0.471168 | 1.325581 | -1.781491 |
| 30 | 1 | 0 | -1.319924 | 3.024963 | -0.633444 |
| 31 | 1 | 0 | -3.611822 | 2.785906 | 0.375113 |
| 32 | 1 | 0 | -5.399882 | 0.709945 | 1.115041 |
| 33 | 1 | 0 | -5.577195 | -1.769185 | 1.104032 |
| 34 | 1 | 0 | -3.731055 | -3.169415 | 0.232897 |
| 35 | 1 | 0 | -1.658544 | -2.138513 | -0.644259 |

---------------------------------------------------------------------

**Conformer 9*R*-11**

---------------------------------------------------------------------

| Center Number | Atomic Number | Atomic Type | Coordinates (Angstroms) X | Y | Z |
|---|---|---|---|---|---|
| 1 | 6 | 0 | -5.648145 | 0.995322 | -0.609675 |
| 2 | 6 | 0 | -4.379780 | 0.713528 | 0.173918 |
| 3 | 6 | 0 | -3.286651 | -0.127794 | -0.502279 |
| 4 | 6 | 0 | -3.848229 | -1.389772 | -1.183074 |
| 5 | 7 | 0 | -2.261686 | -0.469341 | 0.469477 |
| 6 | 6 | 0 | -0.904342 | -0.521034 | -0.061608 |
| 7 | 6 | 0 | 0.096868 | -0.879888 | 1.049553 |

| | | | | | |
|---|---|---|---|---|---|
| 8 | 6 | 0 | 1.518272 | -0.963063 | 0.571201 |
| 9 | 6 | 0 | 2.239145 | -2.108552 | 0.344623 |
| 10 | 7 | 0 | 3.505101 | -1.793892 | -0.117442 |
| 11 | 6 | 0 | 3.631630 | -0.422207 | -0.191651 |
| 12 | 6 | 0 | 4.714232 | 0.370780 | -0.586770 |
| 13 | 6 | 0 | 4.546846 | 1.750318 | -0.551517 |
| 14 | 6 | 0 | 3.330893 | 2.327445 | -0.130867 |
| 15 | 6 | 0 | 2.258499 | 1.535264 | 0.259332 |
| 16 | 6 | 0 | 2.393603 | 0.135689 | 0.233452 |
| 17 | 8 | 0 | -4.217696 | 1.162648 | 1.294600 |
| 18 | 1 | 0 | -2.891236 | 0.535777 | -1.302780 |
| 19 | 1 | 0 | -6.196454 | 1.806693 | -0.126842 |
| 20 | 1 | 0 | -5.431058 | 1.255955 | -1.651589 |
| 21 | 1 | 0 | -6.283604 | 0.101048 | -0.626010 |
| 22 | 1 | 0 | -4.555304 | -1.148455 | -1.983958 |
| 23 | 1 | 0 | -4.343586 | -2.030606 | -0.446347 |
| 24 | 1 | 0 | -3.028053 | -1.963876 | -1.625351 |
| 25 | 1 | 0 | -2.323987 | 0.197928 | 1.237948 |
| 26 | 1 | 0 | -0.854900 | -1.293462 | -0.839843 |
| 27 | 1 | 0 | -0.596299 | 0.425716 | -0.547138 |
| 28 | 1 | 0 | -0.211671 | -1.833974 | 1.492537 |
| 29 | 1 | 0 | 0.016472 | -0.129031 | 1.849938 |

| | | | | | |
|---|---|---|---|---|---|
| 30 | 1 | 0 | 1.947407 | -3.141445 | 0.481407 |
| 31 | 1 | 0 | 4.229737 | -2.460748 | -0.330911 |
| 32 | 1 | 0 | 5.652717 | -0.073788 | -0.908656 |
| 33 | 1 | 0 | 5.368526 | 2.394859 | -0.851867 |
| 34 | 1 | 0 | 3.234817 | 3.409694 | -0.112657 |
| 35 | 1 | 0 | 1.327100 | 1.992965 | 0.583247 |

---------------------------------------------------------------------

**Conformer 9*R*-12**

---------------------------------------------------------------------

| Center Number | Atomic Number | Atomic Type | Coordinates (Angstroms) X | Y | Z |
|---|---|---|---|---|---|
| 1 | 6 | 0 | -4.635797 | 1.713752 | 0.607121 |
| 2 | 6 | 0 | -3.586543 | 0.938942 | -0.172193 |
| 3 | 6 | 0 | -3.537778 | -0.598953 | -0.005967 |
| 4 | 6 | 0 | -3.652202 | -1.056711 | 1.456605 |
| 5 | 7 | 0 | -2.404796 | -1.212584 | -0.692202 |
| 6 | 6 | 0 | -1.095581 | -0.995270 | -0.060229 |
| 7 | 6 | 0 | 0.025078 | -1.427695 | -1.019755 |
| 8 | 6 | 0 | 1.401428 | -1.222958 | -0.455684 |
| 9 | 6 | 0 | 2.266199 | -2.194472 | -0.019280 |
| 10 | 7 | 0 | 3.434297 | -1.618580 | 0.450030 |

| | | | | | |
|---|---|---|---|---|---|
| 11 | 6 | 0 | 3.348014 | -0.247791 | 0.317611 |
| 12 | 6 | 0 | 4.267158 | 0.756786 | 0.638700 |
| 13 | 6 | 0 | 3.895818 | 2.071932 | 0.383285 |
| 14 | 6 | 0 | 2.639880 | 2.380352 | -0.179085 |
| 15 | 6 | 0 | 1.729521 | 1.379639 | -0.495443 |
| 16 | 6 | 0 | 2.074901 | 0.038585 | -0.248839 |
| 17 | 8 | 0 | -2.835886 | 1.500965 | -0.949766 |
| 18 | 1 | 0 | -4.448514 | -0.952011 | -0.520911 |
| 19 | 1 | 0 | -4.724259 | 2.721509 | 0.195700 |
| 20 | 1 | 0 | -5.610021 | 1.211376 | 0.580145 |
| 21 | 1 | 0 | -4.341208 | 1.786116 | 1.661689 |
| 22 | 1 | 0 | -2.859350 | -0.635813 | 2.083688 |
| 23 | 1 | 0 | -3.577121 | -2.147536 | 1.498635 |
| 24 | 1 | 0 | -4.614104 | -0.763738 | 1.887662 |
| 25 | 1 | 0 | -2.362338 | -0.788739 | -1.619094 |
| 26 | 1 | 0 | -0.929586 | 0.051669 | 0.240129 |
| 27 | 1 | 0 | -1.032537 | -1.605173 | 0.848492 |
| 28 | 1 | 0 | -0.077823 | -0.855576 | -1.954416 |
| 29 | 1 | 0 | -0.126843 | -2.481434 | -1.282941 |
| 30 | 1 | 0 | 2.142846 | -3.269351 | -0.008191 |
| 31 | 1 | 0 | 4.231950 | -2.123386 | 0.802407 |
| 32 | 1 | 0 | 5.236184 | 0.519604 | 1.070952 |

| | | | | | |
|---|---|---|---|---|---|
| 33 | 1 | 0 | 4.586849 | 2.876283 | 0.620925 |
| 34 | 1 | 0 | 2.382440 | 3.419342 | -0.366223 |
| 35 | 1 | 0 | 0.763432 | 1.630457 | -0.926395 |

---------------------------------------------------------------------

**Conformer 9*R*-13**

---------------------------------------------------------------------

| Center Number | Atomic Number | Atomic Type | Coordinates (Angstroms) X | Y | Z |
|---|---|---|---|---|---|
| 1 | 6 | 0 | 4.525743 | -0.696483 | -0.755410 |
| 2 | 6 | 0 | 4.091177 | 0.490085 | 0.084713 |
| 3 | 6 | 0 | 2.730667 | 0.409690 | 0.801715 |
| 4 | 6 | 0 | 2.945936 | -0.255866 | 2.171376 |
| 5 | 7 | 0 | 1.747859 | -0.342709 | 0.026755 |
| 6 | 6 | 0 | 1.133710 | 0.419020 | -1.059613 |
| 7 | 6 | 0 | 0.050129 | -0.413717 | -1.768710 |
| 8 | 6 | 0 | -1.113113 | -0.774651 | -0.886605 |
| 9 | 6 | 0 | -1.362167 | -1.999526 | -0.316044 |
| 10 | 7 | 0 | -2.517769 | -1.946694 | 0.442521 |
| 11 | 6 | 0 | -3.047340 | -0.674144 | 0.373955 |
| 12 | 6 | 0 | -4.198944 | -0.136788 | 0.958838 |
| 13 | 6 | 0 | -4.485712 | 1.198803 | 0.701980 |

| | | | | | |
|---|---|---|---|---|---|
| 14 | 6 | 0 | -3.647546 | 1.981123 | -0.118968 |
| 15 | 6 | 0 | -2.504578 | 1.442956 | -0.696462 |
| 16 | 6 | 0 | -2.184136 | 0.095030 | -0.454856 |
| 17 | 8 | 0 | 4.800508 | 1.467224 | 0.236868 |
| 18 | 1 | 0 | 2.426391 | 1.457203 | 0.974502 |
| 19 | 1 | 0 | 4.368628 | -1.639139 | -0.220580 |
| 20 | 1 | 0 | 3.918062 | -0.757556 | -1.665327 |
| 21 | 1 | 0 | 5.577405 | -0.580127 | -1.025829 |
| 22 | 1 | 0 | 3.709252 | 0.280355 | 2.742506 |
| 23 | 1 | 0 | 3.257955 | -1.299935 | 2.056417 |
| 24 | 1 | 0 | 2.014077 | -0.240919 | 2.749496 |
| 25 | 1 | 0 | 1.004876 | -0.665442 | 0.645287 |
| 26 | 1 | 0 | 0.698916 | 1.373516 | -0.708559 |
| 27 | 1 | 0 | 1.912753 | 0.680824 | -1.787980 |
| 28 | 1 | 0 | 0.517178 | -1.327817 | -2.154204 |
| 29 | 1 | 0 | -0.304691 | 0.154393 | -2.639094 |
| 30 | 1 | 0 | -0.803866 | -2.922165 | -0.402844 |
| 31 | 1 | 0 | -2.924114 | -2.723599 | 0.939469 |
| 32 | 1 | 0 | -4.846983 | -0.740052 | 1.589614 |
| 33 | 1 | 0 | -5.372906 | 1.647184 | 1.140658 |
| 34 | 1 | 0 | -3.902917 | 3.021494 | -0.300517 |
| 35 | 1 | 0 | -1.867654 | 2.056623 | -1.328339 |

-----------------------------------------------------------------

**Conformer 9*R*-14**

-----------------------------------------------------------------

| Center Number | Atomic Number | Atomic Type | Coordinates (Angstroms) X | Y | Z |
|---|---|---|---|---|---|
| 1 | 6 | 0 | -5.272026 | 0.025969 | -1.288663 |
| 2 | 6 | 0 | -4.147275 | -0.354398 | -0.340449 |
| 3 | 6 | 0 | -3.431074 | 0.774827 | 0.437727 |
| 4 | 6 | 0 | -3.102406 | 1.999760 | -0.430034 |
| 5 | 7 | 0 | -2.276069 | 0.304406 | 1.197097 |
| 6 | 6 | 0 | -1.082652 | -0.016682 | 0.402449 |
| 7 | 6 | 0 | -0.030144 | -0.697197 | 1.294039 |
| 8 | 6 | 0 | 1.271306 | -0.948546 | 0.587942 |
| 9 | 6 | 0 | 1.713448 | -2.142487 | 0.076265 |
| 10 | 7 | 0 | 2.945031 | -1.975057 | -0.532683 |
| 11 | 6 | 0 | 3.329858 | -0.655104 | -0.420312 |
| 12 | 6 | 0 | 4.485642 | -0.003462 | -0.863476 |
| 13 | 6 | 0 | 4.599762 | 1.355905 | -0.595389 |
| 14 | 6 | 0 | 3.588377 | 2.050281 | 0.099952 |
| 15 | 6 | 0 | 2.442206 | 1.398484 | 0.537757 |
| 16 | 6 | 0 | 2.294319 | 0.023933 | 0.280522 |

| | | | | | |
|---|---|---|---|---|---|
| 17 | 8 | 0 | -3.849210 | -1.520091 | -0.150789 |
| 18 | 1 | 0 | -4.181645 | 1.103909 | 1.177973 |
| 19 | 1 | 0 | -4.858570 | 0.471656 | -2.202141 |
| 20 | 1 | 0 | -5.839618 | -0.867635 | -1.556984 |
| 21 | 1 | 0 | -5.939170 | 0.772627 | -0.841768 |
| 22 | 1 | 0 | -2.566135 | 2.736500 | 0.175696 |
| 23 | 1 | 0 | -4.014594 | 2.469662 | -0.809498 |
| 24 | 1 | 0 | -2.473851 | 1.741876 | -1.288707 |
| 25 | 1 | 0 | -2.565267 | -0.554867 | 1.664777 |
| 26 | 1 | 0 | -0.662397 | 0.914249 | 0.005700 |
| 27 | 1 | 0 | -1.294298 | -0.670903 | -0.459540 |
| 28 | 1 | 0 | -0.438987 | -1.649173 | 1.659840 |
| 29 | 1 | 0 | 0.128111 | -0.065972 | 2.178459 |
| 30 | 1 | 0 | 1.241326 | -3.115761 | 0.098289 |
| 31 | 1 | 0 | 3.485557 | -2.707690 | -0.964441 |
| 32 | 1 | 0 | 5.266982 | -0.538817 | -1.397276 |
| 33 | 1 | 0 | 5.485324 | 1.891850 | -0.926158 |
| 34 | 1 | 0 | 3.711287 | 3.112186 | 0.295350 |
| 35 | 1 | 0 | 1.670834 | 1.943895 | 1.075852 |

---------------------------------------------------------------------
